# Juvenile and adult expression of polyglutamine expanded *huntingtin* produce distinct aggregate distributions in *Drosophila* muscle

**DOI:** 10.1101/2023.04.04.535619

**Authors:** Taylor Barwell, Sehaj Raina, Austin Page, Hayley MacCharles, Laurent Seroude

**Affiliations:** Department of Biology, Queen’s University, Kingston, Ontario, Canada; Department of Biology, York University, Toronto, Ontario, Canada

## Abstract

While Huntington’s disease (HD) is widely recognized as a disease affecting the nervous system, much evidence has accumulated to suggest peripheral or non-neuronal tissues are affected as well. Here, we utilize the UAS/GAL4 system to express a pathogenic HD construct in the muscle of the fly, and characterize the effects. We observe detrimental phenotypes such as reduced lifespan, decreased locomotion, and accumulation of protein aggregates. Strikingly, depending on the GAL4 driver used to express the construct we saw different aggregate distributions and severity of phenotypes. These different aggregate distributions were found to be dependent on expression level and the timing of expression. Hsp70, a well-documented suppressor of polyglutamine aggregates, was found to strongly reduce the accumulation of aggregates in the eye, but in the muscle it did not prevent the reduction of the lifespan. Therefore, the molecular mechanisms underlying the detrimental effects of aggregates in the muscle are distinct from the nervous system.

## INTRODUCTION

The polyglutamine disease family encompasses at least nine hereditary disorders all defined by the abnormal expansion of repeated glutamine tracts (polyQ) in their respective proteins. The accumulation of these mutant proteins leads to formation of aggregates and impaired autophagy, axonal transport, endocytosis and proteostasis ^1–3^. These impairments manifest as motor coordination defects, muscle atrophy, neuropsychiatric symptoms and cognitive decline. The number of glutamine repeats in the polyQ tract is correlated with the severity of the disease and age of onset. In Huntington’s disease (HD), the most thoroughly studied polyQ disease, more than 35 glutamine repeats are necessary for disease symptoms.

*Drosophila melanogaster* have been used extensively to model polyQ diseases and led to numerous discoveries that have profoundly changed our understanding of many diseases ^4–6^. Various different constructs have been used to introduce polyQ repeats into the fly including N-terminal fragments of huntingtin ^7, 8^, C-terminal fragments of ataxin-3 ^9^, naked polyQ tracts ^10^, epitope-tagged polyQ tracts ^10, 11^, and the full-length ataxin-1 gene^12^.

There is much evidence to suggest peripheral tissues are involved in the pathology of HD. In mammals, the *huntingtin* (Htt) gene is expressed almost ubiquitously and polyQ inclusions have been observed in a wide range of tissue outside of the nervous system ^13–20^. In *C. elegans*, the expression of huntingtin constructs with polyQ expansion in the muscle cells results in delayed development, motor impairment, and shortened lifespan^21^. In *Drosophila*, driving a polyQ expanded ataxin-3 construct in the muscle causes larval lethality, whereas animals that express the same construct in the nervous system make it to the adult stage ^9^. Altering the expression of muscle-specific genes (*myostatin* signalling and *Scn4a* sodium channel) in HD mouse models has been shown to affect skeletal muscle atrophy ^22, 23^. Finally, clinical data have shown that myopathy markers are observed and muscle performance declined many years before first signs of chorea were diagnosed ^24^.

In this study, we further characterize the biological effects of overexpressing polyglutamine repeats in the muscle by taking advantage of the vast genetic toolkit of *Drosophila*. The majority of the *Drosophila* polyQ constructs were built under the control of a UAS promoter, since a UAS transgene is not expressed unless bound by the yeast transcription factor, GAL4 ^25^, it is trivial by a simple genetic cross to test such constructs in a variety of tissues. We chose three muscle-specific GAL4 drivers: DJ694, MHC-Geneswitch, and Mef2, which have been previously characterized across age ^26–28^. The drivers were each crossed with two UAS lines each containing a transgene with an N-terminal fragment of human *huntingtin* with either 72 (Q72: pathogenic) or 25 (Q25: non-pathogenic) glutamine repeats that is tagged with eGFP (UAS-Httex1-Qn-eGFP) ^29^. Three different phenotypes were examined to assess the biological effects of the pathogenic transgene in muscles. The longevity and locomotion behaviour were measured, as well as the aggregation and cellular distribution of the *huntingtin* encoded by the transgenes.

We show here that expression of the pathogenic transgene in the muscle causes detrimental effects, in the absence of any expression in the nervous system. Our findings further support that the muscle is a peripheral tissue that contributes to symptoms of the disease. We discover that the aggregates can arrange in distinct distributions depending on the time and level of expression. One of these distributions is associated with more severe symptoms and can only be established during the pre-adult stage. Finally, we demonstrate that the molecular mechanisms at play in the muscle differ from those that have been previously demonstrated in the nervous system.

## RESULTS

### Polyglutamine repeats in the muscle are detrimental and aggregates can have alternate distributions

We expressed Htt transgenes in the muscles using the UAS-GAL4 system ^25^ using three different muscle-specific GAL4 drivers: MHC Geneswitch, DJ694, and Mef2-Gal4 ^28, 30, 31^. MHC and Mef2 are expressed in the muscle at all stages ^26, 32, 33^. DJ694 is expressed in the adult muscle and in the oenocytes during metamorphosis ^28, 32^. We measured the longevity and locomotor behaviour of animals expressing Htt containing normal (Q25) or expanded (Q72) glutamine repeats, as well as animals that do not express Htt to control for the effect of the genetic background (driver, Q72, Q25 alone).

The effect on longevity of females and males can be seen in Figure 1A and S1A, respectively. With Mef2, the expression of the Q72 transgene strongly reduces longevity across all replicates in both sexes when compared to all controls. With MHC, the lifespan is consistently reduced across replicates in females but not males compared to the Q25 control. However, compared to the other controls no consistent significant effect is observed across replicates. With DJ694, no consistent effect is seen compared to either control in both sexes.

**Figure 1.**
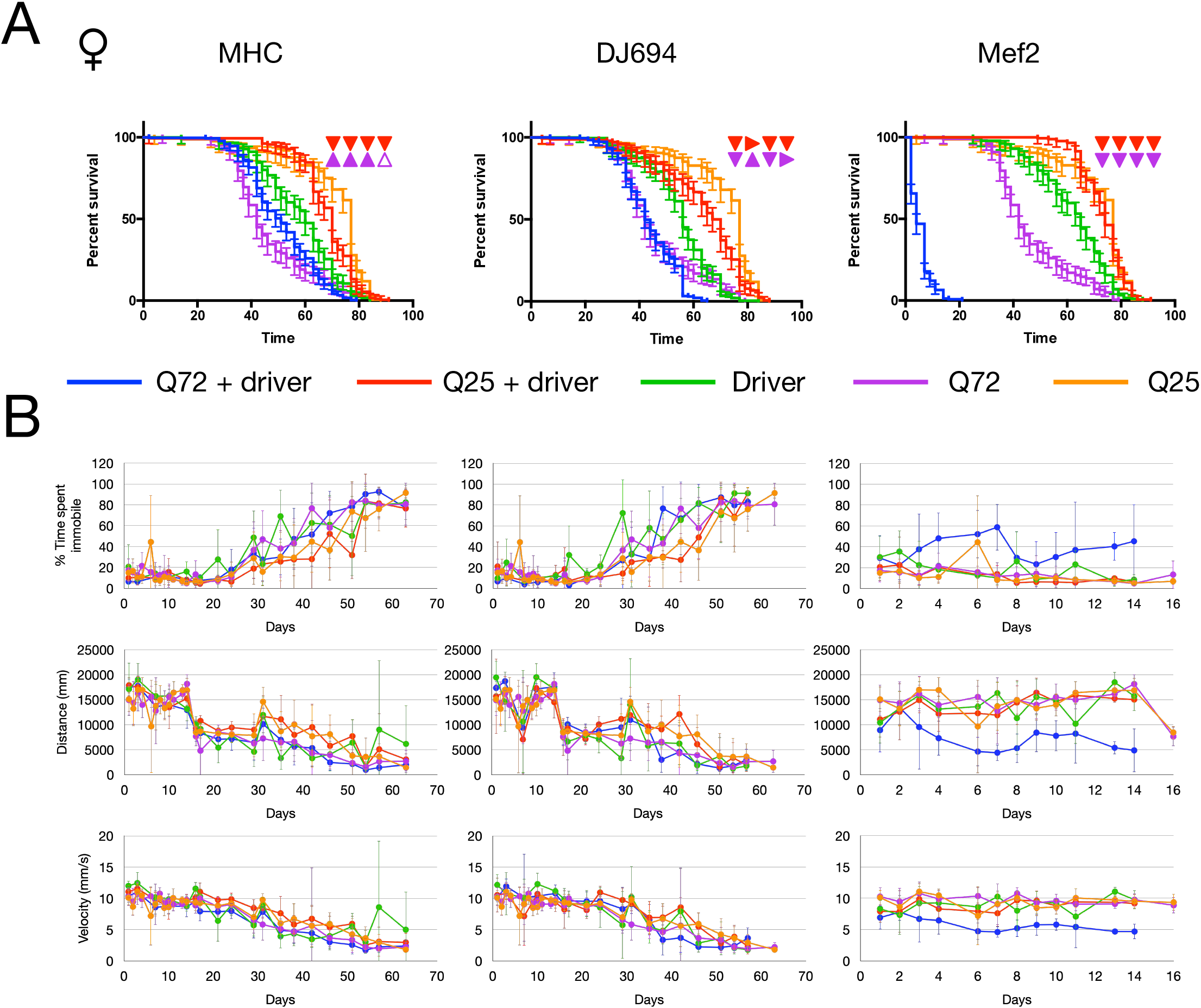
Expression of UAS-Htt-Q72 in the muscle results in declines in longevity and locomotion. All data from female flies. Three muscle specific GAL4 drivers are used: MHC-Geneswitch, DJ694, Mef2. A) Lifespan assay. X-axis denotes time in days at 25℃, y-axis denotes percent survival, error bars represent 95% CI. Triangles represent the result of the logrank comparison of Q72+driver curves to controls from the four independent replicates. Red triangles compare to Q25+driver. Violet triangles compare to Q72 alone. ▴: significant increase,△: insignificant increase,▾: significant decrease, ▽: insigificant decrease, ▶: no change B) Locomotion data measured by three parameters: % time spent immobile (<1mm/s), distance travelled (mm), velocity (mm/s). Error bars represent ± SD. Genotype abbreviations: *Q72+driver (blue)*: Q72/+;;MHC/+, Q72/+;DJ694/+, Q72/+;;Mef2/+ ; *Q25+driver (red)*: Q25/MHC, DJ694/+;Q25/+, Q25/Mef2 ; *Driver (green):* MHC/+, DJ694/+, Mef2/+ ; *Q72 (violet):* Q72/+ ; *Q25 (orange):* Q25/+

The effect on locomotion is seen in Figure 1B and S1B. With the MHC and DJ694 drivers, there is no apparent difference in locomotion in any of the parameters measured (% time spent immobile, distance travelled, velocity). With Mef2-Gal4, in both sexes it is evident they are more immobile, and display a slower velocity that cannot be attributed to exhaustion because they also covered less distance.

Next, Q25 and Q72 expression was visualized in the muscle tissue to assess protein aggregation. In order to exclude artifacts from the fixation ^34^, the GFP intensity and pattern visualized by immunochemistry was systematically confirmed by direct excitation and gave identical results. As expected, no aggregates are present with Q25 resulting in a diffused signal in the muscle with MHC, DJ694, and Mef2 in both sexes (Figure 2 and S2). With the Q72 expressing flies, eGFP-positive puncta (aggregates) are formed with all three GAL4 drivers in both males and females. Interestingly, the pattern or distribution of the aggregates differed depending on the driver. With MHC and DJ694, the aggregates appear randomly dispersed throughout the muscle. In contrast, in the Mef2 genotypes the aggregates have a distinct striped arrangement, which we will hereafter refer to as striated. Notably, the striated distribution correlates with severe longevity and locomotor phenotypes which lead us to investigate the cause of the alternate distribution.

**Figure 2.**
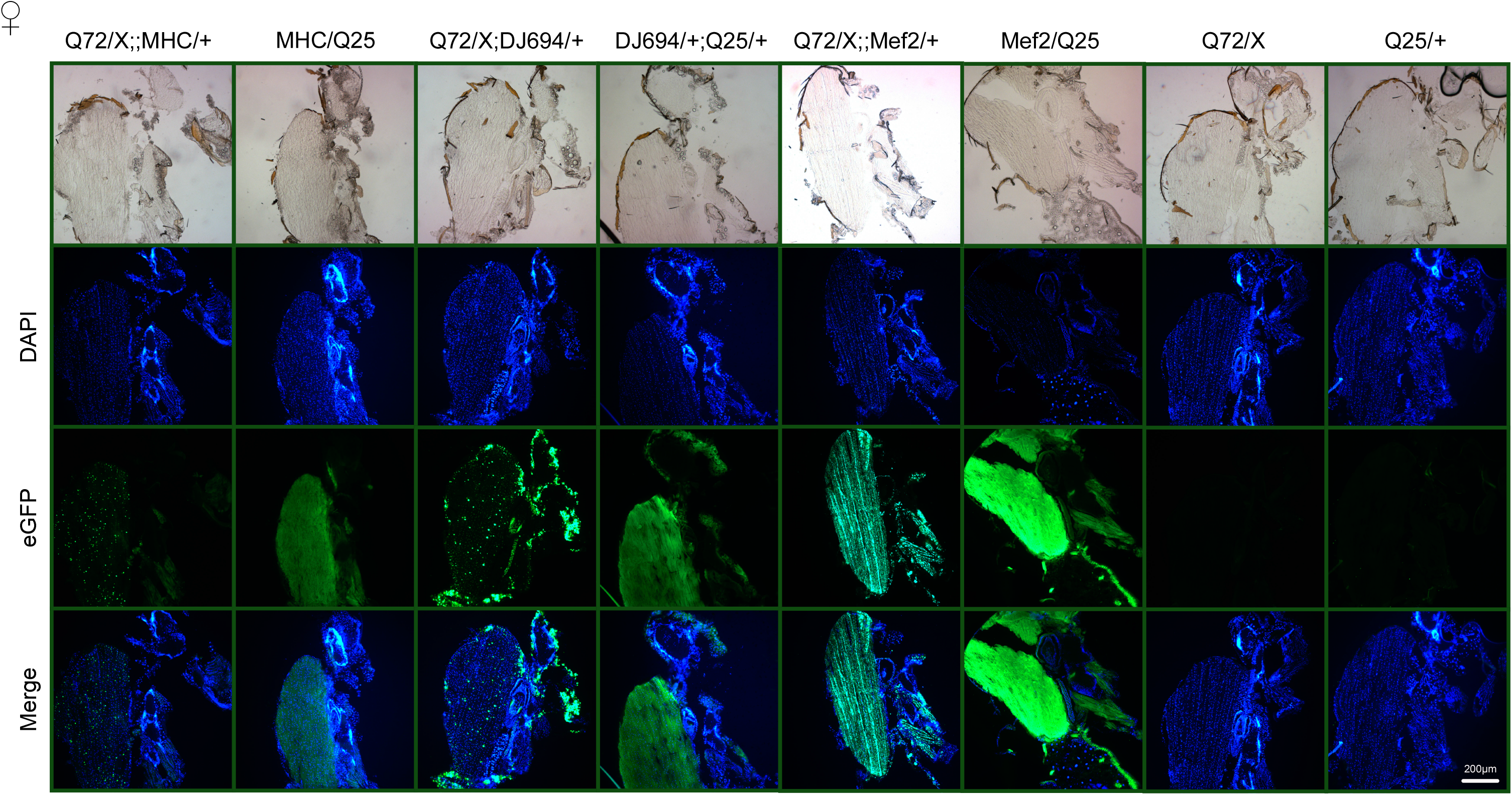
Alternate aggregate distributions with different muscle drivers. Visualization by saggital cryosectioning and anti-eGFP staining. Blue is DAPI. Green is eGFP. All photos taken under identical light exposure parameters. All images are of female flies. Genotypes are indicated. Scale bar represents 200µm.

### Increasing expression level alone is insufficient to phenocopy striated aggregate distribution

The expression profile of each driver was measured to compare expression levels during development and adulthood. It was found that from the late pupal stage and thereafter Mef2 had increasingly higher levels of ß-galactosidase activity in males and females (Figure S3). This led to the first hypothesis that the striated aggregate distribution observed with Mef2, and not with the other two drivers, is due to a higher expression level in adult muscles.

In order to test this, the expression level was increased with MHC and DJ694 to see whether the striation and pathogenic effects could be phenocopied. The expression level was increased in two ways. First, strains were built that were homozygous for both UAS and GAL4 transgenes, allowing for the measurement of genotypes with up to two copies of each transgene. Second, the MHC driver transgene expresses a modified GAL4 protein (Geneswitch) that is a fusion of the GAL4 DNA binding domain, ligand binding domain of human progesterone receptor, and the activation domain of human p65 which allows it to be inducible by feeding antiprogestin, RU486 ^30, 35^. Even in the absence of the inducer there is still some expression, which is why aggregates were visible in the MHC flies without the need to induce ^27^. However, by treating the MHC flies with RU486 we can increase the expression further.

In order to assess whether expression level had been increased to the level of Mef2, a UAS-lacZ reporter line was used to quantify the expression level of DJ694 and MHC genotypes after increasing copy number and RU486 induction (Figure 3A). With MHC, when the transgene copy number is increased and treated with RU486, the level of expression is not significantly different in females and significantly higher in males (p<0.05) to that of Mef2. Likewise, with DJ694 when the number of copies was increased the expression level is not significantly different to that of Mef2 in both males and females. Increasing the expression level aggravated the longevity with both drivers (Figure 3 and S4). With DJ694, the genotypes with two copies of the GAL4 transgene displayed the greatest reduction in lifespan in both males and females (Figure 3B and S4A). In line with this result, increasing the number of copies of GAL4 results in the greatest increase in expression level as measured by UAS-lacZ reporter (Figure S5). Indeed, these genotypes also display more protein aggregates, that tend to also be larger in size. With MHC, increasing the copy number did not result in a strong reduction in longevity, however the genotypes with increased copies of the UAS transgene tended to be shorter lived, in both males and females (Figure 3C and S4B). Consistent with this observation, increasing the number of copies of UAS resulted in the greatest increase in UAS-lacZ expression level (Figure S5) and the most abundant aggregates (Figure 3C). Induction with RU486 resulted in a much shorter lifespan (Figure 3D and S4C), as well as a greater accumulation of aggregates that appear larger in size. Still, neither the longevity of DJ694 nor RU486 treated MHC was as severe as Mef2, and the aggregates remain in the dispersed distribution. Taken together, increasing the expression level equivalent to Mef2 does not recapitulate the striated aggregate distribution, leading to the conclusion that high expression in adult muscle is insufficient for striation. However, increasing the expression level increases the amount of dispersed aggregates and aggravates the longevity, demonstrating that the presence of dispersed aggregates in adult muscle is detrimental.

**Figure 3.**
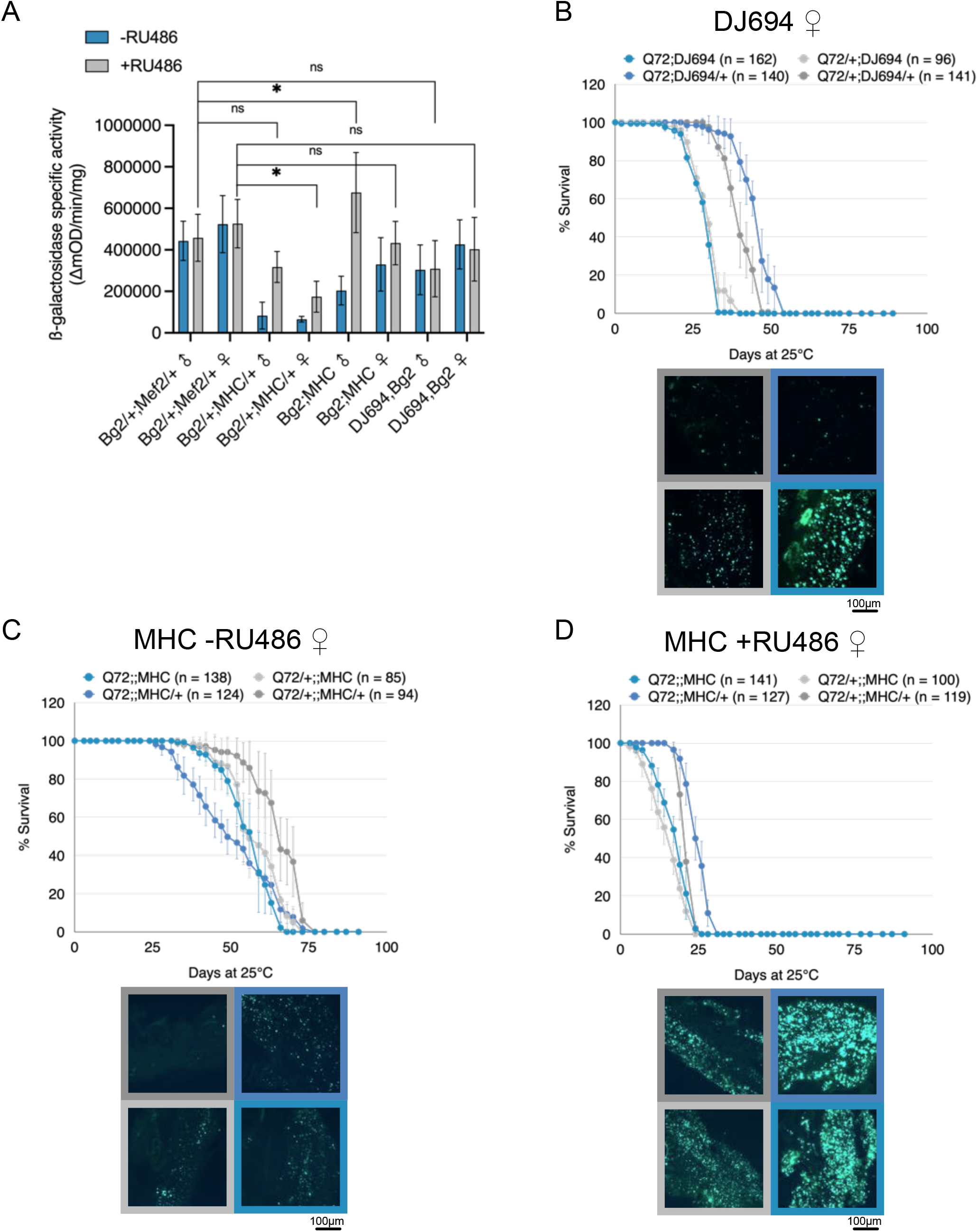
Increased expression level aggravates longevity and amount of aggregates. Expression levels were increased by increasing copy number and treating with RU486 (50µg/ml). A) UAS-lacZ (Bg2) reporter assay to quantify expression level compared to Mef2. Flies measured were 10-12 days old (n=5, 2 independent replicates). Y-axis represents ß-galactosidase specific activity (ΔmOD/min/mg). Error bars specify ± SD. One-way ANOVA was used to compare relative to Mef2 genotype. ns= not significant, *p<0.05. B-D) Effect of increasing expression on longevity and aggregates. Graphs depict representative survival data from one female replicate (n= number of flies in the replicate). Flies were maintained at 25℃. Error bars represent ± SD. Representative images of muscle aggregates are displayed below the graphs. Panels of aggregate images are outlined in a colour corresponding to the legend in the survival graph. Increasing copy number with B) DJ694 and C) MHC. D) Increasing copy number and RU486 (50µg/ml) induction with MHC.

### Striated aggregate distribution is established during metamorphosis

Since Mef2 is also significantly higher in the late pupae we hypothesized that the combination of level and timing of expression is responsible for the striated distribution. Because this hypothesis implies that the striated distribution is established in the late pupal stages, L3 larval fillet dissections were performed and staged early pupae were collected and imaged daily (Figure 4A). With MHC, no aggregates can be detected at any stage, as expected from the low level of expression. With Mef2, in the L3 larvae aggregates are visible all throughout the muscles, and they do not appear to be striated but rather dispersed. The aggregates remain dispersed into the pupae until two days after puparium formation (APF), where in the thorax the aggregates have clearly arranged within the dorsal longitudinal muscle fibres in a striated arrangement (Figure 4A). If the hypothesis is true, then it would be expected that a driver with similar expression level in the late pupal muscle would also display striated aggregates. The 24B driver is expressed in the muscle with a comparable expression level in late pupae to Mef2, but not in adult muscles (Figure S3B) ^32^. With 24B, it can be seen that, like Mef2, L3 larvae have dispersed aggregates (Figure 4A), while late pupae and young adults display striated aggregates (Figure 4B). None of the 24B flies used for imaging survived passed 20 days (Figure 4D). From these results, it is clear that the striated distribution occurs in the late pupal stage. It remains possible that the striation at the end of metamorphosis may cause lethality and the adults obtained are escapers. The developmental lethality was measured with Mef2 and no significant increase in lethality was observed at any stage (Figure 4C).

**Figure 4.**
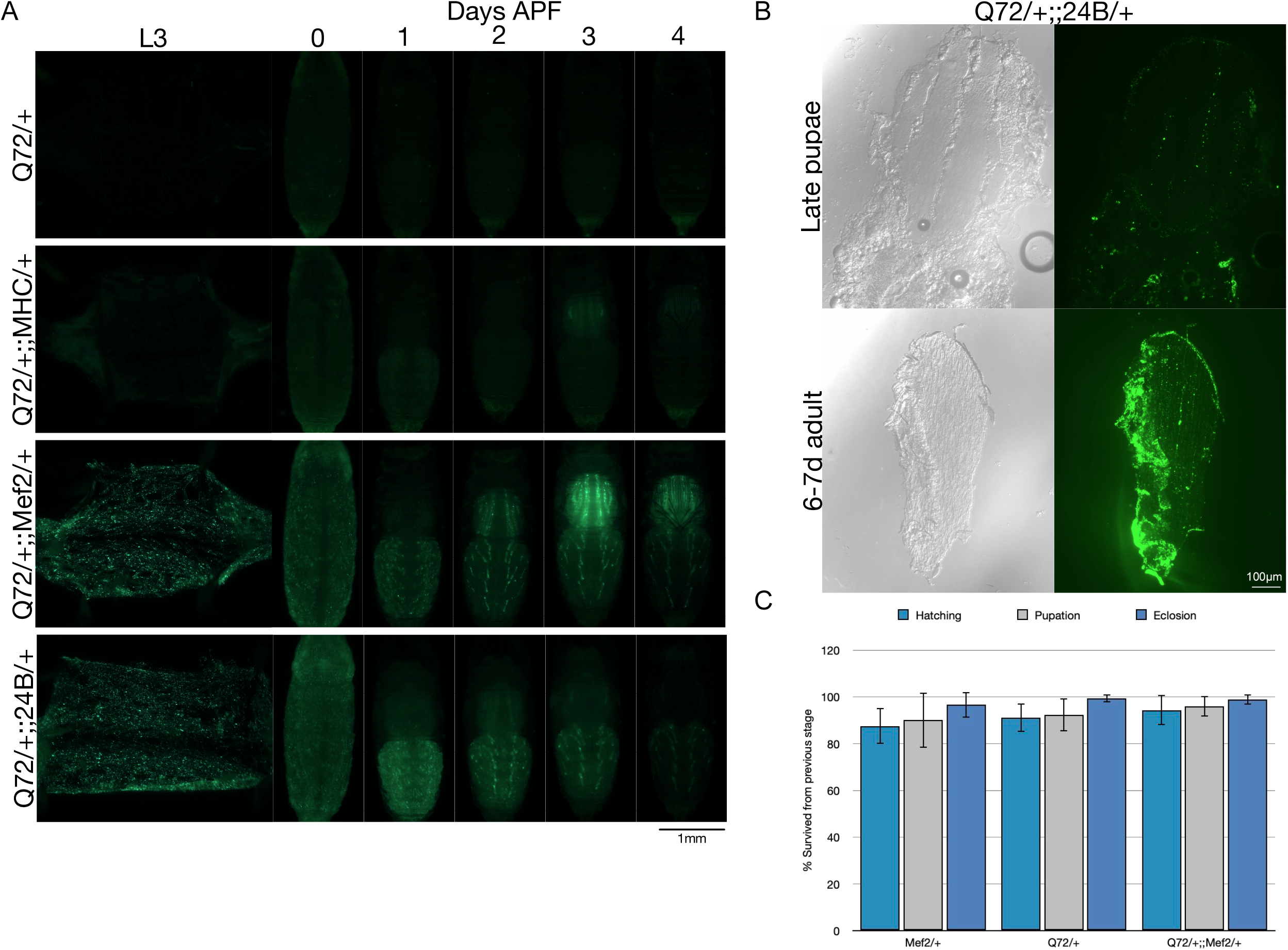
Striated aggregate distribution occurs during metamorphosis and muscle driver 24B also shows striation. A) Imaging of aggregates through development in L3 larval fillet dissections and live whole pupae via fluorescence microscopy. Exposure time is equivalent in animals that have aggregates (Mef2 and 24B). In animals without aggregates (MHC and Q72/+) exposure time was set 2-2.5 times higher to be certain absolutely no fluorescence was detected. B) Cryosectioned Q72/+;;24B/+ late pupae and 6-7d old adults imaged by direct fluorescence and anti-eGFP staining respectively. C) Developmental lethality during hatching (teal), pupation (grey), and eclosion (blue) measured as % survived from previous stage. For each genotype the graph depicts the average of 20 plates of 25 eggs each (n=500). Error bars denote ± SD.

### Striated aggregate distribution can be abolished by temporal repression

From these findings we aimed to test the hypothesis by increasing MHC expression by treating with RU486 in the late pupae to phenocopy the striation. However, treating with RU486 during development caused lethality and morphological defects as previously reported ^36^. We tested lower inducing doses of RU486 but still observed larval and pupal lethality, and the adults that did emerge had defects with their hind tarsi, independent of whether the animal carried the Q72 transgene or not. Alternatively, it has also been documented that GAL4 activation can be influenced by temperature ^37^, where higher temperatures result in higher expression. However, even MHC animals raised at 29℃ did not appear to have any evidence of striation (Figure S6A). Instead, we decided to take an alternate approach where instead of increasing MHC GAL4 activity, we would test the hypothesis by decreasing Mef2 GAL4 activity by using its repressor GAL80. Different transgenes exist to allow for temporal expression of GAL80, including the tetracycline-inducible Tet-off GAL80 transgenes ^32^ and the temperature-inducible GAL80^ts^ ^38^. Each system has caveats ^39^. When measuring longevity, a drug-inducible system would be preferable over temperature, which is known to influence lifespan ^40, 41^, while treatment with a single antibiotic does not ^42^. However, a drug-inducible system would have limitations in induction ability during the pupal stages when the animal is not feeding. Hence both GAL80 systems were used. The repression ability in presence of GAL80 was quantified (Figure 5A) ^39^. In the late pupae, GAL80ts almost completely represses, while with Tet-off GAL80 the activity is reduced but there is still significant expression. Therefore, if the hypothesis is correct it is expected that GAL80ts should eliminate the striation, and the effect of Tet-off GAL80 will depend on the threshold of expression required for striation. In order to conclusively determine whether striation is dependent on the timing of expression, four different treatment groups were used, which induced or repressed across the developmental (preadult) and/or adult stages. The four treatment groups were as follows: no induction, induction only during development, induction only during adulthood, induction across the whole life.

**Figure 5.**
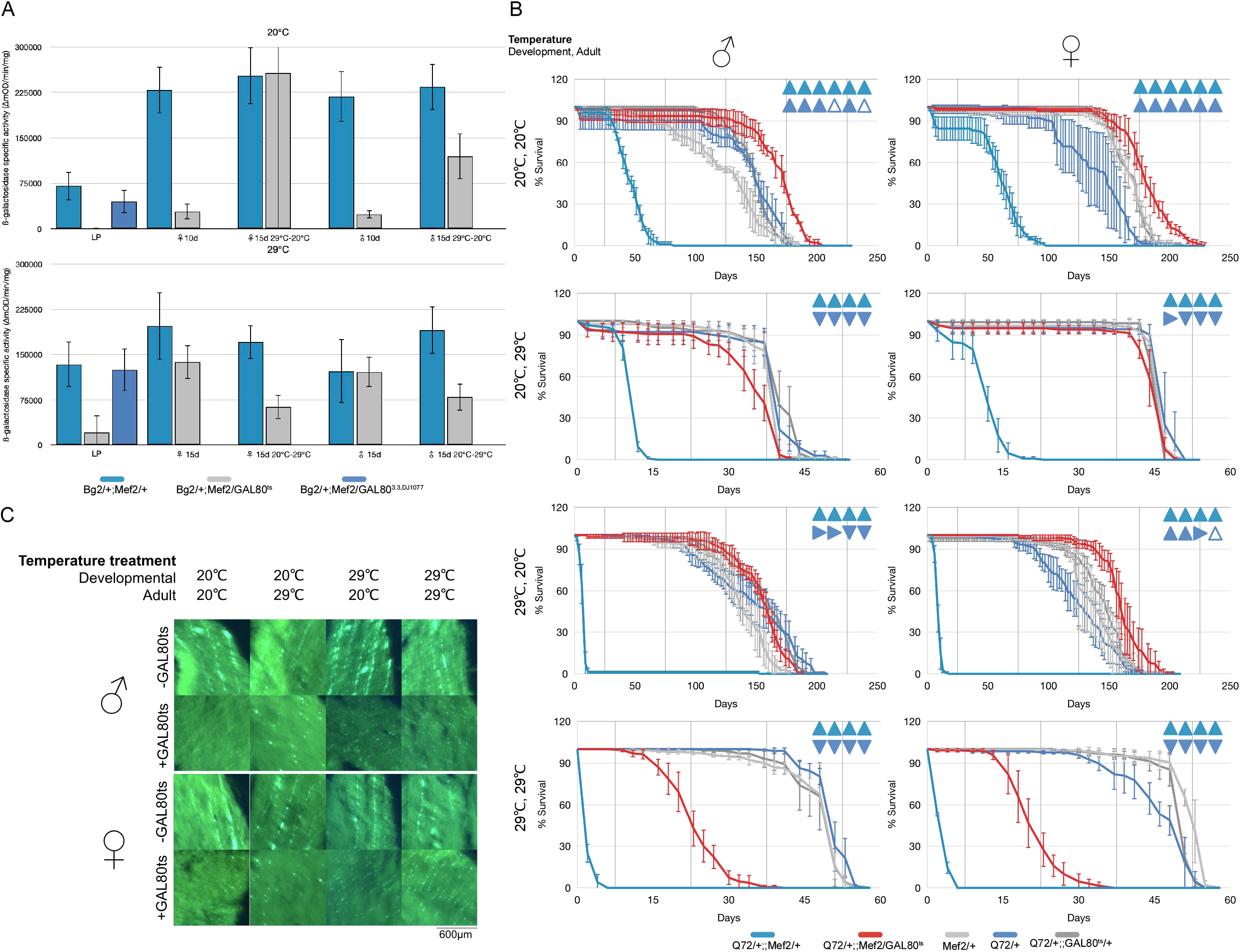
Temporal repression with GAL80^ts^ can rescue shortened lifespan and eliminate striated aggregate distribution. A) UAS-lacZ (Bg2) reporter assay to quantify and compare expression level at permissive and restrictive temperatures with Tet-off GAL80 transgenes and GAL80ts transgenes. LP=late pupae raised at indicated temperature, 10d=10 days old raised and aged at indicated temperature, 15d= 15 days old raised and aged at indicated temperature, 15d 29℃-20℃= 15 days old raised at 29℃ and then aged at 20℃, 15d 20℃-29℃= 15 days old raised at 20℃ and then aged at 29℃. Y-axis represents ß-galactosidase specific activity (ΔmOD/min/mg). Error bars specify ± SD. Bars represent the average of at least four independent replicates of 5 individuals in each (20 total). B)Using the GAL80^ts^ transgene to temporally repress Mef2 mediated expression of Q72. Each graph depicts the survival curve of a different temperature regimen, as indicated. Error bars represent ± SD. Triangles represent the result of the logrank comparision of Q72/+;;Mef2/GAL80^ts^ curves to controls from a minimum of four independent replicates. Teal triangles compare to Q72/+;;Mef2/+. Blue triangles compare to Q72/+. •: significant increase,6: insignificant increase, T: significant decrease, v: insignificant decrease,►: no change C) Cryosectioned flies from B) show striated aggregates in Q72/+;;Mef2/+ flies (-GAL80) and dispersed aggregates in Q72/+;;Mef2/GAL80^ts^ (+GAL80).

A copy of each version of the Tet-off GAL80 transgenes was used. Each version uses a different ubiquitous promoter and it was previously shown that a copy of each version represses the DJ694 driver better than two copies that use the same promoter ^32^. With the addition of GAL80 it was found that the longevity was extended in both males and females (Figure S6B), more so in males than females. The aggregates were also examined in these genotypes (Figure S6C). The addition of GAL80 made an indiscernible difference, all of the aggregates still appear striated. Furthermore, there were no obvious differences between the different treatment groups.

When the GAL80^ts^ transgene is used, it is clear in all temperature treatments and across all replicates that the addition of GAL80 significantly extends lifespan (Figure 5B). The degree of extension can be assessed by comparing to one of the other controls with a wildtype lifespan (Q72/+). The only temperature treatments where the GAL80^ts^ genotype was significantly shorter lived relative to wildtype, across all replicates, were the males raised at 20°C and aged at 29°C, and both the males and females that were maintained at 29°C. The shorter lifespan at 29°C indeed results from the inactivation of GAL80 and the severity of the reduction is in agreement with the level of expression (Figure 5A). However there is a significant extension compared to the control without GAL80 because the inactivation of GAL80 is incomplete (Figure 5A)^39^. The normal longevity of animals raised at 29°C and aged at 20°C contradicts the hypothesis that higher expression during development is more detrimental. Unfortunately, it turns out that GAL80ts cannot be inactivated resulting in a very low level of expression in the muscle of late pupae (Figure 5A) ^39^. Indeed, upon examination of the aggregate distribution it revealed that the aggregates at 29°C are not striated but dispersed like those seen with the MHC and DJ694 drivers (Figure 5C).

Taken together, these results conclusively demonstrate that the striated aggregate distribution relies on the level of expression. Furthermore, these data demonstrate that the severe symptoms are linked to the striated aggregate distribution because when abolished it significantly extends the lifespan back to a level comparable with wildtype. It is unfortunate that the dynamics of temperature induction with GAL80ts makes it impossible to provide the direct demonstration in genetically identical individuals that high expression only during development is required and sufficient for the striated distribution and severe reduction of longevity.

### Molecular mechanisms of toxicity differ in muscle compared to the nervous system

Since the polyglutamine repeats were found to be toxic to the muscle we next wanted to determine the molecular mechanism of toxicity. Indeed, in studies of the nervous system numerous genetic modifiers have been identified that contribute to the understanding of the molecular mechanisms of the disease. The chaperone, Hsp70 was chosen as a candidate gene since the overexpression has been shown to suppress polyglutamine toxicity in a model of SCA3 ^43^, and the deletion enhances toxicity in a model of SCA1 ^12^. Further, we have demonstrated previously that the overexpression of Hsp70 has no effect on lifespan ^44^. We had previously made transgenic lines carrying the *Drosophila* gene under the control of a UAS promoter ^45^. *Drosophila* Hsp70 has never been tested for its ability to modify polyglutamine toxicity in a model of HD and previous studies used a human Hsp70 gene ^12, 43^. Two different insertions of the UAS-hsp70 construct (4.2 and 4.4) were introduced into the background of Q72 (Q72;;UAS-hsp70) and crossed with GMR-Gal4 to drive expression in the eye (Figure 6A). To control for GAL4 titration effects brought on by the addition of another UAS transgene, a UAS-lacZ (Bg3) transgene was also built into the background of Q72 (Q72;;UAS-lacZ), and then crossed with GMR-Gal4. In *Drosophila* neurodegeneration studies, the eye is used to model the nervous system and depending on the strength of the pathogenic transgene, it can cause loss of photoreceptors, the collapse of ommatidial structure and the presence of protein aggregates. Expressing Q72 in the eye did not result in any obvious morphological defects but does result in an accumulation of polyQ aggregates, as previously reported^29^. It was found that the addition of either hsp70 insertion drastically reduced the amount of aggregates that accumulate in the eye.

**Figure 6.**
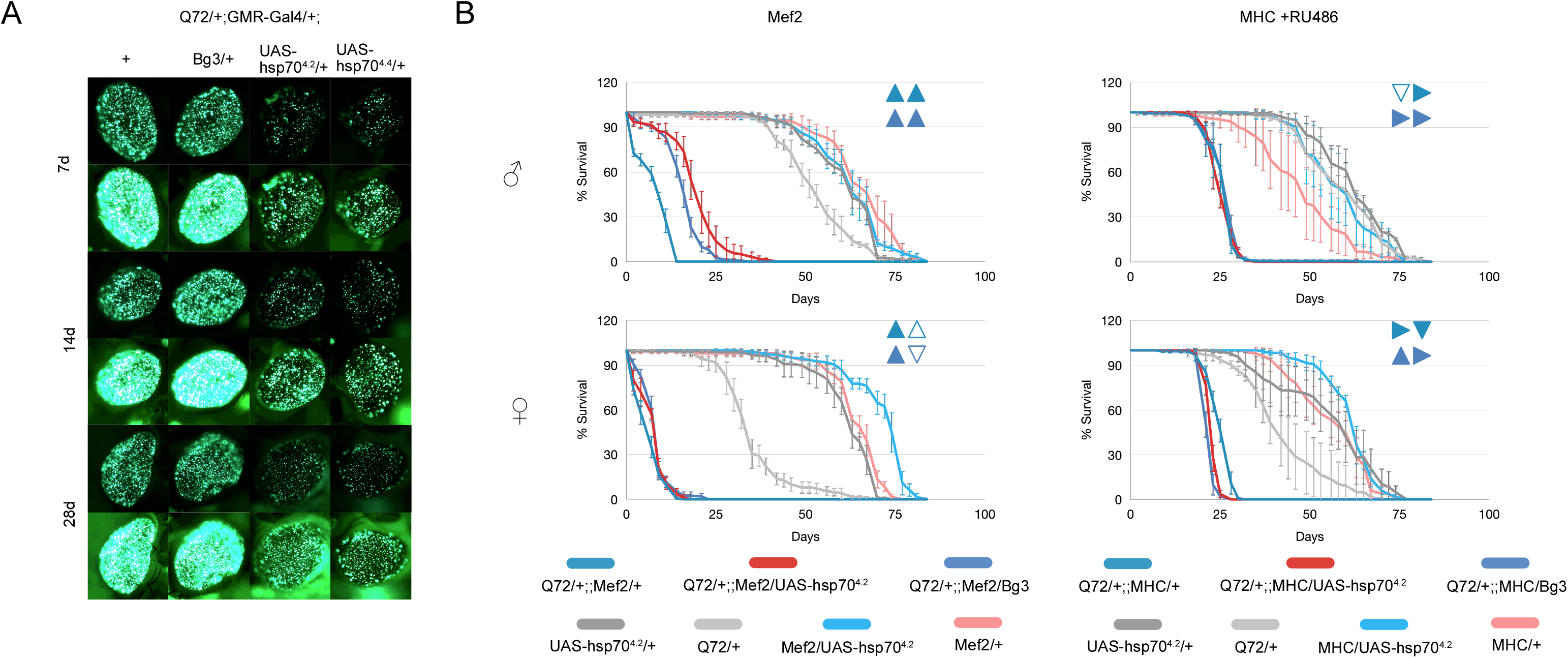
Hsp70 does not suppress the detrimental phenotype caused by polyglutamine in muscle but does in the eye. A) Effect of Hsp70 on polyglutamine aggregates in the eye of the fly. All genotypes carry the Q72 and GMR-Gal4 driver transgenes, and on the third chromosome either a wildtype (+), UAS-lacZ (Bg3/+), or UAS-hsp70 (UAS-hsp70^4.2^/+ & UAS-hsp70^4.4^/+). Flies were collected and allowed to age, and sampled at 7, 14, 28d to dissect and image. Images were taken at a low and high exposure to best capture the amount of aggregates. Relative low and high exposure photos were taken under same conditions. B) Survival curves show the effect of expressing UAS-hsp70^4.2^ in the background of flies expressing Q72 in the muscle driven by Mef2 and MHC drivers. MHC genotypes were treated with RU486 (50µg/ml), Mef2 is maintained on standard fly food. Error bars denote ± SD. Triangles represent the result of the logrank comparision of Q72/+;;Mef2/UAS-hsp70^4.2^ curves to controls from two independent replicates. Teal triangles compare to Q72/+;;Mef2/+. Blue triangles compare to Q72/+;;Mef2/Bg3. ▴: significant increase,△: insignificant increase, ▾: significant decrease, ▽: insignificant decrease, ▶: no change

To test the effect of Hsp70 in the muscle, the Q72;;UAS-hsp70 and Q72;;UAS-lacZ lines were crossed with each MHC and Mef2. Both drivers were tested to gain a more comprehensive understanding of the mechanistic differences between the dispersed and striated aggregate distributions. With MHC, the addition of 4.2 did not result in any significant difference or no consistent effect on longevity across replicates compared to either the control with no hsp70 (Q72/+;;MHC/+) or the control with UAS-lacZ (Q72;;MHC/UAS-Bg3) (Figure 6B). In Mef2 flies, females show no consistent significant effect across replicates, while in males there is a modest extension observed. With 4.4, a significant reduction in lifespan is observed with both drivers and sexes, across all replicates (Figure S7). The control genotype, 4.4/+, also appears to have a shortened lifespan indicating that the reduction is most likely attributed to where the transgene is inserted in the genome, rather than hsp70 being an enhancer of polyglutamine toxicity. We conclude that Hsp70 does not affect the reduced longevity resulting from the presence of either dispersed or striated HD aggregates, which contrasts the strong effect in the nervous system. These findings imply the molecular mechanisms underlying the effects of HD aggregates in muscles are distinct from the nervous system.

## DISCUSSION

We report here that not only are polyglutamine repeats toxic to the muscle, but the protein aggregates can take on distinct distributions depending on the level of expression during development. These alternate distributions are associated with differing severity of symptoms. The dispersed distribution was observed with DJ694 and MHC, drivers that express in the mature animal when the adult muscle has already formed. It was shown that in the dispersed mode by increasing the expression level the effect on longevity and the number of aggregates could be aggravated. Although the phenotype could be exacerbated, it could not be made as severe as that of the striated distribution. The striated distribution was observed with Mef2 and 24B, drivers that have high expression during the juvenile or pre-adult stages. The aggregates were observed to arrange into the striated distribution during metamorphosis, but no developmental lethality was observed indicating the detrimental effects only manifest in the adult animal. This would imply that the perturbations caused by the disease can occur during development, and have a much more severe effect in the adult animal than those that happen during adulthood. This adds to the growing evidence showing that although HD is a late-manifesting disorder, the disease causes abnormalities in presymptomatic carriers during development ^46^. The localization of the aggregates with respect to the sarcomere remains unclear. There are many antibodies and reporter lines that now exist that mark the various stages from myogenesis to sarcomerogenesis ^47, 48^. It would be beneficial to use these markers to determine where the striated aggregates are localizing compared to the dispersed.

Since we express exclusively in the muscle it brings to question, what kills the fly? One might expect that causing damage to a tissue like the muscle would be detrimental to the animal’s ability to move, but not necessarily affect survival. While we can only speculate, it is possible that expressing the toxic protein in the muscle triggers rhabdomyolysis. Rhabdomyolysis is clinically defined by the breakdown and necrosis of muscle which causes a release of intracellular components into the blood stream ^49^. Anecdotally, it was noticed that many of the dead flies expressing Q72 were black in colour, a feature of necrosis (Figure S8). Rhabdomyolysis can be caused by physical trauma or injury, overexertion, genetic disorders, and environmental causes like alcohol, drugs, toxins, or infections. Diagnosis is typically performed by blood analysis showing elevated creatine kinase, myoglobin, and creatinine among other indicators, and urinalysis showing blood in the urine ^49^. Creatine kinase has been measured in HD mouse models via enzymatic activity and microarray analysis and found to be perturbed in brain and muscle ^50, 51^. The functional relationship between the changes in gene expression in nerve and muscle are striking. Also, creatine supplementation has been shown to improve survival, motor symptoms and reduce gross brain atrophy and aggregates ^52, 53^. It should be possible to extract hemolymph from our flies ^54^ and measure creatine kinase levels via enzyme activity for which kits exist, or western blotting with an anti-creatine kinase antibody. So far, no specific connections have been made between rhabdomyolysis and Huntington’s disease but given one of the major symptoms of the disease is muscle atrophy, it may be worth investigating.

It is also unclear what causes the unique striated aggregate distribution. At this time we can only speculate. It cannot be overlooked that the onset of striation coincides with the formation of the indirect flight muscles. During pupation the majority of larval oblique muscles (LOMs) histolyze, except for three mesothoracic muscles which give rise to the dorsal longitudinal muscles (DLMs). Unlike the dorsoventral muscles (DVMs) that form *de novo*, the DLMs are formed from the remaining mesothoracic LOMs which change shape, vacuolate, and split into the six DLM fibres ^55^. It is possible that the aggregates are positioned during this process and get integrated into the muscle fibre. This could explain why the phenotype is so severe.

Alternatively, the striation may be brought about by the aggregates aligning along the recently described autophagy-dependent tubular autolysosomal network (tAL) ^56^. This extensive structural network has been demonstrated to have degradative activity required for regulating muscle remodeling during metamorphosis. If the aggregates interfere with the efficiency of lysosomal tubulation this could exacerbate symptoms. Although tAL was initially studied in abdominal muscles, similar structures have been observed in the indirect flight muscles of the thorax ^56^.

Another possibility is that the aggregates are intercalating between the myofibrils like the mitochondria. A mechanical feedback mechanism involving the coordination of mitochondria and sarcomere morphogenesis has recently been described ^57^, in which the mitochondria of the flight muscle are densely packed taking on an ellipsoid shape from the mechanical tension of the myofibrils. As a result, the mitochondria have a striped appearance similar to that of the striated aggregates. It is possible the aggregates could be interfering with the arrangement of the mitochondria which may cause disruptions in oxidative respiration and in turn contribute to the very severe effect on longevity and locomotion. The leg muscle on the other hand is known to have a cross-striated myofibrillar morphology, with myofibrils aligned laterally forming a tube and mitochondria located peripherally and centrally in the tube ^58^. From our imaging we can see striated aggregates in the legs but they tend to appear more bleb-like compared to the sharper stripes in the thorax. It may be worth investigating whether the aggregates co-localize with the mitochondria and further whether oxidative respiration is impaired.

Research initiatives in HD have traditionally focused on the biological processes mediating the neurological symptoms during adult life. However, our findings indicate that in the striated distribution, aggregates produced in pre-adult stage result in the most severe longevity and behaviour phenotypes. Although HD is most often described as an adult late onset disease, in certain cases HD has been known to affect young individuals before 20 years of age (Juvenile HD) and is associated with the most severe symptoms and devastating progression rates. Although there is a correlation between the age of onset and the length of the repeats there are exceptions ^59^. It is tempting to speculate that the severe symptoms of Juvenile HD (JHD) may also result from aggregates being produced before adulthood.

The fact that driving polyglutamine in the muscle can recapitulate HD symptoms (shortened lifespan, reduced locomotion, aggregate formation) provides a genetic model to elucidate the molecular mechanisms of the symptoms of this disease. We show that Hsp70 suppresses aggregate accumulation in the eye but had no effect in the muscle, indicating that the mechanisms of toxicity in muscle and nervous system differ. Our model can be used to conduct screens on multiple phenotypes and two different aggregate distributions. A genetic approach may reveal whether the alternate distribution is caused by aggregate composition, morphology, or both, which would be very challenging to address biochemically.

## LIMITATIONS OF THE STUDY

Although this study could lead to better treatments and diagnostics, it is obviously limited to identify the mechanisms underlying the symptoms in peripheral tissues and will not reveal molecular targets to prevent all symptoms and cure the disease.

This study expanded our current understanding of the biological consequences of HD aggregates in muscle and will require clinical investigations that have been largely neglected. It remains unknown whether distinct aggregate distributions occur in the muscle of human patients. Aggregates have only been observed in the muscles of two HD patients and Juvenile HD patients have not been examined.

It is unknown whether the alternate aggregate distributions are specific to *huntingtin* and HD, or could apply to any polyglutamine disease or proteopathic disease model. The large number of UAS constructs available in *Drosophila* will make it trivial to test other polyglutamine proteins, naked polyQ tracts, or any other aggregate-prone proteins and will reveal whether the different aggregate distributions are specific to the protein expressed, or whether any aggregated protein would arrange that way.

## METHODS

### RESOURCE AVAILABILITY

#### Lead contact

Further information and requests for resources and reagents should be directed to and will be fulfilled by the lead contact, Laurent Seroude.

#### Materials availability

This study did not generate new unique reagents.

#### Data and code availability

- Source data generated in this study is available in Supplemental Data. Microscopy data reported in this paper will be shared by the lead contact upon request. Requests for resources and reagents used in this study should be directed to the lead contact.
- Original R code for longevity analysis is included in the Supplemental Data.
- Any additional information required to reanalyze the data reported in this paper is available from the lead contact upon request.

### KEY RESOURCE TABLE

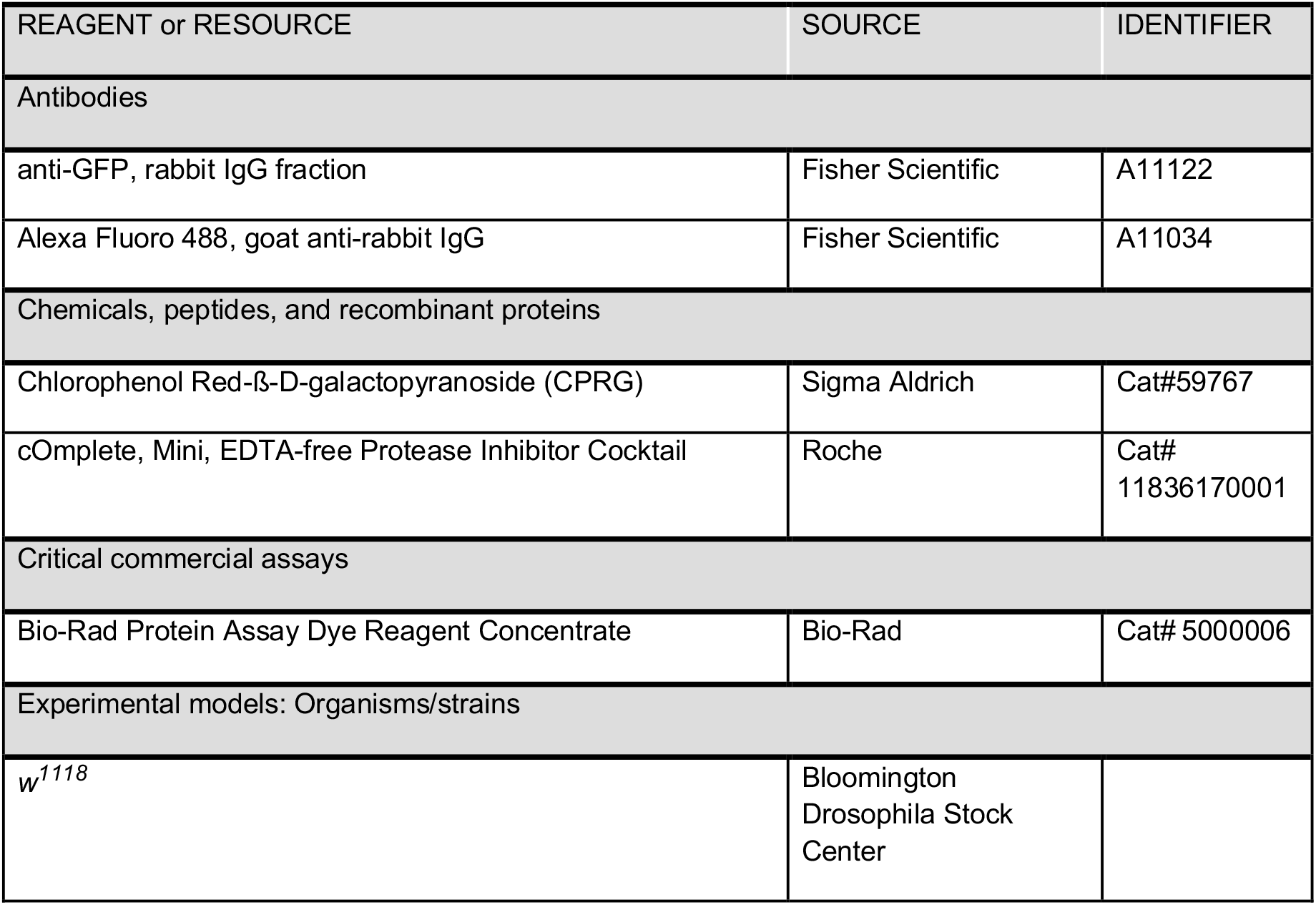
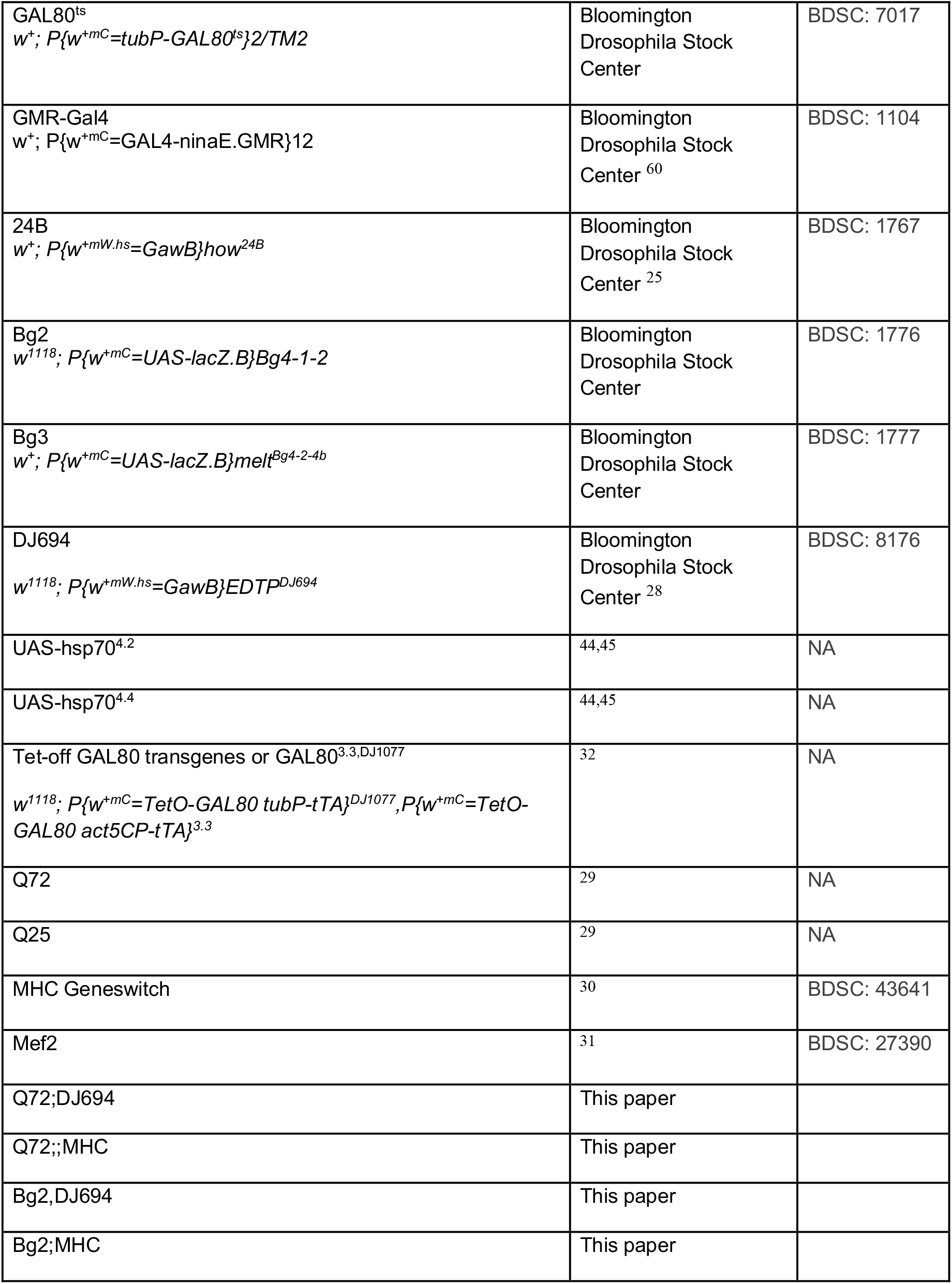
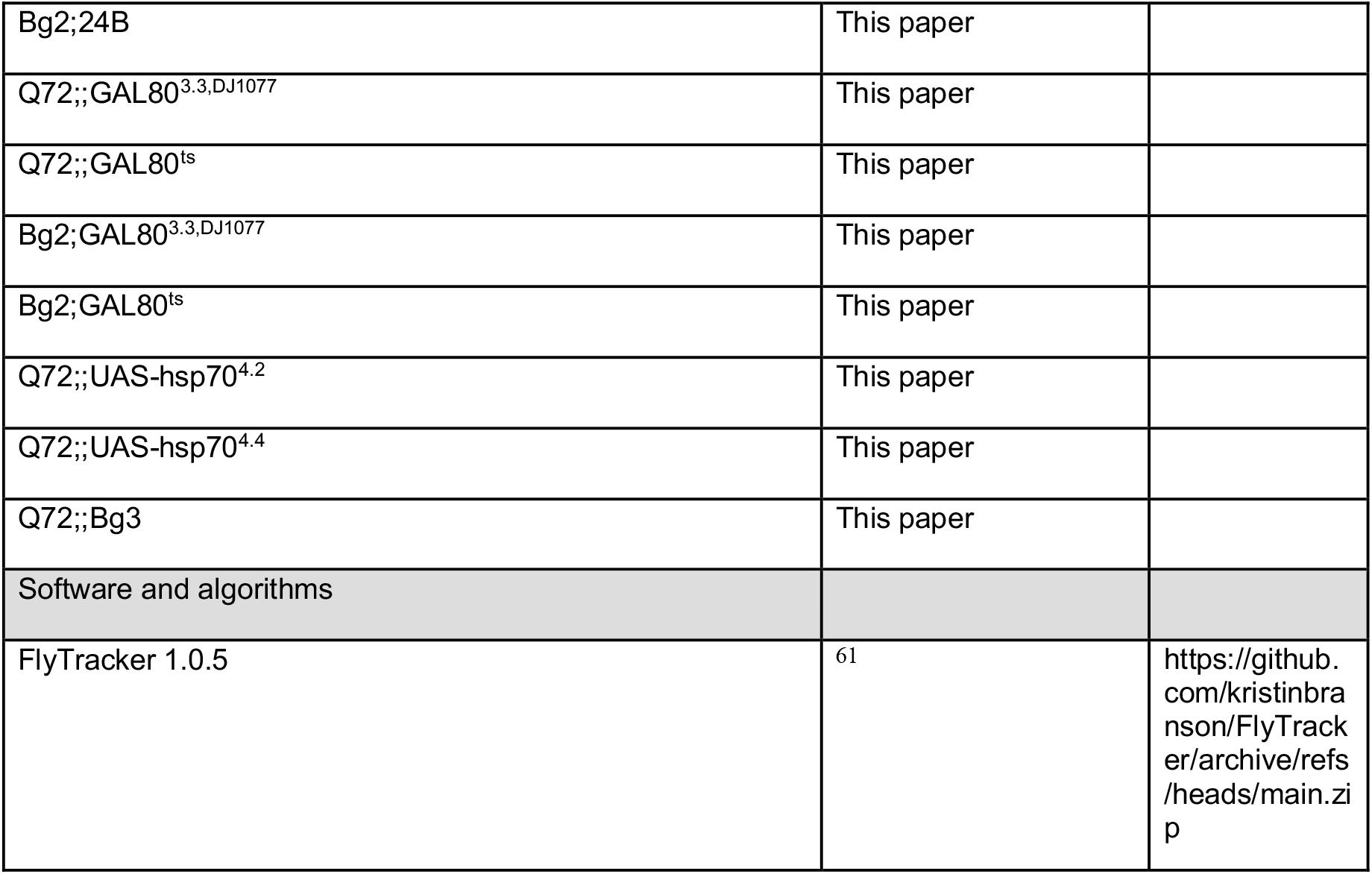

### EXPERIMENTAL MODEL AND SUBJECT DETAILS

All fly lines used in this study are described in the Key Resources Table. For all experiments, male and female flies were reared and aged to specific timepoints as per experimental requirements. Flies were maintained on standard cornmeal medium (0.01% molasses, 8.2% cornmeal, 3.4% yeast, 0.94% agar, 0.18% benzoic acid, 0.66% propionic acid) ^62^ at 25℃ on 12h light/dark cycle unless stated otherwise.

RU486 was administered as follows. A stock solution of RU486 was prepared in 100% EtOH at 25mg/ml. The stock was diluted down to 50µg/ml using water as solvent. 200µl of the 50µg/ml solution was applied to the surface of vials of standard fly food and allowed to absorb for 48-72h. The control vials for -RU486 treatment were prepared simultaneously by applying 200µl of 0.2% EtOH to the surface of vials of standard fly food and allowing to absorb. It was previously described this was the most effective drug delivery method ^27^.

Tetracycline was administered as follows. A stock solution of tetracycline was prepared in water at 100mg/ml. For developmental treatment the stock solution was diluted down to 100µg/ml in melted standard fly food and distributed to bottles. Crosses were set up in these bottles, untreated crosses were set up on bottles of standard fly food. For adult treatment the stock solution was diluted down to 100µg/ml in water and 200µl was applied to the surface of vials of standard fly food and allowed to absorb for 48-72h.

### METHOD DETAILS

#### Longevity assay

Experimental genotypes for Figures 1 and S1 were obtained by crossing females of each UAS line (Q72 & Q25) with males of each driver line (DJ694, MHC, Mef2). Driver control genotypes were obtained by crossing female w^1118^ flies with males of each driver line. UAS control genotypes were obtained by crossing females of each UAS line with w^1118^. Crosses were set up in bottles (Bottle 1) and after 48h flipped into a second set of bottles (Bottle 2). A minimum of four age synchronized vials at a density of approximately 25-30 flies were collected of each sex, from each cross, from each Bottle 1 and 2. Flies were collected under nitrogen anesthesia. Flies were transferred to fresh food and scored for deaths every 2-3 days. The above steps were repeated to generate a completely independent replicate. All longevity assays have a minimum of two independent replicates.

For longevity assays with multiple copies of UAS and GAL4 transgenes (Figures 3, S4), genotypes were initially built by standard genetic crosses that combined Q72 with either MHC or DJ694 (Q72; driver). Virgins were collected of each of these lines and then crossed with sibling males of the same genotype, Q72, males of either MHC or DJ694, and w^1118^. The resultant progeny from these crosses effectively generated genotypes with 2 copies of each UAS and GAL4, 2 copies UAS 1 copy GAL4, 1 copy UAS 2 copy GAL4, and 1 copy of each UAS and GAL4 respectively. Indeed, since Q72 lies on the X chromosome, the resultant male progeny can only carry 1 copy of the UAS, thus only GAL4 copy number can be tested and the genotypes were obtained from two separate crosses.

For longevity assays with Tet-off GAL80 (Figure S6), genotypes were initially built by standard genetic crosses that combined Q72 with GAL80^3.3,DJ1077^ (Q72;;GAL80^3.3,DJ1077^).

Virgin Q72;;GAL80^3.3,DJ1077^ and Q72 were crossed with Mef2 males to obtain experimental genotypes. In order to temporally induce or inhibit GAL80, flies were treated with four possible tetracycline regimens: 0µg/ml tetracycline across development and adulthood, 0µg/ml during development and 100µg/ml during adulthood, 100µg/ml during development and 0µg/ml during adulthood, and 100µg/ml across development and adulthood. In order to administer the appropriate developmental treatment, crosses were done in duplicate where one bottle contained 100µg/ml tetracycline and the other 0µg/ml. Upon eclosion the adult flies from each bottle were collected and half were put on 100µg/ml tetracycline and the other half 0µg/ml.

For longevity assays with GAL80ts (Figure 5), genotypes were initially built that combined Q72 with GAL80ts (Q72;;GAL80ts). Virgin Q72;;GAL80ts and Q72 were collected and crossed with Mef2 and w^1118^ males. A driver control genotype was obtained by crossing w^1118^ females with Mef2 males. In order to temporally induce or inhibit GAL80, flies were treated with four possible temperature regimens: 20℃ during development and adulthood, 20℃ during development and 29℃ during adulthood, 29℃ during development and 20℃ during adulthood, and 29℃ during development and adulthood. In order to administer the appropriate developmental temperature, crosses were done in duplicate where one bottle was put at 20℃ and the other at 29℃. Upon eclosion, the adult progeny from each bottle were split, half were put at 20℃ and the other half 29℃.

For longevity assays with Hsp70 (Figures 6 and S7), lines were initially built in which two different UAS-hsp70 insertions (4.2 and 4.4), as well as UAS-lacZ (Bg3), were each combined with Q72 (Q72;;4.2, Q72;;4.4, Q72;;Bg3). Virgin Q72;;4.2, Q72;;4.4, Q72;;Bg3 and Q72 were crossed with Mef2 and MHC males to obtain experimental genotypes. Control genotypes were generated by crossing 4.2, 4.4 and Q72 females with w^1118^ males, as well 4.2, 4.4 and w^1118^ females were crossed with Mef2 and MHC males. MHC genotypes and associated controls were maintained on RU486 (50µg/ml) containing food.

#### Locomotion behaviour assay

Locomotion assays were performed as described ^62, 63^. Flies were aspirated from the longevity vials, recorded for 3 minutes (30 frames per second), and returned to their vials. In a given video we recorded all five genotypes corresponding to one of the drivers in triplicate. Both Bottle 1 and 2 flies were measured each week. Flies were aspirated from one of the eight longevity vials that were collected, alternating the vial at each recorded time point. The positions of the various genotypes in the arenas were rearranged at each recorded time point to eliminate effects associated with any specific arena. Five different arena conformations were used. Tracking and data analysis was completed using FlyTracker and a custom MatLab script to export the data to Excel.

#### Cryosectioning and Immunochemistry

Whole flies were embedded in Optimal Cutting Temperature compound (OCT, Tissue-Tek) for approximately 20min. A layer of OCT was applied to the sectioning plates from the cryostat microtome. Flies were then positioned sagittally atop this layer and covered with OCT medium and frozen (-15℃). Sections were cut from this block that were 20µm thick and adhered to slides. GFP was visualized in sections by immunochemistry or direct excitation and gave identical expression patterns, indicating no artifacts as a result of fixation ^34^. Sections in Figures 2, S2, adults in 4, 5 were immunostained as indicated below. Sections in Figures 3, pupae in 4, S6 were mounted in 80% glycerol and imaged immediately after sectioning. Sections were fixed with Mirsky’s fixative or 2% glutaraldehyde for 10 minutes at room temperature. Slides were washed (1X PBS, 0.2% Triton X) five times for 5 minutes before blocking (1X PBS, 0.2% Triton X, 5% BSA) for 30 minutes at room temperature. Slides were incubated in anti-GFP, rabbit (1:1000 in blocking) at room temperature overnight. Slides were washed again five times for 5 minutes and then incubated with blocking buffer for 30 minutes. Slides were incubated in Alexa Fluoro 488, goat anti-rabbit IgG (1:1000 in blocking) at room temperature in the dark overnight. Slides were washed again five times for 5 minutes and then mounted with Fluoromount-G with DAPI. Images were obtained using a Zeiss Axioplan II imaging microscrope with a Leica DC500 high-resolution camera and the OpenLab imaging software (Improvision, Lexington, MA, USA). All images were captured with the same exposure time and light intensity.

#### CPRG Assay

Details on the CPRG assay protocol have been previously described ^64^. For single copy measurements (Figure S3), experimental genotypes were obtained by crossing female UAS-lacZ (Bg2) with males of each driver (MHC, Mef2, DJ694, 24B) line and w^1118^. Five individuals were measured per experimental replicate and at least two experimental replicates were performed. Measurements were done on third instar larvae (L3), early pupae (EP) and late pupae (LP), as well as 0-5 and 10-11 day old adults. In order to enrich extracts for muscle tissue, in all adult measurements thoraces were dissected. Specific activity measurements represent the average across all experimental replicates. For measurements of genotypes with multiple copies of UAS and GAL4 (Figure S5), genotypes were initially built by standard genetic crosses that combined Bg2 with each of the driver lines (DJ694, MHC, 24B). Virgin females of these lines were collected and crossed with sibling males, Bg2, the corresponding driver line, and w^1118^ to obtain experimental genotypes. Measurements were done on EP, LP, and 0-2 day old adult thoraces. Two experimental replicates with five individuals each were measured and averaged.

For measurements of genotypes with GAL80 (Figure 5), genotypes were initially built by standard genetic crosses that combined Bg2 with each type of GAL80 transgene (GAL80^ts^ and GAL80^3.3,DJ1077^). Virgin females of these lines and Bg2 were collected and crossed with Mef2 males to obtain experimental genotypes. For these experiments animals were raised and/or aged at different temperatures, in order to inhibit or induce GAL80^ts^, and simulate the temperature treatments in the longevity assay. Four different temperature treatments were used: 20℃ during development and adulthood, 20℃ during development and 29℃ during adulthood, 29℃ during development and 20℃ during adulthood, and 29℃ during development and adulthood. Measurements were done on LP and 10 or 15d adult thoraces for the different temperature treatments. At least two experimental replicates with five individuals each were measured and averaged.

#### Developmental Lethality Assay

Desired genotypes were generated by crossing Q72 females with Mef2 and w1118 males, as well as w1118 females with Mef2 males. Crosses were set up in vials, allowing 48h for mating. The flies were moved to egg collectors (egg collector medium: 2% agar, 5% sucrose, dyed with neutral red; dolloped with yeast paste: yeast with water until peanut butter consistency) with a minimum of 50 females and 50 males and allowed to cross overnight (12-16h). Embryos were aligned onto the disks (25 per disk) of standard fly food and then transferred on top of a standard vial of food to continue development. Hatching was scored 26-30h later by counting empty eggshells. Pupation and eclosion were scored 5-6d and 10-11d respectively from the time the eggs were aligned.

#### Whole animal GFP imaging

To visualize protein aggregates (Figure 4), desired genotypes were obtained by crossing female Q72 with w1118, MHC, Mef2 and 24B males. From these crosses wandering L3 larvae were collected and fillet dissections were performed and imaged. Pre-pupal (immobile L3) individuals were collected with a wet paintbrush and transferred to a microscope slide. The slide was maintained in a petri dish with moist paper towel for humidity. The staged pupae were imaged once per day from the date of collection (0d APF) until the day before eclosion (4d APF). Imaging was performed using a Zeiss dissecting microscope fitted with a Leica DC500 high-resolution camera and the OpenLab imaging software (Improvision, Lexington, MA, USA). All images were captured with identical light intensity and exposure time, except for genotypes in which GFP was negative, in such cases relative exposure time is provided.

#### Eye GFP imaging

To visualize protein aggregates in the eye (Figure 6), desired genotypes were obtained by crossing male GMR-Gal4 with Q72, Q72;;Bg3, Q72;;4.2 and Q72;;4.4 females. Progeny were collected and aged before imaging. Eyes were imaged at 7, 14 and 28 days of age. Eyes were isolated for imaging by decapitating the flies using a fly collar or guillotine ^65^. Subsequently, heads were dissected sagittally so the eye would lay flat for imaging. All images were taken with identical light intensity and exposure time.

### QUANTIFICATION AND STATISTICAL ANALYSIS

All longevity data was analyzed and compiled in R by Logrank and Bonferroni post-hoc tests. When comparing the lifespan of different genotypes the threshold for significance was if the difference in mean lifespan was 5% or greater, and the p value from the Logrank was <0.05. Comparison of ß-galactosidase expression level was done by one way ANOVA and Dunnett’s post test in GraphPad Prism. Two-tailed p values of <0.05 were considered the cutoff for statistical significance. The value of n represents the number of animals measured.

