## Supplemental Figures and Data for "Juvenile and adult expression of polyglutamine expanded *huntingtin* produce distinct aggregate distributions in *Drosophila* muscle"

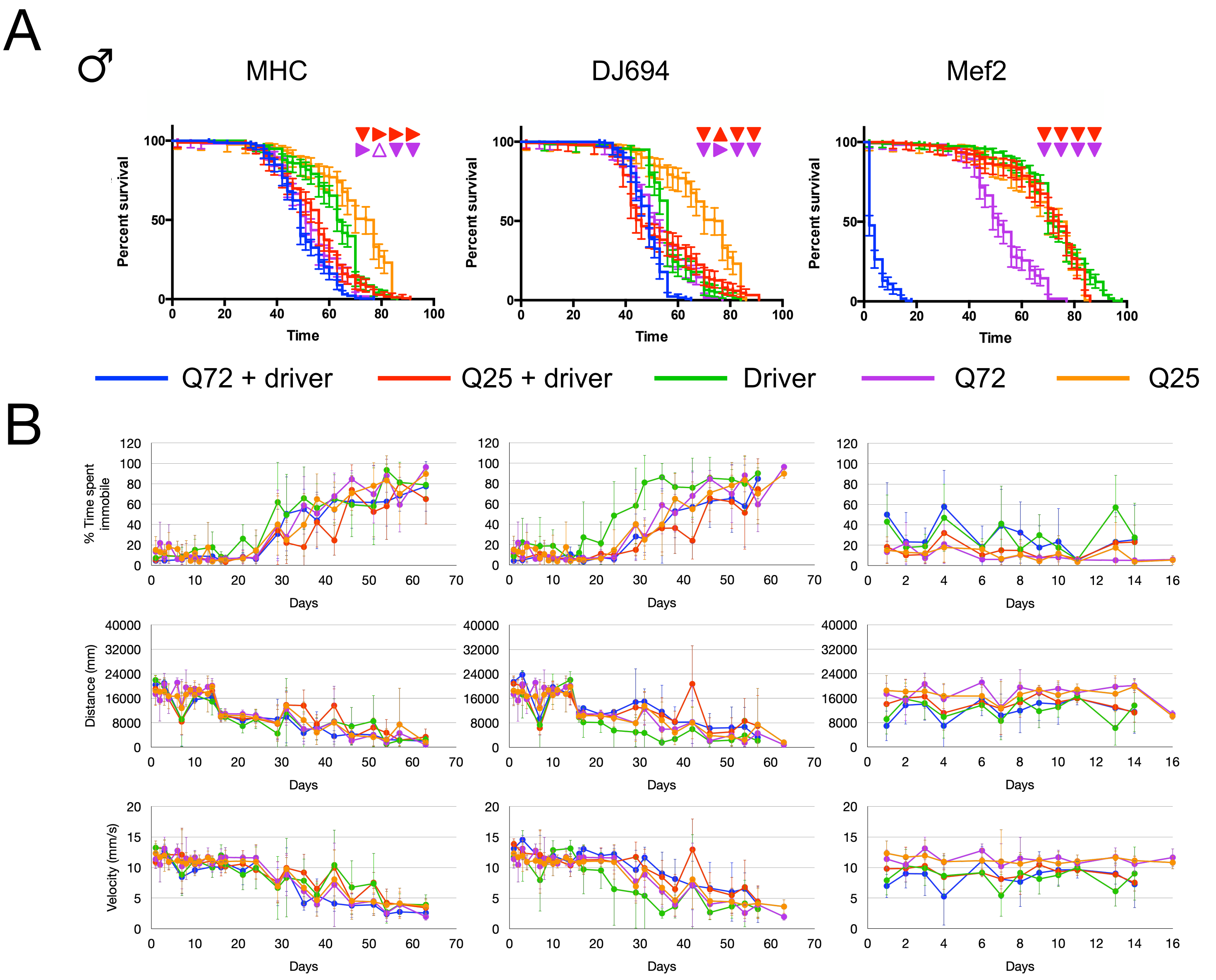

**Figure S1. Expression of UAS-Htt-Q72 in the muscle results in declines in longevity and locomotion.** All data from male flies. Three muscle specific GAL4 drivers are used: MHC-Geneswitch, DJ694, Mef2. A) Lifespan assay. X-axis denotes time in days at 25°C, y-axis denotes percent survival, error bars represent 95% CI. Triangles represent the result of the logrank comparison of Q72+driver curves to controls from the four independent replicates. Red triangles compare to Q25+driver. Violet triangles compare to Q72 alone. ▲: significant increase, △: insignificant increase, ▼: significant decrease, ▽: insigificant decrease, ▶: no change B) Locomotion data measured by three parameters: % time spent immobile (<1mm/s), distance travelled (mm), velocity (mm/s). Error bars represent ± SD. Genotype abbreviations: *Q72+driver (blue)*: Q72/+;;MHC/+, Q72/+;DJ694/+, Q72/+;;Mef2/+ ; *Q25+driver (red)*: Q25/MHC, DJ694/+;Q25/+, Q25/Mef2 ; *Driver (green):* MHC/+, DJ694/+, Mef2/+ ; *Q72 (violet):* Q72/+ ; *Q25 (orange):* Q25/+

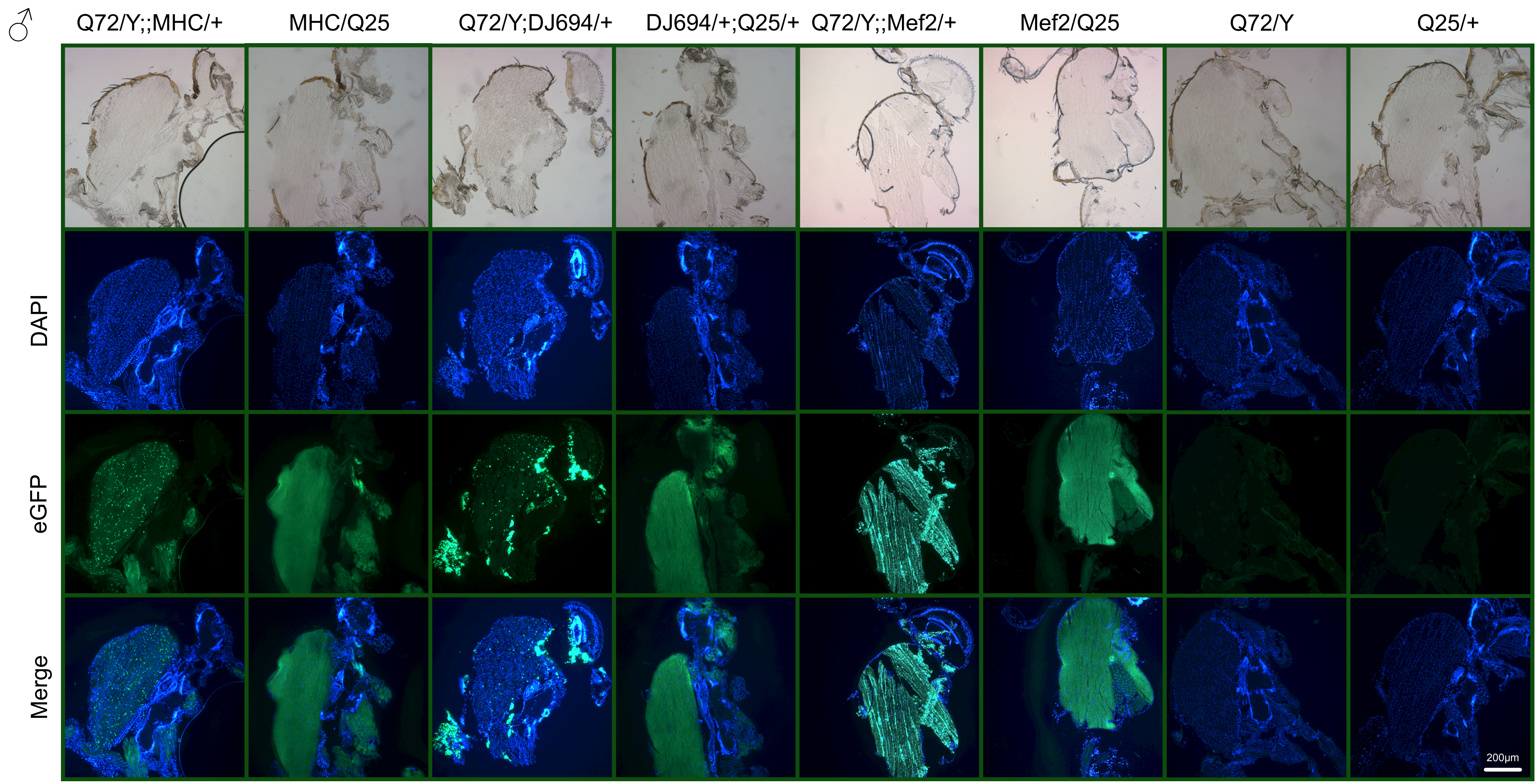

**Figure S2. Alternate aggregate distributions with different muscle drivers** Visualization by saggital cryosectioning and anti-eGFP staining. Blue is DAPI. Green is eGFP. All photos taken under identical light exposure parameters. All images are of male flies. Genotypes are indicated. Scale bar represents 200$\mu$m.

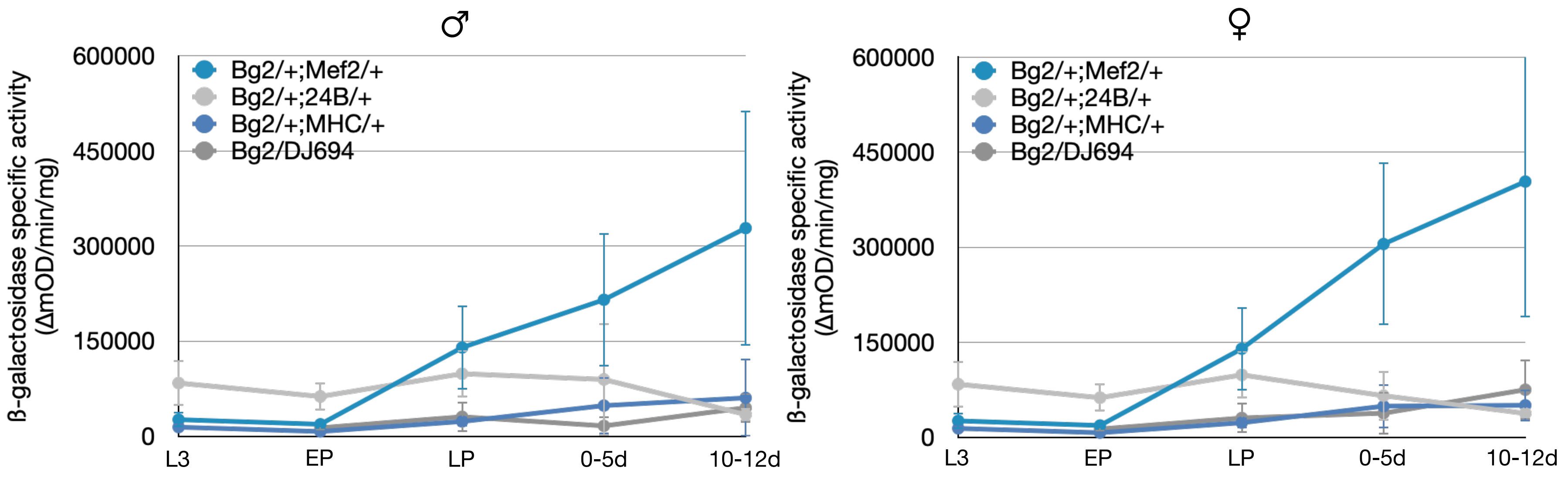

**Figure S3. Comparison of muscle driver expression pattern and level** UAS-lacZ (Bg2) quantitative reporter assay to compare expression of each driver at various time points. Y-axis represents β-galactosidase specific activity (ΔmOD/min/mg). Error bars specify ± SD.

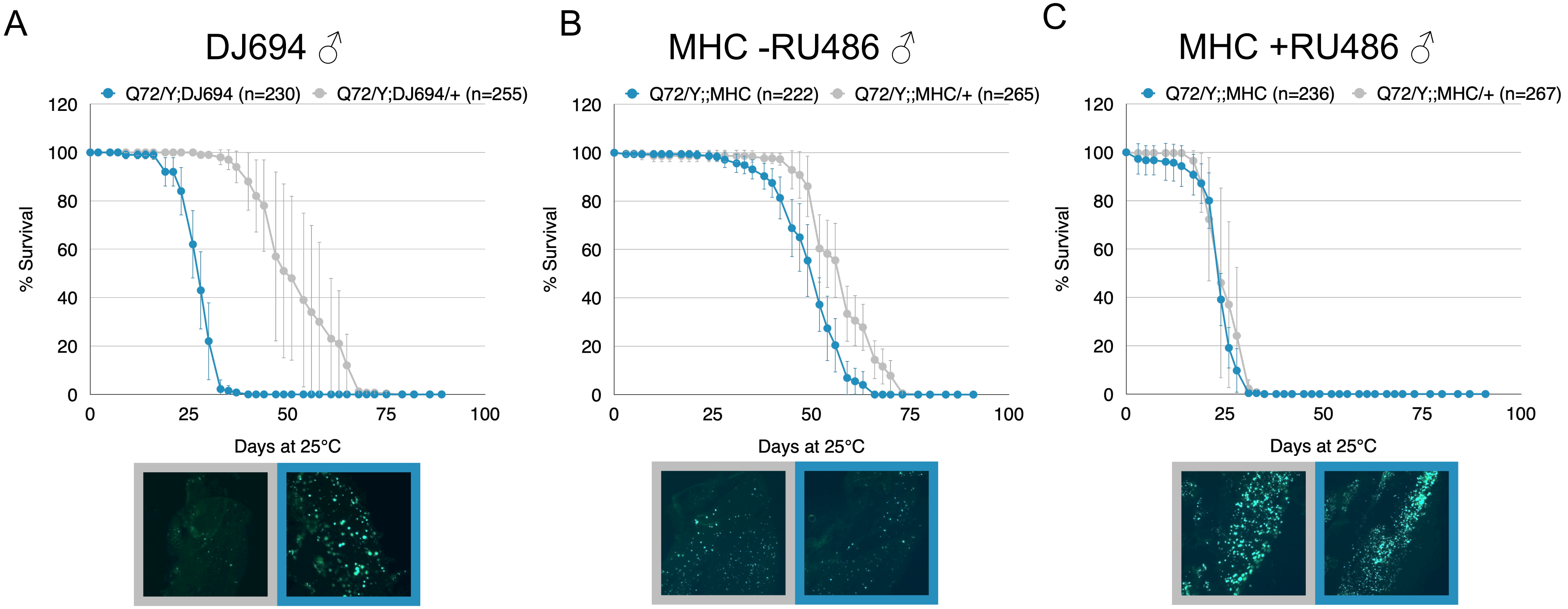

**Figure S4. Increased expression level aggravates longevity and amount of aggregates** Effect of increasing copy number with A) DJ694 driver and B) MHC driver. C) Effect of increasing copy number and treating with RU486 (50μg/ml). Graphs depict survival data for males. Since Q72 resides on the X chromosome, in males only copies of GAL4 can be increased. Flies were maintained at 25°C. Error bars represent ± SD. Bottom images show representative images of muscle aggregates. Panels of aggregate images are outlined in a colour corresponding to the legend in the survival graph.

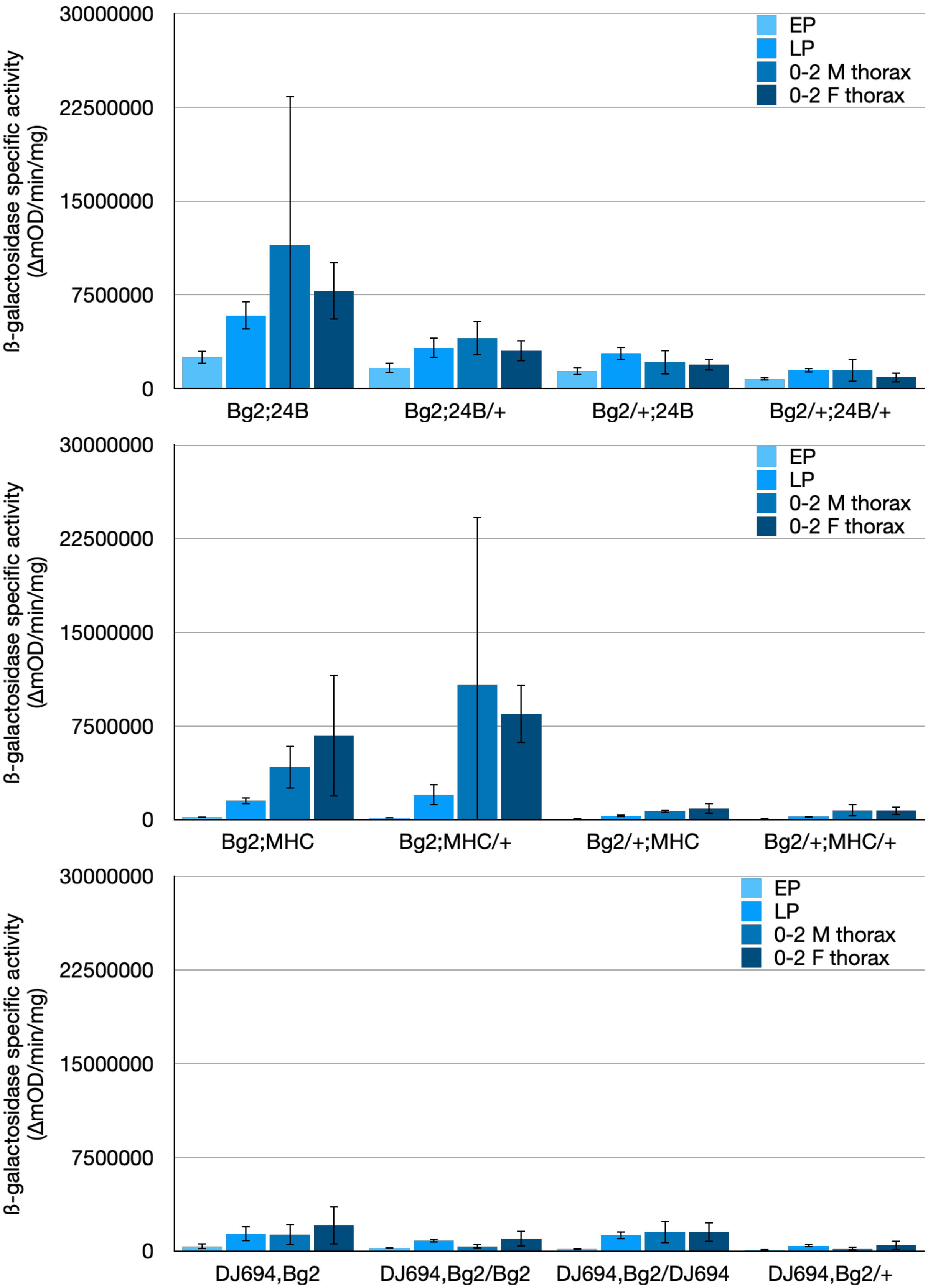

**Figure S5. Effect of increasing UAS and GAL4 copy number of expression level**

Recombinant driver and UAS-lacZ (Bg2) lines were used to generate genotypes with multiple copies of UAS and GAL4 transgenes. Muscle drivers 24B, MHC, and DJ694 were used. Flies were raised at 25°C on standard fly food. Graphs depict the expression level as measured by UAS-lacZ (Bg2) quantitative reporter assay. Early pupae (EP), late pupae (LP), 0-2d old adult thoraces were measured. Y-axis depicts the ß-galactosidase specific activity (ΔmOD/min/mg). Bars represent the average of two independent replicates with 5 individuals in each (10 total). Error bars denote ± SD.

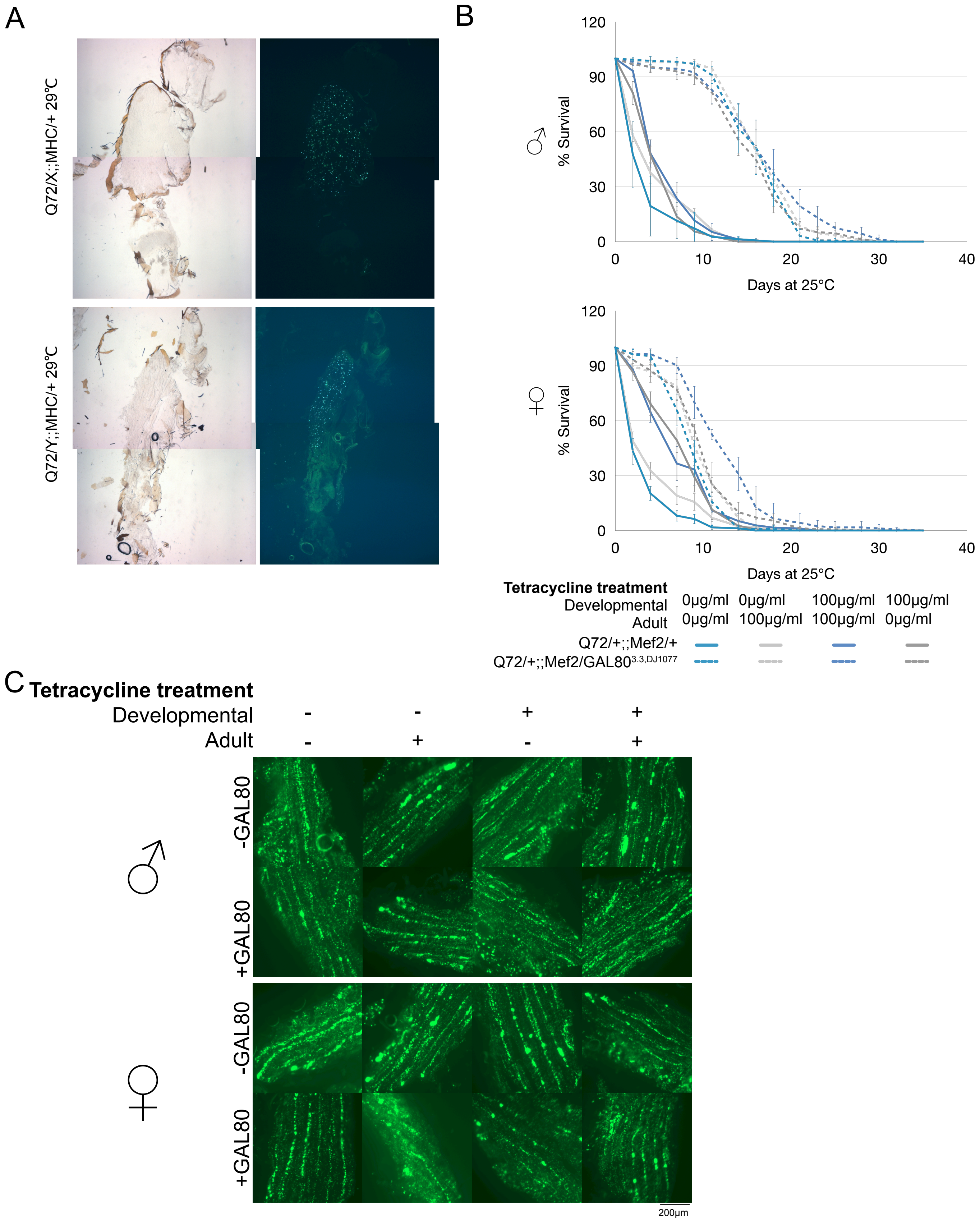

**Figure S6. Temporal repression with Tet-off GAL80 transgenes does not eliminate striation.** A) Cryosectioned male and female Q72/+;;MHC/+ flies raised at 29°C still display dispersed aggregates. B) Using Tet-off GAL80 transgenes (GAL80$^{3.3,DJ1077}$) to temporally repress Q72 expression with the Mef2 driver. Animals were treated with different tetracycline regimens to repress expression (0$\mu$g/ml tetracycline) or induce expression (100$\mu$g/ml tetracycline). Animals were either treated during development (dark grey) or adulthood (light grey), both (blue), or neither (teal). Graphs show the effect on survival of male and female flies. Solid lines represent the control genotype that does not carry the Tet-off GAL80 transgenes (Q72/+;;Mef2/+). Dashed lines represent experimental genotype that carries the Tet-off GAL80 transgenes (Q72/+;;Mef2/GAL80$^{3.3,DJ1077}$). Error bars denote ± SD. C) Cryosectioned flies from B) show striated aggregates present in all genotypes and treatments. Genotype abbreviations: *-GAL80:* Q72/+;Mef2/+ ; *+GAL80:* Q72/+;;Mef2/GAL80$^{3.3,DJ1077}$

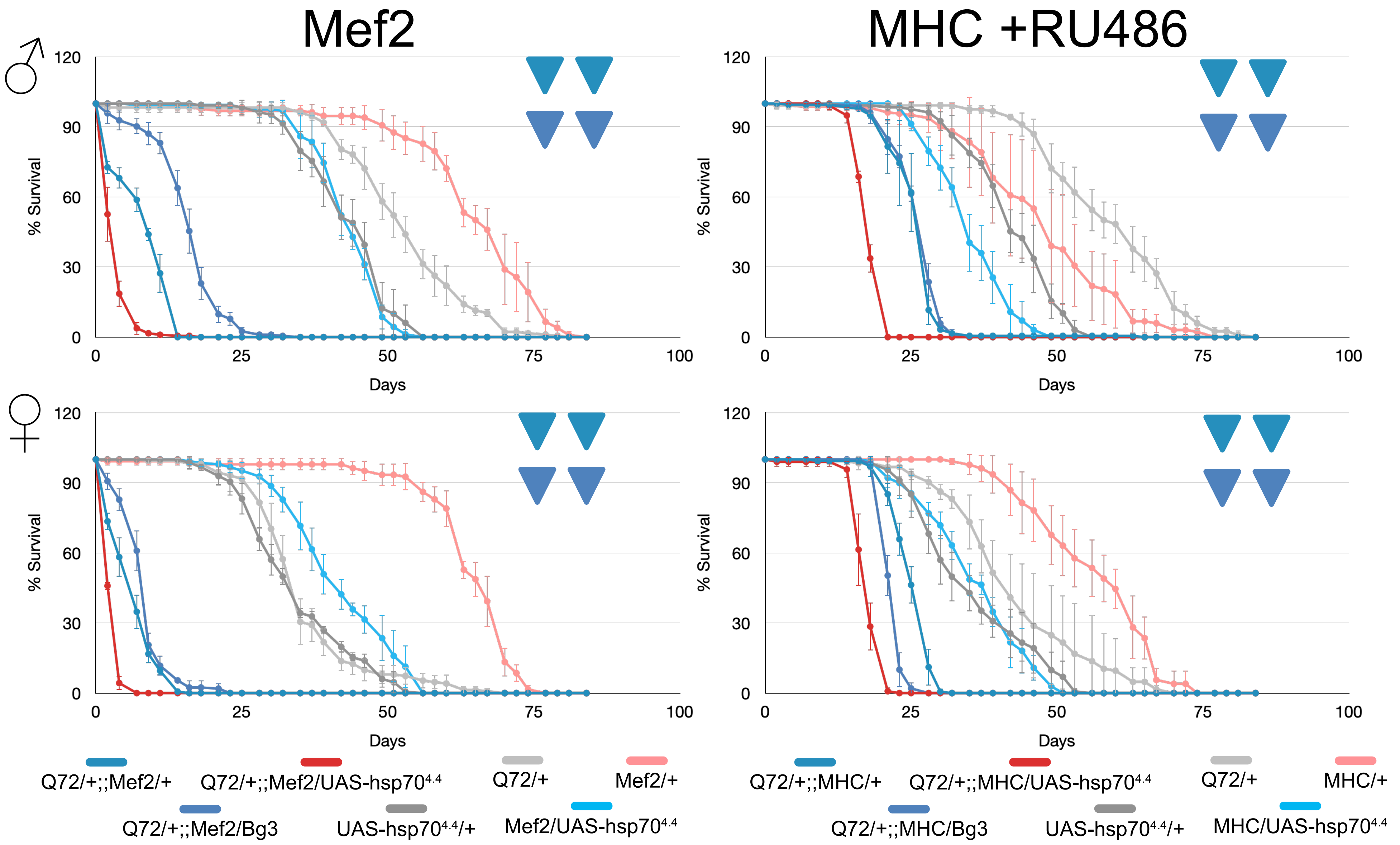

**Figure S7. Hsp70 does not suppress the detrimental phenotype caused by polyglutamine in muscle** Survival curves show the effect of expressing UAS-hsp70$^{4.4}$ in the background of flies expressing Q72 in the muscle driven by Mef2 and MHC drivers. MHC genotypes were treated with RU486 (50$\mu$g/ml), Mef2 is maintained on standard fly food. Error bars denote ± SD.Triangles represent the result of the logrank comparision of Q72/+;;Mef2/UAS-hsp70$^{4.4}$ curves to controls from two independent replicates. Teal triangles compare to Q72/+;;Mef2/+. Blue triangles compare to Q72/+;;Mef2/Bg3. ▲: significant increase, △: insignificant increase, ▼: significant decrease, ▽: insignificant decrease,

▶: no change

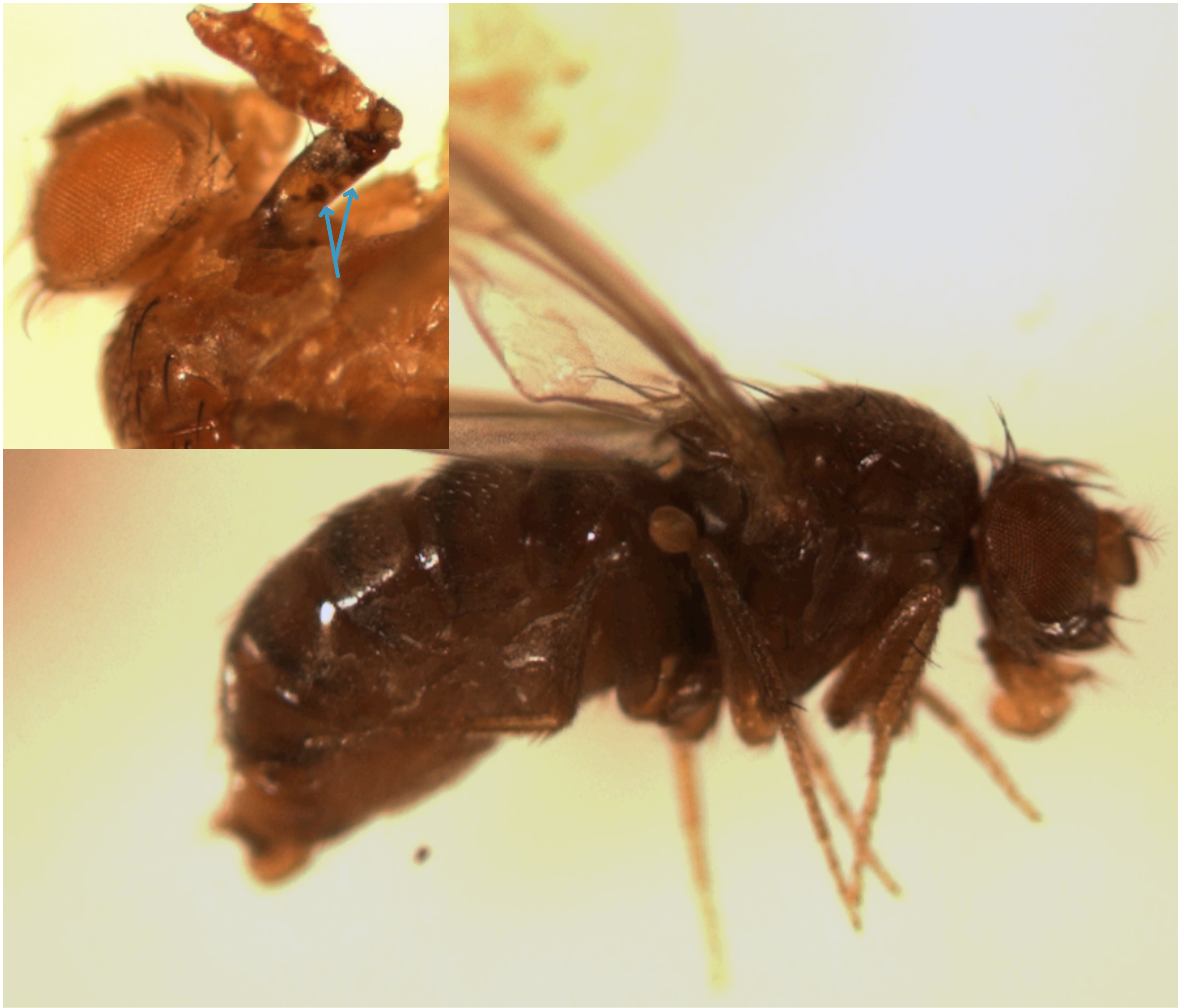

**Figure S8. Flies with black necrotic tissue** When flies that expressed Q72 were scored as dead they were notably black entirely or in parts of the fly (see inset with black lesions throughout the leg).

Table 1 Expression of Q72 in muscle longevity dataset (related to Figure 1)

| Replicate | Genotype | Sex | Treatment | n | Mean | Median | Maximum |
|---|---|---|---|---|---|---|---|
| 1 | MHC/+ | male | -RU486 | 221 | 57.74 | 63.00 | 71.88 |
| 2 | MHC/+ | male | -RU486 | 219 | 59.65 | 62.00 | 76.50 |
| 3 | MHC/+ | male | -RU486 | 156 | 46.89 | 46.00 | 62.25 |
| 4 | MHC/+ | male | -RU486 | 160 | 51.64 | 53.00 | 69.50 |
| 1 | MHC/Q25 | male | -RU486 | 176 | 54.04 | 56.00 | 70.88 |
| 2 | MHC/Q25 | male | -RU486 | 194 | 53.46 | 55.00 | 66.38 |
| 3 | MHC/Q25 | male | -RU486 | 121 | 45.36 | 46.00 | 59.25 |
| 4 | MHC/Q25 | male | -RU486 | 122 | 45.77 | 46.00 | 56.75 |
| 3 | Q25/+ | male | -RU486 | 100 | 65.36 | 66.50 | 80.67 |
| 4 | Q25/+ | male | -RU486 | 124 | 57.01 | 53.00 | 78.00 |
| 3 | Q72/+ | male | -RU486 | 76 | 56.76 | 56.00 | 75.33 |
| 4 | Q72/+ | male | -RU486 | 84 | 58.72 | 55.00 | 77.67 |
| 1 | Q72/+;;MHC/+ | male | -RU486 | 329 | 50.84 | 49.00 | 65.30 |
| 2 | Q72/+;;MHC/+ | male | -RU486 | 201 | 52.97 | 55.00 | 64.13 |
| 3 | Q72/+;;MHC/+ | male | -RU486 | 123 | 43.29 | 42.00 | 55.25 |
| 4 | Q72/+;;MHC/+ | male | -RU486 | 153 | 44.86 | 46.00 | 55.75 |
| 3 | MHC/+ | male | +RU486 | 156 | 44.50 | 45.00 | 56.00 |
| 4 | MHC/+ | male | +RU486 | 146 | 60.38 | 60.00 | 77.50 |
| 3 | MHC/Q25 | male | +RU486 | 127 | 47.93 | 49.00 | 71.00 |
| 4 | MHC/Q25 | male | +RU486 | 125 | 47.29 | 48.00 | 66.00 |
| 3 | Q25/+ | male | +RU486 | 84 | 62.43 | 63.00 | 80.67 |
| 4 | Q25/+ | male | +RU486 | 114 | 55.42 | 54.00 | 74.00 |
| 3 | Q72/+ | male | +RU486 | 80 | 56.38 | 56.00 | 74.67 |
| 4 | Q72/+ | male | +RU486 | 115 | 57.84 | 60.00 | 72.75 |
| 3 | Q72/+;;MHC/+ | male | +RU486 | 134 | 27.41 | 28.00 | 31.50 |
| 4 | Q72/+;;MHC/+ | male | +RU486 | 150 | 28.14 | 29.00 | 32.50 |
| 1 | DJ694/+ | male | none | 140 | 56.36 | 56.00 | 73.88 |
| 2 | DJ694/+ | male | none | 230 | 57.85 | 55.00 | 80.25 |
| 3 | DJ694/+ | male | none | 215 | 44.26 | 46.00 | 61.25 |
| 4 | DJ694/+ | male | none | 176 | 50.03 | 50.00 | 67.00 |
| 1 | DJ694/+;Q25/+ | male | none | 152 | 52.51 | 46.00 | 81.88 |
| 2 | DJ694/+;Q25/+ | male | none | 156 | 41.01 | 34.00 | 61.86 |
| 3 | DJ694/+;Q25/+ | male | none | 120 | 56.64 | 56.00 | 79.75 |
| 4 | DJ694/+;Q25/+ | male | none | 116 | 49.28 | 46.00 | 74.25 |
| 1 | Mef2/+ | male | none | 228 | 71.74 | 70.00 | 90.25 |
| 2 | Mef2/+ | male | none | 222 | 67.11 | 69.00 | 84.00 |
| 3 | Mef2/+ | male | none | 199 | 58.09 | 63.00 | 78.00 |
| 4 | Mef2/+ | male | none | 230 | 54.85 | 60.00 | 72.00 |
| 1 | Mef2/Q25 | male | none | 170 | 69.25 | 74.00 | 84.57 |
| 2 | Mef2/Q25 | male | none | 209 | 67.21 | 73.00 | 85.38 |
| 3 | Mef2/Q25 | male | none | 98 | 64.58 | 70.00 | 83.00 |

| Replicate | Genotype | Sex | Treatment | n | Mean | Median | Maximum |
|---|---|---|---|---|---|---|---|
| 4 | Mef2/Q25 | male | none | 102 | 60.35 | 60.00 | 81.00 |
| 1 | Q25/+ | male | none | 130 | 68.72 | 75.50 | 84.29 |
| 2 | Q25/+ | male | none | 167 | 70.96 | 76.00 | 84.38 |
| 3 | Q25/+ | male | none | 101 | 62.22 | 63.00 | 83.00 |
| 4 | Q25/+ | male | none | 97 | 57.36 | 57.00 | 77.67 |
| 1 | Q72/+ | male | none | 179 | 52.29 | 51.00 | 69.63 |
| 2 | Q72/+ | male | none | 183 | 44.95 | 48.00 | 56.25 |
| 3 | Q72/+ | male | none | 89 | 57.72 | 56.00 | 76.00 |
| 4 | Q72/+ | male | none | 95 | 56.65 | 57.00 | 74.67 |
| 1 | Q72/+;;Mef2/+ | male | none | 239 | 4.42 | 2.00 | 11.75 |
| 2 | Q72/+;;Mef2/+ | male | none | 190 | 7.05 | 6.00 | 14.25 |
| 3 | Q72/+;;Mef2/+ | male | none | 144 | 10.68 | 14.00 | 19.00 |
| 4 | Q72/+;;Mef2/+ | male | none | 128 | 10.91 | 11.00 | 18.50 |
| 1 | Q72/+;DJ694/+ | male | none | 252 | 48.29 | 49.00 | 56.50 |
| 2 | Q72/+;DJ694/+ | male | none | 216 | 46.53 | 45.00 | 57.50 |
| 3 | Q72/+;DJ694/+ | male | none | 128 | 41.58 | 42.00 | 46.00 |
| 4 | Q72/+;DJ694/+ | male | none | 139 | 46.08 | 46.00 | 53.00 |
| 1 | MHC/+ | female | -RU486 | 215 | 54.16 | 56.00 | 70.75 |
| 2 | MHC/+ | female | -RU486 | 275 | 48.80 | 48.00 | 65.88 |
| 3 | MHC/+ | female | -RU486 | 183 | 53.92 | 53.00 | 68.25 |
| 4 | MHC/+ | female | -RU486 | 160 | 60.21 | 60.00 | 76.25 |
| 1 | MHC/Q25 | female | -RU486 | 183 | 68.64 | 70.00 | 79.63 |
| 2 | MHC/Q25 | female | -RU486 | 228 | 66.12 | 69.00 | 78.00 |
| 3 | MHC/Q25 | female | -RU486 | 110 | 77.65 | 77.00 | 84.00 |
| 4 | MHC/Q25 | female | -RU486 | 160 | 72.73 | 74.00 | 79.50 |
| 3 | Q25/+ | female | -RU486 | 120 | 67.76 | 72.00 | 79.75 |
| 4 | Q25/+ | female | -RU486 | 160 | 57.46 | 63.00 | 76.25 |
| 3 | Q72/+ | female | -RU486 | 124 | 47.43 | 49.00 | 64.75 |
| 4 | Q72/+ | female | -RU486 | 135 | 46.13 | 46.00 | 66.25 |
| 1 | Q72/+;;MHC/+ | female | -RU486 | 363 | 52.01 | 51.00 | 69.58 |
| 2 | Q72/+;;MHC/+ | female | -RU486 | 249 | 54.81 | 55.00 | 70.00 |
| 3 | Q72/+;;MHC/+ | female | -RU486 | 139 | 52.31 | 53.00 | 67.75 |
| 4 | Q72/+;;MHC/+ | female | -RU486 | 175 | 48.44 | 46.00 | 64.00 |
| 3 | MHC/+ | female | +RU486 | 165 | 56.84 | 56.00 | 74.00 |
| 4 | MHC/+ | female | +RU486 | 144 | 64.42 | 67.00 | 76.00 |
| 3 | MHC/Q25 | female | +RU486 | 120 | 73.29 | 74.00 | 77.00 |
| 4 | MHC/Q25 | female | +RU486 | 151 | 65.60 | 67.00 | 74.00 |
| 3 | Q25/+ | female | +RU486 | 130 | 69.09 | 74.00 | 81.00 |
| 4 | Q25/+ | female | +RU486 | 151 | 58.68 | 60.00 | 79.75 |
| 3 | Q72/+ | female | +RU486 | 131 | 48.16 | 49.00 | 64.75 |
| 4 | Q72/+ | female | +RU486 | 137 | 46.70 | 46.00 | 69.25 |

| Replicate | Genotype | Sex | Treatment | n | Mean | Median | Maximum |
|---|---|---|---|---|---|---|---|
| 3 | Q72/+;;MHC/+ | female | +RU486 | 123 | 24.49 | 25.00 | 30.50 |
| 4 | Q72/+;;MHC/+ | female | +RU486 | 135 | 24.68 | 25.00 | 28.50 |
| 1 | DJ694/+ | female | none | 239 | 55.49 | 56.00 | 72.00 |
| 2 | DJ694/+ | female | none | 271 | 48.67 | 50.00 | 67.88 |
| 3 | DJ694/+ | female | none | 203 | 46.83 | 49.00 | 57.75 |
| 4 | DJ694/+ | female | none | 162 | 48.92 | 48.00 | 63.50 |
| 1 | DJ694/+; Q25/+ | female | none | 180 | 64.03 | 68.50 | 80.38 |
| 2 | DJ694/+; Q25/+ | female | none | 171 | 47.00 | 43.00 | 57.71 |
| 3 | DJ694/+;Q25/+ | female | none | 132 | 63.94 | 66.00 | 80.00 |
| 4 | DJ694/+;Q25/+ | female | none | 130 | 61.71 | 67.00 | 80.25 |
| 1 | Mef2/+ | female | none | 237 | 55.14 | 58.00 | 72.88 |
| 2 | Mef2/+ | female | none | 250 | 48.50 | 41.00 | 60.25 |
| 3 | Mef2/+ | female | none | 203 | 64.88 | 66.00 | 78.50 |
| 4 | Mef2/+ | female | none | 218 | 61.02 | 60.00 | 74.25 |
| 1 | Mef2/Q25 | female | none | 175 | 73.47 | 74.00 | 84.50 |
| 2 | Mef2/Q25 | female | none | 229 | 70.24 | 73.00 | 81.63 |
| 3 | Mef2/Q25 | female | none | 119 | 71.52 | 74.00 | 81.25 |
| 4 | Mef2/Q25 | female | none | 131 | 72.74 | 74.00 | 81.00 |
| 1 | Q25/+ | female | none | 169 | 70.09 | 77.00 | 82.38 |
| 2 | Q25/+ | female | none | 217 | 70.67 | 76.00 | 80.63 |
| 3 | Q25/+ | female | none | 98 | 68.02 | 72.00 | 82.00 |
| 4 | Q25/+ | female | none | 108 | 57.48 | 60.00 | 78.67 |
| 1 | Q72/+ | female | none | 200 | 46.89 | 42.00 | 69.50 |
| 2 | Q72/+ | female | none | 250 | 44.50 | 41.00 | 65.13 |
| 3 | Q72/+ | female | none | 96 | 47.56 | 44.00 | 66.33 |
| 4 | Q72/+ | female | none | 148 | 45.78 | 43.00 | 61.50 |
| 1 | Q72/+;;Mef2/+ | female | none | 247 | 5.29 | 4.00 | 10.00 |
| 2 | Q72/+;;Mef2/+ | female | none | 231 | 7.36 | 6.00 | 14.25 |
| 3 | Q72/+;;Mef2/+ | female | none | 151 | 7.78 | 7.00 | 17.25 |
| 4 | Q72/+;;Mef2/+ | female | none | 120 | 9.48 | 8.00 | 19.33 |
| 1 | Q72/+;DJ694/+ | female | none | 371 | 43.76 | 42.00 | 57.44 |
| 2 | Q72/+;DJ694/+ | female | none | 232 | 47.01 | 50.00 | 57.63 |
| 3 | Q72/+;DJ694/+ | female | none | 154 | 42.78 | 42.00 | 49.00 |
| 4 | Q72/+;DJ694/+ | female | none | 158 | 44.88 | 46.00 | 53.00 |

Table 2 Expression level of muscle drivers as measured by CPRG assay (related to Figures 3, 5, S3, S5)

| Rep. | Genotype | Stage | Sex | Treatmen | °C | ß-gal ΔmOD/min/ml | Protein mg/ml | Specific Activity ΔmOD/min/mg | Whole/ dissected |
|---|---|---|---|---|---|---|---|---|---|
| 5 | Bg2;24B | 0-1d | F | none | 25 | 53840.00 | 0.09 | 610814.37 | thorax |
| 5 | Bg2;24B | 0-1d | F | none | 25 | 58880.00 | 0.05 | 1147999.13 | thorax |
| 5 | Bg2;24B | 0-1d | F | none | 25 | 58560.00 | 0.08 | 737009.10 | thorax |
| 5 | Bg2;24B | 0-1d | F | none | 25 | 53720.00 | 0.06 | 908402.65 | thorax |
| 5 | Bg2;24B | 0-1d | F | none | 25 | 44120.00 | 0.05 | 965767.85 | thorax |
| 4 | Bg2;24B | 1-2d | F | none | 25 | 43320.00 | 0.08 | 565867.50 | thorax |
| 4 | Bg2;24B | 1-2d | F | none | 25 | 37720.00 | 0.04 | 878654.12 | thorax |
| 4 | Bg2;24B | 1-2d | F | none | 25 | 49960.00 | 0.09 | 553606.83 | thorax |
| 4 | Bg2;24B | 1-2d | F | none | 25 | 106760.00 | 0.11 | 990458.99 | thorax |
| 4 | Bg2;24B | 1-2d | F | none | 25 | 44120.00 | 0.10 | 454116.11 | thorax |
| 5 | Bg2;24B | 0-1d | M | none | 25 | 43040.00 | 0.06 | 667681.39 | thorax |
| 5 | Bg2;24B | 0-1d | M | none | 25 | 50000.00 | 0.05 | 1097846.15 | thorax |
| 5 | Bg2;24B | 0-1d | M | none | 25 | 40600.00 | 0.01 | 4260611.76 | thorax |
| 5 | Bg2;24B | 0-1d | M | none | 25 | 34560.00 | 0.04 | 768287.10 | thorax |
| 5 | Bg2;24B | 0-1d | M | none | 25 | 44960.00 | 0.02 | 1980460.25 | thorax |
| 4 | Bg2;24B | 1-2d | M | none | 25 | 37800.00 | 0.09 | 407460.17 | thorax |
| 4 | Bg2;24B | 1-2d | M | none | 25 | 35440.00 | 0.04 | 942228.13 | thorax |
| 4 | Bg2;24B | 1-2d | M | none | 25 | 34480.00 | 0.07 | 526222.15 | thorax |
| 4 | Bg2;24B | 1-2d | M | none | 25 | 30040.00 | 0.09 | 325210.01 | thorax |
| 4 | Bg2;24B | 1-2d | M | none | 25 | 36720.00 | 0.07 | 551459.64 | thorax |
| 4 | Bg2;24B | EP | ND | none | 25 | 93240.00 | 0.29 | 316232.47 | whole |
| 4 | Bg2;24B | EP | ND | none | 25 | 150360.00 | 0.49 | 307189.55 | whole |
| 4 | Bg2;24B | EP | ND | none | 25 | 72840.00 | 0.32 | 228424.43 | whole |
| 4 | Bg2;24B | EP | ND | none | 25 | 60480.00 | 0.33 | 183600.81 | whole |
| 4 | Bg2;24B | EP | ND | none | 25 | 41080.00 | 0.22 | 185519.65 | whole |
| 5 | Bg2;24B | EP | ND | none | 25 | 127040.00 | 0.48 | 263207.02 | whole |
| 5 | Bg2;24B | EP | ND | none | 25 | 113920.00 | 0.38 | 300669.07 | whole |
| 5 | Bg2;24B | EP | ND | none | 25 | 116960.00 | 0.41 | 285716.05 | whole |
| 5 | Bg2;24B | EP | ND | none | 25 | 101520.00 | 0.52 | 196950.02 | whole |
| 5 | Bg2;24B | EP | ND | none | 25 | 68280.00 | 0.29 | 235630.59 | whole |
| 4 | Bg2;24B | LP | ND | none | 25 | 249160.00 | 0.36 | 696982.09 | whole |
| 4 | Bg2;24B | LP | ND | none | 25 | 241400.00 | 0.34 | 719901.23 | whole |
| 4 | Bg2;24B | LP | ND | none | 25 | 183680.00 | 0.32 | 576995.61 | whole |
| 4 | Bg2;24B | LP | ND | none | 25 | 263160.00 | 0.39 | 667960.18 | whole |
| 4 | Bg2;24B | LP | ND | none | 25 | 185160.00 | 0.26 | 708307.49 | whole |
| 5 | Bg2;24B | LP | ND | none | 25 | 162720.00 | 0.34 | 477843.46 | whole |
| 5 | Bg2;24B | LP | ND | none | 25 | 223360.00 | 0.40 | 555639.52 | whole |
| 5 | Bg2;24B | LP | ND | none | 25 | 112320.00 | 0.20 | 563025.21 | whole |

| Rep. | Genotype | Stage | Sex | Treatmen | °C | ß-gal ΔmOD/min/ ml | Protein mg/ml | Specific Activity ΔmOD/min/ mg | Whole/ dissected |
|---|---|---|---|---|---|---|---|---|---|
| 5 | Bg2;24B | LP | ND | none | 25 | 165280.00 | 0.32 | 517420.34 | whole |
| 5 | Bg2;24B | LP | ND | none | 25 | 78280.00 | 0.21 | 374948.55 | whole |
| 5 | Bg2;24B/+ | 0-1d | F | none | 25 | 37240.00 | 0.10 | 378015.14 | thorax |
| 5 | Bg2;24B/+ | 0-1d | F | none | 25 | 23640.00 | 0.11 | 219084.47 | thorax |
| 5 | Bg2;24B/+ | 0-1d | F | none | 25 | 34160.00 | 0.18 | 191188.83 | thorax |
| 5 | Bg2;24B/+ | 0-1d | F | none | 25 | 33320.00 | 0.07 | 454630.06 | thorax |
| 5 | Bg2;24B/+ | 0-1d | F | none | 25 | 24840.00 | 0.11 | 219107.84 | thorax |
| 4 | Bg2;24B/+ | 1-2d | F | none | 25 | 27520.00 | 0.08 | 336683.71 | thorax |
| 4 | Bg2;24B/+ | 1-2d | F | none | 25 | 20520.00 | 0.08 | 263468.40 | thorax |
| 4 | Bg2;24B/+ | 1-2d | F | none | 25 | 33680.00 | 0.09 | 389260.09 | thorax |
| 4 | Bg2;24B/+ | 1-2d | F | none | 25 | 28080.00 | 0.11 | 258597.21 | thorax |
| 4 | Bg2;24B/+ | 1-2d | F | none | 25 | 27240.00 | 0.08 | 333258.15 | thorax |
| 5 | Bg2;24B/+ | 0-1d | M | none | 25 | 16840.00 | 0.07 | 233794.24 | thorax |
| 5 | Bg2;24B/+ | 0-1d | M | none | 25 | 20760.00 | 0.04 | 550718.81 | thorax |
| 5 | Bg2;24B/+ | 0-1d | M | none | 25 | 33120.00 | 0.06 | 558733.62 | thorax |
| 5 | Bg2;24B/+ | 0-1d | M | none | 25 | 18080.00 | 0.10 | 176980.63 | thorax |
| 5 | Bg2;24B/+ | 0-1d | M | none | 25 | 21360.00 | 0.06 | 357805.07 | thorax |
| 4 | Bg2;24B/+ | 1-2d | M | none | 25 | 32880.00 | 0.08 | 428008.86 | thorax |
| 4 | Bg2;24B/+ | 1-2d | M | none | 25 | 28280.00 | 0.07 | 391857.68 | thorax |
| 4 | Bg2;24B/+ | 1-2d | M | none | 25 | 17160.00 | 0.05 | 314140.73 | thorax |
| 4 | Bg2;24B/+ | 1-2d | M | none | 25 | 55680.00 | 0.10 | 554882.54 | thorax |
| 4 | Bg2;24B/+ | 1-2d | M | none | 25 | 27200.00 | 0.06 | 483812.77 | thorax |
| 4 | Bg2;24B/+ | EP | ND | none | 25 | 28600.00 | 0.26 | 110195.32 | whole |
| 4 | Bg2;24B/+ | EP | ND | none | 25 | 23200.00 | 0.19 | 122076.90 | whole |
| 4 | Bg2;24B/+ | EP | ND | none | 25 | 105760.00 | 0.43 | 244191.85 | whole |
| 4 | Bg2;24B/+ | EP | ND | none | 25 | 47200.00 | 0.32 | 146032.35 | whole |
| 4 | Bg2;24B/+ | EP | ND | none | 25 | 74680.00 | 0.47 | 157229.25 | whole |
| 5 | Bg2;24B/+ | EP | ND | none | 25 | 91320.00 | 0.56 | 164391.25 | whole |
| 5 | Bg2;24B/+ | EP | ND | none | 25 | 60600.00 | 0.34 | 177899.10 | whole |
| 5 | Bg2;24B/+ | EP | ND | none | 25 | 104720.00 | 0.46 | 229640.39 | whole |
| 5 | Bg2;24B/+ | EP | ND | none | 25 | 44160.00 | 0.24 | 184417.17 | whole |
| 5 | Bg2;24B/+ | EP | ND | none | 25 | 48840.00 | 0.34 | 142152.49 | whole |
| 4 | Bg2;24B/+ | LP | ND | none | 25 | 118400.00 | 0.33 | 356762.26 | whole |
| 4 | Bg2;24B/+ | LP | ND | none | 25 | 101880.00 | 0.25 | 406226.29 | whole |
| 4 | Bg2;24B/+ | LP | ND | none | 25 | 92440.00 | 0.24 | 379862.73 | whole |
| 4 | Bg2;24B/+ | LP | ND | none | 25 | 161840.00 | 0.43 | 379237.01 | whole |
| 4 | Bg2;24B/+ | LP | ND | none | 25 | 132840.00 | 0.34 | 395841.14 | whole |
| 5 | Bg2;24B/+ | LP | ND | none | 25 | 35440.00 | 0.20 | 179822.78 | whole |
| 5 | Bg2;24B/+ | LP | ND | none | 25 | 40160.00 | 0.19 | 207580.91 | whole |
| 5 | Bg2;24B/+ | LP | ND | none | 25 | 147320.00 | 0.40 | 363814.80 | whole |

| Rep. | Genotype | Stage | Sex | Treatmen | °C | ß-gal ΔmOD/min/ ml | Protein mg/ml | Specific Activity ΔmOD/min/ mg | Whole/ dissected |
|---|---|---|---|---|---|---|---|---|---|
| 5 | Bg2;24B/+ | LP | ND | none | 25 | 52240.00 | 0.20 | 262656.39 | whole |
| 5 | Bg2;24B/+ | LP | ND | none | 25 | 81480.00 | 0.24 | 333641.70 | whole |
| 6 | Bg2;MHC | 10-11d | F | -RU486 | 25 | 39635.00 | 0.12 | 339748.44 | thorax |
| 6 | Bg2;MHC | 10-11d | F | -RU486 | 25 | 20580.00 | 0.07 | 291038.73 | thorax |
| 6 | Bg2;MHC | 10-11d | F | -RU486 | 25 | 39815.00 | 0.09 | 420164.34 | thorax |
| 6 | Bg2;MHC | 10-11d | F | -RU486 | 25 | 31130.00 | 0.09 | 362145.67 | thorax |
| 6 | Bg2;MHC | 10-11d | F | -RU486 | 25 | 39475.00 | 0.10 | 378557.11 | thorax |
| 7 | Bg2;MHC | 10-12d | F | -RU486 | 25 | 23400.00 | 0.13 | 179138.35 | thorax |
| 7 | Bg2;MHC | 10-12d | F | -RU486 | 25 | 62680.00 | 0.10 | 617584.50 | thorax |
| 7 | Bg2;MHC | 10-12d | F | -RU486 | 25 | 33900.00 | 0.14 | 235017.73 | thorax |
| 7 | Bg2;MHC | 10-12d | F | -RU486 | 25 | 21940.00 | 0.12 | 190011.65 | thorax |
| 7 | Bg2;MHC | 10-12d | F | -RU486 | 25 | 31240.00 | 0.11 | 285796.92 | thorax |
| 6 | Bg2;MHC | 10-11d | F | +RU486 | 25 | 45620.00 | 0.10 | 455826.83 | thorax |
| 6 | Bg2;MHC | 10-11d | F | +RU486 | 25 | 47740.00 | 0.07 | 641699.15 | thorax |
| 6 | Bg2;MHC | 10-11d | F | +RU486 | 25 | 47415.00 | 0.11 | 451158.11 | thorax |
| 6 | Bg2;MHC | 10-11d | F | +RU486 | 25 | 51650.00 | 0.10 | 533534.67 | thorax |
| 6 | Bg2;MHC | 10-11d | F | +RU486 | 25 | 36960.00 | 0.09 | 434102.31 | thorax |
| 7 | Bg2;MHC | 10-12d | F | +RU486 | 25 | 64820.00 | 0.14 | 464240.95 | thorax |
| 7 | Bg2;MHC | 10-12d | F | +RU486 | 25 | 47620.00 | 0.13 | 376854.06 | thorax |
| 7 | Bg2;MHC | 10-12d | F | +RU486 | 25 | 51300.00 | 0.15 | 352463.14 | thorax |
| 7 | Bg2;MHC | 10-12d | F | +RU486 | 25 | 46860.00 | 0.14 | 343477.29 | thorax |
| 7 | Bg2;MHC | 10-12d | F | +RU486 | 25 | 52200.00 | 0.19 | 273774.41 | thorax |
| 5 | Bg2;MHC | 0-1d | F | none | 25 | 43840.00 | 0.14 | 316641.94 | thorax |
| 5 | Bg2;MHC | 0-1d | F | none | 25 | 38120.00 | 0.15 | 255902.46 | thorax |
| 5 | Bg2;MHC | 0-1d | F | none | 25 | 41040.00 | 0.13 | 326489.90 | thorax |
| 5 | Bg2;MHC | 0-1d | F | none | 25 | 89520.00 | 0.14 | 621415.10 | thorax |
| 5 | Bg2;MHC | 0-1d | F | none | 25 | 55000.00 | 0.11 | 520530.50 | thorax |
| 4 | Bg2;MHC | 1-2d | F | none | 25 | 45600.00 | 0.10 | 475859.08 | thorax |
| 4 | Bg2;MHC | 1-2d | F | none | 25 | 63640.00 | 0.13 | 473617.57 | thorax |
| 4 | Bg2;MHC | 1-2d | F | none | 25 | 55320.00 | 0.03 | 1866491.84 | thorax |
| 4 | Bg2;MHC | 1-2d | F | none | 25 | 45160.00 | 0.06 | 773995.08 | thorax |
| 4 | Bg2;MHC | 1-2d | F | none | 25 | 76560.00 | 0.07 | 1084816.27 | thorax |
| 6 | Bg2;MHC | 10-11d | M | -RU486 | 25 | 26630.00 | 0.08 | 321269.58 | thorax |
| 6 | Bg2;MHC | 10-11d | M | -RU486 | 25 | 23115.00 | 0.21 | 111766.34 | thorax |
| 6 | Bg2;MHC | 10-11d | M | -RU486 | 25 | 20985.00 | 0.09 | 243835.22 | thorax |
| 6 | Bg2;MHC | 10-11d | M | -RU486 | 25 | 25455.00 | 0.09 | 285586.98 | thorax |
| 6 | Bg2;MHC | 10-11d | M | -RU486 | 25 | 8430.00 | 0.06 | 144269.63 | thorax |
| 7 | Bg2;MHC | 10-12d | M | -RU486 | 25 | 24540.00 | 0.10 | 254877.93 | thorax |
| 7 | Bg2;MHC | 10-12d | M | -RU486 | 25 | 17940.00 | 0.09 | 199322.84 | thorax |
| 7 | Bg2;MHC | 10-12d | M | -RU486 | 25 | 15800.00 | 0.10 | 157701.18 | thorax |

| Rep. | Genotype | Stage | Sex | Treatmen | °C | ß-gal ΔmOD/min/ ml | Protein mg/ml | Specific Activity ΔmOD/min/ mg | Whole/ dissected |
|---|---|---|---|---|---|---|---|---|---|
| 7 | Bg2;MHC | 10-12d | M | -RU486 | 25 | 14540.00 | 0.10 | 144612.20 | thorax |
| 7 | Bg2;MHC | 10-12d | M | -RU486 | 25 | 17580.00 | 0.10 | 176091.96 | thorax |
| 6 | Bg2;MHC | 10-11d | M | +RU486 | 25 | 53950.00 | 0.07 | 798786.97 | thorax |
| 6 | Bg2;MHC | 10-11d | M | +RU486 | 25 | 61625.00 | 0.11 | 569725.17 | thorax |
| 6 | Bg2;MHC | 10-11d | M | +RU486 | 25 | 50770.00 | 0.05 | 1100054.19 | thorax |
| 6 | Bg2;MHC | 10-11d | M | +RU486 | 25 | 59495.00 | 0.08 | 744411.19 | thorax |
| 6 | Bg2;MHC | 10-11d | M | +RU486 | 25 | 48745.00 | 0.06 | 751318.83 | thorax |
| 7 | Bg2;MHC | 10-12d | M | +RU486 | 25 | 54880.00 | 0.08 | 691651.82 | thorax |
| 7 | Bg2;MHC | 10-12d | M | +RU486 | 25 | 65740.00 | 0.10 | 641002.96 | thorax |
| 7 | Bg2;MHC | 10-12d | M | +RU486 | 25 | 43620.00 | 0.08 | 573718.50 | thorax |
| 7 | Bg2;MHC | 10-12d | M | +RU486 | 25 | 54680.00 | 0.12 | 445673.67 | thorax |
| 7 | Bg2;MHC | 10-12d | M | +RU486 | 25 | 46840.00 | 0.11 | 443404.66 | thorax |
| 5 | Bg2;MHC | 0-1d | M | none | 25 | 32400.00 | 0.08 | 417340.07 | thorax |
| 5 | Bg2;MHC | 0-1d | M | none | 25 | 19640.00 | 0.07 | 286607.44 | thorax |
| 5 | Bg2;MHC | 0-1d | M | none | 25 | 26760.00 | 0.05 | 498588.41 | thorax |
| 5 | Bg2;MHC | 0-1d | M | none | 25 | 20960.00 | 0.07 | 307126.41 | thorax |
| 5 | Bg2;MHC | 0-1d | M | none | 25 | 16840.00 | 0.10 | 167835.53 | thorax |
| 4 | Bg2;MHC | 1-2d | M | none | 25 | 34480.00 | 0.04 | 783768.94 | thorax |
| 4 | Bg2;MHC | 1-2d | M | none | 25 | 26320.00 | 0.07 | 384527.53 | thorax |
| 4 | Bg2;MHC | 1-2d | M | none | 25 | 27120.00 | 0.05 | 523207.38 | thorax |
| 4 | Bg2;MHC | 1-2d | M | none | 25 | 29480.00 | 0.09 | 314620.60 | thorax |
| 4 | Bg2;MHC | 1-2d | M | none | 25 | 30680.00 | 0.06 | 530658.21 | thorax |
| 4 | Bg2;MHC | EP | ND | none | 25 | 8880.00 | 0.49 | 17942.27 | whole |
| 4 | Bg2;MHC | EP | ND | none | 25 | 10480.00 | 0.45 | 23420.42 | whole |
| 4 | Bg2;MHC | EP | ND | none | 25 | 11960.00 | 0.42 | 28286.29 | whole |
| 4 | Bg2;MHC | EP | ND | none | 25 | 18280.00 | 0.62 | 29498.17 | whole |
| 4 | Bg2;MHC | EP | ND | none | 25 | 13600.00 | 0.44 | 30906.31 | whole |
| 5 | Bg2;MHC | EP | ND | none | 25 | 9360.00 | 0.54 | 17391.69 | whole |
| 5 | Bg2;MHC | EP | ND | none | 25 | 6200.00 | 0.28 | 22359.91 | whole |
| 5 | Bg2;MHC | EP | ND | none | 25 | 3400.00 | 0.33 | 10222.15 | whole |
| 5 | Bg2;MHC | EP | ND | none | 25 | 9080.00 | 0.41 | 22334.12 | whole |
| 5 | Bg2;MHC | EP | ND | none | 25 | 9640.00 | 0.42 | 22941.80 | whole |
| 4 | Bg2;MHC | LP | ND | none | 25 | 42160.00 | 0.35 | 121266.32 | whole |
| 4 | Bg2;MHC | LP | ND | none | 25 | 52840.00 | 0.30 | 175496.80 | whole |
| 4 | Bg2;MHC | LP | ND | none | 25 | 44360.00 | 0.36 | 122095.31 | whole |
| 4 | Bg2;MHC | LP | ND | none | 25 | 52880.00 | 0.38 | 137938.28 | whole |
| 4 | Bg2;MHC | LP | ND | none | 25 | 44680.00 | 0.32 | 140353.68 | whole |
| 5 | Bg2;MHC | LP | ND | none | 25 | 28560.00 | 0.21 | 134088.15 | whole |
| 5 | Bg2;MHC | LP | ND | none | 25 | 37800.00 | 0.24 | 157899.70 | whole |
| 5 | Bg2;MHC | LP | ND | none | 25 | 36960.00 | 0.24 | 157080.00 | whole |

| Rep. | Genotype | Stage | Sex | Treatmen | °C | ß-gal ΔmOD/min/ml | Protein mg/ml | Specific Activity ΔmOD/min/mg | Whole/ dissected |
|---|---|---|---|---|---|---|---|---|---|
| 5 | Bg2;MHC | LP | ND | none | 25 | 79840.00 | 0.36 | 220181.22 | whole |
| 5 | Bg2;MHC | LP | ND | none | 25 | 39160.00 | 0.27 | 144122.94 | whole |
| 5 | Bg2;MHC/+ | 0-1d | F | none | 25 | 75000.00 | 0.09 | 822119.82 | thorax |
| 5 | Bg2;MHC/+ | 0-1d | F | none | 25 | 75280.00 | 0.10 | 718179.25 | thorax |
| 5 | Bg2;MHC/+ | 0-1d | F | none | 25 | 78600.00 | 0.10 | 762078.26 | thorax |
| 5 | Bg2;MHC/+ | 0-1d | F | none | 25 | 84880.00 | 0.09 | 900005.47 | thorax |
| 5 | Bg2;MHC/+ | 0-1d | F | none | 25 | 113040.00 | 0.09 | 1298958.84 | thorax |
| 4 | Bg2;MHC/+ | 1-2d | F | none | 25 | 98320.00 | 0.08 | 1198962.20 | thorax |
| 4 | Bg2;MHC/+ | 1-2d | F | none | 25 | 98640.00 | 0.18 | 543712.35 | thorax |
| 4 | Bg2;MHC/+ | 1-2d | F | none | 25 | 116360.00 | 0.15 | 783789.29 | thorax |
| 4 | Bg2;MHC/+ | 1-2d | F | none | 25 | 91320.00 | 0.13 | 725545.60 | thorax |
| 4 | Bg2;MHC/+ | 1-2d | F | none | 25 | 82800.00 | 0.12 | 697634.04 | thorax |
| 5 | Bg2;MHC/+ | 0-1d | M | none | 25 | 77080.00 | 0.12 | 668338.86 | thorax |
| 5 | Bg2;MHC/+ | 0-1d | M | none | 25 | 67320.00 | 0.08 | 854796.30 | thorax |
| 5 | Bg2;MHC/+ | 0-1d | M | none | 25 | 44120.00 | 0.09 | 494254.82 | thorax |
| 5 | Bg2;MHC/+ | 0-1d | M | none | 25 | 60920.00 | 0.12 | 494005.82 | thorax |
| 5 | Bg2;MHC/+ | 0-1d | M | none | 25 | 76400.00 | 0.15 | 505273.77 | thorax |
| 4 | Bg2;MHC/+ | 1-2d | M | none | 25 | 49440.00 | 0.05 | 1053786.29 | thorax |
| 4 | Bg2;MHC/+ | 1-2d | M | none | 25 | 48600.00 | 0.01 | 4875552.00 | thorax |
| 4 | Bg2;MHC/+ | 1-2d | M | none | 25 | 36520.00 | 0.05 | 759051.05 | thorax |
| 4 | Bg2;MHC/+ | 1-2d | M | none | 25 | 47000.00 | 0.09 | 520807.07 | thorax |
| 4 | Bg2;MHC/+ | 1-2d | M | none | 25 | 56320.00 | 0.10 | 547482.79 | thorax |
| 4 | Bg2;MHC/+ | EP | ND | none | 25 | 8360.00 | 0.61 | 13663.41 | whole |
| 4 | Bg2;MHC/+ | EP | ND | none | 25 | 16760.00 | 0.62 | 27213.99 | whole |
| 4 | Bg2;MHC/+ | EP | ND | none | 25 | 8920.00 | 0.61 | 14638.58 | whole |
| 4 | Bg2;MHC/+ | EP | ND | none | 25 | 8520.00 | 0.57 | 14926.82 | whole |
| 4 | Bg2;MHC/+ | EP | ND | none | 25 | 8600.00 | 0.58 | 14914.30 | whole |
| 5 | Bg2;MHC/+ | EP | ND | none | 25 | 4480.00 | 0.46 | 9796.80 | whole |
| 5 | Bg2;MHC/+ | EP | ND | none | 25 | 7240.00 | 0.46 | 15872.15 | whole |
| 5 | Bg2;MHC/+ | EP | ND | none | 25 | 5880.00 | 0.50 | 11853.69 | whole |
| 5 | Bg2;MHC/+ | EP | ND | none | 25 | 6320.00 | 0.47 | 13376.71 | whole |
| 5 | Bg2;MHC/+ | EP | ND | none | 25 | 6720.00 | 0.48 | 14045.21 | whole |
| 4 | Bg2;MHC/+ | LP | ND | none | 25 | 60640.00 | 0.33 | 184118.87 | whole |
| 4 | Bg2;MHC/+ | LP | ND | none | 25 | 62920.00 | 0.33 | 189817.90 | whole |
| 4 | Bg2;MHC/+ | LP | ND | none | 25 | 134000.00 | 0.43 | 311770.30 | whole |
| 4 | Bg2;MHC/+ | LP | ND | none | 25 | 119800.00 | 0.35 | 345905.29 | whole |
| 4 | Bg2;MHC/+ | LP | ND | none | 25 | 64960.00 | 0.35 | 184314.22 | whole |
| 5 | Bg2;MHC/+ | LP | ND | none | 25 | 25600.00 | 0.29 | 88766.99 | whole |
| 5 | Bg2;MHC/+ | LP | ND | none | 25 | 59520.00 | 0.27 | 220950.38 | whole |
| 5 | Bg2;MHC/+ | LP | ND | none | 25 | 63800.00 | 0.24 | 268263.96 | whole |

| Rep. | Genotype | Stage | Sex | Treatmen | °C | ß-gal ΔmOD/min/ ml | Protein mg/ml | Specific Activity ΔmOD/min/ mg | Whole/ dissected |
|---|---|---|---|---|---|---|---|---|---|
| 5 | Bg2;MHC/+ | LP | ND | none | 25 | 37000.00 | 0.29 | 127915.18 | whole |
| 5 | Bg2;MHC/+ | LP | ND | none | 25 | 18720.00 | 0.19 | 97002.57 | whole |
| 6 | Bg2/+ | 10-11d | F | -RU486 | 25 | 0.00 | 0.08 | 0.00 | thorax |
| 6 | Bg2/+ | 10-11d | F | -RU486 | 25 | 10.00 | 0.07 | 149.42 | thorax |
| 6 | Bg2/+ | 10-11d | F | -RU486 | 25 | 2.86 | 0.07 | 38.94 | thorax |
| 6 | Bg2/+ | 10-11d | F | -RU486 | 25 | 0.00 | 0.10 | 0.00 | thorax |
| 6 | Bg2/+ | 10-11d | F | -RU486 | 25 | 5.00 | 0.08 | 63.54 | thorax |
| 7 | Bg2/+ | 10-12d | F | -RU486 | 25 | 110.00 | 0.14 | 770.82 | thorax |
| 7 | Bg2/+ | 10-12d | F | -RU486 | 25 | 57.14 | 0.18 | 323.18 | thorax |
| 7 | Bg2/+ | 10-12d | F | -RU486 | 25 | 1.43 | 0.14 | 10.58 | thorax |
| 7 | Bg2/+ | 10-12d | F | -RU486 | 25 | 25.00 | 0.13 | 189.16 | thorax |
| 7 | Bg2/+ | 10-12d | F | -RU486 | 25 | 27.27 | 0.14 | 192.87 | thorax |
| 6 | Bg2/+ | 10-11d | F | +RU486 | 25 | 0.00 | 0.08 | 0.00 | thorax |
| 6 | Bg2/+ | 10-11d | F | +RU486 | 25 | 5.00 | 0.07 | 73.03 | thorax |
| 6 | Bg2/+ | 10-11d | F | +RU486 | 25 | 6.67 | 0.07 | 91.24 | thorax |
| 6 | Bg2/+ | 10-11d | F | +RU486 | 25 | 53.33 | 0.09 | 574.61 | thorax |
| 6 | Bg2/+ | 10-11d | F | +RU486 | 25 | 0.00 | 0.08 | 0.00 | thorax |
| 7 | Bg2/+ | 10-12d | F | +RU486 | 25 | 28.00 | 0.06 | 468.18 | thorax |
| 7 | Bg2/+ | 10-12d | F | +RU486 | 25 | 15.00 | 0.13 | 112.09 | thorax |
| 7 | Bg2/+ | 10-12d | F | +RU486 | 25 | 30.00 | 0.12 | 252.56 | thorax |
| 7 | Bg2/+ | 10-12d | F | +RU486 | 25 | 13.33 | 0.11 | 122.38 | thorax |
| 7 | Bg2/+ | 10-12d | F | +RU486 | 25 | 30.00 | 0.11 | 266.37 | thorax |
| 1 | Bg2/+ | 0-24h | F | none | 25 | 3.53 | 0.19 | 18.81 | thorax |
| 1 | Bg2/+ | 0-24h | F | none | 25 | 5.99 | 0.19 | 30.83 | thorax |
| 1 | Bg2/+ | 0-24h | F | none | 25 | 0.00 | 0.21 | 0.00 | thorax |
| 1 | Bg2/+ | 0-24h | F | none | 25 | 1.50 | 0.15 | 10.28 | thorax |
| 1 | Bg2/+ | 0-24h | F | none | 25 | 2.72 | 0.21 | 12.70 | thorax |
| 8 | Bg2/+ | 0-4d | F | none | 25 | 20.00 | 0.15 | 136.04 | thorax |
| 8 | Bg2/+ | 0-4d | F | none | 25 | 0.00 | 0.14 | 0.00 | thorax |
| 8 | Bg2/+ | 0-4d | F | none | 25 | 10.00 | 0.18 | 56.68 | thorax |
| 8 | Bg2/+ | 0-4d | F | none | 25 | 5.00 | 0.14 | 36.79 | thorax |
| 8 | Bg2/+ | 0-4d | F | none | 25 | 0.00 | 0.16 | 0.00 | thorax |
| 9 | Bg2/+ | 1-3d | F | none | 25 | 0.00 | 0.11 | 0.00 | thorax |
| 9 | Bg2/+ | 1-3d | F | none | 25 | 20.00 | 0.10 | 195.54 | thorax |
| 9 | Bg2/+ | 1-3d | F | none | 25 | 0.00 | 0.11 | 0.00 | thorax |
| 9 | Bg2/+ | 1-3d | F | none | 25 | 4.00 | 0.13 | 31.95 | thorax |
| 9 | Bg2/+ | 1-3d | F | none | 25 | 20.00 | 0.15 | 130.81 | thorax |
| 1 | Bg2/+ | 10-11d | F | none | 25 | 0.00 | 0.15 | 0.00 | thorax |
| 1 | Bg2/+ | 10-11d | F | none | 25 | 0.00 | 0.23 | 0.00 | thorax |
| 1 | Bg2/+ | 10-11d | F | none | 25 | 0.00 | 0.20 | 0.00 | thorax |

| Rep. | Genotype | Stage | Sex | Treatmen | °C | ß-gal ΔmOD/min/ ml | Protein mg/ml | Specific Activity ΔmOD/min/ mg | Whole/ dissected |
|---|---|---|---|---|---|---|---|---|---|
| 1 | Bg2/+ | 10-11d | F | none | 25 | 0.00 | 0.13 | 0.00 | thorax |
| 1 | Bg2/+ | 10-11d | F | none | 25 | 1.67 | 0.29 | 5.70 | thorax |
| 1 | Bg2/+ | 3-5d | F | none | 25 | 3.00 | 0.20 | 15.02 | thorax |
| 1 | Bg2/+ | 3-5d | F | none | 25 | 1.00 | 0.18 | 5.67 | thorax |
| 1 | Bg2/+ | 3-5d | F | none | 25 | 0.00 | 0.18 | 0.00 | thorax |
| 1 | Bg2/+ | 3-5d | F | none | 25 | 0.00 | 0.14 | 0.00 | thorax |
| 1 | Bg2/+ | 3-5d | F | none | 25 | 0.00 | 0.16 | 0.00 | thorax |
| 6 | Bg2/+ | 10-11d | M | -RU486 | 25 | 3.33 | 0.04 | 77.37 | thorax |
| 6 | Bg2/+ | 10-11d | M | -RU486 | 25 | 5.00 | 0.04 | 131.34 | thorax |
| 6 | Bg2/+ | 10-11d | M | -RU486 | 25 | 0.00 | 0.05 | 0.00 | thorax |
| 6 | Bg2/+ | 10-11d | M | -RU486 | 25 | 0.00 | 0.05 | 0.00 | thorax |
| 6 | Bg2/+ | 10-11d | M | -RU486 | 25 | 4.00 | 0.05 | 74.74 | thorax |
| 7 | Bg2/+ | 10-12d | M | -RU486 | 25 | 37.78 | 0.09 | 407.40 | thorax |
| 7 | Bg2/+ | 10-12d | M | -RU486 | 25 | 62.50 | 0.10 | 624.56 | thorax |
| 7 | Bg2/+ | 10-12d | M | -RU486 | 25 | 64.00 | 0.10 | 669.66 | thorax |
| 7 | Bg2/+ | 10-12d | M | -RU486 | 25 | 13.33 | 0.10 | 136.47 | thorax |
| 7 | Bg2/+ | 10-12d | M | -RU486 | 25 | 22.50 | 0.01 | 2567.43 | thorax |
| 6 | Bg2/+ | 10-11d | M | +RU486 | 25 | 4.44 | 0.06 | 73.36 | thorax |
| 6 | Bg2/+ | 10-11d | M | +RU486 | 25 | 20.00 | 0.04 | 492.29 | thorax |
| 6 | Bg2/+ | 10-11d | M | +RU486 | 25 | 6.67 | 0.05 | 142.24 | thorax |
| 6 | Bg2/+ | 10-11d | M | +RU486 | 25 | 20.00 | 0.05 | 397.24 | thorax |
| 6 | Bg2/+ | 10-11d | M | +RU486 | 25 | 0.00 | 0.05 | 0.00 | thorax |
| 7 | Bg2/+ | 10-12d | M | +RU486 | 25 | 43.64 | 0.13 | 326.08 | thorax |
| 7 | Bg2/+ | 10-12d | M | +RU486 | 25 | 2.86 | 0.24 | 11.73 | thorax |
| 7 | Bg2/+ | 10-12d | M | +RU486 | 25 | 15.00 | 0.14 | 105.90 | thorax |
| 7 | Bg2/+ | 10-12d | M | +RU486 | 25 | 0.00 | 0.10 | 0.00 | thorax |
| 7 | Bg2/+ | 10-12d | M | +RU486 | 25 | 11.43 | 0.10 | 114.75 | thorax |
| 1 | Bg2/+ | 0-24h | M | none | 25 | 19.09 | 0.17 | 111.60 | thorax |
| 1 | Bg2/+ | 0-24h | M | none | 25 | 15.00 | 0.17 | 87.26 | thorax |
| 1 | Bg2/+ | 0-24h | M | none | 25 | 33.14 | 0.16 | 213.72 | thorax |
| 8 | Bg2/+ | 0-4d | M | none | 25 | 120.00 | 0.11 | 1123.29 | thorax |
| 8 | Bg2/+ | 0-4d | M | none | 25 | 0.00 | 0.12 | 0.00 | thorax |
| 8 | Bg2/+ | 0-4d | M | none | 25 | 0.00 | 0.11 | 0.00 | thorax |
| 8 | Bg2/+ | 0-4d | M | none | 25 | 0.00 | 0.14 | 0.00 | thorax |
| 8 | Bg2/+ | 0-4d | M | none | 25 | 33.33 | 0.14 | 233.36 | thorax |
| 9 | Bg2/+ | 1-3d | M | none | 25 | 20.00 | 0.11 | 180.44 | thorax |
| 9 | Bg2/+ | 1-3d | M | none | 25 | 15.00 | 0.08 | 181.07 | thorax |
| 9 | Bg2/+ | 1-3d | M | none | 25 | 4.00 | 0.10 | 38.95 | thorax |
| 9 | Bg2/+ | 1-3d | M | none | 25 | 8.00 | 0.08 | 97.95 | thorax |
| 9 | Bg2/+ | 1-3d | M | none | 25 | 8.00 | 0.10 | 84.12 | thorax |

| Rep. | Genotype | Stage | Sex | Treatmen | °C | ß-gal ΔmOD/min/ml | Protein mg/ml | Specific Activity ΔmOD/min/mg | Whole/ dissected |
|---|---|---|---|---|---|---|---|---|---|
| 1 | Bg2/+ | 10-11d | M | none | 25 | 0.00 | 0.15 | 0.00 | thorax |
| 1 | Bg2/+ | 10-11d | M | none | 25 | 0.00 | 0.12 | 0.00 | thorax |
| 1 | Bg2/+ | 10-11d | M | none | 25 | 1.58 | 0.15 | 10.88 | thorax |
| 1 | Bg2/+ | 10-11d | M | none | 25 | 0.00 | 0.11 | 0.00 | thorax |
| 1 | Bg2/+ | 10-11d | M | none | 25 | 0.00 | 0.13 | 0.00 | thorax |
| 1 | Bg2/+ | 3-5d | M | none | 25 | 18.76 | 0.17 | 108.20 | thorax |
| 1 | Bg2/+ | 3-5d | M | none | 25 | 6.00 | 0.14 | 41.81 | thorax |
| 1 | Bg2/+ | 3-5d | M | none | 25 | 2.00 | 0.14 | 13.84 | thorax |
| 1 | Bg2/+ | 3-5d | M | none | 25 | 0.00 | 0.13 | 0.00 | thorax |
| 1 | Bg2/+ | 3-5d | M | none | 25 | 1.07 | 0.10 | 10.82 | thorax |
| 6 | Bg2/+ | EP | ND | -RU486 | 25 | 6.67 | 0.80 | 8.30 | whole |
| 6 | Bg2/+ | EP | ND | -RU486 | 25 | 0.00 | 1.04 | 0.00 | whole |
| 6 | Bg2/+ | EP | ND | -RU486 | 25 | 0.00 | 0.90 | 0.00 | whole |
| 6 | Bg2/+ | EP | ND | -RU486 | 25 | 18.82 | 0.71 | 26.60 | whole |
| 6 | Bg2/+ | EP | ND | -RU486 | 25 | 60.00 | 0.95 | 63.02 | whole |
| 7 | Bg2/+ | EP | ND | -RU486 | 25 | 3.81 | 0.47 | 8.03 | whole |
| 7 | Bg2/+ | EP | ND | -RU486 | 25 | 12.73 | 0.91 | 14.05 | whole |
| 7 | Bg2/+ | EP | ND | -RU486 | 25 | 29.09 | 0.43 | 67.32 | whole |
| 7 | Bg2/+ | EP | ND | -RU486 | 25 | 0.00 | 0.44 | 0.00 | whole |
| 7 | Bg2/+ | EP | ND | -RU486 | 25 | 21.67 | 0.44 | 49.11 | whole |
| 6 | Bg2/+ | EP | ND | +RU486 | 25 | 17.14 | 0.44 | 39.36 | whole |
| 6 | Bg2/+ | EP | ND | +RU486 | 25 | 24.00 | 0.86 | 27.94 | whole |
| 6 | Bg2/+ | EP | ND | +RU486 | 25 | 90.00 | 0.95 | 95.20 | whole |
| 6 | Bg2/+ | EP | ND | +RU486 | 25 | 22.50 | 0.61 | 37.05 | whole |
| 6 | Bg2/+ | EP | ND | +RU486 | 25 | 4.00 | 0.65 | 6.19 | whole |
| 7 | Bg2/+ | EP | ND | +RU486 | 25 | 47.50 | 0.49 | 97.21 | whole |
| 7 | Bg2/+ | EP | ND | +RU486 | 25 | 6.67 | 0.60 | 11.16 | whole |
| 7 | Bg2/+ | EP | ND | +RU486 | 25 | 25.00 | 0.52 | 47.97 | whole |
| 7 | Bg2/+ | EP | ND | +RU486 | 25 | 0.00 | 0.43 | 0.00 | whole |
| 7 | Bg2/+ | EP | ND | +RU486 | 25 | 4.00 | 0.48 | 8.42 | whole |
| 8 | Bg2/+ | EP | ND | none | 25 | 10.00 | 0.48 | 20.78 | whole |
| 8 | Bg2/+ | EP | ND | none | 25 | 90.00 | 0.46 | 196.05 | whole |
| 8 | Bg2/+ | EP | ND | none | 25 | 20.00 | 0.41 | 48.84 | whole |
| 8 | Bg2/+ | EP | ND | none | 25 | 15.00 | 0.61 | 24.39 | whole |
| 8 | Bg2/+ | EP | ND | none | 25 | 120.00 | 0.45 | 269.37 | whole |
| 9 | Bg2/+ | EP | ND | none | 25 | 0.00 | 0.36 | 0.00 | whole |
| 9 | Bg2/+ | EP | ND | none | 25 | 2.31 | 0.50 | 4.65 | whole |
| 9 | Bg2/+ | EP | ND | none | 25 | 20.00 | 0.28 | 70.66 | whole |
| 9 | Bg2/+ | EP | ND | none | 25 | 40.00 | 0.55 | 73.39 | whole |
| 9 | Bg2/+ | EP | ND | none | 25 | 10.00 | 0.40 | 25.26 | whole |

| Rep. | Genotype | Stage | Sex | Treatmen | °C | ß-gal ΔmOD/min/ ml | Protein mg/ml | Specific Activity ΔmOD/min/ mg | Whole/ dissected |
|---|---|---|---|---|---|---|---|---|---|
| 2 | Bg2/+ | L3 | ND | none | 25 | 480.00 | 0.56 | 852.42 | whole |
| 2 | Bg2/+ | L3 | ND | none | 25 | 160.00 | 0.45 | 356.83 | whole |
| 2 | Bg2/+ | L3 | ND | none | 25 | 133.33 | 0.52 | 254.69 | whole |
| 2 | Bg2/+ | L3 | ND | none | 25 | 80.00 | 0.38 | 207.97 | whole |
| 2 | Bg2/+ | L3 | ND | none | 25 | 80.00 | 0.33 | 241.10 | whole |
| 3 | Bg2/+ | L3 | ND | none | 25 | 240.00 | 0.27 | 885.32 | whole |
| 3 | Bg2/+ | L3 | ND | none | 25 | 160.00 | 0.42 | 377.34 | whole |
| 3 | Bg2/+ | L3 | ND | none | 25 | 400.00 | 0.19 | 2160.18 | whole |
| 3 | Bg2/+ | L3 | ND | none | 25 | 40.00 | 0.30 | 131.26 | whole |
| 3 | Bg2/+ | L3 | ND | none | 25 | 400.00 | 0.48 | 834.67 | whole |
| 1 | Bg2/+ | LP | ND | none | 25 | 0.00 | 0.28 | 0.00 | whole |
| 1 | Bg2/+ | LP | ND | none | 25 | 6.00 | 0.21 | 28.15 | whole |
| 1 | Bg2/+ | LP | ND | none | 25 | 10.00 | 0.14 | 70.86 | whole |
| 1 | Bg2/+ | LP | ND | none | 25 | 0.00 | 0.24 | 0.00 | whole |
| 1 | Bg2/+ | LP | ND | none | 25 | 0.00 | 0.29 | 0.00 | whole |
| 8 | Bg2/+ | LP | ND | none | 25 | 170.00 | 0.28 | 613.13 | whole |
| 8 | Bg2/+ | LP | ND | none | 25 | 30.00 | 0.18 | 164.24 | whole |
| 8 | Bg2/+ | LP | ND | none | 25 | 90.00 | 0.29 | 314.63 | whole |
| 8 | Bg2/+ | LP | ND | none | 25 | 85.00 | 0.24 | 356.30 | whole |
| 8 | Bg2/+ | LP | ND | none | 25 | 15.00 | 0.35 | 42.79 | whole |
| 9 | Bg2/+ | LP | ND | none | 25 | 25.71 | 0.31 | 81.63 | whole |
| 9 | Bg2/+ | LP | ND | none | 25 | 15.00 | 0.44 | 33.73 | whole |
| 9 | Bg2/+ | LP | ND | none | 25 | 0.00 | 0.33 | 0.00 | whole |
| 9 | Bg2/+ | LP | ND | none | 25 | 40.00 | 0.35 | 115.26 | whole |
| 9 | Bg2/+ | LP | ND | none | 25 | 25.00 | 0.27 | 91.93 | whole |
| 5 | Bg2/+;24B | 0-1d | F | none | 25 | 18000.00 | 0.09 | 194030.21 | thorax |
| 5 | Bg2/+;24B | 0-1d | F | none | 25 | 15040.00 | 0.08 | 180987.25 | thorax |
| 5 | Bg2/+;24B | 0-1d | F | none | 25 | 17360.00 | 0.07 | 253854.43 | thorax |
| 5 | Bg2/+;24B | 0-1d | F | none | 25 | 26120.00 | 0.10 | 254982.65 | thorax |
| 5 | Bg2/+;24B | 0-1d | F | none | 25 | 24040.00 | 0.17 | 139019.00 | thorax |
| 4 | Bg2/+;24B | 1-2d | F | none | 25 | 12280.00 | 0.05 | 254530.91 | thorax |
| 4 | Bg2/+;24B | 1-2d | F | none | 25 | 9080.00 | 0.07 | 130129.37 | thorax |
| 4 | Bg2/+;24B | 1-2d | F | none | 25 | 14440.00 | 0.07 | 197180.69 | thorax |
| 4 | Bg2/+;24B | 1-2d | F | none | 25 | 19160.00 | 0.12 | 154181.65 | thorax |
| 4 | Bg2/+;24B | 1-2d | F | none | 25 | 23040.00 | 0.13 | 181902.37 | thorax |
| 5 | Bg2/+;24B | 0-1d | M | none | 25 | 8400.00 | 0.02 | 361098.80 | thorax |
| 5 | Bg2/+;24B | 0-1d | M | none | 25 | 15920.00 | 0.08 | 189341.87 | thorax |
| 5 | Bg2/+;24B | 0-1d | M | none | 25 | 10960.00 | 0.07 | 156421.12 | thorax |
| 5 | Bg2/+;24B | 0-1d | M | none | 25 | 14240.00 | 0.05 | 306074.22 | thorax |
| 5 | Bg2/+;24B | 0-1d | M | none | 25 | 19760.00 | 0.06 | 334934.35 | thorax |

| Rep. | Genotype | Stage | Sex | Treatmen | °C | ß-gal ΔmOD/min/ml | Protein mg/ml | Specific Activity ΔmOD/min/mg | Whole/ dissected |
|---|---|---|---|---|---|---|---|---|---|
| 4 | Bg2/+;24B | 1-2d | M | none | 25 | 12120.00 | 0.16 | 77609.26 | thorax |
| 4 | Bg2/+;24B | 1-2d | M | none | 25 | 13240.00 | 0.05 | 254126.94 | thorax |
| 4 | Bg2/+;24B | 1-2d | M | none | 25 | 13000.00 | 0.14 | 93510.52 | thorax |
| 4 | Bg2/+;24B | 1-2d | M | none | 25 | 14840.00 | 0.07 | 199386.00 | thorax |
| 4 | Bg2/+;24B | 1-2d | M | none | 25 | 14440.00 | 0.09 | 158376.91 | thorax |
| 4 | Bg2/+;24B | EP | ND | none | 25 | 75920.00 | 0.48 | 159299.78 | whole |
| 4 | Bg2/+;24B | EP | ND | none | 25 | 49600.00 | 0.32 | 155665.32 | whole |
| 4 | Bg2/+;24B | EP | ND | none | 25 | 23520.00 | 0.33 | 71778.24 | whole |
| 4 | Bg2/+;24B | EP | ND | none | 25 | 100000.00 | 0.58 | 173021.37 | whole |
| 4 | Bg2/+;24B | EP | ND | none | 25 | 38480.00 | 0.31 | 125094.87 | whole |
| 5 | Bg2/+;24B | EP | ND | none | 25 | 30920.00 | 0.27 | 114457.99 | whole |
| 5 | Bg2/+;24B | EP | ND | none | 25 | 61680.00 | 0.43 | 143864.98 | whole |
| 5 | Bg2/+;24B | EP | ND | none | 25 | 59960.00 | 0.46 | 131266.05 | whole |
| 5 | Bg2/+;24B | EP | ND | none | 25 | 61760.00 | 0.34 | 183085.96 | whole |
| 5 | Bg2/+;24B | EP | ND | none | 25 | 64200.00 | 0.46 | 140118.20 | whole |
| 4 | Bg2/+;24B | LP | ND | none | 25 | 122480.00 | 0.35 | 346474.95 | whole |
| 4 | Bg2/+;24B | LP | ND | none | 25 | 86520.00 | 0.37 | 236837.89 | whole |
| 4 | Bg2/+;24B | LP | ND | none | 25 | 89200.00 | 0.29 | 308920.59 | whole |
| 4 | Bg2/+;24B | LP | ND | none | 25 | 99400.00 | 0.26 | 378704.95 | whole |
| 4 | Bg2/+;24B | LP | ND | none | 25 | 53920.00 | 0.20 | 264704.00 | whole |
| 5 | Bg2/+;24B | LP | ND | none | 25 | 73840.00 | 0.29 | 257131.00 | whole |
| 5 | Bg2/+;24B | LP | ND | none | 25 | 64800.00 | 0.29 | 221767.25 | whole |
| 5 | Bg2/+;24B | LP | ND | none | 25 | 59920.00 | 0.20 | 306660.28 | whole |
| 5 | Bg2/+;24B | LP | ND | none | 25 | 71760.00 | 0.29 | 243339.23 | whole |
| 5 | Bg2/+;24B | LP | ND | none | 25 | 82880.00 | 0.32 | 262281.74 | whole |
| 5 | Bg2/+;24B/+ | 0-1d | F | none | 25 | 6360.00 | 0.16 | 39533.94 | thorax |
| 5 | Bg2/+;24B/+ | 0-1d | F | none | 25 | 7960.00 | 0.07 | 114290.87 | thorax |
| 5 | Bg2/+;24B/+ | 0-1d | F | none | 25 | 13520.00 | 0.11 | 123849.45 | thorax |
| 5 | Bg2/+;24B/+ | 0-1d | F | none | 25 | 4960.00 | 0.10 | 50708.54 | thorax |
| 5 | Bg2/+;24B/+ | 0-1d | F | none | 25 | 8800.00 | 0.10 | 85554.22 | thorax |
| 8 | Bg2/+;24B/+ | 0-4d | F | none | 25 | 4305.00 | 0.11 | 37629.50 | thorax |
| 8 | Bg2/+;24B/+ | 0-4d | F | none | 25 | 6110.00 | 0.15 | 41936.61 | thorax |
| 8 | Bg2/+;24B/+ | 0-4d | F | none | 25 | 6215.00 | 0.14 | 44362.24 | thorax |
| 8 | Bg2/+;24B/+ | 0-4d | F | none | 25 | 13060.00 | 0.19 | 69117.07 | thorax |
| 8 | Bg2/+;24B/+ | 0-4d | F | none | 25 | 5655.00 | 0.14 | 41536.89 | thorax |
| 4 | Bg2/+;24B/+ | 1-2d | F | none | 25 | 7400.00 | 0.06 | 134486.96 | thorax |
| 4 | Bg2/+;24B/+ | 1-2d | F | none | 25 | 4680.00 | 0.08 | 61452.57 | thorax |
| 4 | Bg2/+;24B/+ | 1-2d | F | none | 25 | 6600.00 | 0.08 | 82902.17 | thorax |
| 4 | Bg2/+;24B/+ | 1-2d | F | none | 25 | 10280.00 | 0.06 | 160803.99 | thorax |
| 4 | Bg2/+;24B/+ | 1-2d | F | none | 25 | 5880.00 | 0.09 | 65933.11 | thorax |

| Rep. | Genotype | Stage | Sex | Treatmen | °C | ß-gal ΔmOD/min/ ml | Protein mg/ml | Specific Activity ΔmOD/min/ mg | Whole/ dissected |
|---|---|---|---|---|---|---|---|---|---|
| 9 | Bg2/+;24B/+ | 1-3d | F | none | 25 | 4640.00 | 0.12 | 37628.93 | thorax |
| 9 | Bg2/+;24B/+ | 1-3d | F | none | 25 | 3490.00 | 0.15 | 22953.00 | thorax |
| 9 | Bg2/+;24B/+ | 1-3d | F | none | 25 | 3960.00 | 0.11 | 36036.00 | thorax |
| 9 | Bg2/+;24B/+ | 1-3d | F | none | 25 | 3640.00 | 0.11 | 31867.68 | thorax |
| 9 | Bg2/+;24B/+ | 1-3d | F | none | 25 | 4650.00 | 0.14 | 34434.74 | thorax |
| 10 | Bg2/+;24B/+ | 11-13d | F | none | 25 | 2510.00 | 0.07 | 33518.63 | thorax |
| 10 | Bg2/+;24B/+ | 11-13d | F | none | 25 | 2790.00 | 0.09 | 30960.00 | thorax |
| 10 | Bg2/+;24B/+ | 11-13d | F | none | 25 | 5630.00 | 0.15 | 37767.55 | thorax |
| 10 | Bg2/+;24B/+ | 11-13d | F | none | 25 | 3210.00 | 0.11 | 30369.64 | thorax |
| 10 | Bg2/+;24B/+ | 11-13d | F | none | 25 | 5380.00 | 0.10 | 55610.58 | thorax |
| 11 | Bg2/+;24B/+ | 11-14d | F | none | 25 | 3000.00 | 0.07 | 41165.22 | thorax |
| 11 | Bg2/+;24B/+ | 11-14d | F | none | 25 | 2560.00 | 0.08 | 31934.23 | thorax |
| 11 | Bg2/+;24B/+ | 11-14d | F | none | 25 | 4090.00 | 0.11 | 37163.26 | thorax |
| 11 | Bg2/+;24B/+ | 11-14d | F | none | 25 | 4220.00 | 0.11 | 37063.97 | thorax |
| 11 | Bg2/+;24B/+ | 11-14d | F | none | 25 | 2950.00 | 0.07 | 44617.57 | thorax |
| 5 | Bg2/+;24B/+ | 0-1d | M | none | 25 | 5080.00 | 0.07 | 71783.92 | thorax |
| 5 | Bg2/+;24B/+ | 0-1d | M | none | 25 | 5480.00 | 0.10 | 53642.36 | thorax |
| 5 | Bg2/+;24B/+ | 0-1d | M | none | 25 | 3000.00 | 0.02 | 165953.49 | thorax |
| 5 | Bg2/+;24B/+ | 0-1d | M | none | 25 | 5400.00 | 0.03 | 167540.87 | thorax |
| 5 | Bg2/+;24B/+ | 0-1d | M | none | 25 | 5200.00 | 0.04 | 133000.72 | thorax |
| 8 | Bg2/+;24B/+ | 0-4d | M | none | 25 | 3320.00 | 0.08 | 41479.51 | thorax |
| 8 | Bg2/+;24B/+ | 0-4d | M | none | 25 | 2705.00 | 0.11 | 23644.09 | thorax |
| 8 | Bg2/+;24B/+ | 0-4d | M | none | 25 | 3085.00 | 0.15 | 21174.21 | thorax |
| 8 | Bg2/+;24B/+ | 0-4d | M | none | 25 | 4730.00 | 0.14 | 33762.41 | thorax |
| 8 | Bg2/+;24B/+ | 0-4d | M | none | 25 | 4755.00 | 0.19 | 25164.75 | thorax |
| 4 | Bg2/+;24B/+ | 1-2d | M | none | 25 | 4600.00 | 0.03 | 146654.24 | thorax |
| 4 | Bg2/+;24B/+ | 1-2d | M | none | 25 | 5480.00 | 0.03 | 162970.43 | thorax |
| 4 | Bg2/+;24B/+ | 1-2d | M | none | 25 | 5360.00 | 0.01 | 376903.18 | thorax |
| 4 | Bg2/+;24B/+ | 1-2d | M | none | 25 | 1000.00 | 0.02 | 49500.00 | thorax |
| 4 | Bg2/+;24B/+ | 1-2d | M | none | 25 | 4520.00 | 0.03 | 162720.00 | thorax |
| 9 | Bg2/+;24B/+ | 1-3d | M | none | 25 | 3105.00 | 0.10 | 31837.18 | thorax |
| 9 | Bg2/+;24B/+ | 1-3d | M | none | 25 | 3105.00 | 0.12 | 25180.57 | thorax |
| 9 | Bg2/+;24B/+ | 1-3d | M | none | 25 | 2805.00 | 0.15 | 18447.89 | thorax |
| 9 | Bg2/+;24B/+ | 1-3d | M | none | 25 | 3775.00 | 0.11 | 34352.50 | thorax |
| 9 | Bg2/+;24B/+ | 1-3d | M | none | 25 | 5320.00 | 0.11 | 46575.84 | thorax |
| 10 | Bg2/+;24B/+ | 11-13d | M | none | 25 | 2630.00 | 0.07 | 39611.21 | thorax |
| 10 | Bg2/+;24B/+ | 11-13d | M | none | 25 | 4290.00 | 0.14 | 31107.93 | thorax |
| 10 | Bg2/+;24B/+ | 11-13d | M | none | 25 | 2550.00 | 0.09 | 29240.00 | thorax |
| 10 | Bg2/+;24B/+ | 11-13d | M | none | 25 | 2150.00 | 0.11 | 19712.15 | thorax |
| 10 | Bg2/+;24B/+ | 11-13d | M | none | 25 | 5190.00 | 0.09 | 55932.33 | thorax |

| Rep. | Genotype | Stage | Sex | Treatmen | °C | ß-gal ΔmOD/min/ ml | Protein mg/ml | Specific Activity ΔmOD/min/ mg | Whole/ dissected |
|---|---|---|---|---|---|---|---|---|---|
| 11 | Bg2/+;24B/+ | 11-14d | M | none | 25 | 3100.00 | 0.10 | 29980.39 | thorax |
| 11 | Bg2/+;24B/+ | 11-14d | M | none | 25 | 2160.00 | 0.07 | 31706.79 | thorax |
| 11 | Bg2/+;24B/+ | 11-14d | M | none | 25 | 2300.00 | 0.09 | 25350.87 | thorax |
| 11 | Bg2/+;24B/+ | 11-14d | M | none | 25 | 2510.00 | 0.08 | 33006.50 | thorax |
| 11 | Bg2/+;24B/+ | 11-14d | M | none | 25 | 4750.00 | 0.10 | 47340.00 | thorax |
| 4 | Bg2/+;24B/+ | EP | ND | none | 25 | 29480.00 | 0.39 | 75471.79 | whole |
| 4 | Bg2/+;24B/+ | EP | ND | none | 25 | 29880.00 | 0.30 | 98606.44 | whole |
| 4 | Bg2/+;24B/+ | EP | ND | none | 25 | 35480.00 | 0.46 | 76719.68 | whole |
| 4 | Bg2/+;24B/+ | EP | ND | none | 25 | 38360.00 | 0.36 | 106114.43 | whole |
| 4 | Bg2/+;24B/+ | EP | ND | none | 25 | 33880.00 | 0.46 | 74194.07 | whole |
| 5 | Bg2/+;24B/+ | EP | ND | none | 25 | 24520.00 | 0.35 | 71024.70 | whole |
| 5 | Bg2/+;24B/+ | EP | ND | none | 25 | 25960.00 | 0.32 | 80422.69 | whole |
| 5 | Bg2/+;24B/+ | EP | ND | none | 25 | 21320.00 | 0.36 | 58596.38 | whole |
| 5 | Bg2/+;24B/+ | EP | ND | none | 25 | 29920.00 | 0.46 | 64440.55 | whole |
| 5 | Bg2/+;24B/+ | EP | ND | none | 25 | 25280.00 | 0.33 | 76092.22 | whole |
| 8 | Bg2/+;24B/+ | EP | ND | none | 25 | 28750.00 | 0.44 | 65407.35 | whole |
| 8 | Bg2/+;24B/+ | EP | ND | none | 25 | 33420.00 | 0.34 | 98980.77 | whole |
| 8 | Bg2/+;24B/+ | EP | ND | none | 25 | 36020.00 | 0.29 | 124444.91 | whole |
| 8 | Bg2/+;24B/+ | EP | ND | none | 25 | 22740.00 | 0.41 | 55020.43 | whole |
| 8 | Bg2/+;24B/+ | EP | ND | none | 25 | 31570.00 | 0.57 | 55173.58 | whole |
| 8 | Bg2/+;24B/+ | EP | ND | none | 25 | 17305.00 | 0.34 | 51252.61 | whole |
| 8 | Bg2/+;24B/+ | EP | ND | none | 25 | 14535.00 | 0.29 | 50216.73 | whole |
| 8 | Bg2/+;24B/+ | EP | ND | none | 25 | 20350.00 | 0.41 | 49237.72 | whole |
| 8 | Bg2/+;24B/+ | EP | ND | none | 25 | 34285.00 | 0.57 | 59918.47 | whole |
| 8 | Bg2/+;24B/+ | EP | ND | none | 25 | 22320.00 | 0.35 | 63522.21 | whole |
| 9 | Bg2/+;24B/+ | EP | ND | none | 25 | 38100.00 | 0.41 | 91990.54 | whole |
| 9 | Bg2/+;24B/+ | EP | ND | none | 25 | 42215.00 | 0.64 | 66336.34 | whole |
| 9 | Bg2/+;24B/+ | EP | ND | none | 25 | 19305.00 | 0.69 | 28047.31 | whole |
| 9 | Bg2/+;24B/+ | EP | ND | none | 25 | 29070.00 | 0.41 | 70188.06 | whole |
| 9 | Bg2/+;24B/+ | EP | ND | none | 25 | 30670.00 | 0.40 | 77327.11 | whole |
| 9 | Bg2/+;24B/+ | EP | ND | none | 25 | 48705.00 | 0.69 | 70761.16 | whole |
| 9 | Bg2/+;24B/+ | EP | ND | none | 25 | 19995.00 | 0.51 | 39533.84 | whole |
| 10 | Bg2/+;24B/+ | EP | ND | none | 25 | 16760.00 | 0.36 | 46420.24 | whole |
| 10 | Bg2/+;24B/+ | EP | ND | none | 25 | 15620.00 | 0.61 | 25717.01 | whole |
| 10 | Bg2/+;24B/+ | EP | ND | none | 25 | 16700.00 | 0.39 | 43028.18 | whole |
| 10 | Bg2/+;24B/+ | EP | ND | none | 25 | 21620.00 | 0.48 | 45381.84 | whole |
| 10 | Bg2/+;24B/+ | EP | ND | none | 25 | 17540.00 | 0.40 | 43752.83 | whole |
| 11 | Bg2/+;24B/+ | EP | ND | none | 25 | 18140.00 | 0.40 | 45486.66 | whole |
| 11 | Bg2/+;24B/+ | EP | ND | none | 25 | 22260.00 | 0.53 | 41764.83 | whole |
| 11 | Bg2/+;24B/+ | EP | ND | none | 25 | 22260.00 | 0.47 | 47128.30 | whole |

| Rep. | Genotype | Stage | Sex | Treatmen | °C | ß-gal ΔmOD/min/ml | Protein mg/ml | Specific Activity ΔmOD/min/mg | Whole/ dissected |
|---|---|---|---|---|---|---|---|---|---|
| 11 | Bg2/+;24B/+ | EP | ND | none | 25 | 22910.00 | 0.50 | 45470.83 | whole |
| 11 | Bg2/+;24B/+ | EP | ND | none | 25 | 26800.00 | 0.68 | 39690.88 | whole |
| 2 | Bg2/+;24B/+ | L3 | ND | none | 25 | 61840.00 | 0.60 | 103583.11 | whole |
| 2 | Bg2/+;24B/+ | L3 | ND | none | 25 | 69320.00 | 0.69 | 99882.27 | whole |
| 2 | Bg2/+;24B/+ | L3 | ND | none | 25 | 61240.00 | 0.52 | 117846.71 | whole |
| 2 | Bg2/+;24B/+ | L3 | ND | none | 25 | 54080.00 | 0.40 | 133644.04 | whole |
| 2 | Bg2/+;24B/+ | L3 | ND | none | 25 | 42240.00 | 0.42 | 101045.33 | whole |
| 3 | Bg2/+;24B/+ | L3 | ND | none | 25 | 39720.00 | 0.47 | 85406.94 | whole |
| 3 | Bg2/+;24B/+ | L3 | ND | none | 25 | 9840.00 | 0.32 | 30624.29 | whole |
| 3 | Bg2/+;24B/+ | L3 | ND | none | 25 | 4760.00 | 0.26 | 18596.93 | whole |
| 3 | Bg2/+;24B/+ | L3 | ND | none | 25 | 25600.00 | 0.34 | 75119.13 | whole |
| 3 | Bg2/+;24B/+ | L3 | ND | none | 25 | 35560.00 | 0.48 | 73903.89 | whole |
| 4 | Bg2/+;24B/+ | LP | ND | none | 25 | 36120.00 | 0.23 | 157007.17 | whole |
| 4 | Bg2/+;24B/+ | LP | ND | none | 25 | 37680.00 | 0.23 | 161413.24 | whole |
| 4 | Bg2/+;24B/+ | LP | ND | none | 25 | 43320.00 | 0.27 | 162336.91 | whole |
| 4 | Bg2/+;24B/+ | LP | ND | none | 25 | 38080.00 | 0.22 | 170269.06 | whole |
| 4 | Bg2/+;24B/+ | LP | ND | none | 25 | 56760.00 | 0.35 | 162949.85 | whole |
| 5 | Bg2/+;24B/+ | LP | ND | none | 25 | 32080.00 | 0.20 | 157944.02 | whole |
| 5 | Bg2/+;24B/+ | LP | ND | none | 25 | 45760.00 | 0.32 | 143420.04 | whole |
| 5 | Bg2/+;24B/+ | LP | ND | none | 25 | 30720.00 | 0.22 | 136797.17 | whole |
| 5 | Bg2/+;24B/+ | LP | ND | none | 25 | 19120.00 | 0.18 | 108051.44 | whole |
| 5 | Bg2/+;24B/+ | LP | ND | none | 25 | 29160.00 | 0.23 | 125472.70 | whole |
| 8 | Bg2/+;24B/+ | LP | ND | none | 25 | 31940.00 | 0.33 | 97533.35 | whole |
| 8 | Bg2/+;24B/+ | LP | ND | none | 25 | 24130.00 | 0.32 | 75933.84 | whole |
| 8 | Bg2/+;24B/+ | LP | ND | none | 25 | 20620.00 | 0.25 | 84024.70 | whole |
| 8 | Bg2/+;24B/+ | LP | ND | none | 25 | 35100.00 | 0.24 | 144987.02 | whole |
| 8 | Bg2/+;24B/+ | LP | ND | none | 25 | 33610.00 | 0.33 | 103294.55 | whole |
| 8 | Bg2/+;24B/+ | LP | ND | none | 25 | 32380.00 | 0.32 | 101895.48 | whole |
| 8 | Bg2/+;24B/+ | LP | ND | none | 25 | 24840.00 | 0.25 | 101220.84 | whole |
| 8 | Bg2/+;24B/+ | LP | ND | none | 25 | 25235.00 | 0.24 | 104237.82 | whole |
| 8 | Bg2/+;24B/+ | LP | ND | none | 25 | 30525.00 | 0.33 | 93813.34 | whole |
| 8 | Bg2/+;24B/+ | LP | ND | none | 25 | 30165.00 | 0.38 | 79453.94 | whole |
| 9 | Bg2/+;24B/+ | LP | ND | none | 25 | 25000.00 | 0.41 | 61187.70 | whole |
| 9 | Bg2/+;24B/+ | LP | ND | none | 25 | 21200.00 | 0.38 | 55350.23 | whole |
| 9 | Bg2/+;24B/+ | LP | ND | none | 25 | 22690.00 | 0.40 | 56472.16 | whole |
| 9 | Bg2/+;24B/+ | LP | ND | none | 25 | 24100.00 | 0.48 | 50489.68 | whole |
| 9 | Bg2/+;24B/+ | LP | ND | none | 25 | 26925.00 | 0.39 | 69719.04 | whole |
| 9 | Bg2/+;24B/+ | LP | ND | none | 25 | 19090.00 | 0.38 | 49841.31 | whole |
| 9 | Bg2/+;24B/+ | LP | ND | none | 25 | 17305.00 | 0.40 | 43069.67 | whole |
| 9 | Bg2/+;24B/+ | LP | ND | none | 25 | 29190.00 | 0.48 | 61153.27 | whole |

| Rep. | Genotype | Stage | Sex | Treatmen | °C | ß-gal ΔmOD/min/ ml | Protein mg/ml | Specific Activity ΔmOD/min/ mg | Whole/ dissected |
|---|---|---|---|---|---|---|---|---|---|
| 9 | Bg2/+;24B/+ | LP | ND | none | 25 | 26345.00 | 0.39 | 68217.20 | whole |
| 9 | Bg2/+;24B/+ | LP | ND | none | 25 | 24470.00 | 0.42 | 58103.66 | whole |
| 10 | Bg2/+;24B/+ | LP | ND | none | 25 | 30810.00 | 0.39 | 78334.21 | whole |
| 10 | Bg2/+;24B/+ | LP | ND | none | 25 | 34560.00 | 0.42 | 81491.23 | whole |
| 10 | Bg2/+;24B/+ | LP | ND | none | 25 | 10720.00 | 0.18 | 60993.34 | whole |
| 10 | Bg2/+;24B/+ | LP | ND | none | 25 | 17280.00 | 0.23 | 75581.07 | whole |
| 10 | Bg2/+;24B/+ | LP | ND | none | 25 | 27450.00 | 0.33 | 84190.94 | whole |
| 11 | Bg2/+;24B/+ | LP | ND | none | 25 | 42060.00 | 0.36 | 116866.29 | whole |
| 11 | Bg2/+;24B/+ | LP | ND | none | 25 | 43040.00 | 0.42 | 102185.96 | whole |
| 11 | Bg2/+;24B/+ | LP | ND | none | 25 | 26397.14 | 0.35 | 75039.50 | whole |
| 11 | Bg2/+;24B/+ | LP | ND | none | 25 | 37653.33 | 0.32 | 118210.46 | whole |
| 11 | Bg2/+;24B/+ | LP | ND | none | 25 | 38573.33 | 0.35 | 110489.96 | whole |
| 8 | Bg2/+;24B/3.3,DJ1077 | 0-4d | F | none | 25 | 1595.00 | 0.16 | 9929.77 | thorax |
| 8 | Bg2/+;24B/3.3,DJ1077 | 0-4d | F | none | 25 | 490.00 | 0.11 | 4512.56 | thorax |
| 8 | Bg2/+;24B/3.3,DJ1077 | 0-4d | F | none | 25 | 1370.00 | 0.13 | 10897.78 | thorax |
| 8 | Bg2/+;24B/3.3,DJ1077 | 0-4d | F | none | 25 | 880.00 | 0.12 | 7442.01 | thorax |
| 8 | Bg2/+;24B/3.3,DJ1077 | 0-4d | F | none | 25 | 1875.00 | 0.14 | 13436.27 | thorax |
| 9 | Bg2/+;24B/3.3,DJ1077 | 1-3d | F | none | 25 | 1485.00 | 0.14 | 10598.82 | thorax |
| 9 | Bg2/+;24B/3.3,DJ1077 | 1-3d | F | none | 25 | 1155.00 | 0.14 | 8363.37 | thorax |
| 9 | Bg2/+;24B/3.3,DJ1077 | 1-3d | F | none | 25 | 1585.00 | 0.14 | 11070.44 | thorax |
| 9 | Bg2/+;24B/3.3,DJ1077 | 1-3d | F | none | 25 | 1770.00 | 0.14 | 12576.04 | thorax |
| 9 | Bg2/+;24B/3.3,DJ1077 | 1-3d | F | none | 25 | 1340.00 | 0.16 | 8586.16 | thorax |
| 8 | Bg2/+;24B/3.3,DJ1077 | 0-4d | M | none | 25 | 1055.00 | 0.08 | 13495.70 | thorax |
| 8 | Bg2/+;24B/3.3,DJ1077 | 0-4d | M | none | 25 | 590.00 | 0.10 | 6169.60 | thorax |
| 8 | Bg2/+;24B/3.3,DJ1077 | 0-4d | M | none | 25 | 705.00 | 0.10 | 7095.18 | thorax |
| 8 | Bg2/+;24B/3.3,DJ1077 | 0-4d | M | none | 25 | 705.00 | 0.09 | 7840.22 | thorax |
| 8 | Bg2/+;24B/3.3,DJ1077 | 0-4d | M | none | 25 | 705.00 | 0.11 | 6351.28 | thorax |
| 9 | Bg2/+;24B/3.3,DJ1077 | 1-3d | M | none | 25 | 1070.00 | 0.13 | 8075.34 | thorax |
| 9 | Bg2/+;24B/3.3,DJ1077 | 1-3d | M | none | 25 | 810.00 | 0.10 | 7735.46 | thorax |
| 9 | Bg2/+;24B/3.3,DJ1077 | 1-3d | M | none | 25 | 1110.00 | 0.08 | 13382.22 | thorax |
| 9 | Bg2/+;24B/3.3,DJ1077 | 1-3d | M | none | 25 | 785.00 | 0.12 | 6735.49 | thorax |
| 9 | Bg2/+;24B/3.3,DJ1077 | 1-3d | M | none | 25 | 665.00 | 0.11 | 6039.88 | thorax |
| 8 | Bg2/+;24B/3.3,DJ1077 | EP | ND | none | 25 | 13405.00 | 0.66 | 20257.58 | whole |
| 8 | Bg2/+;24B/3.3,DJ1077 | EP | ND | none | 25 | 10710.00 | 0.59 | 18012.79 | whole |
| 8 | Bg2/+;24B/3.3,DJ1077 | EP | ND | none | 25 | 3760.00 | 0.29 | 12782.28 | whole |
| 8 | Bg2/+;24B/3.3,DJ1077 | EP | ND | none | 25 | 6635.00 | 0.49 | 13517.49 | whole |
| 8 | Bg2/+;24B/3.3,DJ1077 | EP | ND | none | 25 | 10670.00 | 0.57 | 18834.30 | whole |
| 8 | Bg2/+;24B/3.3,DJ1077 | EP | ND | none | 25 | 10325.00 | 0.59 | 17560.80 | whole |
| 8 | Bg2/+;24B/3.3,DJ1077 | EP | ND | none | 25 | 10255.00 | 0.52 | 19637.10 | whole |
| 8 | Bg2/+;24B/3.3,DJ1077 | EP | ND | none | 25 | 3895.00 | 0.30 | 12807.46 | whole |

| Rep. | Genotype | Stage | Sex | Treatmen | °C | ß-gal ΔmOD/min/ ml | Protein mg/ml | Specific Activity ΔmOD/min/ mg | Whole/ dissected |
|---|---|---|---|---|---|---|---|---|---|
| 8 | Bg2/+;24B/3.3,DJ1077 | EP | ND | none | 25 | 7095.00 | 0.44 | 16176.08 | whole |
| 9 | Bg2/+;24B/3.3,DJ1077 | EP | ND | none | 25 | 3280.00 | 0.27 | 12366.44 | whole |
| 9 | Bg2/+;24B/3.3,DJ1077 | EP | ND | none | 25 | 11320.00 | 0.58 | 19577.83 | whole |
| 9 | Bg2/+;24B/3.3,DJ1077 | EP | ND | none | 25 | 12150.00 | 0.58 | 20794.30 | whole |
| 9 | Bg2/+;24B/3.3,DJ1077 | EP | ND | none | 25 | 7145.00 | 0.39 | 18207.99 | whole |
| 9 | Bg2/+;24B/3.3,DJ1077 | EP | ND | none | 25 | 6555.00 | 0.31 | 21466.53 | whole |
| 9 | Bg2/+;24B/3.3,DJ1077 | EP | ND | none | 25 | 6195.00 | 0.51 | 12156.23 | whole |
| 9 | Bg2/+;24B/3.3,DJ1077 | EP | ND | none | 25 | 7235.00 | 0.41 | 17468.55 | whole |
| 8 | Bg2/+;24B/3.3,DJ1077 | LP | ND | none | 25 | 12125.00 | 0.31 | 39357.23 | whole |
| 8 | Bg2/+;24B/3.3,DJ1077 | LP | ND | none | 25 | 1580.00 | 0.19 | 8239.96 | whole |
| 8 | Bg2/+;24B/3.3,DJ1077 | LP | ND | none | 25 | 11830.00 | 0.31 | 37568.38 | whole |
| 8 | Bg2/+;24B/3.3,DJ1077 | LP | ND | none | 25 | 13540.00 | 0.40 | 34030.68 | whole |
| 8 | Bg2/+;24B/3.3,DJ1077 | LP | ND | none | 25 | 4935.00 | 0.24 | 20215.35 | whole |
| 9 | Bg2/+;24B/3.3,DJ1077 | LP | ND | none | 25 | 12140.00 | 0.47 | 25682.05 | whole |
| 9 | Bg2/+;24B/3.3,DJ1077 | LP | ND | none | 25 | 10065.00 | 0.43 | 23663.86 | whole |
| 9 | Bg2/+;24B/3.3,DJ1077 | LP | ND | none | 25 | 7685.00 | 0.15 | 51725.75 | whole |
| 9 | Bg2/+;24B/3.3,DJ1077 | LP | ND | none | 25 | 9740.00 | 0.48 | 20343.82 | whole |
| 9 | Bg2/+;24B/3.3,DJ1077 | LP | ND | none | 25 | 11940.00 | 0.34 | 35512.27 | whole |
| 9 | Bg2/+;24B/3.3,DJ1077 | LP | ND | none | 25 | 10370.00 | 0.42 | 24665.71 | whole |
| 9 | Bg2/+;24B/3.3,DJ1077 | LP | ND | none | 25 | 10875.00 | 0.36 | 30621.59 | whole |
| 12 | Bg2/+;Mef2/+ | 10d | F | none | 20 | 24218.00 | 0.11 | 228189.16 | thorax |
| 12 | Bg2/+;Mef2/+ | 10d | F | none | 20 | 24186.00 | 0.10 | 250749.15 | thorax |
| 12 | Bg2/+;Mef2/+ | 10d | F | none | 20 | 25424.00 | 0.11 | 223786.32 | thorax |
| 12 | Bg2/+;Mef2/+ | 10d | F | none | 20 | 23886.00 | 0.11 | 212220.44 | thorax |
| 13 | Bg2/+;Mef2/+ | 10d | F | none | 20 | 24500.00 | 0.09 | 283910.30 | thorax |
| 13 | Bg2/+;Mef2/+ | 10d | F | none | 20 | 19900.00 | 0.11 | 177290.91 | thorax |
| 13 | Bg2/+;Mef2/+ | 10d | F | none | 20 | 24292.00 | 0.11 | 228035.88 | thorax |
| 13 | Bg2/+;Mef2/+ | 10d | F | none | 20 | 21840.00 | 0.11 | 190032.24 | thorax |
| 13 | Bg2/+;Mef2/+ | 10d | F | none | 20 | 20702.00 | 0.12 | 176881.12 | thorax |
| 14 | Bg2/+;Mef2/+ | 10d | F | none | 20 | 26158.00 | 0.10 | 269595.78 | thorax |
| 14 | Bg2/+;Mef2/+ | 10d | F | none | 20 | 25278.00 | 0.13 | 196458.37 | thorax |
| 14 | Bg2/+;Mef2/+ | 10d | F | none | 20 | 26242.00 | 0.12 | 212821.97 | thorax |
| 14 | Bg2/+;Mef2/+ | 10d | F | none | 20 | 26368.00 | 0.10 | 252675.95 | thorax |
| 14 | Bg2/+;Mef2/+ | 10d | F | none | 20 | 22756.00 | 0.10 | 220357.76 | thorax |
| 15 | Bg2/+;Mef2/+ | 10d | F | none | 20 | 25026.00 | 0.11 | 231896.18 | thorax |
| 15 | Bg2/+;Mef2/+ | 10d | F | none | 20 | 24548.00 | 0.10 | 256145.30 | thorax |
| 15 | Bg2/+;Mef2/+ | 10d | F | none | 20 | 21476.00 | 0.08 | 271196.38 | thorax |
| 15 | Bg2/+;Mef2/+ | 10d | F | none | 20 | 21250.00 | 0.09 | 245565.04 | thorax |
| 15 | Bg2/+;Mef2/+ | 10d | F | none | 20 | 23030.00 | 0.15 | 152634.67 | thorax |
| 6 | Bg2/+;Mef2/+ | 10-11d | F | -RU486 | 25 | 42135.00 | 0.06 | 656687.75 | thorax |

| Rep. | Genotype | Stage | Sex | Treatmen | °C | ß-gal ΔmOD/min/ml | Protein mg/ml | Specific Activity ΔmOD/min/mg | Whole/ dissected |
|---|---|---|---|---|---|---|---|---|---|
| 6 | Bg2/+;Mef2/+ | 10-11d | F | -RU486 | 25 | 42260.00 | 0.09 | 493976.94 | thorax |
| 6 | Bg2/+;Mef2/+ | 10-11d | F | -RU486 | 25 | 58305.00 | 0.07 | 786956.44 | thorax |
| 6 | Bg2/+;Mef2/+ | 10-11d | F | -RU486 | 25 | 49880.00 | 0.10 | 496867.85 | thorax |
| 6 | Bg2/+;Mef2/+ | 10-11d | F | -RU486 | 25 | 45085.00 | 0.07 | 624037.71 | thorax |
| 7 | Bg2/+;Mef2/+ | 10-12d | F | -RU486 | 25 | 53660.00 | 0.13 | 427457.58 | thorax |
| 7 | Bg2/+;Mef2/+ | 10-12d | F | -RU486 | 25 | 36600.00 | 0.12 | 305687.83 | thorax |
| 7 | Bg2/+;Mef2/+ | 10-12d | F | -RU486 | 25 | 54500.00 | 0.10 | 557140.44 | thorax |
| 7 | Bg2/+;Mef2/+ | 10-12d | F | -RU486 | 25 | 44320.00 | 0.10 | 426725.29 | thorax |
| 7 | Bg2/+;Mef2/+ | 10-12d | F | -RU486 | 25 | 50420.00 | 0.11 | 462265.45 | thorax |
| 6 | Bg2/+;Mef2/+ | 10-11d | F | +RU486 | 25 | 54310.00 | 0.09 | 634829.33 | thorax |
| 6 | Bg2/+;Mef2/+ | 10-11d | F | +RU486 | 25 | 40135.00 | 0.07 | 541711.63 | thorax |
| 6 | Bg2/+;Mef2/+ | 10-11d | F | +RU486 | 25 | 50720.00 | 0.10 | 505235.31 | thorax |
| 6 | Bg2/+;Mef2/+ | 10-11d | F | +RU486 | 25 | 48555.00 | 0.07 | 672067.22 | thorax |
| 6 | Bg2/+;Mef2/+ | 10-11d | F | +RU486 | 25 | 47925.00 | 0.07 | 706369.68 | thorax |
| 7 | Bg2/+;Mef2/+ | 10-12d | F | +RU486 | 25 | 52660.00 | 0.12 | 439822.99 | thorax |
| 7 | Bg2/+;Mef2/+ | 10-12d | F | +RU486 | 25 | 51460.00 | 0.10 | 526063.24 | thorax |
| 7 | Bg2/+;Mef2/+ | 10-12d | F | +RU486 | 25 | 50820.00 | 0.10 | 489309.10 | thorax |
| 7 | Bg2/+;Mef2/+ | 10-12d | F | +RU486 | 25 | 41500.00 | 0.11 | 380484.26 | thorax |
| 7 | Bg2/+;Mef2/+ | 10-12d | F | +RU486 | 25 | 41880.00 | 0.11 | 364571.88 | thorax |
| 1 | Bg2/+;Mef2/+ | 0-24h | F | none | 25 | 30735.00 | 0.21 | 147727.63 | thorax |
| 1 | Bg2/+;Mef2/+ | 0-24h | F | none | 25 | 30975.00 | 0.16 | 191211.68 | thorax |
| 1 | Bg2/+;Mef2/+ | 0-24h | F | none | 25 | 22965.00 | 0.20 | 115967.70 | thorax |
| 1 | Bg2/+;Mef2/+ | 0-24h | F | none | 25 | 32415.00 | 0.20 | 160717.59 | thorax |
| 1 | Bg2/+;Mef2/+ | 0-24h | F | none | 25 | 28935.00 | 0.18 | 164285.68 | thorax |
| 8 | Bg2/+;Mef2/+ | 0-4d | F | none | 25 | 54440.00 | 0.13 | 410463.18 | thorax |
| 8 | Bg2/+;Mef2/+ | 0-4d | F | none | 25 | 59905.00 | 0.15 | 410545.33 | thorax |
| 8 | Bg2/+;Mef2/+ | 0-4d | F | none | 25 | 56100.00 | 0.21 | 271209.55 | thorax |
| 8 | Bg2/+;Mef2/+ | 0-4d | F | none | 25 | 51573.33 | 0.11 | 490323.51 | thorax |
| 9 | Bg2/+;Mef2/+ | 1-3d | F | none | 25 | 52980.00 | 0.15 | 346511.90 | thorax |
| 9 | Bg2/+;Mef2/+ | 1-3d | F | none | 25 | 56850.00 | 0.13 | 433544.24 | thorax |
| 9 | Bg2/+;Mef2/+ | 1-3d | F | none | 25 | 56373.33 | 0.14 | 399040.56 | thorax |
| 9 | Bg2/+;Mef2/+ | 1-3d | F | none | 25 | 58520.00 | 0.14 | 426025.60 | thorax |
| 9 | Bg2/+;Mef2/+ | 1-3d | F | none | 25 | 43550.00 | 0.14 | 304624.69 | thorax |
| 1 | Bg2/+;Mef2/+ | 10-11d | F | none | 25 | 14290.00 | 0.12 | 121066.04 | thorax |
| 1 | Bg2/+;Mef2/+ | 10-11d | F | none | 25 | 15830.00 | 0.21 | 77211.63 | thorax |
| 1 | Bg2/+;Mef2/+ | 10-11d | F | none | 25 | 25980.00 | 0.08 | 325408.03 | thorax |
| 1 | Bg2/+;Mef2/+ | 10-11d | F | none | 25 | 16090.00 | 0.13 | 120959.69 | thorax |
| 1 | Bg2/+;Mef2/+ | 10-11d | F | none | 25 | 21200.00 | 0.12 | 172809.77 | thorax |
| 1 | Bg2/+;Mef2/+ | 3-5d | F | none | 25 | 27515.00 | 0.10 | 275150.00 | thorax |
| 1 | Bg2/+;Mef2/+ | 3-5d | F | none | 25 | 34545.00 | 0.14 | 244134.28 | thorax |

| Rep. | Genotype | Stage | Sex | Treatmen | °C | ß-gal ΔmOD/min/ ml | Protein mg/ml | Specific Activity ΔmOD/min/ mg | Whole/ dissected |
|---|---|---|---|---|---|---|---|---|---|
| 1 | Bg2/+;Mef2/+ | 3-5d | F | none | 25 | 29465.00 | 0.16 | 183582.55 | thorax |
| 1 | Bg2/+;Mef2/+ | 3-5d | F | none | 25 | 28245.00 | 0.20 | 139619.38 | thorax |
| 1 | Bg2/+;Mef2/+ | 3-5d | F | none | 25 | 32435.00 | 0.16 | 202213.22 | thorax |
| 12 | Bg2/+;Mef2/+ | 15d | F | none | 29 | 26610.00 | 0.11 | 244556.97 | thorax |
| 12 | Bg2/+;Mef2/+ | 15d | F | none | 29 | 30886.00 | 0.11 | 286261.77 | thorax |
| 12 | Bg2/+;Mef2/+ | 15d | F | none | 29 | 19080.00 | 0.12 | 165279.37 | thorax |
| 12 | Bg2/+;Mef2/+ | 15d | F | none | 29 | 25442.00 | 0.11 | 230910.70 | thorax |
| 12 | Bg2/+;Mef2/+ | 15d | F | none | 29 | 28924.00 | 0.10 | 275553.09 | thorax |
| 13 | Bg2/+;Mef2/+ | 15d | F | none | 29 | 29084.00 | 0.12 | 238698.41 | thorax |
| 13 | Bg2/+;Mef2/+ | 15d | F | none | 29 | 25764.00 | 0.12 | 219014.03 | thorax |
| 13 | Bg2/+;Mef2/+ | 15d | F | none | 29 | 19106.00 | 0.10 | 184837.87 | thorax |
| 13 | Bg2/+;Mef2/+ | 15d | F | none | 29 | 26862.00 | 0.12 | 228881.83 | thorax |
| 13 | Bg2/+;Mef2/+ | 15d | F | none | 29 | 29772.00 | 0.14 | 205757.94 | thorax |
| 14 | Bg2/+;Mef2/+ | 15d | F | none | 29 | 19218.00 | 0.12 | 162861.38 | thorax |
| 14 | Bg2/+;Mef2/+ | 15d | F | none | 29 | 10952.00 | 0.10 | 106235.37 | thorax |
| 14 | Bg2/+;Mef2/+ | 15d | F | none | 29 | 12968.00 | 0.11 | 121219.47 | thorax |
| 14 | Bg2/+;Mef2/+ | 15d | F | none | 29 | 21498.00 | 0.11 | 201557.58 | thorax |
| 14 | Bg2/+;Mef2/+ | 15d | F | none | 29 | 16920.00 | 0.12 | 144676.92 | thorax |
| 15 | Bg2/+;Mef2/+ | 15d | F | none | 29 | 24458.00 | 0.09 | 259209.75 | thorax |
| 15 | Bg2/+;Mef2/+ | 15d | F | none | 29 | 12126.00 | 0.09 | 139099.09 | thorax |
| 15 | Bg2/+;Mef2/+ | 15d | F | none | 29 | 18524.00 | 0.09 | 213499.60 | thorax |
| 15 | Bg2/+;Mef2/+ | 15d | F | none | 29 | 10612.00 | 0.11 | 95757.64 | thorax |
| 15 | Bg2/+;Mef2/+ | 15d | F | none | 29 | 18140.00 | 0.08 | 226635.98 | thorax |
| 12 | Bg2/+;Mef2/+ | 15d | F | none | 20-29 | 25556.00 | 0.18 | 138453.39 | thorax |
| 12 | Bg2/+;Mef2/+ | 15d | F | none | 20-29 | 23836.00 | 0.15 | 157510.00 | thorax |
| 12 | Bg2/+;Mef2/+ | 15d | F | none | 20-29 | 17810.00 | 0.12 | 151417.86 | thorax |
| 12 | Bg2/+;Mef2/+ | 15d | F | none | 20-29 | 19210.00 | 0.12 | 158297.00 | thorax |
| 12 | Bg2/+;Mef2/+ | 15d | F | none | 20-29 | 23208.00 | 0.13 | 182201.96 | thorax |
| 13 | Bg2/+;Mef2/+ | 15d | F | none | 20-29 | 19488.00 | 0.11 | 176773.04 | thorax |
| 13 | Bg2/+;Mef2/+ | 15d | F | none | 20-29 | 17342.00 | 0.11 | 156874.63 | thorax |
| 13 | Bg2/+;Mef2/+ | 15d | F | none | 20-29 | 25928.00 | 0.15 | 168761.05 | thorax |
| 13 | Bg2/+;Mef2/+ | 15d | F | none | 20-29 | 19660.00 | 0.09 | 209415.81 | thorax |
| 13 | Bg2/+;Mef2/+ | 15d | F | none | 20-29 | 23078.00 | 0.10 | 230380.03 | thorax |
| 14 | Bg2/+;Mef2/+ | 15d | F | none | 20-29 | 28308.00 | 0.14 | 208818.91 | thorax |
| 14 | Bg2/+;Mef2/+ | 15d | F | none | 20-29 | 21284.00 | 0.12 | 174569.99 | thorax |
| 14 | Bg2/+;Mef2/+ | 15d | F | none | 20-29 | 21696.00 | 0.14 | 157696.50 | thorax |
| 14 | Bg2/+;Mef2/+ | 15d | F | none | 20-29 | 23116.00 | 0.14 | 170688.64 | thorax |
| 14 | Bg2/+;Mef2/+ | 15d | F | none | 20-29 | 18108.00 | 0.12 | 148084.55 | thorax |
| 15 | Bg2/+;Mef2/+ | 15d | F | none | 20-29 | 19072.00 | 0.12 | 156757.05 | thorax |
| 15 | Bg2/+;Mef2/+ | 15d | F | none | 20-29 | 19426.00 | 0.12 | 159196.85 | thorax |

| Rep. | Genotype | Stage | Sex | Treatmen | °C | ß-gal ΔmOD/min/ml | Protein mg/ml | Specific Activity ΔmOD/min/mg | Whole/ dissected |
|---|---|---|---|---|---|---|---|---|---|
| 15 | Bg2/+;Mef2/+ | 15d | F | none | 20-29 | 24606.00 | 0.11 | 224587.35 | thorax |
| 15 | Bg2/+;Mef2/+ | 15d | F | none | 20-29 | 20534.00 | 0.12 | 167083.00 | thorax |
| 15 | Bg2/+;Mef2/+ | 15d | F | none | 20-29 | 20748.00 | 0.17 | 124354.14 | thorax |
| 12 | Bg2/+;Mef2/+ | 15d | F | none | 29-20 | 29460.00 | 0.08 | 369614.91 | thorax |
| 12 | Bg2/+;Mef2/+ | 15d | F | none | 29-20 | 23890.00 | 0.09 | 254479.02 | thorax |
| 12 | Bg2/+;Mef2/+ | 15d | F | none | 29-20 | 22690.00 | 0.10 | 224731.74 | thorax |
| 12 | Bg2/+;Mef2/+ | 15d | F | none | 29-20 | 24514.00 | 0.12 | 200424.41 | thorax |
| 12 | Bg2/+;Mef2/+ | 15d | F | none | 29-20 | 26940.00 | 0.09 | 294879.70 | thorax |
| 13 | Bg2/+;Mef2/+ | 15d | F | none | 29-20 | 24074.00 | 0.10 | 247437.20 | thorax |
| 13 | Bg2/+;Mef2/+ | 15d | F | none | 29-20 | 24724.00 | 0.08 | 300693.13 | thorax |
| 13 | Bg2/+;Mef2/+ | 15d | F | none | 29-20 | 18538.00 | 0.09 | 208766.40 | thorax |
| 13 | Bg2/+;Mef2/+ | 15d | F | none | 29-20 | 18492.00 | 0.07 | 261726.05 | thorax |
| 13 | Bg2/+;Mef2/+ | 15d | F | none | 29-20 | 23632.00 | 0.08 | 296019.23 | thorax |
| 14 | Bg2/+;Mef2/+ | 15d | F | none | 29-20 | 28082.00 | 0.11 | 261280.59 | thorax |
| 14 | Bg2/+;Mef2/+ | 15d | F | none | 29-20 | 22196.00 | 0.13 | 175118.79 | thorax |
| 14 | Bg2/+;Mef2/+ | 15d | F | none | 29-20 | 27892.00 | 0.11 | 263273.85 | thorax |
| 14 | Bg2/+;Mef2/+ | 15d | F | none | 29-20 | 29680.00 | 0.11 | 271360.00 | thorax |
| 14 | Bg2/+;Mef2/+ | 15d | F | none | 29-20 | 24540.00 | 0.13 | 191780.81 | thorax |
| 15 | Bg2/+;Mef2/+ | 15d | F | none | 29-20 | 29662.00 | 0.11 | 274941.54 | thorax |
| 15 | Bg2/+;Mef2/+ | 15d | F | none | 29-20 | 21844.00 | 0.11 | 194575.03 | thorax |
| 15 | Bg2/+;Mef2/+ | 15d | F | none | 29-20 | 27198.00 | 0.12 | 220936.36 | thorax |
| 15 | Bg2/+;Mef2/+ | 15d | F | none | 29-20 | 28210.00 | 0.10 | 293002.93 | thorax |
| 15 | Bg2/+;Mef2/+ | 15d | F | none | 29-20 | 27632.00 | 0.11 | 247425.40 | thorax |
| 12 | Bg2/+;Mef2/+ | 10d | M | none | 20 | 32984.00 | 0.11 | 298660.32 | thorax |
| 12 | Bg2/+;Mef2/+ | 10d | M | none | 20 | 17498.00 | 0.07 | 263510.42 | thorax |
| 12 | Bg2/+;Mef2/+ | 10d | M | none | 20 | 15868.00 | 0.09 | 185456.73 | thorax |
| 12 | Bg2/+;Mef2/+ | 10d | M | none | 20 | 21308.00 | 0.08 | 268312.26 | thorax |
| 13 | Bg2/+;Mef2/+ | 10d | M | none | 20 | 19710.00 | 0.10 | 203660.86 | thorax |
| 13 | Bg2/+;Mef2/+ | 10d | M | none | 20 | 21154.00 | 0.10 | 209453.73 | thorax |
| 13 | Bg2/+;Mef2/+ | 10d | M | none | 20 | 19692.00 | 0.08 | 234327.44 | thorax |
| 13 | Bg2/+;Mef2/+ | 10d | M | none | 20 | 21716.00 | 0.09 | 245563.78 | thorax |
| 13 | Bg2/+;Mef2/+ | 10d | M | none | 20 | 17782.00 | 0.09 | 192577.85 | thorax |
| 14 | Bg2/+;Mef2/+ | 10d | M | none | 20 | 12178.00 | 0.10 | 121442.18 | thorax |
| 14 | Bg2/+;Mef2/+ | 10d | M | none | 20 | 21814.00 | 0.08 | 268021.19 | thorax |
| 14 | Bg2/+;Mef2/+ | 10d | M | none | 20 | 21494.00 | 0.09 | 239656.06 | thorax |
| 14 | Bg2/+;Mef2/+ | 10d | M | none | 20 | 22420.00 | 0.12 | 181023.59 | thorax |
| 14 | Bg2/+;Mef2/+ | 10d | M | none | 20 | 17370.00 | 0.07 | 232364.53 | thorax |
| 15 | Bg2/+;Mef2/+ | 10d | M | none | 20 | 22534.00 | 0.10 | 230880.18 | thorax |
| 15 | Bg2/+;Mef2/+ | 10d | M | none | 20 | 18078.00 | 0.11 | 162923.82 | thorax |
| 15 | Bg2/+;Mef2/+ | 10d | M | none | 20 | 12076.00 | 0.07 | 163404.90 | thorax |

| Rep. | Genotype | Stage | Sex | Treatmen | °C | ß-gal ΔmOD/min/ ml | Protein mg/ml | Specific Activity ΔmOD/min/ mg | Whole/ dissected |
|---|---|---|---|---|---|---|---|---|---|
| 15 | Bg2/+;Mef2/+ | 10d | M | none | 20 | 16090.00 | 0.06 | 255181.75 | thorax |
| 15 | Bg2/+;Mef2/+ | 10d | M | none | 20 | 16162.00 | 0.07 | 234633.51 | thorax |
| 6 | Bg2/+;Mef2/+ | 10-11d | M | -RU486 | 25 | 31680.00 | 0.06 | 537460.00 | thorax |
| 6 | Bg2/+;Mef2/+ | 10-11d | M | -RU486 | 25 | 29315.00 | 0.05 | 566138.70 | thorax |
| 6 | Bg2/+;Mef2/+ | 10-11d | M | -RU486 | 25 | 31350.00 | 0.06 | 519241.02 | thorax |
| 6 | Bg2/+;Mef2/+ | 10-11d | M | -RU486 | 25 | 34460.00 | 0.06 | 563115.59 | thorax |
| 6 | Bg2/+;Mef2/+ | 10-11d | M | -RU486 | 25 | 35640.00 | 0.10 | 344825.82 | thorax |
| 7 | Bg2/+;Mef2/+ | 10-12d | M | -RU486 | 25 | 29540.00 | 0.09 | 331696.49 | thorax |
| 7 | Bg2/+;Mef2/+ | 10-12d | M | -RU486 | 25 | 34060.00 | 0.09 | 379423.01 | thorax |
| 7 | Bg2/+;Mef2/+ | 10-12d | M | -RU486 | 25 | 34640.00 | 0.08 | 449309.00 | thorax |
| 7 | Bg2/+;Mef2/+ | 10-12d | M | -RU486 | 25 | 32020.00 | 0.09 | 376045.73 | thorax |
| 7 | Bg2/+;Mef2/+ | 10-12d | M | -RU486 | 25 | 36120.00 | 0.10 | 365266.20 | thorax |
| 6 | Bg2/+;Mef2/+ | 10-11d | M | +RU486 | 25 | 33135.00 | 0.06 | 536973.83 | thorax |
| 6 | Bg2/+;Mef2/+ | 10-11d | M | +RU486 | 25 | 33605.00 | 0.06 | 519601.36 | thorax |
| 6 | Bg2/+;Mef2/+ | 10-11d | M | +RU486 | 25 | 34555.00 | 0.06 | 577216.17 | thorax |
| 6 | Bg2/+;Mef2/+ | 10-11d | M | +RU486 | 25 | 33380.00 | 0.06 | 529528.18 | thorax |
| 6 | Bg2/+;Mef2/+ | 10-11d | M | +RU486 | 25 | 32065.00 | 0.05 | 596836.53 | thorax |
| 7 | Bg2/+;Mef2/+ | 10-12d | M | +RU486 | 25 | 29780.00 | 0.06 | 467402.08 | thorax |
| 7 | Bg2/+;Mef2/+ | 10-12d | M | +RU486 | 25 | 28000.00 | 0.09 | 307054.55 | thorax |
| 7 | Bg2/+;Mef2/+ | 10-12d | M | +RU486 | 25 | 29740.00 | 0.09 | 340738.89 | thorax |
| 7 | Bg2/+;Mef2/+ | 10-12d | M | +RU486 | 25 | 37800.00 | 0.09 | 411318.56 | thorax |
| 7 | Bg2/+;Mef2/+ | 10-12d | M | +RU486 | 25 | 31540.00 | 0.11 | 289797.34 | thorax |
| 1 | Bg2/+;Mef2/+ | 0-24h | M | none | 25 | 26085.00 | 0.19 | 140469.86 | thorax |
| 1 | Bg2/+;Mef2/+ | 0-24h | M | none | 25 | 26775.00 | 0.19 | 144448.36 | thorax |
| 1 | Bg2/+;Mef2/+ | 0-24h | M | none | 25 | 23535.00 | 0.15 | 158867.96 | thorax |
| 1 | Bg2/+;Mef2/+ | 0-24h | M | none | 25 | 22725.00 | 0.15 | 150035.69 | thorax |
| 1 | Bg2/+;Mef2/+ | 0-24h | M | none | 25 | 27855.00 | 0.14 | 201263.14 | thorax |
| 8 | Bg2/+;Mef2/+ | 0-4d | M | none | 25 | 43940.00 | 0.09 | 467529.81 | thorax |
| 8 | Bg2/+;Mef2/+ | 0-4d | M | none | 25 | 41000.00 | 0.14 | 283114.48 | thorax |
| 8 | Bg2/+;Mef2/+ | 0-4d | M | none | 25 | 32940.00 | 0.13 | 263173.26 | thorax |
| 9 | Bg2/+;Mef2/+ | 1-3d | M | none | 25 | 45246.67 | 0.12 | 383704.71 | thorax |
| 9 | Bg2/+;Mef2/+ | 1-3d | M | none | 25 | 41050.00 | 0.13 | 328122.64 | thorax |
| 9 | Bg2/+;Mef2/+ | 1-3d | M | none | 25 | 32188.57 | 0.17 | 193540.43 | thorax |
| 9 | Bg2/+;Mef2/+ | 1-3d | M | none | 25 | 35440.00 | 0.12 | 288147.90 | thorax |
| 9 | Bg2/+;Mef2/+ | 1-3d | M | none | 25 | 35720.00 | 0.13 | 267025.34 | thorax |
| 1 | Bg2/+;Mef2/+ | 10-11d | M | none | 25 | 11130.00 | 0.19 | 58813.16 | thorax |
| 1 | Bg2/+;Mef2/+ | 10-11d | M | none | 25 | 15110.00 | 0.10 | 145877.60 | thorax |
| 1 | Bg2/+;Mef2/+ | 10-11d | M | none | 25 | 12740.00 | 0.15 | 82853.33 | thorax |
| 1 | Bg2/+;Mef2/+ | 10-11d | M | none | 25 | 9050.00 | 0.12 | 76818.56 | thorax |
| 1 | Bg2/+;Mef2/+ | 10-11d | M | none | 25 | 12620.00 | 0.10 | 129418.00 | thorax |

| Rep. | Genotype | Stage | Sex | Treatmen | °C | ß-gal ΔmOD/min/ml | Protein mg/ml | Specific Activity ΔmOD/min/mg | Whole/ dissected |
|---|---|---|---|---|---|---|---|---|---|
| 1 | Bg2/+;Mef2/+ | 3-5d | M | none | 25 | 21715.00 | 0.16 | 133713.05 | thorax |
| 1 | Bg2/+;Mef2/+ | 3-5d | M | none | 25 | 17360.00 | 0.13 | 130037.45 | thorax |
| 1 | Bg2/+;Mef2/+ | 3-5d | M | none | 25 | 19880.00 | 0.27 | 74652.65 | thorax |
| 1 | Bg2/+;Mef2/+ | 3-5d | M | none | 25 | 21020.00 | 0.12 | 170894.31 | thorax |
| 1 | Bg2/+;Mef2/+ | 3-5d | M | none | 25 | 21965.00 | 0.22 | 98497.76 | thorax |
| 12 | Bg2/+;Mef2/+ | 15d | M | none | 29 | 15666.00 | 0.10 | 158934.56 | thorax |
| 12 | Bg2/+;Mef2/+ | 15d | M | none | 29 | 19574.00 | 0.09 | 216992.88 | thorax |
| 12 | Bg2/+;Mef2/+ | 15d | M | none | 29 | 16838.00 | 0.08 | 199945.66 | thorax |
| 12 | Bg2/+;Mef2/+ | 15d | M | none | 29 | 15602.00 | 0.10 | 150112.19 | thorax |
| 12 | Bg2/+;Mef2/+ | 15d | M | none | 29 | 15244.00 | 0.07 | 206454.17 | thorax |
| 13 | Bg2/+;Mef2/+ | 15d | M | none | 29 | 18434.00 | 0.11 | 165535.84 | thorax |
| 13 | Bg2/+;Mef2/+ | 15d | M | none | 29 | 14282.00 | 0.10 | 137589.63 | thorax |
| 13 | Bg2/+;Mef2/+ | 15d | M | none | 29 | 15506.00 | 0.12 | 128794.26 | thorax |
| 13 | Bg2/+;Mef2/+ | 15d | M | none | 29 | 13394.00 | 0.10 | 128591.60 | thorax |
| 13 | Bg2/+;Mef2/+ | 15d | M | none | 29 | 16792.00 | 0.11 | 154514.04 | thorax |
| 14 | Bg2/+;Mef2/+ | 15d | M | none | 29 | 11626.00 | 0.09 | 125879.20 | thorax |
| 14 | Bg2/+;Mef2/+ | 15d | M | none | 29 | 6206.00 | 0.07 | 84873.29 | thorax |
| 14 | Bg2/+;Mef2/+ | 15d | M | none | 29 | 5882.00 | 0.08 | 77424.50 | thorax |
| 14 | Bg2/+;Mef2/+ | 15d | M | none | 29 | 4316.00 | 0.06 | 68205.56 | thorax |
| 14 | Bg2/+;Mef2/+ | 15d | M | none | 29 | 3362.00 | 0.07 | 47303.93 | thorax |
| 15 | Bg2/+;Mef2/+ | 15d | M | none | 29 | 4334.00 | 0.07 | 65715.26 | thorax |
| 15 | Bg2/+;Mef2/+ | 15d | M | none | 29 | 5810.00 | 0.08 | 75722.21 | thorax |
| 15 | Bg2/+;Mef2/+ | 15d | M | none | 29 | 2896.00 | 0.05 | 59717.70 | thorax |
| 15 | Bg2/+;Mef2/+ | 15d | M | none | 29 | 4508.00 | 0.07 | 64808.99 | thorax |
| 15 | Bg2/+;Mef2/+ | 15d | M | none | 29 | 9502.00 | 0.07 | 139462.03 | thorax |
| 12 | Bg2/+;Mef2/+ | 15d | M | none | 20-29 | 22128.00 | 0.12 | 178842.52 | thorax |
| 12 | Bg2/+;Mef2/+ | 15d | M | none | 20-29 | 26016.00 | 0.10 | 256517.31 | thorax |
| 12 | Bg2/+;Mef2/+ | 15d | M | none | 20-29 | 16748.00 | 0.10 | 162813.11 | thorax |
| 12 | Bg2/+;Mef2/+ | 15d | M | none | 20-29 | 21432.00 | 0.11 | 195832.00 | thorax |
| 12 | Bg2/+;Mef2/+ | 15d | M | none | 20-29 | 17770.00 | 0.11 | 159559.35 | thorax |
| 13 | Bg2/+;Mef2/+ | 15d | M | none | 20-29 | 21256.00 | 0.07 | 296078.69 | thorax |
| 13 | Bg2/+;Mef2/+ | 15d | M | none | 20-29 | 21154.00 | 0.10 | 220488.65 | thorax |
| 13 | Bg2/+;Mef2/+ | 15d | M | none | 20-29 | 13688.00 | 0.08 | 180836.95 | thorax |
| 13 | Bg2/+;Mef2/+ | 15d | M | none | 20-29 | 20284.00 | 0.11 | 189206.76 | thorax |
| 13 | Bg2/+;Mef2/+ | 15d | M | none | 20-29 | 19020.00 | 0.09 | 217742.26 | thorax |
| 14 | Bg2/+;Mef2/+ | 15d | M | none | 20-29 | 17218.00 | 0.10 | 169279.48 | thorax |
| 14 | Bg2/+;Mef2/+ | 15d | M | none | 20-29 | 16288.00 | 0.10 | 166166.37 | thorax |
| 14 | Bg2/+;Mef2/+ | 15d | M | none | 20-29 | 18388.00 | 0.09 | 209722.59 | thorax |
| 14 | Bg2/+;Mef2/+ | 15d | M | none | 20-29 | 14630.00 | 0.09 | 158852.41 | thorax |
| 14 | Bg2/+;Mef2/+ | 15d | M | none | 20-29 | 12424.00 | 0.08 | 165231.67 | thorax |

| Rep. | Genotype | Stage | Sex | Treatmen | °C | ß-gal ΔmOD/min/ ml | Protein mg/ml | Specific Activity ΔmOD/min/ mg | Whole/ dissected |
|---|---|---|---|---|---|---|---|---|---|
| 15 | Bg2/+;Mef2/+ | 15d | M | none | 20-29 | 11174.00 | 0.10 | 115280.80 | thorax |
| 15 | Bg2/+;Mef2/+ | 15d | M | none | 20-29 | 16402.00 | 0.09 | 183541.81 | thorax |
| 15 | Bg2/+;Mef2/+ | 15d | M | none | 20-29 | 14740.00 | 0.07 | 216068.51 | thorax |
| 15 | Bg2/+;Mef2/+ | 15d | M | none | 20-29 | 18130.00 | 0.11 | 169512.02 | thorax |
| 15 | Bg2/+;Mef2/+ | 15d | M | none | 20-29 | 21584.00 | 0.11 | 202496.49 | thorax |
| 12 | Bg2/+;Mef2/+ | 15d | M | none | 29-20 | 17480.00 | 0.08 | 208150.96 | thorax |
| 12 | Bg2/+;Mef2/+ | 15d | M | none | 29-20 | 9132.00 | 0.06 | 164621.83 | thorax |
| 12 | Bg2/+;Mef2/+ | 15d | M | none | 29-20 | 14142.00 | 0.06 | 218147.06 | thorax |
| 12 | Bg2/+;Mef2/+ | 15d | M | none | 29-20 | 17470.00 | 0.07 | 245984.51 | thorax |
| 13 | Bg2/+;Mef2/+ | 15d | M | none | 29-20 | 18576.00 | 0.09 | 210312.47 | thorax |
| 13 | Bg2/+;Mef2/+ | 15d | M | none | 29-20 | 18008.00 | 0.07 | 257218.41 | thorax |
| 13 | Bg2/+;Mef2/+ | 15d | M | none | 29-20 | 14992.00 | 0.05 | 301535.21 | thorax |
| 13 | Bg2/+;Mef2/+ | 15d | M | none | 29-20 | 18258.00 | 0.07 | 252857.75 | thorax |
| 13 | Bg2/+;Mef2/+ | 15d | M | none | 29-20 | 19468.00 | 0.08 | 229305.44 | thorax |
| 14 | Bg2/+;Mef2/+ | 15d | M | none | 29-20 | 14536.00 | 0.07 | 208279.20 | thorax |
| 14 | Bg2/+;Mef2/+ | 15d | M | none | 29-20 | 18346.00 | 0.09 | 212663.84 | thorax |
| 14 | Bg2/+;Mef2/+ | 15d | M | none | 29-20 | 20014.00 | 0.09 | 223007.67 | thorax |
| 14 | Bg2/+;Mef2/+ | 15d | M | none | 29-20 | 23626.00 | 0.09 | 275238.76 | thorax |
| 14 | Bg2/+;Mef2/+ | 15d | M | none | 29-20 | 16026.00 | 0.09 | 177720.71 | thorax |
| 15 | Bg2/+;Mef2/+ | 15d | M | none | 29-20 | 19578.00 | 0.07 | 264002.58 | thorax |
| 15 | Bg2/+;Mef2/+ | 15d | M | none | 29-20 | 21362.00 | 0.08 | 262244.73 | thorax |
| 15 | Bg2/+;Mef2/+ | 15d | M | none | 29-20 | 17848.00 | 0.10 | 180322.87 | thorax |
| 15 | Bg2/+;Mef2/+ | 15d | M | none | 29-20 | 18514.00 | 0.08 | 233687.82 | thorax |
| 15 | Bg2/+;Mef2/+ | 15d | M | none | 29-20 | 20944.00 | 0.08 | 266964.35 | thorax |
| 12 | Bg2/+;Mef2/+ | LP | ND | none | 20 | 16996.00 | 0.32 | 53905.90 | whole |
| 12 | Bg2/+;Mef2/+ | LP | ND | none | 20 | 27488.00 | 0.67 | 41263.06 | whole |
| 12 | Bg2/+;Mef2/+ | LP | ND | none | 20 | 24262.00 | 0.52 | 46594.96 | whole |
| 12 | Bg2/+;Mef2/+ | LP | ND | none | 20 | 19426.00 | 0.19 | 100967.48 | whole |
| 13 | Bg2/+;Mef2/+ | LP | ND | none | 20 | 15788.00 | 0.37 | 42480.62 | whole |
| 13 | Bg2/+;Mef2/+ | LP | ND | none | 20 | 28286.00 | 0.34 | 83851.96 | whole |
| 13 | Bg2/+;Mef2/+ | LP | ND | none | 20 | 30074.00 | 0.31 | 96581.73 | whole |
| 13 | Bg2/+;Mef2/+ | LP | ND | none | 20 | 29854.00 | 0.28 | 105675.79 | whole |
| 13 | Bg2/+;Mef2/+ | LP | ND | none | 20 | 26988.00 | 0.26 | 105135.86 | whole |
| 14 | Bg2/+;Mef2/+ | LP | ND | none | 20 | 26472.00 | 0.24 | 108145.47 | whole |
| 14 | Bg2/+;Mef2/+ | LP | ND | none | 20 | 15628.00 | 0.21 | 73675.60 | whole |
| 14 | Bg2/+;Mef2/+ | LP | ND | none | 20 | 14454.00 | 0.19 | 78057.11 | whole |
| 14 | Bg2/+;Mef2/+ | LP | ND | none | 20 | 23470.00 | 0.38 | 60982.07 | whole |
| 14 | Bg2/+;Mef2/+ | LP | ND | none | 20 | 16002.00 | 0.22 | 71424.54 | whole |
| 15 | Bg2/+;Mef2/+ | LP | ND | none | 20 | 11272.00 | 0.24 | 47164.07 | whole |
| 15 | Bg2/+;Mef2/+ | LP | ND | none | 20 | 13874.00 | 0.35 | 39863.73 | whole |

| Rep. | Genotype | Stage | Sex | Treatmen | °C | ß-gal ΔmOD/min/ml | Protein mg/ml | Specific Activity ΔmOD/min/mg | Whole/ dissected |
|---|---|---|---|---|---|---|---|---|---|
| 15 | Bg2/+;Mef2/+ | LP | ND | none | 20 | 15850.00 | 0.22 | 72738.62 | whole |
| 15 | Bg2/+;Mef2/+ | LP | ND | none | 20 | 13920.00 | 0.25 | 54626.81 | whole |
| 15 | Bg2/+;Mef2/+ | LP | ND | none | 20 | 18082.00 | 0.25 | 71650.33 | whole |
| 6 | Bg2/+;Mef2/+ | EP | ND | -RU486 | 25 | 7720.00 | 1.58 | 4892.25 | whole |
| 6 | Bg2/+;Mef2/+ | EP | ND | -RU486 | 25 | 8525.00 | 1.53 | 5554.86 | whole |
| 6 | Bg2/+;Mef2/+ | EP | ND | -RU486 | 25 | 11045.00 | 0.80 | 13740.52 | whole |
| 6 | Bg2/+;Mef2/+ | EP | ND | -RU486 | 25 | 5790.00 | 1.25 | 4625.00 | whole |
| 6 | Bg2/+;Mef2/+ | EP | ND | -RU486 | 25 | 8475.00 | 0.92 | 9171.13 | whole |
| 7 | Bg2/+;Mef2/+ | EP | ND | -RU486 | 25 | 8310.00 | 1.58 | 5266.14 | whole |
| 7 | Bg2/+;Mef2/+ | EP | ND | -RU486 | 25 | 9720.00 | 1.53 | 6333.51 | whole |
| 7 | Bg2/+;Mef2/+ | EP | ND | -RU486 | 25 | 4945.00 | 0.80 | 6151.82 | whole |
| 7 | Bg2/+;Mef2/+ | EP | ND | -RU486 | 25 | 6050.00 | 1.25 | 4832.68 | whole |
| 7 | Bg2/+;Mef2/+ | EP | ND | -RU486 | 25 | 3805.00 | 0.92 | 4117.54 | whole |
| 6 | Bg2/+;Mef2/+ | EP | ND | +RU486 | 25 | 7300.00 | 0.80 | 9083.94 | whole |
| 6 | Bg2/+;Mef2/+ | EP | ND | +RU486 | 25 | 4415.00 | 1.06 | 4184.48 | whole |
| 6 | Bg2/+;Mef2/+ | EP | ND | +RU486 | 25 | 9195.00 | 0.70 | 13120.74 | whole |
| 6 | Bg2/+;Mef2/+ | EP | ND | +RU486 | 25 | 8560.00 | 1.31 | 6539.98 | whole |
| 6 | Bg2/+;Mef2/+ | EP | ND | +RU486 | 25 | 5610.00 | 0.74 | 7614.49 | whole |
| 7 | Bg2/+;Mef2/+ | EP | ND | +RU486 | 25 | 7830.00 | 0.80 | 9743.45 | whole |
| 7 | Bg2/+;Mef2/+ | EP | ND | +RU486 | 25 | 10955.00 | 1.06 | 10383.02 | whole |
| 7 | Bg2/+;Mef2/+ | EP | ND | +RU486 | 25 | 10905.00 | 0.70 | 15560.81 | whole |
| 7 | Bg2/+;Mef2/+ | EP | ND | +RU486 | 25 | 8820.00 | 1.31 | 6738.62 | whole |
| 7 | Bg2/+;Mef2/+ | EP | ND | +RU486 | 25 | 10090.00 | 0.74 | 13695.22 | whole |
| 8 | Bg2/+;Mef2/+ | EP | ND | none | 25 | 7513.33 | 0.40 | 18956.88 | whole |
| 8 | Bg2/+;Mef2/+ | EP | ND | none | 25 | 6513.33 | 0.35 | 18543.90 | whole |
| 8 | Bg2/+;Mef2/+ | EP | ND | none | 25 | 6485.00 | 0.41 | 15680.53 | whole |
| 8 | Bg2/+;Mef2/+ | EP | ND | none | 25 | 7713.33 | 0.47 | 16318.61 | whole |
| 8 | Bg2/+;Mef2/+ | EP | ND | none | 25 | 6473.33 | 0.37 | 17428.02 | whole |
| 8 | Bg2/+;Mef2/+ | EP | ND | none | 25 | 9237.50 | 0.43 | 21611.39 | whole |
| 9 | Bg2/+;Mef2/+ | EP | ND | none | 25 | 6150.00 | 0.46 | 13290.65 | whole |
| 9 | Bg2/+;Mef2/+ | EP | ND | none | 25 | 17625.00 | 0.57 | 30875.91 | whole |
| 9 | Bg2/+;Mef2/+ | EP | ND | none | 25 | 5726.67 | 0.26 | 21735.91 | whole |
| 9 | Bg2/+;Mef2/+ | EP | ND | none | 25 | 4553.33 | 0.31 | 14624.77 | whole |
| 9 | Bg2/+;Mef2/+ | EP | ND | none | 25 | 13217.50 | 0.69 | 19247.89 | whole |
| 2 | Bg2/+;Mef2/+ | L3 | ND | none | 25 | 9440.00 | 0.44 | 21474.82 | whole |
| 2 | Bg2/+;Mef2/+ | L3 | ND | none | 25 | 5240.00 | 0.33 | 15784.57 | whole |
| 2 | Bg2/+;Mef2/+ | L3 | ND | none | 25 | 23040.00 | 0.69 | 33389.93 | whole |
| 2 | Bg2/+;Mef2/+ | L3 | ND | none | 25 | 16360.00 | 0.53 | 30806.65 | whole |
| 2 | Bg2/+;Mef2/+ | L3 | ND | none | 25 | 2360.00 | 0.31 | 7560.56 | whole |
| 3 | Bg2/+;Mef2/+ | L3 | ND | none | 25 | 15080.00 | 0.36 | 42177.83 | whole |

| Rep. | Genotype | Stage | Sex | Treatmen | °C | ß-gal ΔmOD/min/ ml | Protein mg/ml | Specific Activity ΔmOD/min/ mg | Whole/ dissected |
|---|---|---|---|---|---|---|---|---|---|
| 3 | Bg2/+;Mef2/+ | L3 | ND | none | 25 | 22200.00 | 0.51 | 43295.10 | whole |
| 3 | Bg2/+;Mef2/+ | L3 | ND | none | 25 | 5400.00 | 0.51 | 10497.17 | whole |
| 3 | Bg2/+;Mef2/+ | L3 | ND | none | 25 | 11160.00 | 0.50 | 22486.51 | whole |
| 3 | Bg2/+;Mef2/+ | L3 | ND | none | 25 | 18280.00 | 0.52 | 35071.53 | whole |
| 1 | Bg2/+;Mef2/+ | LP | ND | none | 25 | 39100.00 | 0.24 | 164674.59 | whole |
| 1 | Bg2/+;Mef2/+ | LP | ND | none | 25 | 9290.00 | 0.11 | 87138.33 | whole |
| 1 | Bg2/+;Mef2/+ | LP | ND | none | 25 | 7750.00 | 0.22 | 35732.57 | whole |
| 1 | Bg2/+;Mef2/+ | LP | ND | none | 25 | 12990.00 | 0.11 | 114983.95 | whole |
| 1 | Bg2/+;Mef2/+ | LP | ND | none | 25 | 14900.00 | 0.07 | 219695.45 | whole |
| 8 | Bg2/+;Mef2/+ | LP | ND | none | 25 | 53270.00 | 0.20 | 272793.72 | whole |
| 8 | Bg2/+;Mef2/+ | LP | ND | none | 25 | 88905.00 | 0.39 | 229653.69 | whole |
| 8 | Bg2/+;Mef2/+ | LP | ND | none | 25 | 55253.33 | 0.33 | 167917.30 | whole |
| 8 | Bg2/+;Mef2/+ | LP | ND | none | 25 | 45600.00 | 0.35 | 130324.77 | whole |
| 9 | Bg2/+;Mef2/+ | LP | ND | none | 25 | 56120.00 | 0.38 | 145861.44 | whole |
| 9 | Bg2/+;Mef2/+ | LP | ND | none | 25 | 27731.43 | 0.46 | 59985.13 | whole |
| 9 | Bg2/+;Mef2/+ | LP | ND | none | 25 | 39285.00 | 0.47 | 82753.07 | whole |
| 9 | Bg2/+;Mef2/+ | LP | ND | none | 25 | 53480.00 | 0.39 | 135788.60 | whole |
| 9 | Bg2/+;Mef2/+ | LP | ND | none | 25 | 51833.33 | 0.32 | 163282.07 | whole |
| 9 | Bg2/+;Mef2/+ | LP | ND | none | 25 | 41710.00 | 0.46 | 90362.97 | whole |
| 12 | Bg2/+;Mef2/+ | LP | ND | none | 29 | 48475.00 | 0.36 | 135422.22 | whole |
| 12 | Bg2/+;Mef2/+ | LP | ND | none | 29 | 58040.00 | 0.33 | 178010.08 | whole |
| 12 | Bg2/+;Mef2/+ | LP | ND | none | 29 | 36540.00 | 0.27 | 134195.06 | whole |
| 12 | Bg2/+;Mef2/+ | LP | ND | none | 29 | 56170.00 | 0.31 | 183508.27 | whole |
| 12 | Bg2/+;Mef2/+ | LP | ND | none | 29 | 51660.00 | 0.31 | 166946.44 | whole |
| 13 | Bg2/+;Mef2/+ | LP | ND | none | 29 | 37080.00 | 0.21 | 174580.95 | whole |
| 13 | Bg2/+;Mef2/+ | LP | ND | none | 29 | 49265.00 | 0.33 | 150760.12 | whole |
| 13 | Bg2/+;Mef2/+ | LP | ND | none | 29 | 65630.00 | 0.41 | 158788.97 | whole |
| 13 | Bg2/+;Mef2/+ | LP | ND | none | 29 | 46375.00 | 0.31 | 149515.27 | whole |
| 13 | Bg2/+;Mef2/+ | LP | ND | none | 29 | 54700.00 | 0.40 | 136036.52 | whole |
| 14 | Bg2/+;Mef2/+ | LP | ND | none | 29 | 38315.00 | 0.33 | 117263.97 | whole |
| 14 | Bg2/+;Mef2/+ | LP | ND | none | 29 | 7305.00 | 0.11 | 69481.93 | whole |
| 14 | Bg2/+;Mef2/+ | LP | ND | none | 29 | 12995.00 | 0.15 | 86305.04 | whole |
| 14 | Bg2/+;Mef2/+ | LP | ND | none | 29 | 33275.00 | 0.29 | 113245.30 | whole |
| 14 | Bg2/+;Mef2/+ | LP | ND | none | 29 | 22040.00 | 0.14 | 152587.10 | whole |
| 15 | Bg2/+;Mef2/+ | LP | ND | none | 29 | 7600.00 | 0.16 | 48337.63 | whole |
| 15 | Bg2/+;Mef2/+ | LP | ND | none | 29 | 27190.00 | 0.31 | 89004.56 | whole |
| 15 | Bg2/+;Mef2/+ | LP | ND | none | 29 | 21425.00 | 0.18 | 118300.00 | whole |
| 15 | Bg2/+;Mef2/+ | LP | ND | none | 29 | 39880.00 | 0.25 | 159790.14 | whole |
| 15 | Bg2/+;Mef2/+ | LP | ND | none | 29 | 51945.00 | 0.33 | 157838.26 | whole |
| 12 | Bg2/+;Mef2/3.3+1077 | 10d | F | none | 20 | 13886.00 | 0.11 | 124787.39 | thorax |

| Rep. | Genotype | Stage | Sex | Treatmen | °C | ß-gal ΔmOD/min/ ml | Protein mg/ml | Specific Activity ΔmOD/min/ mg | Whole/ dissected |
|---|---|---|---|---|---|---|---|---|---|
| 12 | Bg2/+;Mef2/3.3+1077 | 10d | F | none | 20 | 17706.00 | 0.11 | 154772.63 | thorax |
| 12 | Bg2/+;Mef2/3.3+1077 | 10d | F | none | 20 | 18248.00 | 0.11 | 164768.28 | thorax |
| 12 | Bg2/+;Mef2/3.3+1077 | 10d | F | none | 20 | 17302.00 | 0.12 | 141452.09 | thorax |
| 12 | Bg2/+;Mef2/3.3+1077 | 10d | F | none | 20 | 15322.00 | 0.14 | 108554.63 | thorax |
| 13 | Bg2/+;Mef2/3.3+1077 | 10d | F | none | 20 | 16512.00 | 0.11 | 147049.29 | thorax |
| 13 | Bg2/+;Mef2/3.3+1077 | 10d | F | none | 20 | 18248.00 | 0.11 | 173157.98 | thorax |
| 13 | Bg2/+;Mef2/3.3+1077 | 10d | F | none | 20 | 16144.00 | 0.11 | 140794.01 | thorax |
| 13 | Bg2/+;Mef2/3.3+1077 | 10d | F | none | 20 | 14114.00 | 0.11 | 127238.66 | thorax |
| 13 | Bg2/+;Mef2/3.3+1077 | 10d | F | none | 20 | 14528.00 | 0.12 | 124786.02 | thorax |
| 14 | Bg2/+;Mef2/3.3+1077 | 10d | F | none | 20 | 22054.00 | 0.10 | 217776.15 | thorax |
| 14 | Bg2/+;Mef2/3.3+1077 | 10d | F | none | 20 | 21350.00 | 0.14 | 151019.70 | thorax |
| 14 | Bg2/+;Mef2/3.3+1077 | 10d | F | none | 20 | 19266.00 | 0.11 | 172345.31 | thorax |
| 14 | Bg2/+;Mef2/3.3+1077 | 10d | F | none | 20 | 18678.00 | 0.14 | 137264.18 | thorax |
| 14 | Bg2/+;Mef2/3.3+1077 | 10d | F | none | 20 | 19220.00 | 0.13 | 152681.27 | thorax |
| 15 | Bg2/+;Mef2/3.3+1077 | 10d | F | none | 20 | 15504.00 | 0.11 | 143547.70 | thorax |
| 15 | Bg2/+;Mef2/3.3+1077 | 10d | F | none | 20 | 14848.00 | 0.12 | 127471.19 | thorax |
| 15 | Bg2/+;Mef2/3.3+1077 | 10d | F | none | 20 | 16438.00 | 0.08 | 216117.43 | thorax |
| 15 | Bg2/+;Mef2/3.3+1077 | 10d | F | none | 20 | 21516.00 | 0.14 | 152781.24 | thorax |
| 15 | Bg2/+;Mef2/3.3+1077 | 10d | F | none | 20 | 23512.00 | 0.13 | 181487.49 | thorax |
| 8 | Bg2/+;Mef2/3.3,DJ1077 | 0-4d | F | none | 25 | 41286.67 | 0.17 | 247068.96 | thorax |
| 8 | Bg2/+;Mef2/3.3,DJ1077 | 0-4d | F | none | 25 | 47060.00 | 0.15 | 323732.99 | thorax |
| 8 | Bg2/+;Mef2/3.3,DJ1077 | 0-4d | F | none | 25 | 56806.67 | 0.14 | 416582.22 | thorax |
| 8 | Bg2/+;Mef2/3.3,DJ1077 | 0-4d | F | none | 25 | 50080.00 | 0.14 | 354963.92 | thorax |
| 8 | Bg2/+;Mef2/3.3,DJ1077 | 0-4d | F | none | 25 | 44063.33 | 0.15 | 303577.03 | thorax |
| 9 | Bg2/+;Mef2/3.3,DJ1077 | 1-3d | F | none | 25 | 36051.11 | 0.13 | 279662.06 | thorax |
| 9 | Bg2/+;Mef2/3.3,DJ1077 | 1-3d | F | none | 25 | 38020.00 | 0.13 | 297127.40 | thorax |
| 9 | Bg2/+;Mef2/3.3,DJ1077 | 1-3d | F | none | 25 | 38666.67 | 0.12 | 317106.87 | thorax |
| 9 | Bg2/+;Mef2/3.3,DJ1077 | 1-3d | F | none | 25 | 50160.00 | 0.15 | 332432.94 | thorax |
| 9 | Bg2/+;Mef2/3.3,DJ1077 | 1-3d | F | none | 25 | 36583.33 | 0.11 | 347615.13 | thorax |
| 12 | Bg2/+;Mef2/3.3+1077 | 15d | F | none | 29 | 29776.00 | 0.11 | 275856.98 | thorax |
| 12 | Bg2/+;Mef2/3.3+1077 | 15d | F | none | 29 | 17024.00 | 0.12 | 143103.70 | thorax |
| 12 | Bg2/+;Mef2/3.3+1077 | 15d | F | none | 29 | 18662.00 | 0.11 | 163341.06 | thorax |
| 12 | Bg2/+;Mef2/3.3+1077 | 15d | F | none | 29 | 26244.00 | 0.10 | 256618.43 | thorax |
| 13 | Bg2/+;Mef2/3.3+1077 | 15d | F | none | 29 | 27300.00 | 0.12 | 220091.15 | thorax |
| 13 | Bg2/+;Mef2/3.3+1077 | 15d | F | none | 29 | 27938.00 | 0.15 | 190853.57 | thorax |
| 13 | Bg2/+;Mef2/3.3+1077 | 15d | F | none | 29 | 23520.00 | 0.09 | 275584.82 | thorax |
| 13 | Bg2/+;Mef2/3.3+1077 | 15d | F | none | 29 | 25502.00 | 0.12 | 212896.42 | thorax |
| 13 | Bg2/+;Mef2/3.3+1077 | 15d | F | none | 29 | 25242.00 | 0.10 | 247818.18 | thorax |
| 14 | Bg2/+;Mef2/3.3+1077 | 15d | F | none | 29 | 24508.00 | 0.17 | 142645.37 | thorax |
| 14 | Bg2/+;Mef2/3.3+1077 | 15d | F | none | 29 | 16822.00 | 0.12 | 140541.16 | thorax |

| Rep. | Genotype | Stage | Sex | Treatmen | °C | ß-gal ΔmOD/min/ ml | Protein mg/ml | Specific Activity ΔmOD/min/ mg | Whole/ dissected |
|---|---|---|---|---|---|---|---|---|---|
| 14 | Bg2/+;Mef2/3.3+1077 | 15d | F | none | 29 | 15088.00 | 0.15 | 100909.94 | thorax |
| 14 | Bg2/+;Mef2/3.3+1077 | 15d | F | none | 29 | 19674.00 | 0.12 | 157970.01 | thorax |
| 14 | Bg2/+;Mef2/3.3+1077 | 15d | F | none | 29 | 15428.00 | 0.11 | 144091.33 | thorax |
| 15 | Bg2/+;Mef2/3.3+1077 | 15d | F | none | 29 | 16658.00 | 0.13 | 127845.23 | thorax |
| 15 | Bg2/+;Mef2/3.3+1077 | 15d | F | none | 29 | 11624.00 | 0.11 | 102976.96 | thorax |
| 15 | Bg2/+;Mef2/3.3+1077 | 15d | F | none | 29 | 15258.00 | 0.11 | 135555.02 | thorax |
| 15 | Bg2/+;Mef2/3.3+1077 | 15d | F | none | 29 | 11856.00 | 0.11 | 107470.81 | thorax |
| 15 | Bg2/+;Mef2/3.3+1077 | 15d | F | none | 29 | 19146.00 | 0.11 | 170790.76 | thorax |
| 12 | Bg2/+;Mef2/3.3+1077 | 15d | F | none | 20-29 | 13072.00 | 0.14 | 90308.95 | thorax |
| 12 | Bg2/+;Mef2/3.3+1077 | 15d | F | none | 20-29 | 20064.00 | 0.16 | 123501.71 | thorax |
| 12 | Bg2/+;Mef2/3.3+1077 | 15d | F | none | 20-29 | 17886.00 | 0.14 | 125928.38 | thorax |
| 12 | Bg2/+;Mef2/3.3+1077 | 15d | F | none | 20-29 | 19558.00 | 0.16 | 125799.51 | thorax |
| 12 | Bg2/+;Mef2/3.3+1077 | 15d | F | none | 20-29 | 20952.00 | 0.13 | 157605.24 | thorax |
| 13 | Bg2/+;Mef2/3.3+1077 | 15d | F | none | 20-29 | 15376.00 | 0.15 | 104898.49 | thorax |
| 13 | Bg2/+;Mef2/3.3+1077 | 15d | F | none | 20-29 | 15190.00 | 0.12 | 123405.36 | thorax |
| 13 | Bg2/+;Mef2/3.3+1077 | 15d | F | none | 20-29 | 19458.00 | 0.11 | 175396.06 | thorax |
| 13 | Bg2/+;Mef2/3.3+1077 | 15d | F | none | 20-29 | 16298.00 | 0.10 | 165932.80 | thorax |
| 13 | Bg2/+;Mef2/3.3+1077 | 15d | F | none | 20-29 | 16632.00 | 0.13 | 129882.96 | thorax |
| 14 | Bg2/+;Mef2/3.3+1077 | 15d | F | none | 20-29 | 18342.00 | 0.15 | 124890.28 | thorax |
| 14 | Bg2/+;Mef2/3.3+1077 | 15d | F | none | 20-29 | 16910.00 | 0.17 | 102299.65 | thorax |
| 14 | Bg2/+;Mef2/3.3+1077 | 15d | F | none | 20-29 | 19312.00 | 0.17 | 111955.37 | thorax |
| 14 | Bg2/+;Mef2/3.3+1077 | 15d | F | none | 20-29 | 21440.00 | 0.16 | 133117.86 | thorax |
| 14 | Bg2/+;Mef2/3.3+1077 | 15d | F | none | 20-29 | 18176.00 | 0.14 | 126246.76 | thorax |
| 15 | Bg2/+;Mef2/3.3+1077 | 15d | F | none | 20-29 | 20338.00 | 0.17 | 119346.24 | thorax |
| 15 | Bg2/+;Mef2/3.3+1077 | 15d | F | none | 20-29 | 18658.00 | 0.12 | 162029.56 | thorax |
| 15 | Bg2/+;Mef2/3.3+1077 | 15d | F | none | 20-29 | 19388.00 | 0.13 | 150408.78 | thorax |
| 15 | Bg2/+;Mef2/3.3+1077 | 15d | F | none | 20-29 | 20078.00 | 0.12 | 167282.35 | thorax |
| 15 | Bg2/+;Mef2/3.3+1077 | 15d | F | none | 20-29 | 16168.00 | 0.15 | 107864.07 | thorax |
| 12 | Bg2/+;Mef2/3.3+1077 | 15d | F | none | 29-20 | 26100.00 | 0.12 | 225596.46 | thorax |
| 12 | Bg2/+;Mef2/3.3+1077 | 15d | F | none | 29-20 | 25220.00 | 0.11 | 240046.03 | thorax |
| 12 | Bg2/+;Mef2/3.3+1077 | 15d | F | none | 29-20 | 29312.00 | 0.10 | 293044.94 | thorax |
| 12 | Bg2/+;Mef2/3.3+1077 | 15d | F | none | 29-20 | 24760.00 | 0.11 | 229059.34 | thorax |
| 12 | Bg2/+;Mef2/3.3+1077 | 15d | F | none | 29-20 | 21178.00 | 0.10 | 218054.27 | thorax |
| 13 | Bg2/+;Mef2/3.3+1077 | 15d | F | none | 29-20 | 22950.00 | 0.07 | 317531.48 | thorax |
| 13 | Bg2/+;Mef2/3.3+1077 | 15d | F | none | 29-20 | 19860.00 | 0.09 | 230639.88 | thorax |
| 13 | Bg2/+;Mef2/3.3+1077 | 15d | F | none | 29-20 | 21618.00 | 0.10 | 226771.17 | thorax |
| 13 | Bg2/+;Mef2/3.3+1077 | 15d | F | none | 29-20 | 17748.00 | 0.08 | 229811.58 | thorax |
| 13 | Bg2/+;Mef2/3.3+1077 | 15d | F | none | 29-20 | 19034.00 | 0.09 | 220174.03 | thorax |
| 14 | Bg2/+;Mef2/3.3+1077 | 15d | F | none | 29-20 | 27070.00 | 0.17 | 158560.47 | thorax |
| 14 | Bg2/+;Mef2/3.3+1077 | 15d | F | none | 29-20 | 28640.00 | 0.13 | 226199.72 | thorax |

| Rep. | Genotype | Stage | Sex | Treatmen | °C | ß-gal ΔmOD/min/ ml | Protein mg/ml | Specific Activity ΔmOD/min/ mg | Whole/ dissected |
|---|---|---|---|---|---|---|---|---|---|
| 14 | Bg2/+;Mef2/3.3+1077 | 15d | F | none | 29-20 | 28514.00 | 0.13 | 217132.72 | thorax |
| 14 | Bg2/+;Mef2/3.3+1077 | 15d | F | none | 29-20 | 27802.00 | 0.15 | 181426.65 | thorax |
| 14 | Bg2/+;Mef2/3.3+1077 | 15d | F | none | 29-20 | 26754.00 | 0.13 | 213114.88 | thorax |
| 15 | Bg2/+;Mef2/3.3+1077 | 15d | F | none | 29-20 | 28428.00 | 0.13 | 214392.10 | thorax |
| 15 | Bg2/+;Mef2/3.3+1077 | 15d | F | none | 29-20 | 29586.00 | 0.11 | 264708.03 | thorax |
| 15 | Bg2/+;Mef2/3.3+1077 | 15d | F | none | 29-20 | 26976.00 | 0.11 | 250044.60 | thorax |
| 15 | Bg2/+;Mef2/3.3+1077 | 15d | F | none | 29-20 | 29136.00 | 0.15 | 197138.58 | thorax |
| 15 | Bg2/+;Mef2/3.3+1077 | 15d | F | none | 29-20 | 31626.00 | 0.16 | 197872.05 | thorax |
| 12 | Bg2/+;Mef2/3.3+1077 | 10d | M | none | 20 | 17190.00 | 0.09 | 181063.67 | thorax |
| 12 | Bg2/+;Mef2/3.3+1077 | 10d | M | none | 20 | 16786.00 | 0.10 | 170600.26 | thorax |
| 12 | Bg2/+;Mef2/3.3+1077 | 10d | M | none | 20 | 14492.00 | 0.11 | 134807.05 | thorax |
| 12 | Bg2/+;Mef2/3.3+1077 | 10d | M | none | 20 | 13906.00 | 0.11 | 129302.01 | thorax |
| 12 | Bg2/+;Mef2/3.3+1077 | 10d | M | none | 20 | 16558.00 | 0.09 | 177767.20 | thorax |
| 13 | Bg2/+;Mef2/3.3+1077 | 10d | M | none | 20 | 18082.00 | 0.08 | 216672.91 | thorax |
| 13 | Bg2/+;Mef2/3.3+1077 | 10d | M | none | 20 | 16336.00 | 0.09 | 181775.72 | thorax |
| 13 | Bg2/+;Mef2/3.3+1077 | 10d | M | none | 20 | 13314.00 | 0.09 | 142322.51 | thorax |
| 13 | Bg2/+;Mef2/3.3+1077 | 10d | M | none | 20 | 15898.00 | 0.08 | 192051.29 | thorax |
| 13 | Bg2/+;Mef2/3.3+1077 | 10d | M | none | 20 | 13012.00 | 0.11 | 121140.96 | thorax |
| 14 | Bg2/+;Mef2/3.3+1077 | 10d | M | none | 20 | 18246.00 | 0.12 | 148907.63 | thorax |
| 14 | Bg2/+;Mef2/3.3+1077 | 10d | M | none | 20 | 20406.00 | 0.11 | 191925.82 | thorax |
| 14 | Bg2/+;Mef2/3.3+1077 | 10d | M | none | 20 | 15026.00 | 0.12 | 127314.75 | thorax |
| 14 | Bg2/+;Mef2/3.3+1077 | 10d | M | none | 20 | 16662.00 | 0.10 | 167575.37 | thorax |
| 14 | Bg2/+;Mef2/3.3+1077 | 10d | M | none | 20 | 15636.00 | 0.11 | 144290.75 | thorax |
| 15 | Bg2/+;Mef2/3.3+1077 | 10d | M | none | 20 | 19882.00 | 0.12 | 170924.11 | thorax |
| 15 | Bg2/+;Mef2/3.3+1077 | 10d | M | none | 20 | 14738.00 | 0.08 | 177006.62 | thorax |
| 15 | Bg2/+;Mef2/3.3+1077 | 10d | M | none | 20 | 14416.00 | 0.08 | 174836.77 | thorax |
| 15 | Bg2/+;Mef2/3.3+1077 | 10d | M | none | 20 | 16400.00 | 0.08 | 199930.29 | thorax |
| 15 | Bg2/+;Mef2/3.3+1077 | 10d | M | none | 20 | 19504.00 | 0.09 | 205662.64 | thorax |
| 8 | Bg2/+;Mef2/3.3,DJ1077 | 0-4d | M | none | 25 | 32363.33 | 0.12 | 264600.75 | thorax |
| 8 | Bg2/+;Mef2/3.3,DJ1077 | 0-4d | M | none | 25 | 29120.00 | 0.12 | 245806.27 | thorax |
| 8 | Bg2/+;Mef2/3.3,DJ1077 | 0-4d | M | none | 25 | 33865.00 | 0.11 | 322301.38 | thorax |
| 8 | Bg2/+;Mef2/3.3,DJ1077 | 0-4d | M | none | 25 | 40700.00 | 0.10 | 403807.84 | thorax |
| 8 | Bg2/+;Mef2/3.3,DJ1077 | 0-4d | M | none | 25 | 34376.00 | 0.12 | 292067.73 | thorax |
| 9 | Bg2/+;Mef2/3.3,DJ1077 | 1-3d | M | none | 25 | 37050.00 | 0.12 | 312236.15 | thorax |
| 9 | Bg2/+;Mef2/3.3,DJ1077 | 1-3d | M | none | 25 | 29980.00 | 0.11 | 284014.73 | thorax |
| 9 | Bg2/+;Mef2/3.3,DJ1077 | 1-3d | M | none | 25 | 42620.00 | 0.08 | 517122.67 | thorax |
| 9 | Bg2/+;Mef2/3.3,DJ1077 | 1-3d | M | none | 25 | 29340.00 | 0.08 | 352377.87 | thorax |
| 9 | Bg2/+;Mef2/3.3,DJ1077 | 1-3d | M | none | 25 | 30300.00 | 0.12 | 255351.02 | thorax |
| 12 | Bg2/+;Mef2/3.3+1077 | 15d | M | none | 29 | 14784.00 | 0.09 | 163581.67 | thorax |
| 12 | Bg2/+;Mef2/3.3+1077 | 15d | M | none | 29 | 18652.00 | 0.08 | 244752.77 | thorax |

| Rep. | Genotype | Stage | Sex | Treatmen | °C | ß-gal ΔmOD/min/ ml | Protein mg/ml | Specific Activity ΔmOD/min/ mg | Whole/ dissected |
|---|---|---|---|---|---|---|---|---|---|
| 12 | Bg2/+;Mef2/3.3+1077 | 15d | M | none | 29 | 18536.00 | 0.10 | 188393.16 | thorax |
| 12 | Bg2/+;Mef2/3.3+1077 | 15d | M | none | 29 | 17284.00 | 0.08 | 204373.47 | thorax |
| 13 | Bg2/+;Mef2/3.3+1077 | 15d | M | none | 29 | 14934.00 | 0.10 | 152198.83 | thorax |
| 13 | Bg2/+;Mef2/3.3+1077 | 15d | M | none | 29 | 16218.00 | 0.10 | 159962.28 | thorax |
| 13 | Bg2/+;Mef2/3.3+1077 | 15d | M | none | 29 | 16714.00 | 0.10 | 174637.87 | thorax |
| 13 | Bg2/+;Mef2/3.3+1077 | 15d | M | none | 29 | 16362.00 | 0.08 | 206931.18 | thorax |
| 13 | Bg2/+;Mef2/3.3+1077 | 15d | M | none | 29 | 16174.00 | 0.10 | 155081.75 | thorax |
| 14 | Bg2/+;Mef2/3.3+1077 | 15d | M | none | 29 | 16450.00 | 0.08 | 210184.00 | thorax |
| 14 | Bg2/+;Mef2/3.3+1077 | 15d | M | none | 29 | 15832.00 | 0.10 | 158674.82 | thorax |
| 14 | Bg2/+;Mef2/3.3+1077 | 15d | M | none | 29 | 10154.00 | 0.09 | 110100.54 | thorax |
| 14 | Bg2/+;Mef2/3.3+1077 | 15d | M | none | 29 | 6638.00 | 0.07 | 92413.47 | thorax |
| 14 | Bg2/+;Mef2/3.3+1077 | 15d | M | none | 29 | 9638.00 | 0.08 | 114031.05 | thorax |
| 15 | Bg2/+;Mef2/3.3+1077 | 15d | M | none | 29 | 8584.00 | 0.08 | 110592.26 | thorax |
| 15 | Bg2/+;Mef2/3.3+1077 | 15d | M | none | 29 | 8470.00 | 0.04 | 192123.56 | thorax |
| 15 | Bg2/+;Mef2/3.3+1077 | 15d | M | none | 29 | 10184.00 | 0.08 | 129570.48 | thorax |
| 15 | Bg2/+;Mef2/3.3+1077 | 15d | M | none | 29 | 8756.00 | 0.09 | 102622.51 | thorax |
| 15 | Bg2/+;Mef2/3.3+1077 | 15d | M | none | 29 | 6986.00 | 0.08 | 87886.62 | thorax |
| 12 | Bg2/+;Mef2/3.3+1077 | 15d | M | none | 20-29 | 17862.00 | 0.14 | 131203.33 | thorax |
| 12 | Bg2/+;Mef2/3.3+1077 | 15d | M | none | 20-29 | 16060.00 | 0.12 | 139166.33 | thorax |
| 12 | Bg2/+;Mef2/3.3+1077 | 15d | M | none | 20-29 | 18810.00 | 0.10 | 191764.50 | thorax |
| 12 | Bg2/+;Mef2/3.3+1077 | 15d | M | none | 20-29 | 14344.00 | 0.10 | 137567.34 | thorax |
| 12 | Bg2/+;Mef2/3.3+1077 | 15d | M | none | 20-29 | 16730.00 | 0.12 | 144259.89 | thorax |
| 13 | Bg2/+;Mef2/3.3+1077 | 15d | M | none | 20-29 | 12950.00 | 0.10 | 124250.29 | thorax |
| 13 | Bg2/+;Mef2/3.3+1077 | 15d | M | none | 20-29 | 12218.00 | 0.11 | 111729.81 | thorax |
| 13 | Bg2/+;Mef2/3.3+1077 | 15d | M | none | 20-29 | 15646.00 | 0.12 | 131775.24 | thorax |
| 13 | Bg2/+;Mef2/3.3+1077 | 15d | M | none | 20-29 | 17226.00 | 0.10 | 165207.40 | thorax |
| 13 | Bg2/+;Mef2/3.3+1077 | 15d | M | none | 20-29 | 12626.00 | 0.09 | 139233.84 | thorax |
| 12 | Bg2/+;Mef2/3.3+1077 | 15d | M | none | 29-20 | 14182.00 | 0.08 | 171753.07 | thorax |
| 12 | Bg2/+;Mef2/3.3+1077 | 15d | M | none | 29-20 | 18552.00 | 0.08 | 218855.93 | thorax |
| 12 | Bg2/+;Mef2/3.3+1077 | 15d | M | none | 29-20 | 20362.00 | 0.08 | 253196.08 | thorax |
| 12 | Bg2/+;Mef2/3.3+1077 | 15d | M | none | 29-20 | 15720.00 | 0.09 | 176572.75 | thorax |
| 12 | Bg2/+;Mef2/3.3+1077 | 15d | M | none | 29-20 | 13848.00 | 0.08 | 175454.24 | thorax |
| 13 | Bg2/+;Mef2/3.3+1077 | 15d | M | none | 29-20 | 10482.00 | 0.07 | 154970.24 | thorax |
| 13 | Bg2/+;Mef2/3.3+1077 | 15d | M | none | 29-20 | 13492.00 | 0.06 | 226705.43 | thorax |
| 13 | Bg2/+;Mef2/3.3+1077 | 15d | M | none | 29-20 | 20354.00 | 0.07 | 307715.72 | thorax |
| 13 | Bg2/+;Mef2/3.3+1077 | 15d | M | none | 29-20 | 12804.00 | 0.07 | 172294.01 | thorax |
| 13 | Bg2/+;Mef2/3.3+1077 | 15d | M | none | 29-20 | 17512.00 | 0.07 | 259410.03 | thorax |
| 14 | Bg2/+;Mef2/3.3+1077 | 15d | M | none | 29-20 | 23696.00 | 0.12 | 200302.16 | thorax |
| 14 | Bg2/+;Mef2/3.3+1077 | 15d | M | none | 29-20 | 19792.00 | 0.12 | 170331.15 | thorax |
| 14 | Bg2/+;Mef2/3.3+1077 | 15d | M | none | 29-20 | 24362.00 | 0.12 | 203567.37 | thorax |

| Rep. | Genotype | Stage | Sex | Treatmen | °C | ß-gal ΔmOD/min/ ml | Protein mg/ml | Specific Activity ΔmOD/min/ mg | Whole/ dissected |
|---|---|---|---|---|---|---|---|---|---|
| 14 | Bg2/+;Mef2/3.3+1077 | 15d | M | none | 29-20 | 20660.00 | 0.12 | 171403.66 | thorax |
| 14 | Bg2/+;Mef2/3.3+1077 | 15d | M | none | 29-20 | 19126.00 | 0.09 | 202918.58 | thorax |
| 15 | Bg2/+;Mef2/3.3+1077 | 15d | M | none | 29-20 | 21378.00 | 0.09 | 243091.24 | thorax |
| 15 | Bg2/+;Mef2/3.3+1077 | 15d | M | none | 29-20 | 20890.00 | 0.11 | 193357.04 | thorax |
| 15 | Bg2/+;Mef2/3.3+1077 | 15d | M | none | 29-20 | 23008.00 | 0.11 | 205448.74 | thorax |
| 15 | Bg2/+;Mef2/3.3+1077 | 15d | M | none | 29-20 | 19842.00 | 0.09 | 216837.40 | thorax |
| 15 | Bg2/+;Mef2/3.3+1077 | 15d | M | none | 29-20 | 23434.00 | 0.11 | 220943.72 | thorax |
| 12 | Bg2/+;Mef2/3.3+1077 | LP | ND | none | 20 | 22820.00 | 0.32 | 72058.93 | whole |
| 12 | Bg2/+;Mef2/3.3+1077 | LP | ND | none | 20 | 16748.00 | 0.20 | 82850.80 | whole |
| 12 | Bg2/+;Mef2/3.3+1077 | LP | ND | none | 20 | 23570.00 | 0.40 | 58318.87 | whole |
| 12 | Bg2/+;Mef2/3.3+1077 | LP | ND | none | 20 | 14536.00 | 0.54 | 26830.25 | whole |
| 12 | Bg2/+;Mef2/3.3+1077 | LP | ND | none | 20 | 15028.00 | 0.67 | 22375.68 | whole |
| 13 | Bg2/+;Mef2/3.3+1077 | LP | ND | none | 20 | 20240.00 | 0.50 | 40222.65 | whole |
| 13 | Bg2/+;Mef2/3.3+1077 | LP | ND | none | 20 | 10530.00 | 0.34 | 30934.03 | whole |
| 13 | Bg2/+;Mef2/3.3+1077 | LP | ND | none | 20 | 11636.00 | 0.20 | 59594.16 | whole |
| 13 | Bg2/+;Mef2/3.3+1077 | LP | ND | none | 20 | 20856.00 | 0.24 | 88624.91 | whole |
| 13 | Bg2/+;Mef2/3.3+1077 | LP | ND | none | 20 | 10122.00 | 0.18 | 56576.24 | whole |
| 14 | Bg2/+;Mef2/3.3+1077 | LP | ND | none | 20 | 5244.00 | 0.19 | 27210.45 | whole |
| 14 | Bg2/+;Mef2/3.3+1077 | LP | ND | none | 20 | 11888.00 | 0.33 | 36120.60 | whole |
| 14 | Bg2/+;Mef2/3.3+1077 | LP | ND | none | 20 | 13148.00 | 0.44 | 29939.00 | whole |
| 14 | Bg2/+;Mef2/3.3+1077 | LP | ND | none | 20 | 15188.00 | 0.36 | 42745.65 | whole |
| 14 | Bg2/+;Mef2/3.3+1077 | LP | ND | none | 20 | 8252.00 | 0.26 | 31894.20 | whole |
| 15 | Bg2/+;Mef2/3.3+1077 | LP | ND | none | 20 | 10428.00 | 0.28 | 37781.31 | whole |
| 15 | Bg2/+;Mef2/3.3+1077 | LP | ND | none | 20 | 8284.00 | 0.25 | 33276.84 | whole |
| 15 | Bg2/+;Mef2/3.3+1077 | LP | ND | none | 20 | 13600.00 | 0.29 | 46442.73 | whole |
| 15 | Bg2/+;Mef2/3.3+1077 | LP | ND | none | 20 | 8418.00 | 0.30 | 28212.23 | whole |
| 15 | Bg2/+;Mef2/3.3+1077 | LP | ND | none | 20 | 20164.00 | 0.47 | 42919.73 | whole |
| 8 | Bg2/+;Mef2/3.3,DJ1077 | EP | ND | none | 25 | 5206.67 | 0.34 | 15148.89 | whole |
| 8 | Bg2/+;Mef2/3.3,DJ1077 | EP | ND | none | 25 | 4900.00 | 0.30 | 16256.01 | whole |
| 8 | Bg2/+;Mef2/3.3,DJ1077 | EP | ND | none | 25 | 4565.00 | 0.24 | 18929.43 | whole |
| 8 | Bg2/+;Mef2/3.3,DJ1077 | EP | ND | none | 25 | 3540.00 | 0.39 | 9194.10 | whole |
| 8 | Bg2/+;Mef2/3.3,DJ1077 | EP | ND | none | 25 | 6680.00 | 0.44 | 15243.94 | whole |
| 9 | Bg2/+;Mef2/3.3,DJ1077 | EP | ND | none | 25 | 6833.33 | 0.45 | 15246.94 | whole |
| 9 | Bg2/+;Mef2/3.3,DJ1077 | EP | ND | none | 25 | 7460.00 | 0.43 | 17394.84 | whole |
| 9 | Bg2/+;Mef2/3.3,DJ1077 | EP | ND | none | 25 | 9515.00 | 0.47 | 20371.08 | whole |
| 9 | Bg2/+;Mef2/3.3,DJ1077 | EP | ND | none | 25 | 3760.00 | 0.34 | 11155.58 | whole |
| 9 | Bg2/+;Mef2/3.3,DJ1077 | EP | ND | none | 25 | 4375.56 | 0.39 | 11355.13 | whole |
| 9 | Bg2/+;Mef2/3.3,DJ1077 | EP | ND | none | 25 | 4523.33 | 0.25 | 18429.53 | whole |
| 8 | Bg2/+;Mef2/3.3,DJ1077 | LP | ND | none | 25 | 57670.00 | 0.36 | 158239.84 | whole |
| 8 | Bg2/+;Mef2/3.3,DJ1077 | LP | ND | none | 25 | 36535.56 | 0.27 | 133922.74 | whole |

| Rep. | Genotype | Stage | Sex | Treatmen | °C | ß-gal ΔmOD/min/ ml | Protein mg/ml | Specific Activity ΔmOD/min/ mg | Whole/ dissected |
|---|---|---|---|---|---|---|---|---|---|
| 8 | Bg2/+;Mef2/3.3,DJ1077 | LP | ND | none | 25 | 58400.00 | 0.30 | 196071.83 | whole |
| 9 | Bg2/+;Mef2/3.3,DJ1077 | LP | ND | none | 25 | 47430.00 | 0.43 | 110798.02 | whole |
| 9 | Bg2/+;Mef2/3.3,DJ1077 | LP | ND | none | 25 | 62290.00 | 0.57 | 108469.91 | whole |
| 9 | Bg2/+;Mef2/3.3,DJ1077 | LP | ND | none | 25 | 36613.33 | 0.43 | 84194.86 | whole |
| 12 | Bg2/+;Mef2/3.3+1077 | LP | ND | none | 29 | 51890.00 | 0.40 | 129282.38 | whole |
| 12 | Bg2/+;Mef2/3.3+1077 | LP | ND | none | 29 | 41220.00 | 0.26 | 159940.12 | whole |
| 12 | Bg2/+;Mef2/3.3+1077 | LP | ND | none | 29 | 17420.00 | 0.15 | 117899.82 | whole |
| 12 | Bg2/+;Mef2/3.3+1077 | LP | ND | none | 29 | 46070.00 | 0.44 | 104192.58 | whole |
| 13 | Bg2/+;Mef2/3.3+1077 | LP | ND | none | 29 | 20990.00 | 0.16 | 129395.14 | whole |
| 13 | Bg2/+;Mef2/3.3+1077 | LP | ND | none | 29 | 36235.00 | 0.34 | 107437.17 | whole |
| 13 | Bg2/+;Mef2/3.3+1077 | LP | ND | none | 29 | 45895.00 | 0.35 | 131974.56 | whole |
| 13 | Bg2/+;Mef2/3.3+1077 | LP | ND | none | 29 | 17225.00 | 0.18 | 95038.86 | whole |
| 13 | Bg2/+;Mef2/3.3+1077 | LP | ND | none | 29 | 32525.00 | 0.23 | 139873.06 | whole |
| 14 | Bg2/+;Mef2/3.3+1077 | LP | ND | none | 29 | 16550.00 | 0.17 | 95537.19 | whole |
| 14 | Bg2/+;Mef2/3.3+1077 | LP | ND | none | 29 | 23695.00 | 0.11 | 207830.84 | whole |
| 14 | Bg2/+;Mef2/3.3+1077 | LP | ND | none | 29 | 20520.00 | 0.18 | 113039.16 | whole |
| 14 | Bg2/+;Mef2/3.3+1077 | LP | ND | none | 29 | 35035.00 | 0.20 | 173688.44 | whole |
| 14 | Bg2/+;Mef2/3.3+1077 | LP | ND | none | 29 | 34220.00 | 0.19 | 178335.95 | whole |
| 15 | Bg2/+;Mef2/3.3+1077 | LP | ND | none | 29 | 24370.00 | 0.24 | 103055.26 | whole |
| 15 | Bg2/+;Mef2/3.3+1077 | LP | ND | none | 29 | 14700.00 | 0.24 | 61044.67 | whole |
| 15 | Bg2/+;Mef2/3.3+1077 | LP | ND | none | 29 | 27565.00 | 0.25 | 108909.88 | whole |
| 15 | Bg2/+;Mef2/3.3+1077 | LP | ND | none | 29 | 25550.00 | 0.17 | 150282.91 | whole |
| 15 | Bg2/+;Mef2/3.3+1077 | LP | ND | none | 29 | 14810.00 | 0.18 | 83979.53 | whole |
| 12 | Bg2/+;Mef2/GAL80ts | 10d | F | none | 20 | 2452.00 | 0.13 | 18471.64 | thorax |
| 12 | Bg2/+;Mef2/GAL80ts | 10d | F | none | 20 | 3064.00 | 0.11 | 29123.37 | thorax |
| 12 | Bg2/+;Mef2/GAL80ts | 10d | F | none | 20 | 2290.00 | 0.14 | 16124.24 | thorax |
| 12 | Bg2/+;Mef2/GAL80ts | 10d | F | none | 20 | 2952.00 | 0.10 | 28523.87 | thorax |
| 13 | Bg2/+;Mef2/GAL80ts | 10d | F | none | 20 | 2706.00 | 0.11 | 23967.13 | thorax |
| 13 | Bg2/+;Mef2/GAL80ts | 10d | F | none | 20 | 2028.00 | 0.11 | 17830.09 | thorax |
| 13 | Bg2/+;Mef2/GAL80ts | 10d | F | none | 20 | 3780.00 | 0.11 | 35251.06 | thorax |
| 13 | Bg2/+;Mef2/GAL80ts | 10d | F | none | 20 | 2575.00 | 0.11 | 22586.88 | thorax |
| 13 | Bg2/+;Mef2/GAL80ts | 10d | F | none | 20 | 2468.00 | 0.12 | 20374.89 | thorax |
| 14 | Bg2/+;Mef2/GAL80ts | 10d | F | none | 20 | 2742.00 | 0.14 | 19816.52 | thorax |
| 14 | Bg2/+;Mef2/GAL80ts | 10d | F | none | 20 | 2852.00 | 0.14 | 20448.83 | thorax |
| 14 | Bg2/+;Mef2/GAL80ts | 10d | F | none | 20 | 3336.00 | 0.07 | 50965.93 | thorax |
| 14 | Bg2/+;Mef2/GAL80ts | 10d | F | none | 20 | 3124.00 | 0.10 | 30011.27 | thorax |
| 14 | Bg2/+;Mef2/GAL80ts | 10d | F | none | 20 | 3156.00 | 0.11 | 28565.40 | thorax |
| 15 | Bg2/+;Mef2/GAL80ts | 10d | F | none | 20 | 3478.00 | 0.09 | 38639.22 | thorax |
| 15 | Bg2/+;Mef2/GAL80ts | 10d | F | none | 20 | 3160.00 | 0.15 | 21746.24 | thorax |
| 15 | Bg2/+;Mef2/GAL80ts | 10d | F | none | 20 | 5004.00 | 0.07 | 69819.30 | thorax |

| Rep. | Genotype | Stage | Sex | Treatmen | °C | ß-gal ΔmOD/min/ ml | Protein mg/ml | Specific Activity ΔmOD/min/ mg | Whole/ dissected |
|---|---|---|---|---|---|---|---|---|---|
| 15 | Bg2/+;Mef2/GAL80ts | 10d | F | none | 20 | 1692.50 | 0.09 | 19249.15 | thorax |
| 15 | Bg2/+;Mef2/GAL80ts | 10d | F | none | 20 | 2282.00 | 0.08 | 28107.20 | thorax |
| 12 | Bg2/+;Mef2/GAL80ts | 15d | F | none | 29 | 11692.00 | 0.09 | 135542.89 | thorax |
| 12 | Bg2/+;Mef2/GAL80ts | 15d | F | none | 29 | 11466.00 | 0.11 | 107409.01 | thorax |
| 12 | Bg2/+;Mef2/GAL80ts | 15d | F | none | 29 | 14344.00 | 0.09 | 151213.70 | thorax |
| 12 | Bg2/+;Mef2/GAL80ts | 15d | F | none | 29 | 13124.00 | 0.11 | 121277.74 | thorax |
| 12 | Bg2/+;Mef2/GAL80ts | 15d | F | none | 29 | 13320.00 | 0.11 | 117147.42 | thorax |
| 13 | Bg2/+;Mef2/GAL80ts | 15d | F | none | 29 | 19344.00 | 0.12 | 164887.80 | thorax |
| 13 | Bg2/+;Mef2/GAL80ts | 15d | F | none | 29 | 16190.00 | 0.12 | 138218.73 | thorax |
| 13 | Bg2/+;Mef2/GAL80ts | 15d | F | none | 29 | 14916.00 | 0.12 | 128851.61 | thorax |
| 13 | Bg2/+;Mef2/GAL80ts | 15d | F | none | 29 | 14448.00 | 0.12 | 116737.28 | thorax |
| 13 | Bg2/+;Mef2/GAL80ts | 15d | F | none | 29 | 13246.00 | 0.12 | 109992.61 | thorax |
| 14 | Bg2/+;Mef2/GAL80ts | 15d | F | none | 29 | 15724.00 | 0.08 | 189729.32 | thorax |
| 14 | Bg2/+;Mef2/GAL80ts | 15d | F | none | 29 | 12718.00 | 0.12 | 104222.77 | thorax |
| 14 | Bg2/+;Mef2/GAL80ts | 15d | F | none | 29 | 12898.00 | 0.11 | 120205.40 | thorax |
| 14 | Bg2/+;Mef2/GAL80ts | 15d | F | none | 29 | 14692.00 | 0.13 | 109144.41 | thorax |
| 14 | Bg2/+;Mef2/GAL80ts | 15d | F | none | 29 | 18654.00 | 0.13 | 144255.23 | thorax |
| 15 | Bg2/+;Mef2/GAL80ts | 15d | F | none | 29 | 14640.00 | 0.09 | 154707.09 | thorax |
| 15 | Bg2/+;Mef2/GAL80ts | 15d | F | none | 29 | 15760.00 | 0.13 | 121759.31 | thorax |
| 15 | Bg2/+;Mef2/GAL80ts | 15d | F | none | 29 | 11294.00 | 0.07 | 167866.77 | thorax |
| 15 | Bg2/+;Mef2/GAL80ts | 15d | F | none | 29 | 15096.00 | 0.10 | 153515.79 | thorax |
| 15 | Bg2/+;Mef2/GAL80ts | 15d | F | none | 29 | 16634.00 | 0.08 | 202723.40 | thorax |
| 12 | Bg2/+;Mef2/GAL80ts | 15d | F | none | 20-29 | 6746.00 | 0.13 | 52681.00 | thorax |
| 12 | Bg2/+;Mef2/GAL80ts | 15d | F | none | 20-29 | 8246.00 | 0.14 | 58391.67 | thorax |
| 12 | Bg2/+;Mef2/GAL80ts | 15d | F | none | 20-29 | 12926.00 | 0.13 | 101425.74 | thorax |
| 12 | Bg2/+;Mef2/GAL80ts | 15d | F | none | 20-29 | 7972.00 | 0.11 | 74092.33 | thorax |
| 12 | Bg2/+;Mef2/GAL80ts | 15d | F | none | 20-29 | 13038.00 | 0.12 | 106410.03 | thorax |
| 13 | Bg2/+;Mef2/GAL80ts | 15d | F | none | 20-29 | 8192.00 | 0.13 | 60722.99 | thorax |
| 13 | Bg2/+;Mef2/GAL80ts | 15d | F | none | 20-29 | 6858.00 | 0.15 | 44716.59 | thorax |
| 13 | Bg2/+;Mef2/GAL80ts | 15d | F | none | 20-29 | 8040.00 | 0.11 | 74007.83 | thorax |
| 13 | Bg2/+;Mef2/GAL80ts | 15d | F | none | 20-29 | 7792.00 | 0.18 | 42542.76 | thorax |
| 13 | Bg2/+;Mef2/GAL80ts | 15d | F | none | 20-29 | 8338.00 | 0.10 | 79778.87 | thorax |
| 14 | Bg2/+;Mef2/GAL80ts | 15d | F | none | 20-29 | 5440.00 | 0.13 | 43217.62 | thorax |
| 14 | Bg2/+;Mef2/GAL80ts | 15d | F | none | 20-29 | 5336.00 | 0.14 | 39303.42 | thorax |
| 14 | Bg2/+;Mef2/GAL80ts | 15d | F | none | 20-29 | 9978.00 | 0.17 | 60363.45 | thorax |
| 14 | Bg2/+;Mef2/GAL80ts | 15d | F | none | 20-29 | 9556.00 | 0.12 | 81748.04 | thorax |
| 14 | Bg2/+;Mef2/GAL80ts | 15d | F | none | 20-29 | 7414.00 | 0.12 | 60276.62 | thorax |
| 15 | Bg2/+;Mef2/GAL80ts | 15d | F | none | 20-29 | 8530.00 | 0.11 | 75181.23 | thorax |
| 15 | Bg2/+;Mef2/GAL80ts | 15d | F | none | 20-29 | 6934.00 | 0.11 | 62297.36 | thorax |
| 15 | Bg2/+;Mef2/GAL80ts | 15d | F | none | 20-29 | 6802.00 | 0.15 | 44054.43 | thorax |

| Rep. | Genotype | Stage | Sex | Treatmen | °C | ß-gal ΔmOD/min/ ml | Protein mg/ml | Specific Activity ΔmOD/min/ mg | Whole/ dissected |
|---|---|---|---|---|---|---|---|---|---|
| 15 | Bg2/+;Mef2/GAL80ts | 15d | F | none | 20-29 | 8472.00 | 0.12 | 73051.80 | thorax |
| 15 | Bg2/+;Mef2/GAL80ts | 15d | F | none | 20-29 | 5174.00 | 0.13 | 38414.75 | thorax |
| 12 | Bg2/+;Mef2/GAL80ts | 15d | F | none | 29-20 | 9280.00 | 0.11 | 88148.71 | thorax |
| 12 | Bg2/+;Mef2/GAL80ts | 15d | F | none | 29-20 | 10358.00 | 0.11 | 92711.42 | thorax |
| 12 | Bg2/+;Mef2/GAL80ts | 15d | F | none | 29-20 | 13322.00 | 0.10 | 130512.14 | thorax |
| 12 | Bg2/+;Mef2/GAL80ts | 15d | F | none | 29-20 | 6724.00 | 0.10 | 67918.49 | thorax |
| 12 | Bg2/+;Mef2/GAL80ts | 15d | F | none | 29-20 | 11146.00 | 0.09 | 121040.29 | thorax |
| 13 | Bg2/+;Mef2/GAL80ts | 15d | F | none | 29-20 | 8346.00 | 0.09 | 88620.45 | thorax |
| 13 | Bg2/+;Mef2/GAL80ts | 15d | F | none | 29-20 | 11664.00 | 0.08 | 139183.67 | thorax |
| 13 | Bg2/+;Mef2/GAL80ts | 15d | F | none | 29-20 | 7512.00 | 0.10 | 73164.69 | thorax |
| 13 | Bg2/+;Mef2/GAL80ts | 15d | F | none | 29-20 | 10426.00 | 0.00 | 2978275.90 | thorax |
| 13 | Bg2/+;Mef2/GAL80ts | 15d | F | none | 29-20 | 8204.00 | 0.21 | 39260.68 | thorax |
| 14 | Bg2/+;Mef2/GAL80ts | 15d | F | none | 29-20 | 14458.00 | 0.16 | 91927.91 | thorax |
| 14 | Bg2/+;Mef2/GAL80ts | 15d | F | none | 29-20 | 14710.00 | 0.11 | 134825.43 | thorax |
| 14 | Bg2/+;Mef2/GAL80ts | 15d | F | none | 29-20 | 15526.00 | 0.12 | 132744.30 | thorax |
| 14 | Bg2/+;Mef2/GAL80ts | 15d | F | none | 29-20 | 14308.00 | 0.09 | 152325.17 | thorax |
| 14 | Bg2/+;Mef2/GAL80ts | 15d | F | none | 29-20 | 14690.00 | 0.13 | 113490.74 | thorax |
| 15 | Bg2/+;Mef2/GAL80ts | 15d | F | none | 29-20 | 12728.00 | 0.12 | 106117.78 | thorax |
| 15 | Bg2/+;Mef2/GAL80ts | 15d | F | none | 29-20 | 17368.00 | 0.11 | 162482.89 | thorax |
| 15 | Bg2/+;Mef2/GAL80ts | 15d | F | none | 29-20 | 12948.00 | 0.13 | 100450.00 | thorax |
| 15 | Bg2/+;Mef2/GAL80ts | 15d | F | none | 29-20 | 18606.00 | 0.12 | 155534.64 | thorax |
| 15 | Bg2/+;Mef2/GAL80ts | 15d | F | none | 29-20 | 18900.00 | 0.11 | 170616.23 | thorax |
| 12 | Bg2/+;Mef2/GAL80ts | 10d | M | none | 20 | 2918.00 | 0.10 | 30670.33 | thorax |
| 12 | Bg2/+;Mef2/GAL80ts | 10d | M | none | 20 | 1816.00 | 0.11 | 16963.54 | thorax |
| 12 | Bg2/+;Mef2/GAL80ts | 10d | M | none | 20 | 1570.00 | 0.10 | 16412.83 | thorax |
| 12 | Bg2/+;Mef2/GAL80ts | 10d | M | none | 20 | 1942.00 | 0.09 | 21144.75 | thorax |
| 13 | Bg2/+;Mef2/GAL80ts | 10d | M | none | 20 | 2404.00 | 0.08 | 30443.38 | thorax |
| 13 | Bg2/+;Mef2/GAL80ts | 10d | M | none | 20 | 1390.00 | 0.09 | 15622.96 | thorax |
| 13 | Bg2/+;Mef2/GAL80ts | 10d | M | none | 20 | 2072.00 | 0.11 | 18988.79 | thorax |
| 13 | Bg2/+;Mef2/GAL80ts | 10d | M | none | 20 | 3138.00 | 0.09 | 33528.16 | thorax |
| 13 | Bg2/+;Mef2/GAL80ts | 10d | M | none | 20 | 2097.50 | 0.09 | 24184.73 | thorax |
| 14 | Bg2/+;Mef2/GAL80ts | 10d | M | none | 20 | 2336.00 | 0.09 | 25311.02 | thorax |
| 14 | Bg2/+;Mef2/GAL80ts | 10d | M | none | 20 | 2754.00 | 0.09 | 30015.23 | thorax |
| 14 | Bg2/+;Mef2/GAL80ts | 10d | M | none | 20 | 2584.00 | 0.08 | 31018.56 | thorax |
| 14 | Bg2/+;Mef2/GAL80ts | 10d | M | none | 20 | 2684.00 | 0.08 | 35560.73 | thorax |
| 14 | Bg2/+;Mef2/GAL80ts | 10d | M | none | 20 | 1954.00 | 0.09 | 21391.16 | thorax |
| 15 | Bg2/+;Mef2/GAL80ts | 10d | M | none | 20 | 2090.00 | 0.08 | 25256.23 | thorax |
| 15 | Bg2/+;Mef2/GAL80ts | 10d | M | none | 20 | 3202.00 | 0.09 | 33885.55 | thorax |
| 15 | Bg2/+;Mef2/GAL80ts | 10d | M | none | 20 | 1520.00 | 0.08 | 19792.84 | thorax |
| 15 | Bg2/+;Mef2/GAL80ts | 10d | M | none | 20 | 1514.00 | 0.07 | 21979.65 | thorax |

| Rep. | Genotype | Stage | Sex | Treatmen | °C | ß-gal ΔmOD/min/ ml | Protein mg/ml | Specific Activity ΔmOD/min/ mg | Whole/ dissected |
|---|---|---|---|---|---|---|---|---|---|
| 15 | Bg2/+;Mef2/GAL80ts | 10d | M | none | 20 | 1315.00 | 0.07 | 18743.34 | thorax |
| 12 | Bg2/+;Mef2/GAL80ts | 15d | M | none | 29 | 10756.00 | 0.07 | 157295.07 | thorax |
| 12 | Bg2/+;Mef2/GAL80ts | 15d | M | none | 29 | 7938.00 | 0.09 | 86793.98 | thorax |
| 12 | Bg2/+;Mef2/GAL80ts | 15d | M | none | 29 | 6490.00 | 0.07 | 88701.96 | thorax |
| 12 | Bg2/+;Mef2/GAL80ts | 15d | M | none | 29 | 11486.00 | 0.07 | 155747.10 | thorax |
| 12 | Bg2/+;Mef2/GAL80ts | 15d | M | none | 29 | 9776.00 | 0.10 | 94998.42 | thorax |
| 13 | Bg2/+;Mef2/GAL80ts | 15d | M | none | 29 | 9618.00 | 0.10 | 99380.07 | thorax |
| 13 | Bg2/+;Mef2/GAL80ts | 15d | M | none | 29 | 9366.00 | 0.11 | 84821.29 | thorax |
| 13 | Bg2/+;Mef2/GAL80ts | 15d | M | none | 29 | 12144.00 | 0.10 | 126474.08 | thorax |
| 13 | Bg2/+;Mef2/GAL80ts | 15d | M | none | 29 | 11390.00 | 0.09 | 132784.36 | thorax |
| 13 | Bg2/+;Mef2/GAL80ts | 15d | M | none | 29 | 10026.00 | 0.11 | 93098.57 | thorax |
| 14 | Bg2/+;Mef2/GAL80ts | 15d | M | none | 29 | 12490.00 | 0.10 | 128422.82 | thorax |
| 14 | Bg2/+;Mef2/GAL80ts | 15d | M | none | 29 | 10160.00 | 0.08 | 124537.64 | thorax |
| 14 | Bg2/+;Mef2/GAL80ts | 15d | M | none | 29 | 10396.00 | 0.07 | 141144.24 | thorax |
| 14 | Bg2/+;Mef2/GAL80ts | 15d | M | none | 29 | 9622.00 | 0.09 | 111549.63 | thorax |
| 14 | Bg2/+;Mef2/GAL80ts | 15d | M | none | 29 | 9354.00 | 0.08 | 123779.27 | thorax |
| 15 | Bg2/+;Mef2/GAL80ts | 15d | M | none | 29 | 9124.00 | 0.08 | 112144.80 | thorax |
| 15 | Bg2/+;Mef2/GAL80ts | 15d | M | none | 29 | 10904.00 | 0.08 | 128400.75 | thorax |
| 15 | Bg2/+;Mef2/GAL80ts | 15d | M | none | 29 | 11070.00 | 0.07 | 167964.81 | thorax |
| 15 | Bg2/+;Mef2/GAL80ts | 15d | M | none | 29 | 12376.00 | 0.09 | 143329.27 | thorax |
| 15 | Bg2/+;Mef2/GAL80ts | 15d | M | none | 29 | 10300.00 | 0.08 | 125704.78 | thorax |
| 12 | Bg2/+;Mef2/GAL80ts | 15d | M | none | 20-29 | 8050.00 | 0.10 | 78490.94 | thorax |
| 12 | Bg2/+;Mef2/GAL80ts | 15d | M | none | 20-29 | 9928.00 | 0.09 | 113202.02 | thorax |
| 12 | Bg2/+;Mef2/GAL80ts | 15d | M | none | 20-29 | 7334.00 | 0.12 | 59086.35 | thorax |
| 12 | Bg2/+;Mef2/GAL80ts | 15d | M | none | 20-29 | 10722.00 | 0.11 | 96922.80 | thorax |
| 12 | Bg2/+;Mef2/GAL80ts | 15d | M | none | 20-29 | 10576.00 | 0.11 | 93527.91 | thorax |
| 13 | Bg2/+;Mef2/GAL80ts | 15d | M | none | 20-29 | 7312.00 | 0.13 | 58237.93 | thorax |
| 13 | Bg2/+;Mef2/GAL80ts | 15d | M | none | 20-29 | 7700.00 | 0.10 | 78012.08 | thorax |
| 13 | Bg2/+;Mef2/GAL80ts | 15d | M | none | 20-29 | 11492.00 | 0.10 | 116949.81 | thorax |
| 13 | Bg2/+;Mef2/GAL80ts | 15d | M | none | 20-29 | 11476.00 | 0.10 | 118424.43 | thorax |
| 13 | Bg2/+;Mef2/GAL80ts | 15d | M | none | 20-29 | 9866.00 | 0.11 | 87147.76 | thorax |
| 14 | Bg2/+;Mef2/GAL80ts | 15d | M | none | 20-29 | 7672.00 | 0.10 | 80706.79 | thorax |
| 14 | Bg2/+;Mef2/GAL80ts | 15d | M | none | 20-29 | 6698.00 | 0.11 | 63738.47 | thorax |
| 14 | Bg2/+;Mef2/GAL80ts | 15d | M | none | 20-29 | 5114.00 | 0.11 | 47531.39 | thorax |
| 14 | Bg2/+;Mef2/GAL80ts | 15d | M | none | 20-29 | 8668.00 | 0.09 | 93561.53 | thorax |
| 14 | Bg2/+;Mef2/GAL80ts | 15d | M | none | 20-29 | 8354.00 | 0.10 | 81151.03 | thorax |
| 15 | Bg2/+;Mef2/GAL80ts | 15d | M | none | 20-29 | 6936.00 | 0.12 | 60088.27 | thorax |
| 15 | Bg2/+;Mef2/GAL80ts | 15d | M | none | 20-29 | 6500.00 | 0.11 | 57935.01 | thorax |
| 15 | Bg2/+;Mef2/GAL80ts | 15d | M | none | 20-29 | 5458.00 | 0.12 | 44841.02 | thorax |
| 15 | Bg2/+;Mef2/GAL80ts | 15d | M | none | 20-29 | 6446.00 | 0.09 | 75763.81 | thorax |

| Rep. | Genotype | Stage | Sex | Treatmen | °C | ß-gal ΔmOD/min/ ml | Protein mg/ml | Specific Activity ΔmOD/min/ mg | Whole/ dissected |
|---|---|---|---|---|---|---|---|---|---|
| 15 | Bg2/+;Mef2/GAL80ts | 15d | M | none | 20-29 | 9000.00 | 0.09 | 98012.90 | thorax |
| 12 | Bg2/+;Mef2/GAL80ts | 15d | M | none | 29-20 | 5284.00 | 0.08 | 67778.09 | thorax |
| 12 | Bg2/+;Mef2/GAL80ts | 15d | M | none | 29-20 | 6308.00 | 0.08 | 81325.34 | thorax |
| 12 | Bg2/+;Mef2/GAL80ts | 15d | M | none | 29-20 | 10516.00 | 0.09 | 121045.65 | thorax |
| 12 | Bg2/+;Mef2/GAL80ts | 15d | M | none | 29-20 | 4478.00 | 0.07 | 60292.79 | thorax |
| 13 | Bg2/+;Mef2/GAL80ts | 15d | M | none | 29-20 | 6878.00 | 0.10 | 71867.05 | thorax |
| 13 | Bg2/+;Mef2/GAL80ts | 15d | M | none | 29-20 | 7588.00 | 0.06 | 119808.31 | thorax |
| 13 | Bg2/+;Mef2/GAL80ts | 15d | M | none | 29-20 | 9550.00 | 0.05 | 206098.96 | thorax |
| 13 | Bg2/+;Mef2/GAL80ts | 15d | M | none | 29-20 | 6408.00 | 0.09 | 70515.87 | thorax |
| 13 | Bg2/+;Mef2/GAL80ts | 15d | M | none | 29-20 | 7870.00 | 0.07 | 117267.12 | thorax |
| 14 | Bg2/+;Mef2/GAL80ts | 15d | M | none | 29-20 | 9584.00 | 0.07 | 141143.93 | thorax |
| 14 | Bg2/+;Mef2/GAL80ts | 15d | M | none | 29-20 | 11644.00 | 0.10 | 116982.52 | thorax |
| 14 | Bg2/+;Mef2/GAL80ts | 15d | M | none | 29-20 | 14362.00 | 0.10 | 142506.29 | thorax |
| 14 | Bg2/+;Mef2/GAL80ts | 15d | M | none | 29-20 | 12914.00 | 0.09 | 136082.01 | thorax |
| 14 | Bg2/+;Mef2/GAL80ts | 15d | M | none | 29-20 | 15302.00 | 0.09 | 166364.60 | thorax |
| 15 | Bg2/+;Mef2/GAL80ts | 15d | M | none | 29-20 | 10112.00 | 0.10 | 99236.52 | thorax |
| 15 | Bg2/+;Mef2/GAL80ts | 15d | M | none | 29-20 | 13796.00 | 0.09 | 160319.98 | thorax |
| 15 | Bg2/+;Mef2/GAL80ts | 15d | M | none | 29-20 | 11450.00 | 0.08 | 138878.96 | thorax |
| 15 | Bg2/+;Mef2/GAL80ts | 15d | M | none | 29-20 | 11810.00 | 0.08 | 150044.34 | thorax |
| 15 | Bg2/+;Mef2/GAL80ts | 15d | M | none | 29-20 | 11382.00 | 0.11 | 107531.04 | thorax |
| 12 | Bg2/+;Mef2/GAL80ts | LP | ND | none | 20 | 20.00 | 0.22 | 90.48 | whole |
| 12 | Bg2/+;Mef2/GAL80ts | LP | ND | none | 20 | 48.00 | 0.36 | 132.43 | whole |
| 12 | Bg2/+;Mef2/GAL80ts | LP | ND | none | 20 | 54.00 | 0.42 | 128.25 | whole |
| 12 | Bg2/+;Mef2/GAL80ts | LP | ND | none | 20 | 90.00 | 0.18 | 491.16 | whole |
| 13 | Bg2/+;Mef2/GAL80ts | LP | ND | none | 20 | 60.00 | 0.22 | 269.15 | whole |
| 13 | Bg2/+;Mef2/GAL80ts | LP | ND | none | 20 | 40.00 | 0.18 | 222.11 | whole |
| 13 | Bg2/+;Mef2/GAL80ts | LP | ND | none | 20 | 8.00 | 0.18 | 45.67 | whole |
| 13 | Bg2/+;Mef2/GAL80ts | LP | ND | none | 20 | 64.00 | 0.27 | 241.45 | whole |
| 13 | Bg2/+;Mef2/GAL80ts | LP | ND | none | 20 | 50.00 | 0.91 | 54.87 | whole |
| 14 | Bg2/+;Mef2/GAL80ts | LP | ND | none | 20 | 38.00 | 0.23 | 162.63 | whole |
| 14 | Bg2/+;Mef2/GAL80ts | LP | ND | none | 20 | 75.38 | 0.23 | 330.14 | whole |
| 14 | Bg2/+;Mef2/GAL80ts | LP | ND | none | 20 | 24.44 | 0.25 | 98.78 | whole |
| 14 | Bg2/+;Mef2/GAL80ts | LP | ND | none | 20 | 50.00 | 0.27 | 184.31 | whole |
| 14 | Bg2/+;Mef2/GAL80ts | LP | ND | none | 20 | 34.00 | 0.38 | 90.61 | whole |
| 15 | Bg2/+;Mef2/GAL80ts | LP | ND | none | 20 | 58.00 | 0.25 | 235.59 | whole |
| 15 | Bg2/+;Mef2/GAL80ts | LP | ND | none | 20 | 37.14 | 0.30 | 122.76 | whole |
| 15 | Bg2/+;Mef2/GAL80ts | LP | ND | none | 20 | 17.50 | 0.23 | 76.66 | whole |
| 15 | Bg2/+;Mef2/GAL80ts | LP | ND | none | 20 | 17.33 | 0.31 | 56.32 | whole |
| 15 | Bg2/+;Mef2/GAL80ts | LP | ND | none | 20 | 88.00 | 0.21 | 413.35 | whole |
| 12 | Bg2/+;Mef2/GAL80ts | LP | ND | none | 29 | 3225.00 | 0.35 | 9215.82 | whole |

| Rep. | Genotype | Stage | Sex | Treatmen | °C | ß-gal ΔmOD/min/ml | Protein mg/ml | Specific Activity ΔmOD/min/mg | Whole/ dissected |
|---|---|---|---|---|---|---|---|---|---|
| 12 | Bg2/+;Mef2/GAL80ts | LP | ND | none | 29 | 180.00 | 0.18 | 988.30 | whole |
| 12 | Bg2/+;Mef2/GAL80ts | LP | ND | none | 29 | 270.00 | 0.18 | 1475.23 | whole |
| 12 | Bg2/+;Mef2/GAL80ts | LP | ND | none | 29 | 310.00 | 0.18 | 1716.75 | whole |
| 12 | Bg2/+;Mef2/GAL80ts | LP | ND | none | 29 | 2960.00 | 0.24 | 12265.78 | whole |
| 13 | Bg2/+;Mef2/GAL80ts | LP | ND | none | 29 | 865.00 | 0.21 | 4122.28 | whole |
| 13 | Bg2/+;Mef2/GAL80ts | LP | ND | none | 29 | 230.00 | 0.23 | 1006.45 | whole |
| 13 | Bg2/+;Mef2/GAL80ts | LP | ND | none | 29 | 1540.00 | 0.42 | 3641.25 | whole |
| 13 | Bg2/+;Mef2/GAL80ts | LP | ND | none | 29 | 235.00 | 0.22 | 1078.74 | whole |
| 13 | Bg2/+;Mef2/GAL80ts | LP | ND | none | 29 | 240.00 | 0.22 | 1110.20 | whole |
| 14 | Bg2/+;Mef2/GAL80ts | LP | ND | none | 29 | 2415.00 | 0.24 | 10221.63 | whole |
| 14 | Bg2/+;Mef2/GAL80ts | LP | ND | none | 29 | 6835.00 | 0.20 | 33902.75 | whole |
| 14 | Bg2/+;Mef2/GAL80ts | LP | ND | none | 29 | 3610.00 | 0.25 | 14440.00 | whole |
| 14 | Bg2/+;Mef2/GAL80ts | LP | ND | none | 29 | 5125.00 | 0.36 | 14088.03 | whole |
| 14 | Bg2/+;Mef2/GAL80ts | LP | ND | none | 29 | 2325.00 | 0.20 | 11562.69 | whole |
| 15 | Bg2/+;Mef2/GAL80ts | LP | ND | none | 29 | 4340.00 | 0.25 | 17197.57 | whole |
| 15 | Bg2/+;Mef2/GAL80ts | LP | ND | none | 29 | 17670.00 | 0.24 | 74093.43 | whole |
| 15 | Bg2/+;Mef2/GAL80ts | LP | ND | none | 29 | 7450.00 | 0.40 | 18815.21 | whole |
| 15 | Bg2/+;Mef2/GAL80ts | LP | ND | none | 29 | 19520.00 | 0.21 | 92046.48 | whole |
| 15 | Bg2/+;Mef2/GAL80ts | LP | ND | none | 29 | 12425.00 | 0.14 | 88546.84 | whole |
| 5 | Bg2/+;MHC | 0-1d | F | none | 25 | 12040.00 | 0.12 | 100253.72 | thorax |
| 5 | Bg2/+;MHC | 0-1d | F | none | 25 | 12280.00 | 0.12 | 103094.21 | thorax |
| 5 | Bg2/+;MHC | 0-1d | F | none | 25 | 5560.00 | 0.18 | 30685.35 | thorax |
| 5 | Bg2/+;MHC | 0-1d | F | none | 25 | 22480.00 | 0.12 | 192346.86 | thorax |
| 5 | Bg2/+;MHC | 0-1d | F | none | 25 | 18680.00 | 0.16 | 117341.97 | thorax |
| 4 | Bg2/+;MHC | 1-2d | F | none | 25 | 7280.00 | 0.12 | 62032.53 | thorax |
| 4 | Bg2/+;MHC | 1-2d | F | none | 25 | 9200.00 | 0.11 | 82503.93 | thorax |
| 4 | Bg2/+;MHC | 1-2d | F | none | 25 | 4960.00 | 0.07 | 67120.58 | thorax |
| 4 | Bg2/+;MHC | 1-2d | F | none | 25 | 12280.00 | 0.13 | 92210.30 | thorax |
| 4 | Bg2/+;MHC | 1-2d | F | none | 25 | 5640.00 | 0.08 | 69338.82 | thorax |
| 5 | Bg2/+;MHC | 0-1d | M | none | 25 | 7520.00 | 0.11 | 68798.36 | thorax |
| 5 | Bg2/+;MHC | 0-1d | M | none | 25 | 4880.00 | 0.06 | 83310.24 | thorax |
| 5 | Bg2/+;MHC | 0-1d | M | none | 25 | 5280.00 | 0.06 | 85827.06 | thorax |
| 5 | Bg2/+;MHC | 0-1d | M | none | 25 | 4400.00 | 0.13 | 34203.05 | thorax |
| 5 | Bg2/+;MHC | 0-1d | M | none | 25 | 7200.00 | 0.11 | 64954.74 | thorax |
| 4 | Bg2/+;MHC | 1-2d | M | none | 25 | 5760.00 | 0.10 | 56577.34 | thorax |
| 4 | Bg2/+;MHC | 1-2d | M | none | 25 | 6160.00 | 0.09 | 71524.44 | thorax |
| 4 | Bg2/+;MHC | 1-2d | M | none | 25 | 5840.00 | 0.09 | 65976.22 | thorax |
| 4 | Bg2/+;MHC | 1-2d | M | none | 25 | 8040.00 | 0.12 | 69372.66 | thorax |
| 4 | Bg2/+;MHC | 1-2d | M | none | 25 | 3920.00 | 0.05 | 74480.00 | thorax |
| 4 | Bg2/+;MHC | EP | ND | none | 25 | 4000.00 | 0.49 | 8236.73 | whole |

| Rep. | Genotype | Stage | Sex | Treatmen | °C | ß-gal ΔmOD/min/ ml | Protein mg/ml | Specific Activity ΔmOD/min/ mg | Whole/ dissected |
|---|---|---|---|---|---|---|---|---|---|
| 4 | Bg2/+;MHC | EP | ND | none | 25 | 4320.00 | 0.45 | 9590.50 | whole |
| 4 | Bg2/+;MHC | EP | ND | none | 25 | 4600.00 | 0.41 | 11090.56 | whole |
| 4 | Bg2/+;MHC | EP | ND | none | 25 | 6400.00 | 0.58 | 11082.97 | whole |
| 4 | Bg2/+;MHC | EP | ND | none | 25 | 2680.00 | 0.45 | 5938.23 | whole |
| 5 | Bg2/+;MHC | EP | ND | none | 25 | 4040.00 | 0.45 | 8967.11 | whole |
| 5 | Bg2/+;MHC | EP | ND | none | 25 | 2760.00 | 0.45 | 6176.73 | whole |
| 5 | Bg2/+;MHC | EP | ND | none | 25 | 1960.00 | 0.32 | 6204.29 | whole |
| 5 | Bg2/+;MHC | EP | ND | none | 25 | 7280.00 | 0.62 | 11730.70 | whole |
| 5 | Bg2/+;MHC | EP | ND | none | 25 | 4520.00 | 0.38 | 12047.19 | whole |
| 4 | Bg2/+;MHC | LP | ND | none | 25 | 8360.00 | 0.29 | 28481.45 | whole |
| 4 | Bg2/+;MHC | LP | ND | none | 25 | 13120.00 | 0.27 | 48801.74 | whole |
| 4 | Bg2/+;MHC | LP | ND | none | 25 | 13040.00 | 0.35 | 36791.25 | whole |
| 4 | Bg2/+;MHC | LP | ND | none | 25 | 5960.00 | 0.28 | 21156.18 | whole |
| 4 | Bg2/+;MHC | LP | ND | none | 25 | 10400.00 | 0.35 | 29320.76 | whole |
| 5 | Bg2/+;MHC | LP | ND | none | 25 | 2320.00 | 0.13 | 17512.94 | whole |
| 5 | Bg2/+;MHC | LP | ND | none | 25 | 9440.00 | 0.24 | 38597.46 | whole |
| 5 | Bg2/+;MHC | LP | ND | none | 25 | 6920.00 | 0.26 | 26609.36 | whole |
| 5 | Bg2/+;MHC | LP | ND | none | 25 | 10880.00 | 0.32 | 33481.98 | whole |
| 5 | Bg2/+;MHC | LP | ND | none | 25 | 17160.00 | 0.39 | 44128.81 | whole |
| 6 | Bg2/+;MHC/+ | 10-11d | F | -RU486 | 25 | 8300.00 | 0.10 | 79751.82 | thorax |
| 6 | Bg2/+;MHC/+ | 10-11d | F | -RU486 | 25 | 8395.00 | 0.12 | 70116.19 | thorax |
| 6 | Bg2/+;MHC/+ | 10-11d | F | -RU486 | 25 | 10365.00 | 0.13 | 82013.59 | thorax |
| 6 | Bg2/+;MHC/+ | 10-11d | F | -RU486 | 25 | 8235.00 | 0.12 | 69015.80 | thorax |
| 6 | Bg2/+;MHC/+ | 10-11d | F | -RU486 | 25 | 10500.00 | 0.13 | 80223.61 | thorax |
| 7 | Bg2/+;MHC/+ | 10-12d | F | -RU486 | 25 | 8500.00 | 0.15 | 56205.17 | thorax |
| 7 | Bg2/+;MHC/+ | 10-12d | F | -RU486 | 25 | 10500.00 | 0.14 | 72673.77 | thorax |
| 7 | Bg2/+;MHC/+ | 10-12d | F | -RU486 | 25 | 5960.00 | 0.14 | 42220.00 | thorax |
| 7 | Bg2/+;MHC/+ | 10-12d | F | -RU486 | 25 | 6820.00 | 0.13 | 52068.79 | thorax |
| 7 | Bg2/+;MHC/+ | 10-12d | F | -RU486 | 25 | 6820.00 | 0.13 | 53421.22 | thorax |
| 6 | Bg2/+;MHC/+ | 10-11d | F | +RU486 | 25 | 31345.00 | 0.12 | 254193.64 | thorax |
| 6 | Bg2/+;MHC/+ | 10-11d | F | +RU486 | 25 | 25550.00 | 0.11 | 229269.61 | thorax |
| 6 | Bg2/+;MHC/+ | 10-11d | F | +RU486 | 25 | 15595.00 | 0.08 | 191209.96 | thorax |
| 6 | Bg2/+;MHC/+ | 10-11d | F | +RU486 | 25 | 18045.00 | 0.10 | 174589.84 | thorax |
| 6 | Bg2/+;MHC/+ | 10-11d | F | +RU486 | 25 | 30700.00 | 0.10 | 319489.24 | thorax |
| 7 | Bg2/+;MHC/+ | 10-12d | F | +RU486 | 25 | 12500.00 | 0.12 | 105761.52 | thorax |
| 7 | Bg2/+;MHC/+ | 10-12d | F | +RU486 | 25 | 12560.00 | 0.12 | 101392.58 | thorax |
| 7 | Bg2/+;MHC/+ | 10-12d | F | +RU486 | 25 | 19800.00 | 0.14 | 139908.95 | thorax |
| 7 | Bg2/+;MHC/+ | 10-12d | F | +RU486 | 25 | 17240.00 | 0.15 | 118739.45 | thorax |
| 7 | Bg2/+;MHC/+ | 10-12d | F | +RU486 | 25 | 20160.00 | 0.19 | 107333.57 | thorax |
| 5 | Bg2/+;MHC/+ | 0-1d | F | none | 25 | 10080.00 | 0.18 | 55246.45 | thorax |

| Rep. | Genotype | Stage | Sex | Treatmen | °C | ß-gal ΔmOD/min/ml | Protein mg/ml | Specific Activity ΔmOD/min/mg | Whole/ dissected |
|---|---|---|---|---|---|---|---|---|---|
| 5 | Bg2/+;MHC/+ | 0-1d | F | none | 25 | 10840.00 | 0.13 | 83176.60 | thorax |
| 5 | Bg2/+;MHC/+ | 0-1d | F | none | 25 | 7120.00 | 0.13 | 56017.99 | thorax |
| 5 | Bg2/+;MHC/+ | 0-1d | F | none | 25 | 5520.00 | 0.20 | 28297.93 | thorax |
| 5 | Bg2/+;MHC/+ | 0-1d | F | none | 25 | 6160.00 | 0.14 | 45131.17 | thorax |
| 1 | Bg2/+;MHC/+ | 0-24h | F | none | 25 | 4269.66 | 0.18 | 23125.71 | thorax |
| 1 | Bg2/+;MHC/+ | 0-24h | F | none | 25 | 3048.69 | 0.18 | 17183.34 | thorax |
| 1 | Bg2/+;MHC/+ | 0-24h | F | none | 25 | 4217.23 | 0.21 | 19972.79 | thorax |
| 1 | Bg2/+;MHC/+ | 0-24h | F | none | 25 | 4037.45 | 0.19 | 20911.39 | thorax |
| 4 | Bg2/+;MHC/+ | 1-2d | F | none | 25 | 10880.00 | 0.13 | 83276.83 | thorax |
| 4 | Bg2/+;MHC/+ | 1-2d | F | none | 25 | 5640.00 | 0.10 | 56430.00 | thorax |
| 4 | Bg2/+;MHC/+ | 1-2d | F | none | 25 | 9920.00 | 0.08 | 126935.51 | thorax |
| 4 | Bg2/+;MHC/+ | 1-2d | F | none | 25 | 5200.00 | 0.09 | 55812.84 | thorax |
| 4 | Bg2/+;MHC/+ | 1-2d | F | none | 25 | 8800.00 | 0.07 | 129066.67 | thorax |
| 1 | Bg2/+;MHC/+ | 10-11d | F | none | 25 | 2440.00 | 0.21 | 11467.30 | thorax |
| 1 | Bg2/+;MHC/+ | 10-11d | F | none | 25 | 1833.33 | 0.15 | 11950.00 | thorax |
| 1 | Bg2/+;MHC/+ | 10-11d | F | none | 25 | 5260.00 | 0.19 | 27230.47 | thorax |
| 1 | Bg2/+;MHC/+ | 10-11d | F | none | 25 | 4343.33 | 0.15 | 29943.94 | thorax |
| 1 | Bg2/+;MHC/+ | 10-11d | F | none | 25 | 2816.67 | 0.12 | 23817.69 | thorax |
| 1 | Bg2/+;MHC/+ | 3-5d | F | none | 25 | 6850.00 | 0.20 | 35002.55 | thorax |
| 1 | Bg2/+;MHC/+ | 3-5d | F | none | 25 | 6120.00 | 0.21 | 28624.88 | thorax |
| 1 | Bg2/+;MHC/+ | 3-5d | F | none | 25 | 5040.00 | 0.17 | 29857.82 | thorax |
| 1 | Bg2/+;MHC/+ | 3-5d | F | none | 25 | 4360.00 | 0.23 | 18568.99 | thorax |
| 1 | Bg2/+;MHC/+ | 3-5d | F | none | 25 | 5550.00 | 0.21 | 26466.38 | thorax |
| 6 | Bg2/+;MHC/+ | 10-11d | M | -RU486 | 25 | 5865.00 | 0.08 | 76213.80 | thorax |
| 6 | Bg2/+;MHC/+ | 10-11d | M | -RU486 | 25 | 4760.00 | 0.09 | 54466.89 | thorax |
| 6 | Bg2/+;MHC/+ | 10-11d | M | -RU486 | 25 | 4365.00 | 0.06 | 75898.19 | thorax |
| 6 | Bg2/+;MHC/+ | 10-11d | M | -RU486 | 25 | 6005.00 | 0.02 | 264328.20 | thorax |
| 6 | Bg2/+;MHC/+ | 10-11d | M | -RU486 | 25 | 4710.00 | 0.08 | 56061.05 | thorax |
| 7 | Bg2/+;MHC/+ | 10-12d | M | -RU486 | 25 | 5080.00 | 0.08 | 60844.71 | thorax |
| 7 | Bg2/+;MHC/+ | 10-12d | M | -RU486 | 25 | 5160.00 | 0.10 | 50487.88 | thorax |
| 7 | Bg2/+;MHC/+ | 10-12d | M | -RU486 | 25 | 4560.00 | 0.07 | 64605.10 | thorax |
| 7 | Bg2/+;MHC/+ | 10-12d | M | -RU486 | 25 | 5520.00 | 0.07 | 76411.28 | thorax |
| 7 | Bg2/+;MHC/+ | 10-12d | M | -RU486 | 25 | 4820.00 | 0.09 | 51979.67 | thorax |
| 6 | Bg2/+;MHC/+ | 10-11d | M | +RU486 | 25 | 15055.00 | 0.07 | 228089.09 | thorax |
| 6 | Bg2/+;MHC/+ | 10-11d | M | +RU486 | 25 | 23630.00 | 0.08 | 291924.60 | thorax |
| 6 | Bg2/+;MHC/+ | 10-11d | M | +RU486 | 25 | 26220.00 | 0.08 | 346714.26 | thorax |
| 6 | Bg2/+;MHC/+ | 10-11d | M | +RU486 | 25 | 32375.00 | 0.07 | 461178.57 | thorax |
| 6 | Bg2/+;MHC/+ | 10-11d | M | +RU486 | 25 | 26145.00 | 0.07 | 391854.20 | thorax |
| 7 | Bg2/+;MHC/+ | 10-12d | M | +RU486 | 25 | 34400.00 | 0.11 | 311666.95 | thorax |
| 7 | Bg2/+;MHC/+ | 10-12d | M | +RU486 | 25 | 27340.00 | 0.08 | 354078.16 | thorax |

| Rep. | Genotype | Stage | Sex | Treatmen | °C | ß-gal ΔmOD/min/ ml | Protein mg/ml | Specific Activity ΔmOD/min/ mg | Whole/ dissected |
|---|---|---|---|---|---|---|---|---|---|
| 7 | Bg2/+;MHC/+ | 10-12d | M | +RU486 | 25 | 26280.00 | 0.12 | 219711.21 | thorax |
| 7 | Bg2/+;MHC/+ | 10-12d | M | +RU486 | 25 | 28920.00 | 0.11 | 255172.92 | thorax |
| 7 | Bg2/+;MHC/+ | 10-12d | M | +RU486 | 25 | 26940.00 | 0.09 | 313767.39 | thorax |
| 5 | Bg2/+;MHC/+ | 0-1d | M | none | 25 | 3880.00 | 0.06 | 67696.04 | thorax |
| 5 | Bg2/+;MHC/+ | 0-1d | M | none | 25 | 4760.00 | 0.06 | 83663.45 | thorax |
| 5 | Bg2/+;MHC/+ | 0-1d | M | none | 25 | 7120.00 | 0.18 | 40646.66 | thorax |
| 5 | Bg2/+;MHC/+ | 0-1d | M | none | 25 | 4080.00 | 0.09 | 43455.04 | thorax |
| 5 | Bg2/+;MHC/+ | 0-1d | M | none | 25 | 5360.00 | 0.06 | 83513.01 | thorax |
| 1 | Bg2/+;MHC/+ | 0-24h | M | none | 25 | 2812.73 | 0.14 | 20548.81 | thorax |
| 1 | Bg2/+;MHC/+ | 0-24h | M | none | 25 | 2539.33 | 0.17 | 15174.44 | thorax |
| 1 | Bg2/+;MHC/+ | 0-24h | M | none | 25 | 2498.13 | 0.16 | 15442.65 | thorax |
| 1 | Bg2/+;MHC/+ | 0-24h | M | none | 25 | 3082.40 | 0.14 | 21586.50 | thorax |
| 1 | Bg2/+;MHC/+ | 0-24h | M | none | 25 | 5161.05 | 0.16 | 31937.36 | thorax |
| 4 | Bg2/+;MHC/+ | 1-2d | M | none | 25 | 3160.00 | 0.08 | 40712.05 | thorax |
| 4 | Bg2/+;MHC/+ | 1-2d | M | none | 25 | 4520.00 | 0.04 | 114122.42 | thorax |
| 4 | Bg2/+;MHC/+ | 1-2d | M | none | 25 | 7640.00 | 0.04 | 198904.36 | thorax |
| 4 | Bg2/+;MHC/+ | 1-2d | M | none | 25 | 6040.00 | 0.14 | 44597.61 | thorax |
| 4 | Bg2/+;MHC/+ | 1-2d | M | none | 25 | 7120.00 | 0.14 | 51609.71 | thorax |
| 1 | Bg2/+;MHC/+ | 10-11d | M | none | 25 | 2280.00 | 0.10 | 22220.85 | thorax |
| 1 | Bg2/+;MHC/+ | 10-11d | M | none | 25 | 2186.67 | 0.18 | 12369.55 | thorax |
| 1 | Bg2/+;MHC/+ | 10-11d | M | none | 25 | 1923.33 | 0.09 | 20381.23 | thorax |
| 1 | Bg2/+;MHC/+ | 10-11d | M | none | 25 | 1896.67 | 0.15 | 12251.44 | thorax |
| 1 | Bg2/+;MHC/+ | 10-11d | M | none | 25 | 1710.00 | 0.17 | 9957.93 | thorax |
| 1 | Bg2/+;MHC/+ | 3-5d | M | none | 25 | 2965.00 | 0.11 | 28051.09 | thorax |
| 1 | Bg2/+;MHC/+ | 3-5d | M | none | 25 | 4875.00 | 0.19 | 25752.77 | thorax |
| 1 | Bg2/+;MHC/+ | 3-5d | M | none | 25 | 2130.00 | 0.12 | 17588.77 | thorax |
| 1 | Bg2/+;MHC/+ | 3-5d | M | none | 25 | 1920.00 | 0.14 | 14004.38 | thorax |
| 1 | Bg2/+;MHC/+ | 3-5d | M | none | 25 | 1585.00 | 0.19 | 8161.69 | thorax |
| 6 | Bg2/+;MHC/+ | EP | ND | -RU486 | 25 | 7480.00 | 1.17 | 6412.20 | whole |
| 6 | Bg2/+;MHC/+ | EP | ND | -RU486 | 25 | 10040.00 | 1.49 | 6752.01 | whole |
| 6 | Bg2/+;MHC/+ | EP | ND | -RU486 | 25 | 5350.00 | 1.18 | 4538.01 | whole |
| 6 | Bg2/+;MHC/+ | EP | ND | -RU486 | 25 | 11245.00 | 1.33 | 8458.20 | whole |
| 6 | Bg2/+;MHC/+ | EP | ND | -RU486 | 25 | 6185.00 | 0.68 | 9112.72 | whole |
| 7 | Bg2/+;MHC/+ | EP | ND | -RU486 | 25 | 5320.00 | 1.17 | 4560.55 | whole |
| 7 | Bg2/+;MHC/+ | EP | ND | -RU486 | 25 | 7930.00 | 1.49 | 5333.01 | whole |
| 7 | Bg2/+;MHC/+ | EP | ND | -RU486 | 25 | 6775.00 | 1.18 | 5746.73 | whole |
| 7 | Bg2/+;MHC/+ | EP | ND | -RU486 | 25 | 10410.00 | 1.33 | 7830.14 | whole |
| 7 | Bg2/+;MHC/+ | EP | ND | -RU486 | 25 | 6770.00 | 0.68 | 9974.63 | whole |
| 6 | Bg2/+;MHC/+ | EP | ND | +RU486 | 25 | 8910.00 | 1.23 | 7263.62 | whole |
| 6 | Bg2/+;MHC/+ | EP | ND | +RU486 | 25 | 14720.00 | 1.40 | 10510.18 | whole |

| Rep. | Genotype | Stage | Sex | Treatmen | °C | ß-gal ΔmOD/min/ml | Protein mg/ml | Specific Activity ΔmOD/min/mg | Whole/ dissected |
|---|---|---|---|---|---|---|---|---|---|
| 6 | Bg2/+;MHC/+ | EP | ND | +RU486 | 25 | 16510.00 | 1.22 | 13542.87 | whole |
| 6 | Bg2/+;MHC/+ | EP | ND | +RU486 | 25 | 10775.00 | 1.41 | 7657.79 | whole |
| 6 | Bg2/+;MHC/+ | EP | ND | +RU486 | 25 | 12955.00 | 1.55 | 8344.26 | whole |
| 7 | Bg2/+;MHC/+ | EP | ND | +RU486 | 25 | 4925.00 | 1.23 | 4014.96 | whole |
| 7 | Bg2/+;MHC/+ | EP | ND | +RU486 | 25 | 7270.00 | 1.40 | 5190.83 | whole |
| 7 | Bg2/+;MHC/+ | EP | ND | +RU486 | 25 | 10035.00 | 1.22 | 8231.54 | whole |
| 7 | Bg2/+;MHC/+ | EP | ND | +RU486 | 25 | 8070.00 | 1.41 | 5735.34 | whole |
| 7 | Bg2/+;MHC/+ | EP | ND | +RU486 | 25 | 10285.00 | 1.55 | 6624.52 | whole |
| 4 | Bg2/+;MHC/+ | EP | ND | none | 25 | 5440.00 | 0.48 | 11282.55 | whole |
| 4 | Bg2/+;MHC/+ | EP | ND | none | 25 | 4040.00 | 0.45 | 8907.64 | whole |
| 4 | Bg2/+;MHC/+ | EP | ND | none | 25 | 3480.00 | 0.41 | 8458.46 | whole |
| 4 | Bg2/+;MHC/+ | EP | ND | none | 25 | 4000.00 | 0.41 | 9658.39 | whole |
| 4 | Bg2/+;MHC/+ | EP | ND | none | 25 | 3000.00 | 0.49 | 6150.88 | whole |
| 5 | Bg2/+;MHC/+ | EP | ND | none | 25 | 3480.00 | 0.45 | 7726.33 | whole |
| 5 | Bg2/+;MHC/+ | EP | ND | none | 25 | 3200.00 | 0.45 | 7104.67 | whole |
| 5 | Bg2/+;MHC/+ | EP | ND | none | 25 | 3640.00 | 0.45 | 8139.16 | whole |
| 5 | Bg2/+;MHC/+ | EP | ND | none | 25 | 5280.00 | 0.46 | 11384.37 | whole |
| 5 | Bg2/+;MHC/+ | EP | ND | none | 25 | 600.00 | 0.22 | 2724.93 | whole |
| 2 | Bg2/+;MHC/+ | L3 | ND | none | 25 | 6120.00 | 0.53 | 11649.24 | whole |
| 2 | Bg2/+;MHC/+ | L3 | ND | none | 25 | 18960.00 | 0.87 | 21812.39 | whole |
| 2 | Bg2/+;MHC/+ | L3 | ND | none | 25 | 4640.00 | 0.55 | 8418.92 | whole |
| 2 | Bg2/+;MHC/+ | L3 | ND | none | 25 | 4320.00 | 0.50 | 8571.62 | whole |
| 2 | Bg2/+;MHC/+ | L3 | ND | none | 25 | 6280.00 | 0.68 | 9255.85 | whole |
| 3 | Bg2/+;MHC/+ | L3 | ND | none | 25 | 15960.00 | 0.75 | 21345.13 | whole |
| 3 | Bg2/+;MHC/+ | L3 | ND | none | 25 | 12440.00 | 0.65 | 19132.34 | whole |
| 3 | Bg2/+;MHC/+ | L3 | ND | none | 25 | 6440.00 | 0.45 | 14395.57 | whole |
| 3 | Bg2/+;MHC/+ | L3 | ND | none | 25 | 7440.00 | 0.40 | 18516.54 | whole |
| 3 | Bg2/+;MHC/+ | L3 | ND | none | 25 | 4000.00 | 0.38 | 10655.23 | whole |
| 1 | Bg2/+;MHC/+ | LP | ND | none | 25 | 3600.00 | 0.22 | 16008.83 | whole |
| 1 | Bg2/+;MHC/+ | LP | ND | none | 25 | 1810.00 | 0.18 | 9848.84 | whole |
| 1 | Bg2/+;MHC/+ | LP | ND | none | 25 | 3070.00 | 0.19 | 16192.52 | whole |
| 1 | Bg2/+;MHC/+ | LP | ND | none | 25 | 1700.00 | 0.10 | 17260.87 | whole |
| 1 | Bg2/+;MHC/+ | LP | ND | none | 25 | 4470.00 | 0.15 | 30212.33 | whole |
| 4 | Bg2/+;MHC/+ | LP | ND | none | 25 | 11520.00 | 0.38 | 30312.40 | whole |
| 4 | Bg2/+;MHC/+ | LP | ND | none | 25 | 7640.00 | 0.32 | 23791.34 | whole |
| 4 | Bg2/+;MHC/+ | LP | ND | none | 25 | 15080.00 | 0.42 | 35591.26 | whole |
| 4 | Bg2/+;MHC/+ | LP | ND | none | 25 | 5840.00 | 0.27 | 21333.13 | whole |
| 4 | Bg2/+;MHC/+ | LP | ND | none | 25 | 8440.00 | 0.28 | 30564.07 | whole |
| 5 | Bg2/+;MHC/+ | LP | ND | none | 25 | 4600.00 | 0.20 | 23498.52 | whole |
| 5 | Bg2/+;MHC/+ | LP | ND | none | 25 | 10400.00 | 0.39 | 26820.88 | whole |

| Rep. | Genotype | Stage | Sex | Treatmen | °C | ß-gal ΔmOD/min/ ml | Protein mg/ml | Specific Activity ΔmOD/min/ mg | Whole/ dissected |
|---|---|---|---|---|---|---|---|---|---|
| 5 | Bg2/+;MHC/+ | LP | ND | none | 25 | 6440.00 | 0.30 | 21349.69 | whole |
| 5 | Bg2/+;MHC/+ | LP | ND | none | 25 | 2440.00 | 0.23 | 10407.32 | whole |
| 5 | Bg2/+;MHC/+ | LP | ND | none | 25 | 11200.00 | 0.31 | 36245.49 | whole |
| 1 | Bg2/DJ694 | 0-24h | F | none | 25 | 1335.00 | 0.20 | 6682.53 | thorax |
| 1 | Bg2/DJ694 | 0-24h | F | none | 25 | 1800.00 | 0.17 | 10731.12 | thorax |
| 1 | Bg2/DJ694 | 0-24h | F | none | 25 | 2220.00 | 0.24 | 9184.07 | thorax |
| 1 | Bg2/DJ694 | 0-24h | F | none | 25 | 780.00 | 0.19 | 4204.19 | thorax |
| 1 | Bg2/DJ694 | 0-24h | F | none | 25 | 750.00 | 0.21 | 3579.68 | thorax |
| 1 | Bg2/DJ694 | 10-11d | F | none | 25 | 11060.00 | 0.21 | 52932.97 | thorax |
| 1 | Bg2/DJ694 | 10-11d | F | none | 25 | 8830.00 | 0.23 | 37615.21 | thorax |
| 1 | Bg2/DJ694 | 10-11d | F | none | 25 | 9920.00 | 0.22 | 44126.50 | thorax |
| 1 | Bg2/DJ694 | 10-11d | F | none | 25 | 8040.00 | 0.22 | 36198.93 | thorax |
| 1 | Bg2/DJ694 | 10-11d | F | none | 25 | 13360.00 | 0.61 | 21736.38 | thorax |
| 1 | Bg2/DJ694 | 3-5d | F | none | 25 | 10990.00 | 0.13 | 81588.72 | thorax |
| 1 | Bg2/DJ694 | 3-5d | F | none | 25 | 9080.00 | 0.18 | 50982.59 | thorax |
| 1 | Bg2/DJ694 | 3-5d | F | none | 25 | 7615.00 | 0.38 | 20055.31 | thorax |
| 1 | Bg2/DJ694 | 3-5d | F | none | 25 | 11910.00 | 0.17 | 68843.93 | thorax |
| 1 | Bg2/DJ694 | 3-5d | F | none | 25 | 6915.00 | 0.18 | 37725.04 | thorax |
| 1 | Bg2/DJ694 | 0-24h | M | none | 25 | 450.00 | 0.14 | 3287.54 | thorax |
| 1 | Bg2/DJ694 | 0-24h | M | none | 25 | 615.00 | 0.16 | 3807.04 | thorax |
| 1 | Bg2/DJ694 | 0-24h | M | none | 25 | 495.00 | 0.14 | 3458.38 | thorax |
| 1 | Bg2/DJ694 | 0-24h | M | none | 25 | 1485.00 | 0.16 | 9329.18 | thorax |
| 1 | Bg2/DJ694 | 0-24h | M | none | 25 | 510.00 | 0.16 | 3244.13 | thorax |
| 1 | Bg2/DJ694 | 10-11d | M | none | 25 | 7400.00 | 0.15 | 48732.95 | thorax |
| 1 | Bg2/DJ694 | 10-11d | M | none | 25 | 4320.00 | 0.15 | 27999.46 | thorax |
| 1 | Bg2/DJ694 | 10-11d | M | none | 25 | 4070.00 | 0.15 | 26275.21 | thorax |
| 1 | Bg2/DJ694 | 10-11d | M | none | 25 | 3480.00 | 0.15 | 22760.87 | thorax |
| 1 | Bg2/DJ694 | 10-11d | M | none | 25 | 3370.00 | 0.05 | 64929.64 | thorax |
| 1 | Bg2/DJ694 | 3-5d | M | none | 25 | 2070.00 | 0.11 | 18784.03 | thorax |
| 1 | Bg2/DJ694 | 3-5d | M | none | 25 | 1930.00 | 0.12 | 15539.45 | thorax |
| 1 | Bg2/DJ694 | 3-5d | M | none | 25 | 2860.00 | 0.10 | 27875.24 | thorax |
| 1 | Bg2/DJ694 | 3-5d | M | none | 25 | 3260.00 | 0.13 | 26080.00 | thorax |
| 1 | Bg2/DJ694 | 3-5d | M | none | 25 | 1840.00 | 0.25 | 7269.85 | thorax |
| 1 | Bg2/DJ694 | LP | ND | none | 25 | 310.00 | 0.15 | 2020.09 | whole |
| 1 | Bg2/DJ694 | LP | ND | none | 25 | 130.00 | 0.10 | 1331.87 | whole |
| 1 | Bg2/DJ694 | LP | ND | none | 25 | 160.00 | 0.15 | 1038.95 | whole |
| 1 | Bg2/DJ694 | LP | ND | none | 25 | 65.11 | 0.10 | 639.01 | whole |
| 1 | Bg2/DJ694 | LP | ND | none | 25 | 150.00 | 0.12 | 1201.82 | whole |
| 6 | DJ694,Bg2 | 10-11d | F | -RU486 | 25 | 55070.00 | 0.11 | 493257.60 | thorax |
| 6 | DJ694,Bg2 | 10-11d | F | -RU486 | 25 | 68995.00 | 0.11 | 609050.71 | thorax |

| Rep. | Genotype | Stage | Sex | Treatmen | °C | ß-gal ΔmOD/min/ml | Protein mg/ml | Specific Activity ΔmOD/min/mg | Whole/ dissected |
|---|---|---|---|---|---|---|---|---|---|
| 6 | DJ694,Bg2 | 10-11d | F | -RU486 | 25 | 62145.00 | 0.12 | 508184.89 | thorax |
| 6 | DJ694,Bg2 | 10-11d | F | -RU486 | 25 | 38695.00 | 0.07 | 537112.98 | thorax |
| 6 | DJ694,Bg2 | 10-11d | F | -RU486 | 25 | 43265.00 | 0.11 | 395126.71 | thorax |
| 7 | DJ694,Bg2 | 10-12d | F | -RU486 | 25 | 65946.67 | 0.15 | 430002.82 | thorax |
| 7 | DJ694,Bg2 | 10-12d | F | -RU486 | 25 | 33760.00 | 0.12 | 279479.84 | thorax |
| 7 | DJ694,Bg2 | 10-12d | F | -RU486 | 25 | 36460.00 | 0.16 | 226707.10 | thorax |
| 7 | DJ694,Bg2 | 10-12d | F | -RU486 | 25 | 49060.00 | 0.11 | 438373.16 | thorax |
| 7 | DJ694,Bg2 | 10-12d | F | -RU486 | 25 | 38860.00 | 0.11 | 342877.58 | thorax |
| 6 | DJ694,Bg2 | 10-11d | F | +RU486 | 25 | 68645.00 | 0.11 | 619389.60 | thorax |
| 6 | DJ694,Bg2 | 10-11d | F | +RU486 | 25 | 30015.00 | 0.08 | 355954.59 | thorax |
| 6 | DJ694,Bg2 | 10-11d | F | +RU486 | 25 | 62490.00 | 0.09 | 681531.56 | thorax |
| 6 | DJ694,Bg2 | 10-11d | F | +RU486 | 25 | 30440.00 | 0.10 | 316783.47 | thorax |
| 6 | DJ694,Bg2 | 10-11d | F | +RU486 | 25 | 39010.00 | 0.08 | 517942.55 | thorax |
| 7 | DJ694,Bg2 | 10-12d | F | +RU486 | 25 | 33100.00 | 0.09 | 358788.70 | thorax |
| 7 | DJ694,Bg2 | 10-12d | F | +RU486 | 25 | 38000.00 | 0.11 | 340990.44 | thorax |
| 7 | DJ694,Bg2 | 10-12d | F | +RU486 | 25 | 19560.00 | 0.08 | 256069.21 | thorax |
| 7 | DJ694,Bg2 | 10-12d | F | +RU486 | 25 | 45220.00 | 0.12 | 369282.09 | thorax |
| 7 | DJ694,Bg2 | 10-12d | F | +RU486 | 25 | 32680.00 | 0.15 | 214413.30 | thorax |
| 5 | DJ694,Bg2 | 0-1d | F | none | 25 | 7560.00 | 0.12 | 65471.07 | thorax |
| 5 | DJ694,Bg2 | 0-1d | F | none | 25 | 19480.00 | 0.13 | 150932.99 | thorax |
| 5 | DJ694,Bg2 | 0-1d | F | none | 25 | 7040.00 | 0.12 | 56958.55 | thorax |
| 5 | DJ694,Bg2 | 0-1d | F | none | 25 | 14760.00 | 0.25 | 59139.45 | thorax |
| 5 | DJ694,Bg2 | 0-1d | F | none | 25 | 24280.00 | 0.14 | 172571.79 | thorax |
| 5 | DJ694,Bg2 | 0-1d | F | none | 25 | 54760.00 | 0.07 | 754377.14 | abdomen |
| 5 | DJ694,Bg2 | 0-1d | F | none | 25 | 58160.00 | 0.13 | 455076.49 | abdomen |
| 5 | DJ694,Bg2 | 0-1d | F | none | 25 | 50040.00 | 0.14 | 358519.52 | abdomen |
| 5 | DJ694,Bg2 | 0-1d | F | none | 25 | 69720.00 | 0.17 | 409820.36 | abdomen |
| 5 | DJ694,Bg2 | 0-1d | F | none | 25 | 83640.00 | 0.13 | 664649.27 | abdomen |
| 4 | DJ694,Bg2 | 1-2d | F | none | 25 | 17240.00 | 0.10 | 179162.65 | thorax |
| 4 | DJ694,Bg2 | 1-2d | F | none | 25 | 18840.00 | 0.11 | 164636.66 | thorax |
| 4 | DJ694,Bg2 | 1-2d | F | none | 25 | 33280.00 | 0.08 | 391860.28 | thorax |
| 4 | DJ694,Bg2 | 1-2d | F | none | 25 | 33360.00 | 0.06 | 524009.69 | thorax |
| 4 | DJ694,Bg2 | 1-2d | F | none | 25 | 37800.00 | 0.12 | 304504.50 | thorax |
| 4 | DJ694,Bg2 | 1-2d | F | none | 25 | 56880.00 | 0.22 | 253383.73 | abdomen |
| 4 | DJ694,Bg2 | 1-2d | F | none | 25 | 56960.00 | 0.15 | 391028.32 | abdomen |
| 4 | DJ694,Bg2 | 1-2d | F | none | 25 | 54560.00 | 0.10 | 526294.15 | abdomen |
| 4 | DJ694,Bg2 | 1-2d | F | none | 25 | 49480.00 | 0.10 | 508589.51 | abdomen |
| 4 | DJ694,Bg2 | 1-2d | F | none | 25 | 46000.00 | 0.18 | 258866.12 | abdomen |
| 4 | DJ694,Bg2 | 9-12d | F | none | 25 | 63960.00 | 0.19 | 336662.52 | thorax |
| 4 | DJ694,Bg2 | 9-12d | F | none | 25 | 64200.00 | 0.16 | 407995.01 | thorax |

| Rep. | Genotype | Stage | Sex | Treatmen | °C | ß-gal ΔmOD/min/ ml | Protein mg/ml | Specific Activity ΔmOD/min/ mg | Whole/ dissected |
|---|---|---|---|---|---|---|---|---|---|
| 4 | DJ694,Bg2 | 9-12d | F | none | 25 | 67360.00 | 0.12 | 551641.68 | thorax |
| 4 | DJ694,Bg2 | 9-12d | F | none | 25 | 67560.00 | 0.16 | 413850.16 | thorax |
| 4 | DJ694,Bg2 | 9-12d | F | none | 25 | 85000.00 | 0.14 | 603829.46 | thorax |
| 4 | DJ694,Bg2 | 9-12d | F | none | 25 | 62200.00 | 0.11 | 583419.45 | abdomen |
| 4 | DJ694,Bg2 | 9-12d | F | none | 25 | 20120.00 | 0.19 | 103700.61 | abdomen |
| 4 | DJ694,Bg2 | 9-12d | F | none | 25 | 54720.00 | 0.10 | 548636.85 | abdomen |
| 4 | DJ694,Bg2 | 9-12d | F | none | 25 | 65400.00 | 0.14 | 457849.96 | abdomen |
| 4 | DJ694,Bg2 | 9-12d | F | none | 25 | 100640.00 | 0.22 | 466969.60 | abdomen |
| 6 | DJ694,Bg2 | 10-11d | M | -RU486 | 25 | 20690.00 | 0.06 | 355329.84 | thorax |
| 6 | DJ694,Bg2 | 10-11d | M | -RU486 | 25 | 21275.00 | 0.07 | 308455.93 | thorax |
| 6 | DJ694,Bg2 | 10-11d | M | -RU486 | 25 | 29455.00 | 0.06 | 455433.96 | thorax |
| 6 | DJ694,Bg2 | 10-11d | M | -RU486 | 25 | 31425.00 | 0.06 | 498514.77 | thorax |
| 6 | DJ694,Bg2 | 10-11d | M | -RU486 | 25 | 24505.00 | 0.07 | 352151.26 | thorax |
| 7 | DJ694,Bg2 | 10-12d | M | -RU486 | 25 | 15260.00 | 0.11 | 135352.35 | thorax |
| 7 | DJ694,Bg2 | 10-12d | M | -RU486 | 25 | 20400.00 | 0.13 | 160090.71 | thorax |
| 7 | DJ694,Bg2 | 10-12d | M | -RU486 | 25 | 26160.00 | 0.10 | 249881.27 | thorax |
| 7 | DJ694,Bg2 | 10-12d | M | -RU486 | 25 | 28380.00 | 0.14 | 198542.44 | thorax |
| 7 | DJ694,Bg2 | 10-12d | M | -RU486 | 25 | 34880.00 | 0.11 | 325084.68 | thorax |
| 6 | DJ694,Bg2 | 10-11d | M | +RU486 | 25 | 22705.00 | 0.07 | 306454.78 | thorax |
| 6 | DJ694,Bg2 | 10-11d | M | +RU486 | 25 | 26040.00 | 0.09 | 298315.22 | thorax |
| 6 | DJ694,Bg2 | 10-11d | M | +RU486 | 25 | 23710.00 | 0.05 | 478706.86 | thorax |
| 6 | DJ694,Bg2 | 10-11d | M | +RU486 | 25 | 25130.00 | 0.07 | 336859.20 | thorax |
| 6 | DJ694,Bg2 | 10-11d | M | +RU486 | 25 | 22190.00 | 0.04 | 597357.25 | thorax |
| 7 | DJ694,Bg2 | 10-12d | M | +RU486 | 25 | 16260.00 | 0.08 | 194200.06 | thorax |
| 7 | DJ694,Bg2 | 10-12d | M | +RU486 | 25 | 19600.00 | 0.10 | 198444.12 | thorax |
| 7 | DJ694,Bg2 | 10-12d | M | +RU486 | 25 | 21200.00 | 0.10 | 217512.52 | thorax |
| 7 | DJ694,Bg2 | 10-12d | M | +RU486 | 25 | 18080.00 | 0.10 | 173683.19 | thorax |
| 7 | DJ694,Bg2 | 10-12d | M | +RU486 | 25 | 19680.00 | 0.07 | 288503.33 | thorax |
| 5 | DJ694,Bg2 | 0-1d | M | none | 25 | 21800.00 | 0.18 | 123660.41 | thorax |
| 5 | DJ694,Bg2 | 0-1d | M | none | 25 | 15360.00 | 0.11 | 143467.23 | thorax |
| 5 | DJ694,Bg2 | 0-1d | M | none | 25 | 9200.00 | 0.12 | 76249.94 | thorax |
| 5 | DJ694,Bg2 | 0-1d | M | none | 25 | 8560.00 | 0.06 | 148262.52 | thorax |
| 5 | DJ694,Bg2 | 0-1d | M | none | 25 | 6040.00 | 0.14 | 43058.38 | thorax |
| 5 | DJ694,Bg2 | 0-1d | M | none | 25 | 43400.00 | 0.03 | 1540807.96 | abdomen |
| 5 | DJ694,Bg2 | 0-1d | M | none | 25 | 60560.00 | 0.07 | 883754.93 | abdomen |
| 5 | DJ694,Bg2 | 0-1d | M | none | 25 | 66160.00 | 0.12 | 571571.14 | abdomen |
| 5 | DJ694,Bg2 | 0-1d | M | none | 25 | 50160.00 | 0.14 | 353697.39 | abdomen |
| 5 | DJ694,Bg2 | 0-1d | M | none | 25 | 47560.00 | 0.05 | 961439.55 | abdomen |
| 4 | DJ694,Bg2 | 1-2d | M | none | 25 | 9120.00 | 0.12 | 74022.52 | thorax |
| 4 | DJ694,Bg2 | 1-2d | M | none | 25 | 11080.00 | 0.10 | 114199.89 | thorax |

| Rep. | Genotype | Stage | Sex | Treatmen | °C | ß-gal ΔmOD/min/ ml | Protein mg/ml | Specific Activity ΔmOD/min/ mg | Whole/ dissected |
|---|---|---|---|---|---|---|---|---|---|
| 4 | DJ694,Bg2 | 1-2d | M | none | 25 | 11120.00 | 0.12 | 92757.07 | thorax |
| 4 | DJ694,Bg2 | 1-2d | M | none | 25 | 13400.00 | 0.08 | 168036.00 | thorax |
| 4 | DJ694,Bg2 | 1-2d | M | none | 25 | 20160.00 | 0.06 | 350309.10 | thorax |
| 4 | DJ694,Bg2 | 1-2d | M | none | 25 | 38920.00 | 0.05 | 745124.89 | abdomen |
| 4 | DJ694,Bg2 | 1-2d | M | none | 25 | 17720.00 | 0.02 | 913186.85 | abdomen |
| 4 | DJ694,Bg2 | 1-2d | M | none | 25 | 16800.00 | 0.01 | 1663200.00 | abdomen |
| 4 | DJ694,Bg2 | 1-2d | M | none | 25 | 11680.00 | 0.04 | 332880.00 | abdomen |
| 4 | DJ694,Bg2 | 1-2d | M | none | 25 | 30880.00 | 0.04 | 764280.00 | abdomen |
| 4 | DJ694,Bg2 | 9-12d | M | none | 25 | 16920.00 | 0.11 | 154591.11 | thorax |
| 4 | DJ694,Bg2 | 9-12d | M | none | 25 | 24880.00 | 0.11 | 236269.76 | thorax |
| 4 | DJ694,Bg2 | 9-12d | M | none | 25 | 21400.00 | 0.10 | 214093.45 | thorax |
| 4 | DJ694,Bg2 | 9-12d | M | none | 25 | 9280.00 | 0.11 | 86336.97 | thorax |
| 4 | DJ694,Bg2 | 9-12d | M | none | 25 | 25880.00 | 0.10 | 259479.56 | thorax |
| 4 | DJ694,Bg2 | 9-12d | M | none | 25 | 50760.00 | 0.13 | 404843.03 | abdomen |
| 4 | DJ694,Bg2 | 9-12d | M | none | 25 | 72040.00 | 0.14 | 525198.54 | abdomen |
| 4 | DJ694,Bg2 | 9-12d | M | none | 25 | 30840.00 | 0.09 | 336850.73 | abdomen |
| 4 | DJ694,Bg2 | 9-12d | M | none | 25 | 32440.00 | 0.08 | 392190.18 | abdomen |
| 4 | DJ694,Bg2 | 9-12d | M | none | 25 | 55080.00 | 0.06 | 856966.25 | abdomen |
| 4 | DJ694,Bg2 | EP | ND | none | 25 | 31360.00 | 0.64 | 49342.37 | whole |
| 4 | DJ694,Bg2 | EP | ND | none | 25 | 27720.00 | 0.38 | 73603.89 | whole |
| 4 | DJ694,Bg2 | EP | ND | none | 25 | 16640.00 | 0.49 | 34186.33 | whole |
| 4 | DJ694,Bg2 | EP | ND | none | 25 | 18520.00 | 0.45 | 41502.90 | whole |
| 4 | DJ694,Bg2 | EP | ND | none | 25 | 38120.00 | 0.48 | 79244.05 | whole |
| 5 | DJ694,Bg2 | EP | ND | none | 25 | 3600.00 | 0.25 | 14406.98 | whole |
| 5 | DJ694,Bg2 | EP | ND | none | 25 | 3840.00 | 0.28 | 13678.91 | whole |
| 5 | DJ694,Bg2 | EP | ND | none | 25 | 4320.00 | 0.30 | 14346.35 | whole |
| 5 | DJ694,Bg2 | EP | ND | none | 25 | 5840.00 | 0.23 | 25620.24 | whole |
| 5 | DJ694,Bg2 | EP | ND | none | 25 | 8240.00 | 0.20 | 41260.00 | whole |
| 4 | DJ694,Bg2 | LP | ND | none | 25 | 85400.00 | 0.40 | 214096.61 | whole |
| 4 | DJ694,Bg2 | LP | ND | none | 25 | 25000.00 | 0.34 | 73393.07 | whole |
| 4 | DJ694,Bg2 | LP | ND | none | 25 | 36320.00 | 0.38 | 96443.81 | whole |
| 4 | DJ694,Bg2 | LP | ND | none | 25 | 46440.00 | 0.36 | 128194.81 | whole |
| 4 | DJ694,Bg2 | LP | ND | none | 25 | 46360.00 | 0.25 | 187893.15 | whole |
| 5 | DJ694,Bg2 | LP | ND | none | 25 | 30200.00 | 0.43 | 70968.69 | whole |
| 5 | DJ694,Bg2 | LP | ND | none | 25 | 27880.00 | 0.25 | 110705.19 | whole |
| 5 | DJ694,Bg2 | LP | ND | none | 25 | 23240.00 | 0.17 | 134730.29 | whole |
| 5 | DJ694,Bg2 | LP | ND | none | 25 | 63320.00 | 0.24 | 260437.64 | whole |
| 5 | DJ694,Bg2 | LP | ND | none | 25 | 29680.00 | 0.22 | 133600.26 | whole |
| 5 | DJ694,Bg2/+ | 0-1d | F | none | 25 | 32240.00 | 0.12 | 280224.90 | abdomen |
| 5 | DJ694,Bg2/+ | 0-1d | F | none | 25 | 25600.00 | 0.08 | 330346.47 | abdomen |

| Rep. | Genotype | Stage | Sex | Treatmen | °C | ß-gal ΔmOD/min/ ml | Protein mg/ml | Specific Activity ΔmOD/min/ mg | Whole/ dissected |
|---|---|---|---|---|---|---|---|---|---|
| 5 | DJ694,Bg2/+ | 0-1d | F | none | 25 | 27560.00 | 0.13 | 219251.01 | abdomen |
| 5 | DJ694,Bg2/+ | 0-1d | F | none | 25 | 12880.00 | 0.11 | 119832.70 | abdomen |
| 5 | DJ694,Bg2/+ | 0-1d | F | none | 25 | 20480.00 | 0.04 | 541278.81 | abdomen |
| 4 | DJ694,Bg2/+ | 1-2d | F | none | 25 | 26440.00 | 0.10 | 257021.40 | abdomen |
| 4 | DJ694,Bg2/+ | 1-2d | F | none | 25 | 26840.00 | 0.17 | 157154.99 | abdomen |
| 4 | DJ694,Bg2/+ | 1-2d | F | none | 25 | 23080.00 | 0.09 | 260350.70 | abdomen |
| 4 | DJ694,Bg2/+ | 1-2d | F | none | 25 | 17760.00 | 0.14 | 129734.21 | abdomen |
| 4 | DJ694,Bg2/+ | 1-2d | F | none | 25 | 26960.00 | 0.10 | 275233.43 | abdomen |
| 4 | DJ694,Bg2/+ | 9-12d | F | none | 25 | 9400.00 | 0.09 | 105825.06 | abdomen |
| 4 | DJ694,Bg2/+ | 9-12d | F | none | 25 | 7360.00 | 0.10 | 71599.83 | abdomen |
| 4 | DJ694,Bg2/+ | 9-12d | F | none | 25 | 46120.00 | 0.25 | 187758.19 | abdomen |
| 4 | DJ694,Bg2/+ | 9-12d | F | none | 25 | 6920.00 | 0.10 | 66403.02 | abdomen |
| 4 | DJ694,Bg2/+ | 9-12d | F | none | 25 | 11440.00 | 0.12 | 96179.96 | abdomen |
| 5 | DJ694,Bg2/+ | 0-1d | F | none | 25 | 1840.00 | 0.12 | 15502.05 | thorax |
| 5 | DJ694,Bg2/+ | 0-1d | F | none | 25 | 3920.00 | 0.13 | 29727.01 | thorax |
| 5 | DJ694,Bg2/+ | 0-1d | F | none | 25 | 1720.00 | 0.12 | 14372.27 | thorax |
| 5 | DJ694,Bg2/+ | 0-1d | F | none | 25 | 4040.00 | 0.13 | 32139.84 | thorax |
| 5 | DJ694,Bg2/+ | 0-1d | F | none | 25 | 2000.00 | 0.06 | 31786.19 | thorax |
| 4 | DJ694,Bg2/+ | 1-2d | F | none | 25 | 1240.00 | 0.09 | 13560.70 | thorax |
| 4 | DJ694,Bg2/+ | 1-2d | F | none | 25 | 5880.00 | 0.10 | 57085.32 | thorax |
| 4 | DJ694,Bg2/+ | 1-2d | F | none | 25 | 11840.00 | 0.10 | 114947.30 | thorax |
| 4 | DJ694,Bg2/+ | 1-2d | F | none | 25 | 4920.00 | 0.09 | 53186.90 | thorax |
| 4 | DJ694,Bg2/+ | 1-2d | F | none | 25 | 10040.00 | 0.09 | 111253.25 | thorax |
| 4 | DJ694,Bg2/+ | 9-12d | F | none | 25 | 9160.00 | 0.12 | 73827.83 | thorax |
| 4 | DJ694,Bg2/+ | 9-12d | F | none | 25 | 16120.00 | 0.14 | 113459.05 | thorax |
| 4 | DJ694,Bg2/+ | 9-12d | F | none | 25 | 21280.00 | 0.12 | 174896.79 | thorax |
| 4 | DJ694,Bg2/+ | 9-12d | F | none | 25 | 14400.00 | 0.15 | 94191.01 | thorax |
| 4 | DJ694,Bg2/+ | 9-12d | F | none | 25 | 15000.00 | 0.14 | 104611.87 | thorax |
| 5 | DJ694,Bg2/+ | 0-1d | M | none | 25 | 10320.00 | 0.03 | 299363.90 | abdomen |
| 5 | DJ694,Bg2/+ | 0-1d | M | none | 25 | 15560.00 | 0.04 | 375122.16 | abdomen |
| 5 | DJ694,Bg2/+ | 0-1d | M | none | 25 | 15920.00 | 0.06 | 263584.97 | abdomen |
| 5 | DJ694,Bg2/+ | 0-1d | M | none | 25 | 10320.00 | 0.05 | 201211.80 | abdomen |
| 5 | DJ694,Bg2/+ | 0-1d | M | none | 25 | 18160.00 | 0.10 | 177520.22 | abdomen |
| 4 | DJ694,Bg2/+ | 1-2d | M | none | 25 | 15400.00 | 0.05 | 280555.93 | abdomen |
| 4 | DJ694,Bg2/+ | 1-2d | M | none | 25 | 17280.00 | 0.13 | 128727.45 | abdomen |
| 4 | DJ694,Bg2/+ | 1-2d | M | none | 25 | 12040.00 | 0.06 | 186781.36 | abdomen |
| 4 | DJ694,Bg2/+ | 1-2d | M | none | 25 | 13320.00 | 0.03 | 396125.22 | abdomen |
| 4 | DJ694,Bg2/+ | 1-2d | M | none | 25 | 15840.00 | 0.08 | 207269.84 | abdomen |
| 4 | DJ694,Bg2/+ | 9-12d | M | none | 25 | 17320.00 | 0.10 | 164990.10 | abdomen |
| 4 | DJ694,Bg2/+ | 9-12d | M | none | 25 | 39560.00 | 0.12 | 342007.40 | abdomen |

| Rep. | Genotype | Stage | Sex | Treatmen | °C | ß-gal ΔmOD/min/ ml | Protein mg/ml | Specific Activity ΔmOD/min/ mg | Whole/ dissected |
|---|---|---|---|---|---|---|---|---|---|
| 4 | DJ694,Bg2/+ | 9-12d | M | none | 25 | 25760.00 | 0.08 | 313915.74 | abdomen |
| 4 | DJ694,Bg2/+ | 9-12d | M | none | 25 | 22600.00 | 0.09 | 248031.62 | abdomen |
| 4 | DJ694,Bg2/+ | 9-12d | M | none | 25 | 31400.00 | 0.11 | 284900.59 | abdomen |
| 5 | DJ694,Bg2/+ | 0-1d | M | none | 25 | 920.00 | 0.09 | 10131.36 | thorax |
| 5 | DJ694,Bg2/+ | 0-1d | M | none | 25 | 1520.00 | 0.10 | 15385.42 | thorax |
| 5 | DJ694,Bg2/+ | 0-1d | M | none | 25 | 280.00 | 0.07 | 4154.01 | thorax |
| 5 | DJ694,Bg2/+ | 0-1d | M | none | 25 | 600.00 | 0.02 | 27098.73 | thorax |
| 5 | DJ694,Bg2/+ | 0-1d | M | none | 25 | 560.00 | 0.07 | 7684.92 | thorax |
| 4 | DJ694,Bg2/+ | 1-2d | M | none | 25 | 2880.00 | 0.07 | 41671.38 | thorax |
| 4 | DJ694,Bg2/+ | 1-2d | M | none | 25 | 120.00 | 0.08 | 1465.71 | thorax |
| 4 | DJ694,Bg2/+ | 1-2d | M | none | 25 | 1120.00 | 0.09 | 12284.08 | thorax |
| 4 | DJ694,Bg2/+ | 1-2d | M | none | 25 | 2720.00 | 0.07 | 40686.44 | thorax |
| 4 | DJ694,Bg2/+ | 1-2d | M | none | 25 | 2280.00 | 0.05 | 48323.15 | thorax |
| 4 | DJ694,Bg2/+ | 9-12d | M | none | 25 | 6720.00 | 0.07 | 100296.55 | thorax |
| 4 | DJ694,Bg2/+ | 9-12d | M | none | 25 | 4640.00 | 0.12 | 39480.93 | thorax |
| 4 | DJ694,Bg2/+ | 9-12d | M | none | 25 | 7080.00 | 0.23 | 30346.64 | thorax |
| 4 | DJ694,Bg2/+ | 9-12d | M | none | 25 | 4920.00 | 0.12 | 40041.63 | thorax |
| 4 | DJ694,Bg2/+ | 9-12d | M | none | 25 | 6640.00 | 0.14 | 48562.62 | thorax |
| 4 | DJ694,Bg2/+ | EP | ND | none | 25 | 7560.00 | 0.64 | 11832.87 | whole |
| 4 | DJ694,Bg2/+ | EP | ND | none | 25 | 8160.00 | 0.47 | 17406.85 | whole |
| 4 | DJ694,Bg2/+ | EP | ND | none | 25 | 8520.00 | 0.50 | 16898.94 | whole |
| 4 | DJ694,Bg2/+ | EP | ND | none | 25 | 5760.00 | 0.32 | 17841.41 | whole |
| 4 | DJ694,Bg2/+ | EP | ND | none | 25 | 4880.00 | 0.41 | 11915.11 | whole |
| 5 | DJ694,Bg2/+ | EP | ND | none | 25 | 3240.00 | 0.35 | 9161.70 | whole |
| 5 | DJ694,Bg2/+ | EP | ND | none | 25 | 3960.00 | 0.35 | 11411.55 | whole |
| 5 | DJ694,Bg2/+ | EP | ND | none | 25 | 9560.00 | 0.57 | 16752.66 | whole |
| 5 | DJ694,Bg2/+ | EP | ND | none | 25 | 2640.00 | 0.31 | 8521.88 | whole |
| 5 | DJ694,Bg2/+ | EP | ND | none | 25 | 5960.00 | 0.51 | 11600.75 | whole |
| 4 | DJ694,Bg2/+ | LP | ND | none | 25 | 9080.00 | 0.30 | 30479.68 | whole |
| 4 | DJ694,Bg2/+ | LP | ND | none | 25 | 9720.00 | 0.18 | 55450.92 | whole |
| 4 | DJ694,Bg2/+ | LP | ND | none | 25 | 8840.00 | 0.18 | 49308.48 | whole |
| 4 | DJ694,Bg2/+ | LP | ND | none | 25 | 13120.00 | 0.23 | 56029.24 | whole |
| 4 | DJ694,Bg2/+ | LP | ND | none | 25 | 9400.00 | 0.30 | 31581.99 | whole |
| 5 | DJ694,Bg2/+ | LP | ND | none | 25 | 15960.00 | 0.34 | 47139.83 | whole |
| 5 | DJ694,Bg2/+ | LP | ND | none | 25 | 16240.00 | 0.30 | 54146.62 | whole |
| 5 | DJ694,Bg2/+ | LP | ND | none | 25 | 6200.00 | 0.17 | 36145.61 | whole |
| 5 | DJ694,Bg2/+ | LP | ND | none | 25 | 10000.00 | 0.35 | 28295.73 | whole |
| 5 | DJ694,Bg2/+ | LP | ND | none | 25 | 21080.00 | 0.32 | 66426.04 | whole |
| 5 | DJ694,Bg2/Bg2 | 0-1d | F | none | 25 | 64800.00 | 0.10 | 675055.18 | abdomen |
| 5 | DJ694,Bg2/Bg2 | 0-1d | F | none | 25 | 59640.00 | 0.11 | 559252.35 | abdomen |

| Rep. | Genotype | Stage | Sex | Treatmen | °C | ß-gal ΔmOD/min/ml | Protein mg/ml | Specific Activity ΔmOD/min/mg | Whole/ dissected |
|---|---|---|---|---|---|---|---|---|---|
| 5 | DJ694,Bg2/Bg2 | 0-1d | F | none | 25 | 35960.00 | 0.05 | 675290.95 | abdomen |
| 5 | DJ694,Bg2/Bg2 | 0-1d | F | none | 25 | 39840.00 | 0.10 | 404983.25 | abdomen |
| 5 | DJ694,Bg2/Bg2 | 0-1d | F | none | 25 | 39480.00 | 0.07 | 554585.20 | abdomen |
| 4 | DJ694,Bg2/Bg2 | 1-2d | F | none | 25 | 33040.00 | 0.07 | 491290.43 | abdomen |
| 4 | DJ694,Bg2/Bg2 | 1-2d | F | none | 25 | 46840.00 | 0.09 | 532362.78 | abdomen |
| 4 | DJ694,Bg2/Bg2 | 1-2d | F | none | 25 | 37240.00 | 0.04 | 953040.00 | abdomen |
| 4 | DJ694,Bg2/Bg2 | 1-2d | F | none | 25 | 46520.00 | 0.09 | 538486.89 | abdomen |
| 4 | DJ694,Bg2/Bg2 | 1-2d | F | none | 25 | 39920.00 | 0.06 | 631004.37 | abdomen |
| 4 | DJ694,Bg2/Bg2 | 9-12d | F | none | 25 | 28520.00 | 0.14 | 206606.55 | abdomen |
| 4 | DJ694,Bg2/Bg2 | 9-12d | F | none | 25 | 76160.00 | 0.10 | 736213.33 | abdomen |
| 4 | DJ694,Bg2/Bg2 | 9-12d | F | none | 25 | 77160.00 | 0.16 | 492748.60 | abdomen |
| 4 | DJ694,Bg2/Bg2 | 9-12d | F | none | 25 | 75680.00 | 0.12 | 649374.08 | abdomen |
| 4 | DJ694,Bg2/Bg2 | 9-12d | F | none | 25 | 77960.00 | 0.19 | 419756.43 | abdomen |
| 5 | DJ694,Bg2/Bg2 | 0-1d | F | none | 25 | 11600.00 | 0.19 | 62238.80 | thorax |
| 5 | DJ694,Bg2/Bg2 | 0-1d | F | none | 25 | 5360.00 | 0.14 | 37352.50 | thorax |
| 5 | DJ694,Bg2/Bg2 | 0-1d | F | none | 25 | 5160.00 | 0.08 | 63595.44 | thorax |
| 5 | DJ694,Bg2/Bg2 | 0-1d | F | none | 25 | 7800.00 | 0.13 | 59466.67 | thorax |
| 5 | DJ694,Bg2/Bg2 | 0-1d | F | none | 25 | 10200.00 | 0.14 | 71640.94 | thorax |
| 4 | DJ694,Bg2/Bg2 | 1-2d | F | none | 25 | 14600.00 | 0.14 | 105019.50 | thorax |
| 4 | DJ694,Bg2/Bg2 | 1-2d | F | none | 25 | 13080.00 | 0.09 | 142422.46 | thorax |
| 4 | DJ694,Bg2/Bg2 | 1-2d | F | none | 25 | 13400.00 | 0.13 | 100821.60 | thorax |
| 4 | DJ694,Bg2/Bg2 | 1-2d | F | none | 25 | 16320.00 | 0.06 | 255816.00 | thorax |
| 4 | DJ694,Bg2/Bg2 | 1-2d | F | none | 25 | 13440.00 | 0.12 | 112986.10 | thorax |
| 4 | DJ694,Bg2/Bg2 | 9-12d | F | none | 25 | 41680.00 | 0.12 | 351385.02 | thorax |
| 4 | DJ694,Bg2/Bg2 | 9-12d | F | none | 25 | 32640.00 | 0.14 | 236640.00 | thorax |
| 4 | DJ694,Bg2/Bg2 | 9-12d | F | none | 25 | 43720.00 | 0.13 | 336115.84 | thorax |
| 4 | DJ694,Bg2/Bg2 | 9-12d | F | none | 25 | 37720.00 | 0.11 | 338888.31 | thorax |
| 4 | DJ694,Bg2/Bg2 | 9-12d | F | none | 25 | 21760.00 | 0.10 | 214880.00 | thorax |
| 5 | DJ694,Bg2/Bg2 | 0-1d | M | none | 25 | 46600.00 | 0.11 | 412066.42 | abdomen |
| 5 | DJ694,Bg2/Bg2 | 0-1d | M | none | 25 | 42880.00 | 0.03 | 1264428.43 | abdomen |
| 5 | DJ694,Bg2/Bg2 | 0-1d | M | none | 25 | 29800.00 | 0.07 | 423611.16 | abdomen |
| 5 | DJ694,Bg2/Bg2 | 0-1d | M | none | 25 | 17880.00 | 0.03 | 650977.96 | abdomen |
| 5 | DJ694,Bg2/Bg2 | 0-1d | M | none | 25 | 47080.00 | 0.11 | 442638.84 | abdomen |
| 4 | DJ694,Bg2/Bg2 | 1-2d | M | none | 25 | 29840.00 | 0.07 | 446354.19 | abdomen |
| 4 | DJ694,Bg2/Bg2 | 1-2d | M | none | 25 | 35880.00 | 0.05 | 780234.45 | abdomen |
| 4 | DJ694,Bg2/Bg2 | 1-2d | M | none | 25 | 37040.00 | 0.07 | 555157.29 | abdomen |
| 4 | DJ694,Bg2/Bg2 | 1-2d | M | none | 25 | 23920.00 | 0.04 | 534047.72 | abdomen |
| 4 | DJ694,Bg2/Bg2 | 1-2d | M | none | 25 | 30000.00 | 0.06 | 542596.15 | abdomen |
| 4 | DJ694,Bg2/Bg2 | 9-12d | M | none | 25 | 60160.00 | 0.09 | 640309.22 | abdomen |
| 4 | DJ694,Bg2/Bg2 | 9-12d | M | none | 25 | 63680.00 | 0.13 | 491215.08 | abdomen |

| Rep. | Genotype | Stage | Sex | Treatmen | °C | ß-gal ΔmOD/min/ml | Protein mg/ml | Specific Activity ΔmOD/min/mg | Whole/ dissected |
|---|---|---|---|---|---|---|---|---|---|
| 4 | DJ694,Bg2/Bg2 | 9-12d | M | none | 25 | 65760.00 | 0.11 | 577226.67 | abdomen |
| 4 | DJ694,Bg2/Bg2 | 9-12d | M | none | 25 | 36120.00 | 0.09 | 389416.09 | abdomen |
| 4 | DJ694,Bg2/Bg2 | 9-12d | M | none | 25 | 52600.00 | 0.10 | 518307.96 | abdomen |
| 5 | DJ694,Bg2/Bg2 | 0-1d | M | none | 25 | 920.00 | 0.04 | 21177.81 | thorax |
| 5 | DJ694,Bg2/Bg2 | 0-1d | M | none | 25 | 6040.00 | 0.10 | 62195.44 | thorax |
| 5 | DJ694,Bg2/Bg2 | 0-1d | M | none | 25 | 2560.00 | 0.08 | 33458.17 | thorax |
| 5 | DJ694,Bg2/Bg2 | 0-1d | M | none | 25 | 2640.00 | 0.11 | 25118.72 | thorax |
| 5 | DJ694,Bg2/Bg2 | 0-1d | M | none | 25 | 1160.00 | 0.04 | 29044.77 | thorax |
| 4 | DJ694,Bg2/Bg2 | 1-2d | M | none | 25 | 2680.00 | 0.11 | 24650.76 | thorax |
| 4 | DJ694,Bg2/Bg2 | 1-2d | M | none | 25 | 2640.00 | 0.09 | 28704.28 | thorax |
| 4 | DJ694,Bg2/Bg2 | 1-2d | M | none | 25 | 3560.00 | 0.06 | 60600.54 | thorax |
| 4 | DJ694,Bg2/Bg2 | 1-2d | M | none | 25 | 3160.00 | 0.04 | 72266.99 | thorax |
| 4 | DJ694,Bg2/Bg2 | 1-2d | M | none | 25 | 3760.00 | 0.10 | 39183.16 | thorax |
| 4 | DJ694,Bg2/Bg2 | 9-12d | M | none | 25 | 10960.00 | 0.12 | 89277.72 | thorax |
| 4 | DJ694,Bg2/Bg2 | 9-12d | M | none | 25 | 15000.00 | 0.12 | 128467.29 | thorax |
| 4 | DJ694,Bg2/Bg2 | 9-12d | M | none | 25 | 15240.00 | 0.11 | 134806.33 | thorax |
| 4 | DJ694,Bg2/Bg2 | 9-12d | M | none | 25 | 5280.00 | 0.09 | 56393.85 | thorax |
| 4 | DJ694,Bg2/Bg2 | 9-12d | M | none | 25 | 11920.00 | 0.13 | 91871.22 | thorax |
| 4 | DJ694,Bg2/Bg2 | EP | ND | none | 25 | 12720.00 | 0.46 | 27825.43 | whole |
| 4 | DJ694,Bg2/Bg2 | EP | ND | none | 25 | 10120.00 | 0.42 | 24247.15 | whole |
| 4 | DJ694,Bg2/Bg2 | EP | ND | none | 25 | 15080.00 | 0.52 | 28849.95 | whole |
| 4 | DJ694,Bg2/Bg2 | EP | ND | none | 25 | 8560.00 | 0.40 | 21405.30 | whole |
| 4 | DJ694,Bg2/Bg2 | EP | ND | none | 25 | 10200.00 | 0.43 | 23809.83 | whole |
| 5 | DJ694,Bg2/Bg2 | EP | ND | none | 25 | 23720.00 | 0.58 | 40716.82 | whole |
| 5 | DJ694,Bg2/Bg2 | EP | ND | none | 25 | 9600.00 | 0.34 | 28610.33 | whole |
| 5 | DJ694,Bg2/Bg2 | EP | ND | none | 25 | 3520.00 | 0.19 | 18211.31 | whole |
| 5 | DJ694,Bg2/Bg2 | EP | ND | none | 25 | 24200.00 | 0.63 | 38382.93 | whole |
| 5 | DJ694,Bg2/Bg2 | EP | ND | none | 25 | 17680.00 | 0.67 | 26216.01 | whole |
| 4 | DJ694,Bg2/Bg2 | LP | ND | none | 25 | 32560.00 | 0.36 | 91556.78 | whole |
| 4 | DJ694,Bg2/Bg2 | LP | ND | none | 25 | 24720.00 | 0.34 | 71788.02 | whole |
| 4 | DJ694,Bg2/Bg2 | LP | ND | none | 25 | 35080.00 | 0.35 | 99309.87 | whole |
| 4 | DJ694,Bg2/Bg2 | LP | ND | none | 25 | 39520.00 | 0.37 | 107789.62 | whole |
| 4 | DJ694,Bg2/Bg2 | LP | ND | none | 25 | 47000.00 | 0.47 | 101024.53 | whole |
| 5 | DJ694,Bg2/Bg2 | LP | ND | none | 25 | 40920.00 | 0.39 | 103886.59 | whole |
| 5 | DJ694,Bg2/Bg2 | LP | ND | none | 25 | 11520.00 | 0.14 | 83321.64 | whole |
| 5 | DJ694,Bg2/Bg2 | LP | ND | none | 25 | 18720.00 | 0.24 | 77341.20 | whole |
| 5 | DJ694,Bg2/Bg2 | LP | ND | none | 25 | 14480.00 | 0.20 | 71503.62 | whole |
| 5 | DJ694,Bg2/Bg2 | LP | ND | none | 25 | 12960.00 | 0.24 | 54549.04 | whole |
| 5 | DJ694,Bg2/DJ694 | 0-1d | F | none | 25 | 31400.00 | 0.08 | 382372.70 | abdomen |
| 5 | DJ694,Bg2/DJ694 | 0-1d | F | none | 25 | 29560.00 | 0.07 | 398000.30 | abdomen |

| Rep. | Genotype | Stage | Sex | Treatmen | °C | ß-gal ΔmOD/min/ ml | Protein mg/ml | Specific Activity ΔmOD/min/ mg | Whole/ dissected |
|---|---|---|---|---|---|---|---|---|---|
| 5 | DJ694,Bg2/DJ694 | 0-1d | F | none | 25 | 26720.00 | 0.07 | 361810.09 | abdomen |
| 5 | DJ694,Bg2/DJ694 | 0-1d | F | none | 25 | 33720.00 | 0.13 | 255712.99 | abdomen |
| 5 | DJ694,Bg2/DJ694 | 0-1d | F | none | 25 | 30600.00 | 0.10 | 311056.41 | abdomen |
| 4 | DJ694,Bg2/DJ694 | 1-2d | F | none | 25 | 24080.00 | 0.15 | 165459.29 | abdomen |
| 4 | DJ694,Bg2/DJ694 | 1-2d | F | none | 25 | 21040.00 | 0.05 | 395762.40 | abdomen |
| 4 | DJ694,Bg2/DJ694 | 1-2d | F | none | 25 | 28120.00 | 0.09 | 322522.68 | abdomen |
| 4 | DJ694,Bg2/DJ694 | 1-2d | F | none | 25 | 28880.00 | 0.17 | 166380.64 | abdomen |
| 4 | DJ694,Bg2/DJ694 | 1-2d | F | none | 25 | 22720.00 | 0.12 | 182633.85 | abdomen |
| 4 | DJ694,Bg2/DJ694 | 9-12d | F | none | 25 | 36200.00 | 0.21 | 170208.72 | abdomen |
| 4 | DJ694,Bg2/DJ694 | 9-12d | F | none | 25 | 32920.00 | 0.10 | 317891.34 | abdomen |
| 4 | DJ694,Bg2/DJ694 | 9-12d | F | none | 25 | 41920.00 | 0.14 | 294146.16 | abdomen |
| 4 | DJ694,Bg2/DJ694 | 9-12d | F | none | 25 | 17240.00 | 0.20 | 84939.44 | abdomen |
| 4 | DJ694,Bg2/DJ694 | 9-12d | F | none | 25 | 32920.00 | 0.24 | 137189.12 | abdomen |
| 5 | DJ694,Bg2/DJ694 | 0-1d | F | none | 25 | 9680.00 | 0.10 | 97017.53 | thorax |
| 5 | DJ694,Bg2/DJ694 | 0-1d | F | none | 25 | 14760.00 | 0.16 | 91668.72 | thorax |
| 5 | DJ694,Bg2/DJ694 | 0-1d | F | none | 25 | 13920.00 | 0.12 | 116451.49 | thorax |
| 5 | DJ694,Bg2/DJ694 | 0-1d | F | none | 25 | 9520.00 | 0.22 | 43325.71 | thorax |
| 5 | DJ694,Bg2/DJ694 | 0-1d | F | none | 25 | 11840.00 | 0.11 | 105480.95 | thorax |
| 4 | DJ694,Bg2/DJ694 | 1-2d | F | none | 25 | 34520.00 | 0.13 | 273398.40 | thorax |
| 4 | DJ694,Bg2/DJ694 | 1-2d | F | none | 25 | 28080.00 | 0.12 | 241455.91 | thorax |
| 4 | DJ694,Bg2/DJ694 | 1-2d | F | none | 25 | 20280.00 | 0.15 | 138715.20 | thorax |
| 4 | DJ694,Bg2/DJ694 | 1-2d | F | none | 25 | 21240.00 | 0.09 | 226039.26 | thorax |
| 4 | DJ694,Bg2/DJ694 | 1-2d | F | none | 25 | 27640.00 | 0.12 | 226047.13 | thorax |
| 4 | DJ694,Bg2/DJ694 | 9-12d | F | none | 25 | 54400.00 | 0.18 | 305092.78 | thorax |
| 4 | DJ694,Bg2/DJ694 | 9-12d | F | none | 25 | 34160.00 | 0.11 | 302456.27 | thorax |
| 4 | DJ694,Bg2/DJ694 | 9-12d | F | none | 25 | 50280.00 | 0.14 | 356354.15 | thorax |
| 4 | DJ694,Bg2/DJ694 | 9-12d | F | none | 25 | 37800.00 | 0.18 | 209811.75 | thorax |
| 4 | DJ694,Bg2/DJ694 | 9-12d | F | none | 25 | 63680.00 | 0.13 | 501774.31 | thorax |
| 5 | DJ694,Bg2/DJ694 | 0-1d | M | none | 25 | 28680.00 | 0.10 | 288254.20 | abdomen |
| 5 | DJ694,Bg2/DJ694 | 0-1d | M | none | 25 | 18240.00 | 0.04 | 445755.62 | abdomen |
| 5 | DJ694,Bg2/DJ694 | 0-1d | M | none | 25 | 21480.00 | 0.04 | 598755.00 | abdomen |
| 5 | DJ694,Bg2/DJ694 | 0-1d | M | none | 25 | 22240.00 | 0.04 | 523777.69 | abdomen |
| 5 | DJ694,Bg2/DJ694 | 0-1d | M | none | 25 | 17600.00 | 0.04 | 470388.01 | abdomen |
| 4 | DJ694,Bg2/DJ694 | 1-2d | M | none | 25 | 24280.00 | 0.08 | 311214.17 | abdomen |
| 4 | DJ694,Bg2/DJ694 | 1-2d | M | none | 25 | 20520.00 | 0.07 | 298054.98 | abdomen |
| 4 | DJ694,Bg2/DJ694 | 1-2d | M | none | 25 | 22200.00 | 0.08 | 276544.37 | abdomen |
| 4 | DJ694,Bg2/DJ694 | 1-2d | M | none | 25 | 17680.00 | 0.07 | 237543.43 | abdomen |
| 4 | DJ694,Bg2/DJ694 | 9-12d | M | none | 25 | 14560.00 | 0.13 | 111748.61 | abdomen |
| 4 | DJ694,Bg2/DJ694 | 9-12d | M | none | 25 | 16240.00 | 0.10 | 170278.44 | abdomen |
| 4 | DJ694,Bg2/DJ694 | 9-12d | M | none | 25 | 25240.00 | 0.10 | 248708.99 | abdomen |

| Rep. | Genotype | Stage | Sex | Treatmen | °C | ß-gal ΔmOD/min/ ml | Protein mg/ml | Specific Activity ΔmOD/min/ mg | Whole/ dissected |
|---|---|---|---|---|---|---|---|---|---|
| 4 | DJ694,Bg2/DJ694 | 9-12d | M | none | 25 | 15400.00 | 0.11 | 143712.42 | abdomen |
| 4 | DJ694,Bg2/DJ694 | 9-12d | M | none | 25 | 25280.00 | 0.10 | 262065.52 | abdomen |
| 5 | DJ694,Bg2/DJ694 | 0-1d | M | none | 25 | 9600.00 | 0.09 | 104909.04 | thorax |
| 5 | DJ694,Bg2/DJ694 | 0-1d | M | none | 25 | 7680.00 | 0.10 | 75696.80 | thorax |
| 5 | DJ694,Bg2/DJ694 | 0-1d | M | none | 25 | 8440.00 | 0.07 | 112575.40 | thorax |
| 5 | DJ694,Bg2/DJ694 | 0-1d | M | none | 25 | 4320.00 | 0.07 | 64627.92 | thorax |
| 5 | DJ694,Bg2/DJ694 | 0-1d | M | none | 25 | 6600.00 | 0.05 | 133420.96 | thorax |
| 4 | DJ694,Bg2/DJ694 | 1-2d | M | none | 25 | 12160.00 | 0.06 | 192614.40 | thorax |
| 4 | DJ694,Bg2/DJ694 | 1-2d | M | none | 25 | 15920.00 | 0.20 | 81539.88 | thorax |
| 4 | DJ694,Bg2/DJ694 | 1-2d | M | none | 25 | 18480.00 | 0.06 | 328708.09 | thorax |
| 4 | DJ694,Bg2/DJ694 | 1-2d | M | none | 25 | 18080.00 | 0.07 | 252382.04 | thorax |
| 4 | DJ694,Bg2/DJ694 | 9-12d | M | none | 25 | 14760.00 | 0.12 | 123188.20 | thorax |
| 4 | DJ694,Bg2/DJ694 | 9-12d | M | none | 25 | 19880.00 | 0.09 | 210613.09 | thorax |
| 4 | DJ694,Bg2/DJ694 | 9-12d | M | none | 25 | 27160.00 | 0.11 | 241879.73 | thorax |
| 4 | DJ694,Bg2/DJ694 | 9-12d | M | none | 25 | 25480.00 | 0.11 | 226477.90 | thorax |
| 4 | DJ694,Bg2/DJ694 | 9-12d | M | none | 25 | 22120.00 | 0.12 | 184279.71 | thorax |
| 4 | DJ694,Bg2/DJ694 | EP | ND | none | 25 | 6720.00 | 0.33 | 20262.92 | whole |
| 4 | DJ694,Bg2/DJ694 | EP | ND | none | 25 | 7800.00 | 0.32 | 24451.11 | whole |
| 4 | DJ694,Bg2/DJ694 | EP | ND | none | 25 | 11360.00 | 0.43 | 26162.03 | whole |
| 4 | DJ694,Bg2/DJ694 | EP | ND | none | 25 | 12240.00 | 0.49 | 25044.68 | whole |
| 4 | DJ694,Bg2/DJ694 | EP | ND | none | 25 | 5040.00 | 0.26 | 19446.88 | whole |
| 5 | DJ694,Bg2/DJ694 | EP | ND | none | 25 | 4040.00 | 0.26 | 15390.85 | whole |
| 5 | DJ694,Bg2/DJ694 | EP | ND | none | 25 | 8720.00 | 0.42 | 20987.93 | whole |
| 5 | DJ694,Bg2/DJ694 | EP | ND | none | 25 | 7760.00 | 0.38 | 20168.80 | whole |
| 5 | DJ694,Bg2/DJ694 | EP | ND | none | 25 | 4720.00 | 0.30 | 15788.35 | whole |
| 5 | DJ694,Bg2/DJ694 | EP | ND | none | 25 | 6960.00 | 0.30 | 23045.27 | whole |
| 4 | DJ694,Bg2/DJ694 | LP | ND | none | 25 | 21280.00 | 0.31 | 68974.66 | whole |
| 4 | DJ694,Bg2/DJ694 | LP | ND | none | 25 | 28480.00 | 0.31 | 92991.89 | whole |
| 4 | DJ694,Bg2/DJ694 | LP | ND | none | 25 | 43240.00 | 0.30 | 142357.64 | whole |
| 4 | DJ694,Bg2/DJ694 | LP | ND | none | 25 | 25640.00 | 0.26 | 98332.34 | whole |
| 4 | DJ694,Bg2/DJ694 | LP | ND | none | 25 | 45280.00 | 0.38 | 118771.35 | whole |
| 5 | DJ694,Bg2/DJ694 | LP | ND | none | 25 | 15720.00 | 0.12 | 130282.84 | whole |
| 5 | DJ694,Bg2/DJ694 | LP | ND | none | 25 | 35720.00 | 0.21 | 169139.91 | whole |
| 5 | DJ694,Bg2/DJ694 | LP | ND | none | 25 | 30960.00 | 0.20 | 156041.41 | whole |
| 5 | DJ694,Bg2/DJ694 | LP | ND | none | 25 | 29560.00 | 0.21 | 140936.64 | whole |
| 5 | DJ694,Bg2/DJ694 | LP | ND | none | 25 | 33160.00 | 0.23 | 142906.68 | whole |

Table 3 Longevity of genotypes with multiple copies of Q72 and muscle driver transgenes (related to Figures 3, S4). EDTP data are not used in this report.

| Replicate | Genotype | Sex | Treatment | n | Mean | Median | Maximum |
|---|---|---|---|---|---|---|---|
| 1 | MHC/+ | male | -RU486 | 126 | 59.66 | 61.00 | 69.75 |
| 2 | MHC/+ | male | -RU486 | 87 | 57.06 | 59.00 | 68.00 |
| 3 | MHC/+ | male | -RU486 | 113 | 58.78 | 58.00 | 73.00 |
| 4 | MHC/+ | male | -RU486 | 122 | 68.52 | 69.00 | 92.75 |
| 1 | Q72;;MHC | male | -RU486 | 136 | 48.07 | 51.00 | 62.00 |
| 2 | Q72;;MHC | male | -RU486 | 137 | 49.75 | 52.00 | 62.25 |
| 3 | Q72;;MHC | male | -RU486 | 87 | 51.41 | 54.00 | 58.25 |
| 4 | Q72;;MHC | male | -RU486 | 87 | 54.09 | 54.00 | 61.25 |
| 5 | Q72;;MHC | male | -RU486 | 47 | 48.69 | 51.00 | 56.00 |
| 6 | Q72;;MHC | male | -RU486 | 56 | 49.23 | 52.00 | 59.00 |
| 7 | Q72;;MHC | male | -RU486 | 153 | 45.64 | 48.00 | 58.25 |
| 8 | Q72;;MHC | male | -RU486 | 153 | 47.55 | 50.00 | 58.75 |
| 1 | Q72;;MHC/+ | male | -RU486 | 144 | 55.25 | 56.00 | 71.50 |
| 2 | Q72;;MHC/+ | male | -RU486 | 144 | 55.94 | 57.50 | 69.75 |
| 3 | Q72;;MHC/+ | male | -RU486 | 127 | 56.52 | 58.00 | 73.25 |
| 4 | Q72;;MHC/+ | male | -RU486 | 78 | 56.58 | 59.00 | 73.00 |
| 5 | Q72;;MHC/+ | male | -RU486 | 83 | 54.39 | 56.00 | 66.00 |
| 6 | Q72;;MHC/+ | male | -RU486 | 92 | 55.05 | 56.00 | 66.67 |
| 7 | Q72;;MHC/+ | male | -RU486 | 95 | 53.69 | 52.00 | 70.00 |
| 8 | Q72;;MHC/+ | male | -RU486 | 125 | 53.91 | 53.00 | 68.00 |
| 1 | Q72/+;;MHC | male | -RU486 | 141 | 46.37 | 47.00 | 55.00 |
| 2 | Q72/+;;MHC | male | -RU486 | 85 | 50.11 | 52.00 | 60.75 |
| 3 | Q72/+;;MHC | male | -RU486 | 63 | 49.68 | 51.00 | 59.00 |
| 4 | Q72/+;;MHC | male | -RU486 | 101 | 51.59 | 52.00 | 60.00 |
| 5 | Q72/+;;MHC | male | -RU486 | 73 | 46.93 | 49.00 | 56.33 |
| 6 | Q72/+;;MHC | male | -RU486 | 75 | 47.67 | 49.00 | 59.00 |
| 7 | Q72/+;;MHC | male | -RU486 | 122 | 44.99 | 46.50 | 53.75 |
| 8 | Q72/+;;MHC | male | -RU486 | 142 | 46.78 | 48.00 | 56.75 |
| 1 | Q72/+;;MHC/+ | male | -RU486 | 118 | 57.43 | 56.00 | 69.50 |
| 2 | Q72/+;;MHC/+ | male | -RU486 | 121 | 59.25 | 59.00 | 73.00 |
| 3 | Q72/+;;MHC/+ | male | -RU486 | 82 | 64.40 | 68.00 | 76.75 |
| 4 | Q72/+;;MHC/+ | male | -RU486 | 98 | 68.98 | 68.00 | 80.00 |
| 5 | Q72/+;;MHC/+ | male | -RU486 | 57 | 57.30 | 58.00 | 65.67 |
| 6 | Q72/+;;MHC/+ | male | -RU486 | 70 | 57.16 | 59.00 | 66.00 |
| 7 | Q72/+;;MHC/+ | male | -RU486 | 112 | 57.13 | 59.00 | 69.00 |
| 8 | Q72/+;;MHC/+ | male | -RU486 | 141 | 58.67 | 60.00 | 74.00 |
| 1 | Q72/+;;MHC/UAS-EDTP | male | -RU486 | 144 | 54.64 | 56.00 | 63.00 |
| 2 | Q72/+;;MHC/UAS-EDTP | male | -RU486 | 98 | 53.30 | 56.00 | 67.75 |
| 3 | Q72/+;;MHC/UAS-EDTP | male | -RU486 | 99 | 59.70 | 61.00 | 72.00 |

| Replicate | Genotype | Sex | Treatment | n | Mean | Median | Maximum |
|---|---|---|---|---|---|---|---|
| 4 | Q72/+;;MHC/UAS-EDTP | male | -RU486 | 95 | 67.52 | 70.00 | 81.00 |
| 1 | UAS-EDTP/+ | male | -RU486 | 152 | 50.14 | 47.00 | 73.25 |
| 2 | UAS-EDTP/+ | male | -RU486 | 117 | 55.46 | 54.00 | 68.75 |
| 3 | UAS-EDTP/+ | male | -RU486 | 69 | 72.76 | 75.00 | 95.25 |
| 4 | UAS-EDTP/+ | male | -RU486 | 99 | 70.29 | 73.00 | 90.50 |
| 1 | UAS-EDTP/MHC | male | -RU486 | 150 | 53.09 | 54.00 | 68.50 |
| 2 | UAS-EDTP/MHC | male | -RU486 | 129 | 52.35 | 52.00 | 65.75 |
| 3 | UAS-EDTP/MHC | male | -RU486 | 74 | 50.50 | 54.00 | 63.67 |
| 4 | UAS-EDTP/MHC | male | -RU486 | 91 | 57.59 | 56.00 | 81.00 |
| 1 | MHC/+ | male | +RU486 | 138 | 57.03 | 54.00 | 69.00 |
| 2 | MHC/+ | male | +RU486 | 89 | 55.10 | 59.00 | 67.00 |
| 3 | MHC/+ | male | +RU486 | 100 | 64.19 | 65.00 | 79.00 |
| 4 | MHC/+ | male | +RU486 | 93 | 64.11 | 63.00 | 85.25 |
| 1 | Q72;;MHC | male | +RU486 | 129 | 22.41 | 23.00 | 28.00 |
| 2 | Q72;;MHC | male | +RU486 | 142 | 23.62 | 24.00 | 30.25 |
| 3 | Q72;;MHC | male | +RU486 | 97 | 22.82 | 23.00 | 26.50 |
| 4 | Q72;;MHC | male | +RU486 | 89 | 24.68 | 24.00 | 27.50 |
| 5 | Q72;;MHC | male | +RU486 | 71 | 24.76 | 26.00 | 26.67 |
| 6 | Q72;;MHC | male | +RU486 | 95 | 24.02 | 24.00 | 26.67 |
| 7 | Q72;;MHC | male | +RU486 | 123 | 22.71 | 22.00 | 28.00 |
| 8 | Q72;;MHC | male | +RU486 | 155 | 22.41 | 25.00 | 27.25 |
| 1 | Q72;;MHC/+ | male | +RU486 | 138 | 27.90 | 28.00 | 32.25 |
| 2 | Q72;;MHC/+ | male | +RU486 | 135 | 28.12 | 28.00 | 32.00 |
| 3 | Q72;;MHC/+ | male | +RU486 | 114 | 29.51 | 30.00 | 34.00 |
| 4 | Q72;;MHC/+ | male | +RU486 | 90 | 28.85 | 28.00 | 36.00 |
| 5 | Q72;;MHC/+ | male | +RU486 | 90 | 27.70 | 28.00 | 32.25 |
| 6 | Q72;;MHC/+ | male | +RU486 | 119 | 28.22 | 28.00 | 32.00 |
| 7 | Q72;;MHC/+ | male | +RU486 | 119 | 26.21 | 27.00 | 30.75 |
| 8 | Q72;;MHC/+ | male | +RU486 | 116 | 26.80 | 27.00 | 32.00 |
| 1 | Q72/+;;MHC | male | +RU486 | 161 | 20.73 | 19.00 | 29.50 |
| 2 | Q72/+;;MHC | male | +RU486 | 94 | 24.13 | 24.00 | 29.50 |
| 3 | Q72/+;;MHC | male | +RU486 | 60 | 24.60 | 26.00 | 27.33 |
| 4 | Q72/+;;MHC | male | +RU486 | 110 | 24.98 | 24.00 | 27.25 |
| 5 | Q72/+;;MHC | male | +RU486 | 96 | 26.95 | 26.00 | 30.75 |
| 6 | Q72/+;;MHC | male | +RU486 | 93 | 26.43 | 26.00 | 30.25 |
| 7 | Q72/+;;MHC | male | +RU486 | 118 | 21.44 | 20.00 | 27.50 |
| 8 | Q72/+;;MHC | male | +RU486 | 125 | 17.26 | 18.00 | 25.50 |
| 1 | Q72/+;;MHC/+ | male | +RU486 | 132 | 21.44 | 21.00 | 26.00 |
| 2 | Q72/+;;MHC/+ | male | +RU486 | 132 | 22.37 | 21.00 | 25.50 |
| 3 | Q72/+;;MHC/+ | male | +RU486 | 71 | 24.69 | 26.00 | 28.50 |
| 4 | Q72/+;;MHC/+ | male | +RU486 | 99 | 24.94 | 24.00 | 30.00 |

| Replicate | Genotype | Sex | Treatment | n | Mean | Median | Maximum |
|---|---|---|---|---|---|---|---|
| 5 | Q72/+;;MHC/+ | male | +RU486 | 82 | 24.68 | 26.00 | 28.75 |
| 6 | Q72/+;;MHC/+ | male | +RU486 | 68 | 25.13 | 24.00 | 30.00 |
| 7 | Q72/+;;MHC/+ | male | +RU486 | 106 | 23.96 | 22.00 | 28.00 |
| 8 | Q72/+;;MHC/+ | male | +RU486 | 128 | 23.78 | 25.00 | 27.50 |
| 1 | Q72/+;;MHC/UAS-EDTP | male | +RU486 | 134 | 24.68 | 26.00 | 28.00 |
| 2 | Q72/+;;MHC/UAS-EDTP | male | +RU486 | 92 | 24.84 | 24.00 | 30.25 |
| 3 | Q72/+;;MHC/UAS-EDTP | male | +RU486 | 51 | 26.48 | 26.00 | 30.00 |
| 4 | Q72/+;;MHC/UAS-EDTP | male | +RU486 | 97 | 26.42 | 26.00 | 32.00 |
| 1 | UAS-EDTP/+ | male | +RU486 | 158 | 47.43 | 48.00 | 66.25 |
| 2 | UAS-EDTP/+ | male | +RU486 | 125 | 58.72 | 52.00 | 69.25 |
| 3 | UAS-EDTP/+ | male | +RU486 | 94 | 71.59 | 75.00 | 90.00 |
| 4 | UAS-EDTP/+ | male | +RU486 | 105 | 65.62 | 61.00 | 80.50 |
| 1 | UAS-EDTP/MHC | male | +RU486 | 142 | 53.69 | 54.00 | 72.50 |
| 2 | UAS-EDTP/MHC | male | +RU486 | 128 | 55.31 | 54.00 | 71.00 |
| 3 | UAS-EDTP/MHC | male | +RU486 | 62 | 48.35 | 49.00 | 64.67 |
| 4 | UAS-EDTP/MHC | male | +RU486 | 62 | 59.39 | 59.00 | 73.00 |
| 1 | DJ694 | male | none | 146 | 42.36 | 44.00 | 50.75 |
| 2 | DJ694 | male | none | 114 | 42.19 | 45.00 | 47.50 |
| 3 | DJ694 | male | none | 114 | 43.11 | 47.00 | 50.50 |
| 4 | DJ694 | male | none | 98 | 46.19 | 47.00 | 52.00 |
| 7 | DJ694 | male | none | 91 | 40.45 | 41.00 | 48.00 |
| 8 | DJ694 | male | none | 125 | 42.88 | 46.00 | 49.25 |
| 1 | DJ694;UAS-EDTP/+ | male | none | 124 | 47.92 | 52.50 | 59.00 |
| 2 | DJ694;UAS-EDTP/+ | male | none | 139 | 49.00 | 49.00 | 66.00 |
| 3 | DJ694;UAS-EDTP/+ | male | none | 144 | 57.14 | 57.00 | 77.75 |
| 4 | DJ694;UAS-EDTP/+ | male | none | 159 | 49.86 | 45.00 | 65.25 |
| 7 | DJ694;UAS-EDTP/+ | male | none | 152 | 44.09 | 48.00 | 56.00 |
| 8 | DJ694;UAS-EDTP/+ | male | none | 153 | 47.87 | 46.00 | 66.00 |
| 1 | DJ694/+ | male | none | 112 | 51.25 | 54.00 | 57.25 |
| 2 | DJ694/+ | male | none | 103 | 54.74 | 52.00 | 66.00 |
| 3 | DJ694/+ | male | none | 51 | 63.58 | 63.00 | 85.50 |
| 4 | DJ694/+ | male | none | 81 | 61.28 | 59.00 | 89.75 |
| 1 | Q72;DJ694 | male | none | 137 | 28.47 | 30.00 | 34.00 |
| 2 | Q72;DJ694 | male | none | 113 | 27.93 | 28.00 | 32.00 |
| 3 | Q72;DJ694 | male | none | 98 | 30.67 | 33.00 | 34.50 |
| 4 | Q72;DJ694 | male | none | 77 | 27.49 | 31.00 | 35.00 |
| 5 | Q72;DJ694 | male | none | 109 | 30.24 | 31.00 | 34.75 |
| 6 | Q72;DJ694 | male | none | 154 | 29.59 | 30.00 | 35.80 |
| 7 | Q72;DJ694 | male | none | 135 | 26.03 | 27.00 | 34.00 |
| 8 | Q72;DJ694 | male | none | 89 | 27.59 | 29.00 | 33.00 |
| 1 | Q72;DJ694/+ | male | none | 133 | 59.58 | 61.00 | 68.00 |

| Replicate | Genotype | Sex | Treatment | n | Mean | Median | Maximum |
|---|---|---|---|---|---|---|---|
| 2 | Q72;DJ694/+ | male | none | 149 | 54.52 | 56.00 | 66.00 |
| 3 | Q72;DJ694/+ | male | none | 132 | 53.48 | 54.00 | 69.00 |
| 4 | Q72;DJ694/+ | male | none | 85 | 52.39 | 54.00 | 67.00 |
| 5 | Q72;DJ694/+ | male | none | 65 | 61.84 | 61.00 | 71.67 |
| 6 | Q72;DJ694/+ | male | none | 83 | 58.83 | 59.00 | 68.33 |
| 7 | Q72;DJ694/+ | male | none | 146 | 51.64 | 52.00 | 62.50 |
| 8 | Q72;DJ694/+ | male | none | 164 | 50.20 | 50.00 | 61.75 |
| 1 | Q72/+;DJ694 | male | none | 93 | 27.45 | 28.00 | 30.75 |
| 2 | Q72/+;DJ694 | male | none | 104 | 25.91 | 26.00 | 29.50 |
| 3 | Q72/+;DJ694 | male | none | 75 | 26.82 | 26.00 | 32.75 |
| 4 | Q72/+;DJ694 | male | none | 96 | 26.91 | 26.00 | 28.75 |
| 5 | Q72/+;DJ694 | male | none | 99 | 27.90 | 28.00 | 30.75 |
| 6 | Q72/+;DJ694 | male | none | 20 | 27.65 | 28.00 | 31.00 |
| 6 | Q72/+;DJ694 | male | none | 111 | 28.06 | 28.00 | 33.25 |
| 7 | Q72/+;DJ694 | male | none | 45 | 23.14 | 24.00 | 27.00 |
| 8 | Q72/+;DJ694 | male | none | 25 | 24.36 | 25.00 | 29.00 |
| 1 | Q72/+;DJ694;UAS-EDTP/+ | male | none | 96 | 32.19 | 35.00 | 40.00 |
| 2 | Q72/+;DJ694;UAS-EDTP/+ | male | none | 96 | 32.26 | 32.00 | 37.25 |
| 3 | Q72/+;DJ694;UAS-EDTP/+ | male | none | 84 | 34.77 | 35.00 | 40.25 |
| 4 | Q72/+;DJ694;UAS-EDTP/+ | male | none | 150 | 36.28 | 38.00 | 42.50 |
| 7 | Q72/+;DJ694;UAS-EDTP/+ | male | none | 58 | 32.53 | 34.00 | 37.50 |
| 8 | Q72/+;DJ694;UAS-EDTP/+ | male | none | 60 | 35.49 | 35.00 | 40.00 |
| 1 | Q72/+;DJ694/+ | male | none | 122 | 45.67 | 47.00 | 52.75 |
| 2 | Q72/+;DJ694/+ | male | none | 177 | 38.91 | 38.00 | 47.75 |
| 3 | Q72/+;DJ694/+ | male | none | 89 | 45.87 | 51.00 | 57.00 |
| 4 | Q72/+;DJ694/+ | male | none | 78 | 51.31 | 52.00 | 60.25 |
| 5 | Q72/+;DJ694/+ | male | none | 102 | 44.59 | 47.00 | 51.25 |
| 6 | Q72/+;DJ694/+ | male | none | 80 | 47.94 | 49.00 | 56.50 |
| 7 | Q72/+;DJ694/+ | male | none | 173 | 39.98 | 48.00 | 54.25 |
| 8 | Q72/+;DJ694/+ | male | none | 179 | 46.17 | 46.00 | 53.50 |
| 1 | UAS-EDTP/+ | male | none | 134 | 49.05 | 44.00 | 69.00 |
| 2 | UAS-EDTP/+ | male | none | 137 | 58.57 | 66.00 | 71.50 |
| 3 | UAS-EDTP/+ | male | none | 70 | 70.67 | 73.50 | 94.75 |
| 4 | UAS-EDTP/+ | male | none | 101 | 73.25 | 84.00 | 96.75 |
| 1 | MHC/+ | female | -RU486 | 127 | 54.16 | 51.00 | 72.50 |
| 2 | MHC/+ | female | -RU486 | 97 | 60.93 | 66.00 | 74.00 |
| 3 | MHC/+ | female | -RU486 | 103 | 66.77 | 68.00 | 85.50 |
| 4 | MHC/+ | female | -RU486 | 149 | 75.34 | 75.00 | 88.50 |
| 1 | Q72;;MHC | female | -RU486 | 139 | 54.58 | 56.00 | 67.25 |
| 2 | Q72;;MHC | female | -RU486 | 138 | 55.97 | 59.00 | 66.50 |
| 3 | Q72;;MHC | female | -RU486 | 109 | 55.41 | 56.00 | 66.50 |

| Replicate | Genotype | Sex | Treatment | n | Mean | Median | Maximum |
|---|---|---|---|---|---|---|---|
| 4 | Q72;;MHC | female | -RU486 | 85 | 47.48 | 38.00 | 57.25 |
| 5 | Q72;;MHC | female | -RU486 | 70 | 58.34 | 61.00 | 65.33 |
| 6 | Q72;;MHC | female | -RU486 | 61 | 60.16 | 61.00 | 66.00 |
| 7 | Q72;;MHC | female | -RU486 | 123 | 56.95 | 59.00 | 66.75 |
| 8 | Q72;;MHC | female | -RU486 | 174 | 56.56 | 60.00 | 67.75 |
| 1 | Q72;;MHC/+ | female | -RU486 | 136 | 47.11 | 44.00 | 70.00 |
| 2 | Q72;;MHC/+ | female | -RU486 | 124 | 51.15 | 52.00 | 71.25 |
| 3 | Q72;;MHC/+ | female | -RU486 | 116 | 58.51 | 61.00 | 74.25 |
| 4 | Q72;;MHC/+ | female | -RU486 | 121 | 57.19 | 56.00 | 74.00 |
| 5 | Q72;;MHC/+ | female | -RU486 | 118 | 49.68 | 47.00 | 71.75 |
| 6 | Q72;;MHC/+ | female | -RU486 | 99 | 66.30 | 68.00 | 76.00 |
| 7 | Q72;;MHC/+ | female | -RU486 | 124 | 52.97 | 52.00 | 69.75 |
| 8 | Q72;;MHC/+ | female | -RU486 | 120 | 49.81 | 46.00 | 66.00 |
| 1 | Q72/+;;MHC | female | -RU486 | 131 | 56.70 | 56.00 | 70.25 |
| 2 | Q72/+;;MHC | female | -RU486 | 85 | 58.39 | 56.00 | 69.75 |
| 3 | Q72/+;;MHC | female | -RU486 | 66 | 66.50 | 68.00 | 75.00 |
| 4 | Q72/+;;MHC | female | -RU486 | 119 | 69.38 | 73.00 | 80.75 |
| 5 | Q72/+;;MHC | female | -RU486 | 105 | 61.92 | 63.00 | 70.00 |
| 6 | Q72/+;;MHC | female | -RU486 | 104 | 66.85 | 68.00 | 72.25 |
| 7 | Q72/+;;MHC | female | -RU486 | 143 | 62.52 | 62.00 | 69.75 |
| 8 | Q72/+;;MHC | female | -RU486 | 148 | 64.60 | 67.00 | 72.00 |
| 1 | Q72/+;;MHC/+ | female | -RU486 | 126 | 48.34 | 49.00 | 66.75 |
| 2 | Q72/+;;MHC/+ | female | -RU486 | 94 | 65.52 | 66.00 | 74.00 |
| 3 | Q72/+;;MHC/+ | female | -RU486 | 106 | 66.84 | 72.00 | 85.50 |
| 4 | Q72/+;;MHC/+ | female | -RU486 | 99 | 81.41 | 84.00 | 90.75 |
| 5 | Q72/+;;MHC/+ | female | -RU486 | 112 | 52.05 | 54.00 | 66.00 |
| 6 | Q72/+;;MHC/+ | female | -RU486 | 65 | 59.22 | 61.00 | 71.33 |
| 7 | Q72/+;;MHC/+ | female | -RU486 | 124 | 53.94 | 55.00 | 74.75 |
| 8 | Q72/+;;MHC/+ | female | -RU486 | 163 | 46.95 | 43.00 | 69.75 |
| 1 | Q72/+;;MHC/UAS-EDTP | female | -RU486 | 154 | 52.28 | 54.00 | 70.25 |
| 2 | Q72/+;;MHC/UAS-EDTP | female | -RU486 | 92 | 45.94 | 45.00 | 68.75 |
| 3 | Q72/+;;MHC/UAS-EDTP | female | -RU486 | 101 | 56.91 | 54.00 | 80.25 |
| 4 | Q72/+;;MHC/UAS-EDTP | female | -RU486 | 104 | 74.48 | 77.00 | 88.00 |
| 1 | UAS-EDTP/+ | female | -RU486 | 144 | 35.19 | 35.00 | 46.25 |
| 2 | UAS-EDTP/+ | female | -RU486 | 114 | 47.02 | 45.00 | 63.00 |
| 3 | UAS-EDTP/+ | female | -RU486 | 106 | 54.28 | 54.00 | 73.50 |
| 4 | UAS-EDTP/+ | female | -RU486 | 95 | 59.76 | 56.00 | 81.50 |
| 1 | UAS-EDTP/MHC | female | -RU486 | 124 | 55.15 | 54.00 | 70.75 |
| 2 | UAS-EDTP/MHC | female | -RU486 | 128 | 59.68 | 62.00 | 72.25 |
| 3 | UAS-EDTP/MHC | female | -RU486 | 78 | 62.90 | 68.00 | 73.75 |
| 4 | UAS-EDTP/MHC | female | -RU486 | 90 | 68.53 | 73.00 | 83.75 |

| Replicate | Genotype | Sex | Treatment | n | Mean | Median | Maximum |
|---|---|---|---|---|---|---|---|
| 1 | MHC/+ | female | +RU486 | 124 | 49.87 | 51.00 | 71.50 |
| 2 | MHC/+ | female | +RU486 | 93 | 59.47 | 63.00 | 75.00 |
| 3 | MHC/+ | female | +RU486 | 97 | 65.17 | 65.00 | 82.50 |
| 4 | MHC/+ | female | +RU486 | 120 | 73.26 | 77.00 | 87.50 |
| 1 | Q72;;MHC | female | +RU486 | 136 | 18.88 | 19.00 | 24.50 |
| 2 | Q72;;MHC | female | +RU486 | 141 | 17.75 | 19.00 | 24.50 |
| 3 | Q72;;MHC | female | +RU486 | 96 | 22.68 | 23.00 | 26.00 |
| 4 | Q72;;MHC | female | +RU486 | 93 | 18.94 | 19.00 | 25.50 |
| 5 | Q72;;MHC | female | +RU486 | 77 | 23.96 | 26.00 | 27.33 |
| 6 | Q72;;MHC | female | +RU486 | 90 | 23.46 | 24.00 | 29.33 |
| 7 | Q72;;MHC | female | +RU486 | 116 | 18.08 | 20.00 | 25.50 |
| 8 | Q72;;MHC | female | +RU486 | 188 | 15.89 | 18.00 | 25.00 |
| 1 | Q72;;MHC/+ | female | +RU486 | 134 | 24.11 | 26.00 | 27.50 |
| 2 | Q72;;MHC/+ | female | +RU486 | 127 | 25.17 | 26.00 | 30.25 |
| 3 | Q72;;MHC/+ | female | +RU486 | 119 | 28.52 | 28.00 | 32.25 |
| 4 | Q72;;MHC/+ | female | +RU486 | 90 | 26.36 | 28.00 | 32.00 |
| 5 | Q72;;MHC/+ | female | +RU486 | 102 | 24.15 | 26.00 | 27.00 |
| 6 | Q72;;MHC/+ | female | +RU486 | 124 | 25.27 | 26.00 | 29.50 |
| 7 | Q72;;MHC/+ | female | +RU486 | 129 | 24.90 | 27.00 | 29.00 |
| 8 | Q72;;MHC/+ | female | +RU486 | 130 | 23.73 | 25.00 | 27.50 |
| 1 | Q72/+;;MHC | female | +RU486 | 132 | 15.82 | 16.00 | 25.75 |
| 2 | Q72/+;;MHC | female | +RU486 | 100 | 15.64 | 17.00 | 24.00 |
| 3 | Q72/+;;MHC | female | +RU486 | 82 | 21.86 | 21.00 | 26.50 |
| 4 | Q72/+;;MHC | female | +RU486 | 116 | 16.98 | 17.00 | 26.00 |
| 5 | Q72/+;;MHC | female | +RU486 | 105 | 23.34 | 26.00 | 32.25 |
| 6 | Q72/+;;MHC | female | +RU486 | 110 | 22.21 | 24.00 | 31.75 |
| 7 | Q72/+;;MHC | female | +RU486 | 129 | 15.92 | 20.00 | 24.75 |
| 8 | Q72/+;;MHC | female | +RU486 | 141 | 10.80 | 6.00 | 21.75 |
| 1 | Q72/+;;MHC/+ | female | +RU486 | 111 | 22.55 | 23.00 | 26.00 |
| 2 | Q72/+;;MHC/+ | female | +RU486 | 119 | 21.59 | 21.00 | 24.50 |
| 3 | Q72/+;;MHC/+ | female | +RU486 | 106 | 22.63 | 23.00 | 26.50 |
| 4 | Q72/+;;MHC/+ | female | +RU486 | 86 | 23.43 | 24.00 | 25.50 |
| 5 | Q72/+;;MHC/+ | female | +RU486 | 104 | 23.81 | 23.00 | 26.50 |
| 6 | Q72/+;;MHC/+ | female | +RU486 | 97 | 24.05 | 24.00 | 27.25 |
| 7 | Q72/+;;MHC/+ | female | +RU486 | 133 | 20.44 | 20.00 | 25.50 |
| 8 | Q72/+;;MHC/+ | female | +RU486 | 151 | 20.66 | 20.00 | 24.25 |
| 1 | Q72/+;;MHC/UAS-EDTP | female | +RU486 | 145 | 22.84 | 23.00 | 26.00 |
| 2 | Q72/+;;MHC/UAS-EDTP | female | +RU486 | 103 | 22.36 | 24.00 | 25.50 |
| 3 | Q72/+;;MHC/UAS-EDTP | female | +RU486 | 79 | 24.59 | 26.00 | 28.00 |
| 4 | Q72/+;;MHC/UAS-EDTP | female | +RU486 | 94 | 25.86 | 26.00 | 30.25 |
| 1 | UAS-EDTP/+ | female | +RU486 | 134 | 36.98 | 35.00 | 57.25 |

| Replicate | Genotype | Sex | Treatment | n | Mean | Median | Maximum |
|---|---|---|---|---|---|---|---|
| 2 | UAS-EDTP/+ | female | +RU486 | 116 | 46.58 | 45.00 | 66.00 |
| 3 | UAS-EDTP/+ | female | +RU486 | 96 | 54.57 | 54.00 | 76.00 |
| 4 | UAS-EDTP/+ | female | +RU486 | 98 | 60.81 | 60.00 | 85.00 |
| 1 | UAS-EDTP/MHC | female | +RU486 | 117 | 56.73 | 56.00 | 75.00 |
| 2 | UAS-EDTP/MHC | female | +RU486 | 112 | 58.59 | 61.00 | 72.25 |
| 3 | UAS-EDTP/MHC | female | +RU486 | 70 | 65.22 | 68.00 | 78.00 |
| 4 | UAS-EDTP/MHC | female | +RU486 | 107 | 63.34 | 66.00 | 79.25 |
| 1 | DJ694 | female | none | 142 | 40.59 | 42.00 | 51.50 |
| 2 | DJ694 | female | none | 129 | 39.49 | 40.00 | 49.75 |
| 3 | DJ694 | female | none | 78 | 39.64 | 40.00 | 52.00 |
| 4 | DJ694 | female | none | 99 | 42.97 | 45.00 | 52.50 |
| 7 | DJ694 | female | none | 121 | 45.01 | 41.00 | 48.00 |
| 8 | DJ694 | female | none | 120 | 41.89 | 43.00 | 49.00 |
| 1 | DJ694;UAS-EDTP/+ | female | none | 148 | 39.30 | 38.50 | 53.00 |
| 2 | DJ694;UAS-EDTP/+ | female | none | 154 | 39.04 | 38.00 | 54.75 |
| 3 | DJ694;UAS-EDTP/+ | female | none | 118 | 46.27 | 47.00 | 63.75 |
| 4 | DJ694;UAS-EDTP/+ | female | none | 147 | 46.49 | 45.00 | 68.00 |
| 7 | DJ694;UAS-EDTP/+ | female | none | 145 | 37.28 | 34.00 | 51.75 |
| 8 | DJ694;UAS-EDTP/+ | female | none | 171 | 40.79 | 43.00 | 50.00 |
| 1 | DJ694/+ | female | none | 126 | 48.41 | 49.00 | 57.25 |
| 2 | DJ694/+ | female | none | 141 | 46.60 | 47.00 | 60.00 |
| 3 | DJ694/+ | female | none | 101 | 54.46 | 54.00 | 71.25 |
| 4 | DJ694/+ | female | none | 91 | 56.82 | 56.00 | 79.25 |
| 1 | Q72;DJ694 | female | none | 162 | 28.73 | 30.00 | 33.00 |
| 2 | Q72;DJ694 | female | none | 89 | 28.24 | 31.00 | 31.00 |
| 3 | Q72;DJ694 | female | none | 120 | 30.40 | 30.00 | 35.50 |
| 4 | Q72;DJ694 | female | none | 82 | 32.69 | 33.00 | 38.75 |
| 5 | Q72;DJ694 | female | none | 149 | 29.66 | 31.00 | 34.50 |
| 6 | Q72;DJ694 | female | none | 206 | 31.01 | 31.00 | 37.00 |
| 7 | Q72;DJ694 | female | none | 156 | 27.22 | 27.00 | 33.25 |
| 8 | Q72;DJ694 | female | none | 128 | 27.29 | 27.00 | 33.00 |
| 1 | Q72;DJ694/+ | female | none | 140 | 45.44 | 47.00 | 53.25 |
| 2 | Q72;DJ694/+ | female | none | 134 | 44.16 | 45.00 | 52.00 |
| 3 | Q72;DJ694/+ | female | none | 132 | 50.41 | 51.00 | 54.50 |
| 4 | Q72;DJ694/+ | female | none | 90 | 47.35 | 49.00 | 53.33 |
| 5 | Q72;DJ694/+ | female | none | 89 | 55.58 | 56.00 | 61.00 |
| 6 | Q72;DJ694/+ | female | none | 98 | 56.30 | 59.00 | 68.00 |
| 7 | Q72;DJ694/+ | female | none | 151 | 45.09 | 45.00 | 51.75 |
| 8 | Q72;DJ694/+ | female | none | 139 | 44.13 | 46.00 | 52.25 |
| 1 | Q72/+;DJ694 | female | none | 96 | 30.22 | 30.00 | 36.75 |
| 2 | Q72/+;DJ694 | female | none | 88 | 32.07 | 31.00 | 38.00 |

| Replicate | Genotype | Sex | Treatment | n | Mean | Median | Maximum |
|---|---|---|---|---|---|---|---|
| 3 | Q72/+;DJ694 | female | none | 84 | 31.51 | 33.00 | 37.50 |
| 4 | Q72/+;DJ694 | female | none | 96 | 35.57 | 38.00 | 45.50 |
| 5 | Q72/+;DJ694 | female | none | 106 | 32.17 | 33.00 | 36.75 |
| 6 | Q72/+;DJ694 | female | none | 24 | 37.83 | 38.00 | 45.00 |
| 6 | Q72/+;DJ694 | female | none | 102 | 38.11 | 38.50 | 44.25 |
| 7 | Q72/+;DJ694 | female | none | 58 | 27.03 | 27.00 | 30.50 |
| 8 | Q72/+;DJ694 | female | none | 33 | 30.70 | 32.00 | 39.00 |
| 1 | Q72/+;DJ694;UAS-EDTP/+ | female | none | 86 | 39.98 | 40.00 | 47.00 |
| 2 | Q72/+;DJ694;UAS-EDTP/+ | female | none | 120 | 36.39 | 38.00 | 41.75 |
| 3 | Q72/+;DJ694;UAS-EDTP/+ | female | none | 81 | 41.00 | 40.00 | 47.25 |
| 4 | Q72/+;DJ694;UAS-EDTP/+ | female | none | 160 | 41.35 | 45.00 | 48.00 |
| 7 | Q72/+;DJ694;UAS-EDTP/+ | female | none | 69 | 35.90 | 34.00 | 41.00 |
| 8 | Q72/+;DJ694;UAS-EDTP/+ | female | none | 56 | 40.36 | 41.00 | 48.00 |
| 1 | Q72/+;DJ694/+ | female | none | 141 | 40.45 | 40.00 | 47.00 |
| 2 | Q72/+;DJ694/+ | female | none | 195 | 35.58 | 38.00 | 41.00 |
| 3 | Q72/+;DJ694/+ | female | none | 84 | 42.22 | 47.00 | 49.50 |
| 4 | Q72/+;DJ694/+ | female | none | 88 | 46.94 | 45.00 | 51.25 |
| 5 | Q72/+;DJ694/+ | female | none | 152 | 42.21 | 40.00 | 47.00 |
| 6 | Q72/+;DJ694/+ | female | none | 112 | 44.94 | 45.00 | 53.00 |
| 7 | Q72/+;DJ694/+ | female | none | 166 | 39.98 | 41.00 | 48.00 |
| 8 | Q72/+;DJ694/+ | female | none | 185 | 39.60 | 39.00 | 47.50 |
| 1 | UAS-EDTP/+ | female | none | 145 | 36.85 | 37.00 | 49.75 |
| 2 | UAS-EDTP/+ | female | none | 127 | 39.86 | 42.00 | 48.00 |
| 3 | UAS-EDTP/+ | female | none | 94 | 56.13 | 54.00 | 79.50 |
| 4 | UAS-EDTP/+ | female | none | 96 | 63.58 | 66.00 | 80.50 |

Table 4 Temporal repression of Q72 using GAL80ts longevity dataset (related to Figure 5)

| 20-20°C | | | | | | |
|---|---|---|---|---|---|---|
| Replicate | Genotype | Sex | n | Mean | Median | Maximum |
| 1 | Mef2/+ | male | 122 | 121.52 | 133.00 | 170.25 |
| 2 | Mef2/+ | male | 116 | 135.41 | 142.00 | 168.25 |
| 3 | Mef2/+ | male | 122 | 111.91 | 120.00 | 168.75 |
| 4 | Mef2/+ | male | 83 | 139.18 | 143.00 | 187.75 |
| 5 | Mef2/+ | male | 121 | 120.37 | 130.00 | 175.50 |
| 6 | Mef2/+ | male | 119 | 115.82 | 136.00 | 177.50 |
| 1 | Q72/+ | male | 118 | 136.47 | 149.00 | 175.75 |
| 2 | Q72/+ | male | 111 | 140.92 | 149.00 | 174.50 |
| 3 | Q72/+ | male | 80 | 137.83 | 148.00 | 182.25 |
| 4 | Q72/+ | male | 93 | 145.74 | 146.00 | 182.50 |
| 5 | Q72/+ | male | 48 | 136.79 | 137.00 | 172.00 |
| 6 | Q72/+ | male | 97 | 125.14 | 143.00 | 175.75 |
| 1 | Q72/+;;GAL80ts/+ | male | 128 | 146.49 | 149.00 | 169.75 |
| 2 | Q72/+;;GAL80ts/+ | male | 116 | 150.77 | 153.00 | 182.75 |
| 3 | Q72/+;;GAL80ts/+ | male | 96 | 147.75 | 155.00 | 176.75 |
| 4 | Q72/+;;GAL80ts/+ | male | 74 | 160.52 | 167.00 | 189.75 |
| 5 | Q72/+;;GAL80ts/+ | male | 105 | 128.15 | 143.00 | 169.50 |
| 6 | Q72/+;;GAL80ts/+ | male | 108 | 133.75 | 143.00 | 173.25 |
| 1 | Q72/+;;Mef2/+ | male | 104 | 45.44 | 44.00 | 65.00 |
| 2 | Q72/+;;Mef2/+ | male | 67 | 53.81 | 51.00 | 70.00 |
| 3 | Q72/+;;Mef2/+ | male | 98 | 45.80 | 46.00 | 63.75 |
| 4 | Q72/+;;Mef2/+ | male | 56 | 48.67 | 48.00 | 70.50 |
| 5 | Q72/+;;Mef2/+ | male | 87 | 50.96 | 54.00 | 65.67 |
| 6 | Q72/+;;Mef2/+ | male | 175 | 52.17 | 52.00 | 65.25 |
| 1 | Q72/+;;Mef2/GAL80ts | male | 153 | 160.34 | 173.00 | 196.00 |
| 2 | Q72/+;;Mef2/GAL80ts | male | 128 | 168.81 | 170.00 | 194.75 |
| 3 | Q72/+;;Mef2/GAL80ts | male | 98 | 153.43 | 160.00 | 182.25 |
| 4 | Q72/+;;Mef2/GAL80ts | male | 75 | 154.66 | 153.00 | 182.00 |
| 5 | Q72/+;;Mef2/GAL80ts | male | 92 | 161.02 | 165.00 | 190.25 |
| 6 | Q72/+;;Mef2/GAL80ts | male | 125 | 134.84 | 154.00 | 178.25 |
| 1 | Mef2/+ | female | 129 | 162.90 | 171.00 | 190.75 |
| 2 | Mef2/+ | female | 137 | 147.59 | 160.00 | 173.50 |
| 3 | Mef2/+ | female | 122 | 152.31 | 155.00 | 178.75 |
| 4 | Mef2/+ | female | 113 | 73.98 | 3.00 | 184.50 |
| 5 | Mef2/+ | female | 146 | 123.46 | 147.00 | 180.75 |
| 6 | Mef2/+ | female | 139 | 141.39 | 154.00 | 180.25 |
| 1 | Q72/+ | female | 124 | 135.83 | 149.00 | 170.25 |
| 2 | Q72/+ | female | 136 | 151.92 | 156.00 | 170.00 |
| 3 | Q72/+ | female | 100 | 149.67 | 151.00 | 175.75 |
| 4 | Q72/+ | female | 98 | 147.64 | 146.00 | 174.00 |

| | | | | | | |
|---|---|---|---|---|---|---|
| 5 | Q72/+ | female | 112 | 147.70 | 149.00 | 179.75 |
| 6 | Q72/+ | female | 106 | 131.70 | 147.00 | 165.00 |
| 1 | Q72/+;;GAL80ts/+ | female | 133 | 166.68 | 171.00 | 187.00 |
| 2 | Q72/+;;GAL80ts/+ | female | 159 | 164.76 | 168.00 | 191.25 |
| 3 | Q72/+;;GAL80ts/+ | female | 109 | 161.79 | 162.00 | 179.50 |
| 4 | Q72/+;;GAL80ts/+ | female | 117 | 140.03 | 167.00 | 187.00 |
| 5 | Q72/+;;GAL80ts/+ | female | 121 | 146.91 | 161.00 | 183.25 |
| 6 | Q72/+;;GAL80ts/+ | female | 112 | 153.63 | 161.00 | 182.75 |
| 1 | Q72/+;;Mef2/+ | female | 135 | 56.18 | 61.00 | 90.00 |
| 2 | Q72/+;;Mef2/+ | female | 120 | 64.43 | 63.50 | 83.75 |
| 3 | Q72/+;;Mef2/+ | female | 113 | 57.09 | 57.00 | 71.75 |
| 4 | Q72/+;;Mef2/+ | female | 125 | 57.46 | 58.00 | 74.00 |
| 5 | Q72/+;;Mef2/+ | female | 168 | 54.38 | 56.00 | 68.50 |
| 6 | Q72/+;;Mef2/+ | female | 270 | 57.17 | 59.00 | 68.00 |
| 1 | Q72/+;;Mef2/GAL80ts | female | 169 | 178.79 | 180.00 | 215.25 |
| 2 | Q72/+;;Mef2/GAL80ts | female | 185 | 169.69 | 170.00 | 190.00 |
| 3 | Q72/+;;Mef2/GAL80ts | female | 109 | 180.18 | 183.00 | 203.50 |
| 4 | Q72/+;;Mef2/GAL80ts | female | 108 | 163.48 | 179.50 | 200.25 |
| 5 | Q72/+;;Mef2/GAL80ts | female | 99 | 161.70 | 176.00 | 199.00 |
| 6 | Q72/+;;Mef2/GAL80ts | female | 102 | 156.18 | 172.00 | 190.50 |

| 20-29°C | | | | | | |
|---|---|---|---|---|---|---|
| Replicate | Genotype | Sex | n | Mean | Median | Maximum |
| 1 | Mef2/+ | male | 131 | 37.71 | 40.00 | 41.75 |
| 2 | Mef2/+ | male | 109 | 37.15 | 37.00 | 40.50 |
| 3 | Mef2/+ | male | 132 | 21.29 | 23.00 | 34.50 |
| 4 | Mef2/+ | male | 99 | 27.83 | 27.00 | 44.50 |
| 1 | Q72/+ | male | 112 | 37.51 | 40.00 | 47.25 |
| 2 | Q72/+ | male | 74 | 40.46 | 41.00 | 44.67 |
| 3 | Q72/+ | male | 82 | 29.60 | 32.00 | 43.25 |
| 4 | Q72/+ | male | 79 | 36.70 | 37.00 | 51.00 |
| 1 | Q72/+;;GAL80ts/+ | male | 118 | 39.40 | 40.00 | 44.25 |
| 2 | Q72/+;;GAL80ts/+ | male | 121 | 39.14 | 41.00 | 46.75 |
| 3 | Q72/+;;GAL80ts/+ | male | 107 | 31.39 | 32.00 | 42.50 |
| 4 | Q72/+;;GAL80ts/+ | male | 115 | 33.57 | 30.00 | 50.25 |
| 1 | Q72/+;;Mef2/+ | male | 97 | 11.34 | 12.00 | 13.50 |
| 2 | Q72/+;;Mef2/+ | male | 50 | 12.14 | 13.00 | 17.00 |
| 3 | Q72/+;;Mef2/+ | male | 101 | 10.31 | 11.00 | 13.00 |
| 4 | Q72/+;;Mef2/+ | male | 48 | 11.58 | 10.00 | 16.00 |
| 1 | Q72/+;;Mef2/GAL80ts | male | 133 | 33.16 | 35.00 | 40.50 |
| 2 | Q72/+;;Mef2/GAL80ts | male | 110 | 34.81 | 37.00 | 39.50 |
| 3 | Q72/+;;Mef2/GAL80ts | male | 123 | 20.81 | 18.00 | 37.50 |
| 4 | Q72/+;;Mef2/GAL80ts | male | 99 | 27.97 | 30.00 | 34.25 |

| | | | | | | |
|---|---|---|---|---|---|---|
| 1 | Mef2/+ | female | 117 | 45.26 | 47.00 | 48.50 |
| 2 | Mef2/+ | female | 136 | 43.92 | 46.00 | 49.50 |
| 3 | Mef2/+ | female | 114 | 52.40 | 53.00 | 60.00 |
| 4 | Mef2/+ | female | 118 | 27.68 | 3.00 | 58.00 |
| 1 | Q72/+ | female | 104 | 45.51 | 47.00 | 49.00 |
| 2 | Q72/+ | female | 85 | 47.28 | 48.00 | 50.25 |
| 3 | Q72/+ | female | 110 | 52.80 | 53.00 | 60.25 |
| 4 | Q72/+ | female | 132 | 48.43 | 51.00 | 58.00 |
| 1 | Q72/+;;GAL80ts/+ | female | 116 | 45.77 | 47.00 | 47.00 |
| 2 | Q72/+;;GAL80ts/+ | female | 116 | 45.44 | 46.00 | 47.50 |
| 3 | Q72/+;;GAL80ts/+ | female | 120 | 54.47 | 53.00 | 61.00 |
| 4 | Q72/+;;GAL80ts/+ | female | 120 | 44.09 | 55.00 | 58.00 |
| 1 | Q72/+;;Mef2/+ | female | 137 | 11.88 | 12.00 | 16.25 |
| 2 | Q72/+;;Mef2/+ | female | 143 | 13.08 | 13.00 | 16.25 |
| 3 | Q72/+;;Mef2/+ | female | 125 | 12.30 | 13.00 | 15.25 |
| 4 | Q72/+;;Mef2/+ | female | 129 | 11.83 | 11.00 | 15.25 |
| 1 | Q72/+;;Mef2/GAL80ts | female | 131 | 43.26 | 47.00 | 48.00 |
| 2 | Q72/+;;Mef2/GAL80ts | female | 102 | 43.99 | 44.00 | 47.00 |
| 3 | Q72/+;;Mef2/GAL80ts | female | 133 | 36.75 | 36.00 | 51.00 |
| 4 | Q72/+;;Mef2/GAL80ts | female | 146 | 34.99 | 39.00 | 50.00 |
| | | 29-20°C | | | | |
| **Replicate** | **Genotype** | **Sex** | **n** | **Mean** | **Median** | **Maximum** |
| 1 | Mef2/+ | male | 160 | 135.34 | 137.00 | 165.75 |
| 2 | Mef2/+ | male | 171 | 141.22 | 144.00 | 176.25 |
| 3 | Mef2/+ | male | 121 | 127.50 | 133.00 | 181.00 |
| 4 | Mef2/+ | male | 125 | 135.91 | 151.00 | 185.00 |
| 1 | Q72/+ | male | 162 | 145.41 | 150.00 | 193.25 |
| 2 | Q72/+ | male | 139 | 147.22 | 158.00 | 198.00 |
| 3 | Q72/+ | male | 138 | 145.70 | 158.00 | 189.75 |
| 4 | Q72/+ | male | 123 | 145.43 | 158.00 | 199.00 |
| 1 | Q72/+;;GAL80ts/+ | male | 154 | 154.00 | 163.00 | 186.25 |
| 2 | Q72/+;;GAL80ts/+ | male | 125 | 167.62 | 172.00 | 194.50 |
| 3 | Q72/+;;GAL80ts/+ | male | 107 | 163.28 | 175.00 | 197.75 |
| 4 | Q72/+;;GAL80ts/+ | male | 118 | 162.44 | 175.00 | 196.50 |
| 1 | Q72/+;;Mef2/+ | male | 126 | 8.77 | 9.00 | 10.50 |
| 2 | Q72/+;;Mef2/+ | male | 126 | 6.71 | 7.00 | 10.50 |
| 3 | Q72/+;;Mef2/+ | male | 78 | 7.28 | 7.00 | 10.00 |
| 4 | Q72/+;;Mef2/+ | male | 130 | 5.38 | 4.00 | 9.50 |
| 1 | Q72/+;;Mef2/GAL80ts | male | 144 | 152.62 | 158.00 | 181.00 |
| 2 | Q72/+;;Mef2/GAL80ts | male | 168 | 152.68 | 157.00 | 193.75 |
| 3 | Q72/+;;Mef2/GAL80ts | male | 111 | 128.09 | 133.00 | 167.00 |
| 4 | Q72/+;;Mef2/GAL80ts | male | 127 | 91.66 | 91.00 | 150.25 |

| | | | | | | |
|---|---|---|---|---|---|---|
| 1 | Mef2/+ | female | 182 | 137.03 | 137.00 | 163.50 |
| 2 | Mef2/+ | female | 151 | 152.05 | 154.00 | 180.75 |
| 3 | Mef2/+ | female | 120 | 114.10 | 140.00 | 174.75 |
| 4 | Mef2/+ | female | 117 | 139.83 | 147.00 | 175.25 |
| 1 | Q72/+ | female | 148 | 124.58 | 125.00 | 163.50 |
| 2 | Q72/+ | female | 159 | 111.16 | 112.00 | 159.50 |
| 3 | Q72/+ | female | 135 | 128.82 | 136.00 | 164.75 |
| 4 | Q72/+ | female | 168 | 112.73 | 116.00 | 159.50 |
| 1 | Q72/+;;GAL80ts/+ | female | 148 | 141.05 | 149.00 | 170.75 |
| 2 | Q72/+;;GAL80ts/+ | female | 138 | 145.18 | 151.00 | 175.50 |
| 3 | Q72/+;;GAL80ts/+ | female | 112 | 121.26 | 133.00 | 161.75 |
| 4 | Q72/+;;GAL80ts/+ | female | 125 | 135.20 | 151.00 | 188.25 |
| 1 | Q72/+;;Mef2/+ | female | 184 | 9.67 | 9.00 | 12.50 |
| 2 | Q72/+;;Mef2/+ | female | 153 | 7.82 | 7.00 | 11.00 |
| 3 | Q72/+;;Mef2/+ | female | 124 | 8.35 | 7.00 | 12.00 |
| 4 | Q72/+;;Mef2/+ | female | 176 | 6.98 | 7.00 | 12.00 |
| 1 | Q72/+;;Mef2/GAL80ts | female | 154 | 159.87 | 160.00 | 186.25 |
| 2 | Q72/+;;Mef2/GAL80ts | female | 164 | 161.99 | 158.00 | 184.00 |
| 3 | Q72/+;;Mef2/GAL80ts | female | 110 | 130.49 | 140.00 | 170.50 |
| 4 | Q72/+;;Mef2/GAL80ts | female | 130 | 121.53 | 122.00 | 158.25 |
| | | | 29-29°C | | | |
| **Replicate** | **Genotype** | **Sex** | **n** | **Mean** | **Median** | **Maximum** |
| 1 | Mef2/+ | male | 162 | 47.73 | 51.00 | 53.50 |
| 2 | Mef2/+ | male | 142 | 45.98 | 49.00 | 50.50 |
| 3 | Mef2/+ | male | 123 | 33.67 | 33.00 | 38.75 |
| 4 | Mef2/+ | male | 121 | 33.63 | 33.00 | 45.00 |
| 1 | Q72/+ | male | 150 | 50.82 | 51.00 | 55.00 |
| 2 | Q72/+ | male | 118 | 46.04 | 49.00 | 51.00 |
| 3 | Q72/+ | male | 141 | 38.54 | 38.00 | 48.75 |
| 4 | Q72/+ | male | 161 | 37.60 | 42.00 | 48.25 |
| 1 | Q72/+;;GAL80ts/+ | male | 132 | 48.41 | 51.00 | 53.50 |
| 2 | Q72/+;;GAL80ts/+ | male | 129 | 46.34 | 49.00 | 50.00 |
| 3 | Q72/+;;GAL80ts/+ | male | 114 | 44.52 | 47.00 | 54.25 |
| 4 | Q72/+;;GAL80ts/+ | male | 136 | 43.27 | 44.00 | 53.50 |
| 1 | Q72/+;;Mef2/+ | male | 121 | 2.58 | 2.00 | 4.50 |
| 2 | Q72/+;;Mef2/+ | male | 119 | 2.54 | 2.00 | 4.75 |
| 3 | Q72/+;;Mef2/+ | male | 82 | 3.48 | 3.00 | 5.00 |
| 4 | Q72/+;;Mef2/+ | male | 108 | 3.00 | 2.00 | 4.00 |
| 1 | Q72/+;;Mef2/GAL80ts | male | 157 | 23.72 | 23.00 | 32.25 |
| 2 | Q72/+;;Mef2/GAL80ts | male | 188 | 20.86 | 21.00 | 28.00 |
| 3 | Q72/+;;Mef2/GAL80ts | male | 115 | 25.80 | 26.00 | 36.75 |
| 4 | Q72/+;;Mef2/GAL80ts | male | 116 | 22.59 | 21.00 | 32.00 |

| | | | | | | |
|---|---|---|---|---|---|---|
| 1 | Mef2/+ | female | 167 | 52.08 | 53.00 | 55.75 |
| 2 | Mef2/+ | female | 164 | 47.66 | 49.00 | 52.50 |
| 3 | Mef2/+ | female | 132 | 47.03 | 56.00 | 59.00 |
| 4 | Mef2/+ | female | 129 | 49.45 | 49.00 | 54.75 |
| 1 | Q72/+ | female | 146 | 45.92 | 48.00 | 53.00 |
| 2 | Q72/+ | female | 139 | 40.54 | 44.00 | 49.00 |
| 3 | Q72/+ | female | 136 | 46.98 | 52.00 | 56.25 |
| 4 | Q72/+ | female | 167 | 39.66 | 42.00 | 46.50 |
| 1 | Q72/+;;GAL80ts/+ | female | 157 | 50.06 | 51.00 | 54.00 |
| 2 | Q72/+;;GAL80ts/+ | female | 149 | 45.43 | 49.00 | 51.00 |
| 3 | Q72/+;;GAL80ts/+ | female | 112 | 47.36 | 47.00 | 55.50 |
| 4 | Q72/+;;GAL80ts/+ | female | 135 | 47.07 | 49.00 | 56.00 |
| 1 | Q72/+;;Mef2/+ | female | 174 | 3.35 | 4.00 | 6.00 |
| 2 | Q72/+;;Mef2/+ | female | 155 | 2.82 | 2.00 | 4.00 |
| 3 | Q72/+;;Mef2/+ | female | 133 | 3.78 | 3.00 | 6.00 |
| 4 | Q72/+;;Mef2/+ | female | 132 | 3.71 | 4.00 | 5.50 |
| 1 | Q72/+;;Mef2/GAL80ts | female | 152 | 21.51 | 20.00 | 31.25 |
| 2 | Q72/+;;Mef2/GAL80ts | female | 171 | 22.40 | 21.00 | 32.75 |
| 3 | Q72/+;;Mef2/GAL80ts | female | 107 | 19.80 | 21.00 | 25.75 |
| 4 | Q72/+;;Mef2/GAL80ts | female | 117 | 18.30 | 18.00 | 23.50 |

Table 5 Expressing Hsp70 in polyQ flies longevity dataset (related to Figure 6, S7)

| Replicate | Genotype | Sex | Treatment | n | Mean | Median | Maximum |
|---|---|---|---|---|---|---|---|
| 1 | 4.2/+ | male | +RU486 | 98 | 72.27 | 74.00 | 81.00 |
| 2 | 4.2/+ | male | +RU486 | 132 | 62.55 | 63.00 | 77.00 |
| 1 | 4.4/+ | male | +RU486 | 112 | 46.85 | 49.00 | 57.50 |
| 2 | 4.4/+ | male | +RU486 | 134 | 42.55 | 42.00 | 53.75 |
| 1 | MHC/+ | male | +RU486 | 94 | 58.58 | 59.00 | 72.75 |
| 2 | MHC/+ | male | +RU486 | 144 | 47.59 | 49.00 | 64.00 |
| 1 | MHC/4.2 | male | +RU486 | 65 | 58.18 | 59.00 | 68.67 |
| 2 | MHC/4.2 | male | +RU486 | 118 | 58.76 | 58.00 | 74.50 |
| 1 | MHC/4.4 | male | +RU486 | 94 | 36.52 | 36.50 | 46.75 |
| 2 | MHC/4.4 | male | +RU486 | 164 | 35.41 | 35.00 | 44.75 |
| 1 | Q72/+ | male | +RU486 | 112 | 63.43 | 66.00 | 78.50 |
| 2 | Q72/+ | male | +RU486 | 119 | 59.05 | 58.00 | 74.50 |
| 1 | Q72/+;;MHC/+ | male | +RU486 | 91 | 28.27 | 28.00 | 31.00 |
| 2 | Q72/+;;MHC/+ | male | +RU486 | 182 | 26.21 | 28.00 | 29.50 |
| 1 | Q72/+;;MHC/4.2 | male | +RU486 | 86 | 26.84 | 26.00 | 31.00 |
| 2 | Q72/+;;MHC/4.2 | male | +RU486 | 116 | 25.77 | 25.00 | 31.00 |
| 1 | Q72/+;;MHC/4.4 | male | +RU486 | 68 | 18.87 | 19.00 | 24.00 |
| 2 | Q72/+;;MHC/4.4 | male | +RU486 | 118 | 18.26 | 18.00 | 21.00 |
| 1 | Q72/+;;MHC/Bg3 | male | +RU486 | 107 | 26.71 | 26.00 | 32.50 |
| 2 | Q72/+;;MHC/Bg3 | male | +RU486 | 167 | 26.54 | 28.00 | 31.00 |
| 1 | 4.2/+ | male | none | 106 | 68.22 | 73.00 | 78.25 |
| 2 | 4.2/+ | male | none | 155 | 60.95 | 63.00 | 70.00 |
| 1 | 4.4/+ | male | none | 94 | 46.28 | 49.00 | 55.67 |
| 2 | 4.4/+ | male | none | 126 | 43.39 | 44.00 | 51.75 |
| 1 | Mef2/+ | male | none | 111 | 65.62 | 68.00 | 80.25 |
| 2 | Mef2/+ | male | none | 124 | 64.63 | 66.00 | 78.25 |
| 1 | Mef2/4.2 | male | none | 70 | 60.18 | 61.00 | 74.33 |
| 2 | Mef2/4.2 | male | none | 144 | 62.33 | 63.00 | 80.25 |
| 1 | Mef2/4.4 | male | none | 115 | 40.21 | 42.00 | 49.50 |
| 2 | Mef2/4.4 | male | none | 153 | 43.41 | 44.00 | 51.00 |
| 1 | Q72/+ | male | none | 93 | 57.99 | 59.00 | 73.25 |
| 2 | Q72/+ | male | none | 184 | 52.68 | 53.00 | 70.00 |
| 1 | Q72/+;;Mef2/+ | male | none | 91 | 9.21 | 10.00 | 17.00 |
| 2 | Q72/+;;Mef2/+ | male | none | 150 | 8.37 | 9.00 | 14.00 |
| 1 | Q72/+;;Mef2/4.2 | male | none | 103 | 20.40 | 21.00 | 31.50 |
| 2 | Q72/+;;Mef2/4.2 | male | none | 151 | 19.57 | 21.00 | 32.50 |
| 1 | Q72/+;;Mef2/4.4 | male | none | 79 | 5.02 | 5.00 | 9.00 |
| 2 | Q72/+;;Mef2/4.4 | male | none | 188 | 3.77 | 4.00 | 8.00 |
| 1 | Q72/+;;Mef2/Bg3 | male | none | 146 | 16.53 | 17.00 | 27.25 |
| 2 | Q72/+;;Mef2/Bg3 | male | none | 200 | 16.10 | 16.00 | 24.00 |
| 1 | 4.2/+ | female | +RU486 | 94 | 60.97 | 66.00 | 75.67 |

| Replicate | Genotype | Sex | Treatment | n | Mean | Median | Maximum |
|---|---|---|---|---|---|---|---|
| 2 | 4.2/+ | female | +RU486 | 132 | 55.74 | 60.00 | 71.50 |
| 1 | 4.4/+ | female | +RU486 | 113 | 33.92 | 33.00 | 54.50 |
| 2 | 4.4/+ | female | +RU486 | 131 | 35.58 | 32.00 | 51.50 |
| 1 | MHC/+ | female | +RU486 | 106 | 55.22 | 56.00 | 71.50 |
| 2 | MHC/+ | female | +RU486 | 116 | 56.47 | 58.00 | 69.50 |
| 1 | MHC/4.2 | female | +RU486 | 110 | 58.87 | 59.00 | 70.00 |
| 2 | MHC/4.2 | female | +RU486 | 118 | 61.75 | 63.00 | 71.00 |
| 1 | MHC/4.4 | female | +RU486 | 114 | 35.97 | 38.00 | 47.00 |
| 2 | MHC/4.4 | female | +RU486 | 138 | 36.18 | 35.00 | 48.25 |
| 1 | Q72/+ | female | +RU486 | 94 | 41.03 | 47.00 | 64.25 |
| 2 | Q72/+ | female | +RU486 | 124 | 42.66 | 42.00 | 58.25 |
| 1 | Q72/+;;MHC/+ | female | +RU486 | 124 | 23.99 | 24.00 | 29.75 |
| 2 | Q72/+;;MHC/+ | female | +RU486 | 159 | 25.50 | 25.00 | 29.50 |
| 1 | Q72/+;;MHC/4.2 | female | +RU486 | 120 | 23.80 | 24.00 | 27.00 |
| 2 | Q72/+;;MHC/4.2 | female | +RU486 | 135 | 23.17 | 23.00 | 26.50 |
| 1 | Q72/+;;MHC/4.4 | female | +RU486 | 111 | 18.73 | 19.00 | 22.50 |
| 2 | Q72/+;;MHC/4.4 | female | +RU486 | 121 | 17.89 | 18.00 | 21.00 |
| 1 | Q72/+;;MHC/Bg3 | female | +RU486 | 123 | 22.48 | 24.00 | 26.00 |
| 2 | Q72/+;;MHC/Bg3 | female | +RU486 | 148 | 22.21 | 23.00 | 25.75 |
| 1 | 4.2/+ | female | none | 135 | 59.67 | 63.00 | 71.50 |
| 2 | 4.2/+ | female | none | 161 | 61.48 | 63.00 | 70.00 |
| 1 | 4.4/+ | female | none | 96 | 40.54 | 40.00 | 58.00 |
| 2 | 4.4/+ | female | none | 152 | 34.49 | 33.50 | 50.50 |
| 1 | Mef2/+ | female | none | 118 | 55.30 | 66.00 | 72.25 |
| 2 | Mef2/+ | female | none | 113 | 64.07 | 65.00 | 73.00 |
| 1 | Mef2/4.2 | female | none | 113 | 70.99 | 75.00 | 80.00 |
| 2 | Mef2/4.2 | female | none | 127 | 71.03 | 74.00 | 80.75 |
| 1 | Mef2/4.4 | female | none | 130 | 38.72 | 40.00 | 54.50 |
| 2 | Mef2/4.4 | female | none | 154 | 41.62 | 42.00 | 55.25 |
| 1 | Q72/+ | female | none | 130 | 39.99 | 46.00 | 62.75 |
| 2 | Q72/+ | female | none | 153 | 35.72 | 35.00 | 55.75 |
| 1 | Q72/+;;Mef2/+ | female | none | 145 | 10.54 | 10.00 | 17.00 |
| 2 | Q72/+;;Mef2/+ | female | none | 159 | 6.55 | 7.00 | 14.00 |
| 1 | Q72/+;;Mef2/4.2 | female | none | 127 | 13.47 | 14.00 | 22.00 |
| 2 | Q72/+;;Mef2/4.2 | female | none | 158 | 7.73 | 9.00 | 14.50 |
| 1 | Q72/+;;Mef2/4.4 | female | none | 124 | 4.49 | 5.00 | 6.50 |
| 2 | Q72/+;;Mef2/4.4 | female | none | 181 | 3.05 | 2.00 | 5.50 |
| 1 | Q72/+;;Mef2/Bg3 | female | none | 140 | 10.99 | 10.00 | 19.00 |
| 2 | Q72/+;;Mef2/Bg3 | female | none | 203 | 8.54 | 9.00 | 16.75 |

Table 6 Compiled Logrank analysis of expression of Q72 in muscle longevity (related to Figure 1). Each p-value is from one experimental replicate

| Males | | | | | | | | |
|---|---|---|---|---|---|---|---|---|
| | Q72/+ | Q25/+ | DJ694/+ | Mef2/+ | MHC/+ | DJ694/+;Q25/+ | Mef2/Q25 | MHC/Q25 |
| Q72/+; DJ694/+ | 9.70e-09 | 4.44e-47 | 1.08e-22 | | | 3.88e-04 | | |
| | 2.29e-03 | 1.85e-65 | 1.60e-36 | | | 6.47e-02 | | |
| | 2.57e-20 | 7.09e-31 | 1.00e+00 | | | 5.43e-11 | | |
| | 5.45e-19 | 2.47e-14 | 8.02e-04 | | | 1.00e+00 | | |
| Q72/+;; Mef2/+ | 9.72e-100 | 5.36e-80 | | 2.83e-119 | | | 4.03e-98 | |
| | 5.91e-92 | 1.60e-84 | | 6.28e-106 | | | 3.94e-96 | |
| | 9.05e-49 | 1.99e-54 | | 8.00e-84 | | | 4.83e-53 | |
| | 1.62e-48 | 6.12e-50 | | 1.38e-92 | | | 8.48e-52 | |
| Q72/+;; MHC/+ | 6.03e-03 | 1.19e-44 | | | 5.75e-25 | | | 3.97e-06 |
| | 1.00e+00 | 3.16e-49 | | | 8.29e-12 | | | 1.00e+00 |
| | 2.21e-18 | 3.91e-35 | | | 4.91e-04 | | | 2.35e-02 |
| | 3.11e-20 | 4.82e-20 | | | 3.18e-10 | | | 1.00e+00 |
| Q72/+;; MHC/+ | 6.78e-42 | 4.16e-42 | | | 2.90e-57 | | | 2.14e-52 |
| +RU486 | 1.79e-53 | 1.72e-52 | | | 6.67e-68 | | | 1.60e-53 |
| Females | | | | | | | | |
| | Q72/+ | Q25/+ | DJ694/+ | Mef2/+ | MHC/+ | DJ694/+;Q25/+ | Mef2/Q25 | MHC/Q25 |
| Q72/+; DJ694/+ | 1.26e-43 | 3.43e-02 | 1.11e-04 | | | 5.05e-22 | | |
| | 3.53e-71 | 1.00e+00 | 5.83e-01 | | | 5.19e-03 | | |
| | 4.64e-04 | 4.56e-40 | 2.60e-07 | | | 1.66e-40 | | |
| | 1.00e+00 | 1.61e-14 | 2.41e-02 | | | 4.17e-28 | | |
| Q72/+;; Mef2/+ | 2.67e-93 | 4.40e-171 | | 8.23e-122 | | | 2.84e-97 | |
| | 6.70e-108 | 1.31e-115 | | 6.10e-124 | | | 1.34e-114 | |
| | 2.80e-54 | 2.21e-55 | | 9.69e-98 | | | 4.56e-63 | |
| | 7.54e-73 | 2.97e-57 | | 2.47e-97 | | | 1.04e-66 | |
| Q72/+;; MHC/+ | 2.28e-55 | 1.21e-20 | | | 6.57e-03 | | | 1.25e-40 |
| | 4.02e-57 | 6.04e-14 | | | 2.20e-01 | | | 1.71e-30 |
| | 1.83e-02 | 1.40e-26 | | | 1.00e+00 | | | 2.41e-52 |
| | 1.00e+00 | 3.03e-15 | | | 7.80e-20 | | | 1.99e-75 |
| Q72/+;; MHC/+ | 1.74e-58 | 3.59e-64 | | | 6.28e-76 | | | 1.04e-60 |
| +RU486 | 1.11e-57 | 1.58e-67 | | | 2.24e-69 | | | 2.61e-69 |

Table 7 Compiled Logrank analysis of longevity with multiple copies of Q72 and muscle driver transgenes (related to Figure 3, S4). Each p-value is from one experimental replicate

| MHC Males +RU486 (50μg/ml) | | | | | MHC Males -RU486 (0μg/ml) | | | |
|---|---|---|---|---|---|---|---|---|
| | Q72;;MHC | Q72;;MHC/+ | Q72/+;;MHC | | | Q72;;MHC | Q72;;MHC/+ | Q72/+;;MHC |
| Q72; MHC/+ | 6.09e-29 | | | | Q72; MHC/+ | 8.15e-09 | | |
| | 2.05e-16 | | | | | 2.28e-09 | | |
| | 1.35e-36 | | | | | 7.68e-08 | | |
| | 1.73e-13 | | | | | 1.83e-05 | | |
| | 4.66e-11 | | | | | 1.21e-09 | | |
| | 6.37e-16 | | | | | 9.74e-10 | | |
| Q72/+; MHC | 4.85e-01 | 9.47e-30 | | | Q72/+; MHC | 3.75e-03 | 4.17e-22 | |
| | 1.00e+00 | 7.49e-11 | | | | 1.00e+00 | 7.26e-07 | |
| | 2.17e-03 | 8.70e-21 | | | | 1.00e+00 | 4.34e-07 | |
| | 1.00e+00 | 1.10e-13 | | | | 3.22e-01 | 1.09e-08 | |
| | 1.56e-01 | 4.29e-17 | | | | 7.08e-02 | 2.22e-12 | |
| | 6.72e-06 | 3.22e-23 | | | | 3.60e-01 | 9.58e-13 | |
| Q72/+; MHC/+ | 9.02e-02 | 3.32e-41 | 1.00e+00 | | Q72/+; MHC/+ | 3.93e-12 | 1.00e+00 | 3.60e-26 |
| | 7.37e-06 | 2.36e-32 | 2.28e-08 | | | 1.70e-15 | 3.18e-01 | 2.34e-13 |
| | 6.32e-07 | 4.29e-17 | 1.00e+00 | | | 3.63e-24 | 1.06e-08 | 1.48e-21 |
| | 1.00e+00 | 5.15e-12 | 1.00e+00 | | | 3.33e-30 | 3.69e-07 | 9.82e-37 |
| | 1.81e-01 | 3.22e-06 | 6.57e-05 | | | 5.65e-21 | 1.00e+00 | 2.28e-28 |
| | 4.52e-01 | 1.87e-14 | 1.49e-08 | | | 1.51e-23 | 1.11e-05 | 5.25e-27 |

| MHC Females +RU486 (50μg/ml) | | | | | MHC Females -RU486 (0μg/ml) | | | |
|---|---|---|---|---|---|---|---|---|
| | Q72;;MHC | Q72;;MHC/+ | Q72/+;;MHC | | | Q72;;MHC | Q72;;MHC/+ | Q72/+;;MHC |
| Q72; MHC/+ | 7.65e-23 | | | | Q72; MHC/+ | 1.17e-01 | | |
| | 1.77e-30 | | | | | 1.00e+00 | | |
| | 4.72e-34 | | | | | 6.96e-04 | | |
| | 2.51e-24 | | | | | 1.01e-08 | | |
| | 1.31e-22 | | | | | 1.00e+00 | | |
| | 8.68e-29 | | | | | 4.57e-02 | | |
| Q72/+; MHC | 5.63e-01 | 2.56e-23 | | | Q72/+; MHC | 3.97e-01 | 3.77e-03 | |
| | 1.19e-01 | 1.54e-35 | | | | 5.41e-02 | 1.00e+00 | |
| | 1.00e+00 | 1.28e-31 | | | | 5.78e-22 | 7.31e-04 | |
| | 1.00e+00 | 3.33e-23 | | | | 7.71e-36 | 7.40e-01 | |
| | 2.50e-01 | 2.15e-31 | | | | 3.62e-09 | 1.23e-04 | |
| | 7.63e-11 | 1.58e-48 | | | | 5.51e-19 | 2.60e-18 | |
| Q72/+; MHC/+ | 2.32e-08 | 4.55e-06 | 2.80e-10 | | Q72/+; MHC/+ | 1.30e-05 | 1.00e+00 | 5.94e-07 |
| | 1.39e-06 | 1.52e-18 | 3.27e-15 | | | 2.46e-18 | 1.19e-11 | 5.65e-08 |

| | | | |
|---|---|---|---|
| | 1.00e+00 | 8.95e-35 | 1.00e+00 |
| | 5.16e-06 | 4.75e-19 | 2.24e-04 |
| | 1.00e+00 | 2.07e-25 | 1.26e-02 |
| | 3.51e-05 | 1.57e-20 | 2.35e-33 |

| | | | |
|---|---|---|---|
| | 1.80e-19 | 1.70e-09 | 1.17e-01 |
| | 2.99e-42 | 3.21e-24 | 9.86e-27 |
| | 1.00e+00 | 1.00e+00 | 1.37e-01 |
| | 2.33e-08 | 5.02e-01 | 2.74e-26 |

DJ694 Males

| | Q72;DJ694 | Q72;DJ694/+ | Q72/+;DJ694 |
|---|---|---|---|
| Q72; DJ694/+ | 2.67e-63 | | |
| | 1.07e-64 | | |
| | 3.39e-55 | | |
| | 2.42e-31 | | |
| | 3.36e-67 | | |
| | 3.27e-67 | | |
| Q72/+; DJ694 | 7.74e-03 | 7.94e-61 | |
| | 1.25e-06 | 3.18e-66 | |
| | 8.05e-14 | 3.09e-57 | |
| | 2.51e-07 | 1.11e-31 | |
| | 2.36e-07 | 1.17e-57 | |
| | 7.59e-05 | 9.87e-57 | |
| Q72/+; DJ694/+ | 3.80e-53 | 1.31e-35 | 8.24e-53 |
| | 3.58e-45 | 4.74e-45 | 5.73e-57 |
| | 5.43e-21 | 7.94e-05 | 1.88e-29 |
| | 1.95e-38 | 1.53e-02 | 3.22e-37 |
| | 5.97e-73 | 2.30e-16 | 7.04e-58 |
| | 7.10e-69 | 3.51e-08 | 3.17e-57 |

DJ694 Females

| | Q72;DJ694 | Q72;DJ694/+ | Q72/+;DJ694 |
|---|---|---|---|
| Q72; DJ694/+ | 5.49e-58 | | |
| | 1.45e-52 | | |
| | 2.41e-63 | | |
| | 1.89e-36 | | |
| | 5.35e-73 | | |
| | 5.03e-60 | | |
| Q72/+; DJ694 | 5.09e-01 | 6.51e-50 | |
| | 1.44e-08 | 2.03e-44 | |
| | 2.10e-02 | 3.49e-59 | |
| | 4.15e-03 | 2.63e-29 | |
| | 1.00e+00 | 3.46e-55 | |
| | 3.62e-04 | 6.63e-34 | |
| Q72/+; DJ694/+ | 6.45e-54 | 1.74e-14 | 6.22e-36 |
| | 9.30e-34 | 5.13e-37 | 1.27e-07 |
| | 3.17e-38 | 3.75e-06 | 1.60e-31 |
| | 2.14e-40 | 1.00e+00 | 2.16e-26 |
| | 9.87e-62 | 6.15e-11 | 7.55e-45 |
| | 8.22e-60 | 3.49e-10 | 1.25e-18 |

Table 8 One-way ANOVA of CPRG expression level measurements of genotypes with multiple copies of Q72 and muscle driver transgenes (related to Figure 3)

| | | | |
|---|---|---|---|
| Males -RU486 (0μg/ml) | | | |
| one- way ANOVA | | | |
| F | p value | R squared | |
| 28.85 | <0.0001 | 0.7062 | |
| Dunnett's multiple comparisions test | | | |
| | | | p value |
| Bg2/+;Mef2/+ | vs | Bg2/+;MHC/+ | <0.0001 |
| | | Bg2;MHC | <0.0001 |
| | | DJ694,Bg2 | 0.0039 |
| Females -RU486 (0μg/ml) | | | |
| one- way ANOVA | | | |
| F | p value | R squared | |
| 31.25 | <0.0001 | 0.7226 | |
| Dunnett's multiple comparisions test | | | |
| | | | p value |
| Bg2/+;Mef2/+ | vs | Bg2/+;MHC/+ | <0.0001 |
| | | Bg2;MHC | 0.0012 |
| | | DJ694,Bg2 | 0.1409 |
| Males +RU486 (50μg/ml) | | | |
| one- way ANOVA | | | |
| F | p value | R squared | |
| 15.86 | <0.0001 | 0.5693 | |
| Dunnett's multiple comparisions test | | | |
| | | | p value |
| Bg2/+;Mef2/+ | vs | Bg2/+;MHC/+ | 0.0696 |
| | | Bg2;MHC | 0.0028 |
| | | DJ694,Bg2 | 0.0512 |
| Females +RU486 (50μg/ml) | | | |
| one- way ANOVA | | | |
| F | p value | R squared | |
| 16.67 | <0.0001 | 0.5815 | |
| Dunnett's multiple comparisions test | | | |
| | | | p value |
| Bg2/+;Mef2/+ | vs | Bg2/+;MHC/+ | <0.0001 |
| | | Bg2;MHC | 0.1899 |
| | | DJ694,Bg2 | 0.0593 |

Table 9 Compiled Logrank analysis of temporal repression of Q72 using GAL80ts longevity (related to Figure 5). Each p-value is from one experimental replicate

| Males 20-20°C | | | | |
|---|---|---|---|---|
| | Q72/+;;Mef2/+ | Mef2/+ | Q72/+ | Q72/+;;GAL80ts/+ |
| Q72/+;;Mef2/ GAL80ts | 1.01e-57 | 2.44e-27 | 3.55e-18 | 1.04e-24 |
| | 3.11e-49 | 1.56e-24 | 5.72e-15 | 7.39e-12 |
| | 1.77e-32 | 1.29e-13 | 7.50e-09 | 2.78e-11 |
| | 8.46e-46 | 2.73e-02 | 1.00e+00 | 1.00e+00 |
| | 9.69e-46 | 2.52e-10 | 3.75e-02 | 4.24e-01 |
| | 7.44e-34 | 1.00e+00 | 4.79e-01 | 3.98e-01 |
| Q72/+;;Mef2/+ | | 6.33e-50 | 2.16e-43 | 2.22e-63 |
| | | 3.73e-37 | 3.26e-38 | 3.50e-41 |
| | | 6.65e-39 | 5.81e-27 | 1.77e-32 |
| | | 3.13e-40 | 3.04e-33 | 8.46e-46 |
| | | 5.34e-34 | 3.65e-37 | 9.69e-46 |
| | | 3.36e-28 | 2.26e-42 | 7.44e-34 |
| Females 20-20°C | | | | |
| | Q72/+;;Mef2/+ | Mef2/+ | Q72/+ | Q72/+;;GAL80ts/+ |
| Q72/+;;Mef2/ GAL80ts | 6.02e-81 | 3.96e-11 | 3.85e-39 | 8.32e-13 |
| | 1.72e-91 | 1.27e-16 | 1.57e-12 | 9.92e-01 |
| | 1.06e-49 | 1.08e-10 | 2.00e-13 | 8.96e-07 |
| | 6.87e-60 | 8.29e-08 | 1.34e-14 | 2.80e-03 |
| | 4.02e-56 | 9.68e-27 | 3.74e-28 | 2.46e-19 |
| | 1.59e-33 | 7.67e-17 | 1.06e-18 | 1.65e-10 |
| Q72/+;;Mef2/+ | | 1.65e-62 | 9.50e-56 | 5.21e-71 |
| | | 4.20e-55 | 3.48e-70 | 1.49e-79 |
| | | 6.59e-34 | 3.50e-60 | 1.06e-49 |
| | | 4.91e-64 | 1.39e-52 | 6.87e-60 |
| | | 5.46e-56 | 2.01e-48 | 4.02e-56 |
| | | 1.43e-03 | 4.05e-51 | 1.59e-33 |
| Males 20-29°C | | | | |
| | Q72/+;;Mef2/+ | Mef2/+ | Q72/+ | Q72/+;;GAL80ts/+ |
| Q72/+;;Mef2/ GAL80ts | 3.61e-38 | 6.49e-08 | 8.83e-13 | 4.69e-16 |
| | 7.35e-45 | 7.43e-05 | 8.78e-21 | 8.61e-20 |
| | 1.53e-41 | 3.63e-01 | 1.88e-10 | 3.89e-15 |
| | 2.87e-31 | 1.00e+00 | 5.76e-09 | 2.76e-04 |
| Q72/+;;Mef2/+ | | 5.88e-50 | 1.90e-37 | 8.53e-50 |
| | | 2.46e-41 | 1.01e-33 | 8.57e-45 |
| | | 1.02e-25 | 1.71e-36 | 1.53e-41 |
| | | 5.42e-18 | 5.06e-26 | 2.87e-31 |
| Females 20-29°C | | | | |
| | Q72/+;;Mef2/+ | Mef2/+ | Q72/+ | Q72/+;;GAL80ts/+ |

| | | | | |
|---|---|---|---|---|
| Q72/+;;Mef2/ GAL80ts | 2.59e-55 | 1.95e-03 | 1.55e-08 | 1.00e+00 |
| | 1.01e-52 | 3.33e-06 | 6.39e-18 | 1.22e-03 |
| | 1.90e-58 | 9.34e-38 | 5.57e-34 | 7.46e-45 |
| | 3.47e-28 | 9.85e-06 | 1.85e-21 | 3.02e-30 |
| Q72/+;;Mef2/+ | | 1.73e-54 | 1.38e-49 | 7.37e-59 |
| | | 1.69e-54 | 4.17e-46 | 4.74e-58 |
| | | 2.58e-53 | 1.20e-56 | 1.90e-58 |
| | | 5.64e-04 | 1.87e-64 | 3.47e-28 |
| Males 29-20°C | | | | |
| | Q72/+;;Mef2/+ | Mef2/+ | Q72/+ | Q72/+;;GAL80ts/+ |
| Q72/+;;Mef2/ GAL80ts | 8.23e-63 | 2.33e-11 | 7.52e-01 | 1.38e-01 |
| | 2.63e-77 | 2.78e-05 | 9.77e-01 | 4.92e-03 |
| | 2.20e-44 | 3.99e-01 | 2.48e-08 | 2.88e-22 |
| | 1.16e-63 | 9.16e-23 | 3.06e-26 | 3.18e-41 |
| Q72/+;;Mef2/+ | | 3.54e-65 | 1.08e-67 | 4.91e-66 |
| | | 2.46e-78 | 2.91e-67 | 2.49e-62 |
| | | 1.85e-44 | 1.13e-52 | 2.20e-44 |
| | | 2.29e-62 | 2.79e-63 | 1.16e-63 |
| Females 29-20°C | | | | |
| | Q72/+;;Mef2/+ | Mef2/+ | Q72/+ | Q72/+;;GAL80ts/+ |
| Q72/+;;Mef2/ GAL80ts | 3.20e-80 | 7.28e-19 | 1.09e-25 | 2.37e-12 |
| | 5.63e-80 | 2.43e-04 | 1.65e-50 | 5.86e-07 |
| | 3.56e-55 | 1.00e+00 | 3.49e-01 | 4.84e-02 |
| | 1.19e-68 | 1.10e-12 | 1.00e+00 | 3.47e-12 |
| Q72/+;;Mef2/+ | | 8.71e-89 | 5.91e-78 | 8.42e-73 |
| | | 3.26e-75 | 3.76e-78 | 1.32e-68 |
| | | 5.87e-24 | 7.34e-65 | 3.56e-55 |
| | | 7.33e-60 | 3.55e-83 | 1.19e-68 |
| Males 29-29°C | | | | |
| | Q72/+;;Mef2/+ | Mef2/+ | Q72/+ | Q72/+;;GAL80ts/+ |
| Q72/+;;Mef2/ GAL80ts | 5.72e-72 | 6.38e-72 | 2.35e-77 | 3.23e-67 |
| | 2.37e-80 | 2.69e-72 | 3.42e-63 | 4.07e-68 |
| | 1.40e-48 | 4.18e-24 | 4.89e-34 | 3.79e-42 |
| | 1.57e-55 | 7.43e-34 | 2.00e-49 | 1.84e-54 |
| Q72/+;;Mef2/+ | | 1.99e-73 | 6.48e-70 | 1.49e-64 |
| | | 2.21e-66 | 3.41e-60 | 8.32e-64 |
| | | 3.72e-51 | 1.70e-57 | 1.40e-48 |
| | | 3.42e-50 | 5.43e-57 | 1.57e-55 |
| Females 29-29°C | | | | |
| | Q72/+;;Mef2/+ | Mef2/+ | Q72/+ | Q72/+;;GAL80ts/+ |
| Q72/+;;Mef2/ GAL80ts | 9.12e-76 | 2.38e-83 | 6.12e-69 | 2.46e-79 |

| | | | | |
|---|---|---|---|---|
| | 4.72e-79 | 6.05e-77 | 2.73e-50 | 7.16e-70 |
| | 2.23e-59 | 3.03e-34 | 9.80e-59 | 6.87e-50 |
| | 7.78e-61 | 3.40e-60 | 3.32e-71 | 3.45e-58 |
| Q72/+;;Mef2/+ | | 2.06e-82 | 4.71e-74 | 7.09e-79 |
| | | 2.24e-74 | 1.16e-68 | 9.78e-71 |
| | | 4.52e-47 | 2.63e-67 | 2.23e-59 |
| | | 1.81e-58 | 2.34e-70 | 7.78e-61 |

Table 10 Compiled Logrank analysis of Hsp70 expression in polyQ flies longevity (related to Figure 6,S7). Each p-value is from one experimental replicate

| MHC Males +RU486 (50$\mu$g/ml) | | | | | | |
|---|---|---|---|---|---|---|
| | Q72/+;;MHC/+ | Q72/+;;MHC/Bg3 | Q72/+ | MHC/+ | UAS-hsp70$^{4.2}$/+ | UAS-hsp70$^{4.4}$/+ |
| Q72/+;;MHC/ UAS-hsp70$^{4.2}$ | 0.14 | 1 | 2.47e-47 | 1.55e-44 | 4.64e-46 | |
| | 1 | 1 | 9.56e-57 | 6.29e-51 | 2.25e-62 | |
| Q72/+;;MHC/ UAS-hsp70$^{4.4}$ | 8.86e-35 | 1.03e-31 | 4.51e-48 | 9.11e-43 | | 2.20e-46 |
| | 4.32e-52 | 2.71e-52 | 2.69e-52 | 1.41e-56 | | 4.91e-57 |
| MHC/UAS-hsp70$^{4.2}$ | 4.52e-32 | 6.06e-36 | 5.36e-05 | 1 | 5.34e-23 | |
| | 4.55e-59 | 3.01e-62 | 1 | 1.20e-08 | 9.52e-02 | |
| MHC/UAS-hsp70$^{4.4}$ | 2.15e-16 | 2.57e-22 | 9.12e-44 | 1.40e-37 | | 1.08e-17 |
| | 8.91e-39 | 1.56e-40 | 9.27e-54 | 3.96e-24 | | 6.52e-17 |
| MHC Females +RU486 (50$\mu$g/ml) | | | | | | |
| | Q72/+;;MHC/+ | Q72/+;;MHC/Bg3 | Q72/+ | MHC/+ | UAS-hsp70$^{4.2}$/+ | UAS-hsp70$^{4.4}$/+ |
| Q72/+;;MHC/ UAS-hsp70$^{4.2}$ | 0.0835 | 0.0029 | 1.35e-20 | 1.68e-47 | 6.06e-48 | |
| | 2.44e-14 | 8.89e-05 | 1.05e-51 | 6.42e-58 | 2.75e-62 | |
| Q72/+;;MHC/ UAS-hsp70$^{4.4}$ | 1.27e-30 | 4.19e-22 | 3.42e-21 | 6.67e-47 | | 6.35e-37 |
| | 2.82e-54 | 8.19e-38 | 1.31e-53 | 6.86e-55 | | 1.68e-55 |
| MHC/UAS-hsp70$^{4.2}$ | 1.25e-55 | 2.43e-54 | 1.76e-08 | 1 | 0.0022 | |
| | 3.50e-60 | 2.36e-61 | 7.45e-26 | 0.31 | 1 | |
| MHC/UAS-hsp70$^{4.4}$ | 7.37e-29 | 4.13e-36 | 1.05e-11 | 4.87e-36 | | 1 |
| | 9.20e-36 | 5.43e-50 | 3.49e-06 | 3.05e-38 | | 1 |
| Mef2 Males | | | | | | |
| | Q72/+;;Mef2/+ | Q72/+;;Mef2/Bg3 | Q72/+ | Mef2/+ | UAS-hsp70$^{4.2}$/+ | UAS-hsp70$^{4.4}$/+ |
| Q72/+;;Mef2/ UAS-hsp70$^{4.2}$ | 1.82e-35 | 0.0006 | 9.72e-46 | 5.11e-55 | 2.70e-54 | |
| | 9.71e-45 | 3.52e-08 | 2.43e-83 | 6.80e-61 | 5.43e-77 | |
| Q72/+;;Mef2/ UAS-hsp70$^{4.4}$ | 2.46e-09 | 2.75e-37 | 5.52e-46 | 1.73e-52 | | 2.38e-46 |
| | 1.52e-23 | 6.35e-74 | 3.39e-88 | 6.59e-70 | | 8.57e-71 |
| Mef2/UAS-hsp70$^{4.2}$ | 6.26e-37 | 4.32e-42 | 1 | 0.0125 | 4.06e-05 | |

| | | | | | | |
|---|---|---|---|---|---|---|
| | 1.54e-68 | 8.36e-77 | 9.11e-13 | 1 | 1 | |
| Mef2/UAS-hsp70$^{4.4}$ | 1.67e-54 | 4.77e-56 | 5.74e-28 | 3.65e-45 | | 1.10e-09 |
| | 1.04e-70 | 5.60e-79 | 4.07e-26 | 3.50e-49 | | 1 |
| Mef2 Females | | | | | | |
| | Q72/+;;Mef2/+ | Q72/+;;Mef2/Bg3 | Q72/+ | Mef2/+ | UAS-hsp70$^{4.2}$/+ | UAS-hsp70$^{4.4}$/+ |
| Q72/+;;Mef2/UAS-hsp70$^{4.2}$ | 1.37e-06 | 0.0038 | 1.10e-27 | 2.81e-38 | 1.21e-67 | |
| | 0.19 | 1 | 1.26e-75 | 1.20e-59 | 2.50e-78 | |
| Q72/+;;Mef2/UAS-hsp70$^{4.4}$ | 8.57e-30 | 1.13e-29 | 8.74e-27 | 2.46e-36 | | 1.88e-48 |
| | 3.49e-24 | 6.75e-57 | 3.57e-76 | 3.34e-61 | | 7.73e-76 |
| Mef2/UAS-hsp70$^{4.2}$ | 2.01e-56 | 2.46e-57 | 5.43e-39 | 7.37e-24 | 3.21e-18 | |
| | 1.87e-67 | 7.60e-72 | 4.02e-59 | 2.47e-17 | 1.99e-27 | |
| Mef2/UAS-hsp70$^{4.4}$ | 2.10e-53 | 7.51e-54 | 3.66e-06 | 8.18e-33 | | 1 |
| | 2.09e-78 | 6.97e-82 | 1.56e-04 | 2.23e-50 | | 2.03e-08 |

```
#writing the survival data df
library (openxlsx)
library(survival)
library(ggplot2)
library(survminer)
datafile <- file.choose()
df <- read.xlsx(datafile, sheet=16, rows = c(5:203))
lastcol<-match("Sample.size",names(df))
df <- read.xlsx(datafile, sheet=16, rows = c(5:203), cols = c(1:lastcol))
survivaldata<-data.frame(matrix(ncol=6,nrow=sum(df$Sample.size)))
names(survivaldata)[1]<-"Time"
names(survivaldata)[2]<-"Dead"
names(survivaldata)[3]<-"Genotype"
names(survivaldata)[4]<-"Sex"
names(survivaldata)[5]<-"Vial"
names(survivaldata)[6]<-"Treatment"
lastday <- match ("Sample.size",names(df)) -1
loop <- dim(df)[1]
scoringdays <- colnames(df)

writeindex <- 1

for (j in 1:loop){
  deathcount <- 0

  for (day in 6:lastday){death <- df[j,day]
  dayvalue=scoringdays[day]
  deathcount <- deathcount + as.numeric(death)

  if (death>0){
    for (i in 1:death){
      survivaldata[writeindex,1] <- as.numeric(dayvalue)
      survivaldata[writeindex,2] <- 1
      survivaldata[writeindex,3] <- df[j,1]
      survivaldata[writeindex,4] <- df[j,2]
      survivaldata[writeindex,5] <- df[j,4]
      survivaldata[writeindex,6] <- df[j,3]
      writeindex <- writeindex+1
    }
  }

  print (deathcount)
  if (deathcount==df[j,lastday+1]){break}
  }

}
genotype.table<-table(survivaldata$Genotype)

#subsetting survival data by sex, treatment
survivaldatam<-subset(survivaldata, Sex=="male")
survivaldataf<-subset(survivaldata, Sex=="female")
survivaldatamRU<-subset(survivaldatam, Treatment==" +RU486")
survivaldatamnoRU<-subset(survivaldatam, Treatment==" -RU486")
```

```
survivaldatamnone<-subset(survivaldatam, Treatment=="none")
survivaldatafRU<-subset(survivaldataf, Treatment==" +RU486")
survivaldatafnoRU<-subset(survivaldataf, Treatment==" -RU486")
survivaldatafnone<-subset(survivaldataf, Treatment=="none")
#making a list of data frames by genotype
survivaldatamgRU<-split(survivaldatamRU, f=survivaldatamRU$Genotype)
survivaldatamgnoRU<-split(survivaldatamnoRU, f=survivaldatamnoRU$Genotype)
survivaldatamgnone<-split(survivaldatamnone, f=survivaldatamnone$Genotype)
survivaldatafgRU<-split(survivaldatafRU, f=survivaldatafRU$Genotype)
survivaldatafgnoRU<-split(survivaldatafnoRU, f=survivaldatafnoRU$Genotype)
survivaldatafgnone<-split(survivaldatafnone, f=survivaldatafnone$Genotype)
#running fit by vial
fitmRU<-surv_fit(Surv(Time, Dead)~Vial, data=survivaldatamgRU)
fitmnoRU<-surv_fit(Surv(Time, Dead)~Vial, data=survivaldatamgnoRU)
fitmnone<-surv_fit(Surv(Time, Dead)~Vial, data=survivaldatamgnone)
fitfRU<-surv_fit(Surv(Time, Dead)~Vial, data=survivaldatafgRU)
fitfnoRU<-surv_fit(Surv(Time, Dead)~Vial, data=survivaldatafgnoRU)
fitfnone<-surv_fit(Surv(Time, Dead)~Vial, data=survivaldatafgnone)
#calculating median survival by vial
medmRU<-surv_median(fitmRU)
medmnoRU<-surv_median(fitmnoRU)
medmnone<-surv_median(fitmnone)
medfRU<-surv_median(fitfRU)
medfnoRU<-surv_median(fitfnoRU)
medfnone<-surv_median(fitfnone)
#running fit of averaged vials
fitmRU2<-surv_fit(Surv(Time, Dead)~Genotype, data=survivaldatamgRU)
fitmnoRU2<-surv_fit(Surv(Time, Dead)~Genotype, data=survivaldatamgnoRU)
fitmnone2<-surv_fit(Surv(Time, Dead)~Genotype, data=survivaldatamgnone)
fitfRU2<-surv_fit(Surv(Time, Dead)~Genotype, data=survivaldatafgRU)
fitfnoRU2<-surv_fit(Surv(Time, Dead)~Genotype, data=survivaldatafgnoRU)
fitfnone2<-surv_fit(Surv(Time, Dead)~Genotype, data=survivaldatafgnone)
#finding medians of averaged vials
medmRU2<-surv_median(fitmRU2)
medmnoRU2<-surv_median(fitmnoRU2)
medmnone2<-surv_median(fitmnone2)
medfRU2<-surv_median(fitfRU2)
medfnoRU2<-surv_median(fitfnoRU2)
medfnone2<-surv_median(fitfnone2)

#writing Compilation Table (by vial)
library(openxlsx)
library(readxl)
datafile <- file.choose()
df <- read.xlsx(datafile, sheet=1, rows = c(5:203))
df2 <- read_excel(datafile, sheet=16,range = cell_rows(207:608))
df2<-as.data.frame(df2)
rowwithdays<-colnames(df2)
loop2<-dim(df2)[1];
lastcol<-dim(df2)[2]
CompilationTable<-data.frame(matrix(nrow = loop2, ncol = 8))
names(CompilationTable)[1]<-"Genotype"
names(CompilationTable)[2]<-"Sex"
names(CompilationTable)[3]<-"Treatment"
```

```
names(CompilationTable)[4]<-"Vial"
names(CompilationTable)[5]<-"Sample size"
names(CompilationTable)[6]<-"Average lifespan"
names(CompilationTable)[7]<-"Median lifespan"
names(CompilationTable)[8]<-"Maximum lifespan"
nacounter<-0
for(counter in 1:loop2){
  value<-df2[counter,1]
  if (is.na(value)==TRUE){nacounter<-nacounter+1}
  else{
    if (as.character(value)=="Processed data"){break}
    else{
      CompilationTable[counter-nacounter,1]<-df2[counter,1]
      CompilationTable[counter-nacounter,2]<-df2[counter,2]
      CompilationTable[counter-nacounter,3]<-df2[counter,3]
      CompilationTable[counter-nacounter,4]<-df2[counter,4]
      CompilationTable[counter-nacounter,5]<-df[counter,lastcol]
      CompilationTable[counter-nacounter,6]<-df2[counter,lastcol]

      for (counter2 in 5:lastcol){
        value2<-as.numeric(df2[counter,counter2])
        if (value2<=5){
          maxls<-as.numeric(rowwithdays[counter2])
          CompilationTable[counter-nacounter,8]<-maxls
          break
        }
      }
    }
  }
}

#finding the position in Compilation Table to write the medians
loopcount<-0
for (g in names(survivaldatamgRU)){
  loopcount<-loopcount+1
  genpos<-which(CompilationTable==g, arr.in=TRUE)
  malecount<-vector()
  treat<-vector()
  for (h in 1:dim(genpos)[1]){
    if (CompilationTable[genpos[h],2]=="male"){
      malecount<-c(malecount,genpos[h])}

  }
  for (k in 1:length(malecount)){
    if (CompilationTable[malecount[k],3]=="+RU486"){
      treat<-c(treat,malecount[k])
    }
  }
  for (m in 1:length(treat)){
    CompilationTable[treat[m],7]<-medmRU[[loopcount]][m,2]
  }

}
loopcount<-0
```

```
for (g in names(survivaldatamgnoRU)){
  loopcount<-loopcount+1
  genpos<-which(CompilationTable==g, arr.in=TRUE)
  malecount<-vector()
  treat<-vector()
  for (h in 1:dim(genpos)[1]){
    if (CompilationTable[genpos[h],2]=="male"){
      malecount<-c(malecount,genpos[h])}

  }
  for (k in 1:length(malecount)){
    if (CompilationTable[malecount[k],3]=="-RU486"){
      treat<-c(treat,malecount[k])
    }
  }
  for (m in 1:length(treat)){
    CompilationTable[treat[m],7]<-medmnoRU[[loopcount]][m,2]
  }

}
loopcount<-0
for (g in names(survivaldatamgnone)){
  loopcount<-loopcount+1
  genpos<-which(CompilationTable==g, arr.in=TRUE)
  malecount<-vector()
  treat<-vector()
  for (h in 1:dim(genpos)[1]){
    if (CompilationTable[genpos[h],2]=="male"){
      malecount<-c(malecount,genpos[h])}

  }
  for (k in 1:length(malecount)){
    if (CompilationTable[malecount[k],3]=="none"){
      treat<-c(treat,malecount[k])
    }
  }
  for (m in 1:length(treat)){
    CompilationTable[treat[m],7]<-medmnone[[loopcount]][m,2]
  }

}
loopcount<-0
for (g in names(survivaldatafgRU)){
  loopcount<-loopcount+1
  genpos<-which(CompilationTable==g, arr.in=TRUE)
  malecount<-vector()
  treat<-vector()
  for (h in 1:dim(genpos)[1]){
    if (CompilationTable[genpos[h],2]=="female"){
      malecount<-c(malecount,genpos[h])}

  }
  for (k in 1:length(malecount)){
    if (CompilationTable[malecount[k],3]=="+RU486"){
```

```
      treat<-c(treat,malecount[k])
    }
  }
  for (m in 1:length(treat)){
    CompilationTable[treat[m],7]<-medfRU[[loopcount]][m,2]
  }

}
loopcount<-0
for (g in names(survivaldatafgnoRU)){
  loopcount<-loopcount+1
  genpos<-which(CompilationTable==g, arr.in=TRUE)
  malecount<-vector()
  treat<-vector()
  for (h in 1:dim(genpos)[1]){
    if (CompilationTable[genpos[h],2]=="female"){
      malecount<-c(malecount,genpos[h])}

  }
  for (k in 1:length(malecount)){
    if (CompilationTable[malecount[k],3]=="-RU486"){
      treat<-c(treat,malecount[k])
    }
  }
  for (m in 1:length(treat)){
    CompilationTable[treat[m],7]<-medfnoRU[[loopcount]][m,2]
  }

}
loopcount<-0
for (g in names(survivaldatafgnone)){
  loopcount<-loopcount+1
  genpos<-which(CompilationTable==g, arr.in=TRUE)
  malecount<-vector()
  treat<-vector()
  for (h in 1:dim(genpos)[1]){
    if (CompilationTable[genpos[h],2]=="female"){
      malecount<-c(malecount,genpos[h])}

  }
  for (k in 1:length(malecount)){
    if (CompilationTable[malecount[k],3]=="none"){
      treat<-c(treat,malecount[k])
    }
  }
  for (m in 1:length(treat)){
    CompilationTable[treat[m],7]<-medfnone[[loopcount]][m,2]
  }

}

#write Compilation table of averaged vials
#subsetting CompilationTable (by vial) by sex, treatment
CompTablem<-subset(CompilationTable, Sex=="male")
```

```
CompTablemRU<-subset(CompTablem, Treatment=="+RU486")
CompTablemnoRU<-subset(CompTablem, Treatment=="-RU486")
CompTablemnone<-subset(CompTablem, Treatment=="none")
CompTablef<-subset(CompilationTable, Sex=="female")
CompTablefRU<-subset(CompTablef, Treatment=="+RU486")
CompTablefnoRU<-subset(CompTablef, Treatment=="-RU486")
CompTablefnone<-subset(CompTablef, Treatment=="none")

#CompilationTableavg is written in 6 separate tables and then bound together
at the end
CompilationTablemRU<-data.frame(matrix(nrow =length(survivaldatamgRU), ncol =
7))
names(CompilationTablemRU)[1]<-"Genotype"
names(CompilationTablemRU)[2]<-"Sex"
names(CompilationTablemRU)[3]<-"Treatment"
names(CompilationTablemRU)[4]<-"Sample size"
names(CompilationTablemRU)[5]<-"Average lifespan"
names(CompilationTablemRU)[6]<-"Median lifespan"
names(CompilationTablemRU)[7]<-"Maximum lifespan"

for (z in 1:length(survivaldatamgRU)){
  CompilationTablemRU[z,1]<-names(survivaldatamgRU)[z]
  CompilationTablemRU[z,2]<-"male"
  CompilationTablemRU[z,3]<-survivaldatamgRU[[z]][1,6]
  CompilationTablemRU[z,4]<-sum(survivaldatamgRU[[z]]$Dead)
  CompilationTablemRU[z,6]<-medmRU2[[z]][1,2]

  y<-names(survivaldatamgRU)[z]
  genpos2<-which(CompTablemRU==y, arr.in=TRUE)
  average1<-as.numeric(CompTablemRU[genpos2[1:dim(genpos2)[1]],6])
  max1<-as.numeric(CompTablemRU[genpos2[1:dim(genpos2)[1]],8])
  averagels<-mean(average1)
  averagemax<-mean(max1)
  print(averagels)
  print(averagemax)
  CompilationTablemRU[z,5]<-averagels
  CompilationTablemRU[z,7]<-averagemax

}

CompilationTablemnoRU<-data.frame(matrix(nrow =length(survivaldatamgnoRU),
ncol = 7))
names(CompilationTablemnoRU)[1]<-"Genotype"
names(CompilationTablemnoRU)[2]<-"Sex"
names(CompilationTablemnoRU)[3]<-"Treatment"
names(CompilationTablemnoRU)[4]<-"Sample size"
names(CompilationTablemnoRU)[5]<-"Average lifespan"
names(CompilationTablemnoRU)[6]<-"Median lifespan"
names(CompilationTablemnoRU)[7]<-"Maximum lifespan"

for (z in 1:length(survivaldatamgnoRU)){
  CompilationTablemnoRU[z,1]<-names(survivaldatamgnoRU)[z]
  CompilationTablemnoRU[z,2]<-"male"
  CompilationTablemnoRU[z,3]<-survivaldatamgnoRU[[z]][1,6]
```

```
  CompilationTablemnoRU[z,4]<-sum(survivaldatamgnoRU[[z]]$Dead)
  CompilationTablemnoRU[z,6]<-medmnoRU2[[z]][1,2]

  y<-names(survivaldatamgnoRU)[z]
  genpos2<-which(CompTablemnoRU==y, arr.in=TRUE)
  average1<-as.numeric(CompTablemnoRU[genpos2[1:dim(genpos2)[1]],6])
  max1<-as.numeric(CompTablemnoRU[genpos2[1:dim(genpos2)[1]],8])
  averagels<-mean(average1)
  averagemax<-mean(max1)
  print(averagels)
  print(averagemax)
  CompilationTablemnoRU[z,5]<-averagels
  CompilationTablemnoRU[z,7]<-averagemax

}

CompilationTablemnone<-data.frame(matrix(nrow =length(survivaldatamgnone),
ncol = 7))
names(CompilationTablemnone)[1]<-"Genotype"
names(CompilationTablemnone)[2]<-"Sex"
names(CompilationTablemnone)[3]<-"Treatment"
names(CompilationTablemnone)[4]<-"Sample size"
names(CompilationTablemnone)[5]<-"Average lifespan"
names(CompilationTablemnone)[6]<-"Median lifespan"
names(CompilationTablemnone)[7]<-"Maximum lifespan"

for (z in 1:length(survivaldatamgnone)){
  CompilationTablemnone[z,1]<-names(survivaldatamgnone)[z]
  CompilationTablemnone[z,2]<-"male"
  CompilationTablemnone[z,3]<-survivaldatamgnone[[z]][1,6]
  CompilationTablemnone[z,4]<-sum(survivaldatamgnone[[z]]$Dead)
  CompilationTablemnone[z,6]<-medmnone2[[z]][1,2]

  y<-names(survivaldatamgnone)[z]
  genpos2<-which(CompTablemnone==y, arr.in=TRUE)
  average1<-as.numeric(CompTablemnone[genpos2[1:dim(genpos2)[1]],6])
  max1<-as.numeric(CompTablemnone[genpos2[1:dim(genpos2)[1]],8])
  averagels<-mean(average1)
  averagemax<-mean(max1)
  print(averagels)
  print(averagemax)
  CompilationTablemnone[z,5]<-averagels
  CompilationTablemnone[z,7]<-averagemax

}

CompilationTablefRU<-data.frame(matrix(nrow =length(survivaldatafgRU), ncol =
7))
names(CompilationTablefRU)[1]<-"Genotype"
names(CompilationTablefRU)[2]<-"Sex"
names(CompilationTablefRU)[3]<-"Treatment"
names(CompilationTablefRU)[4]<-"Sample size"
names(CompilationTablefRU)[5]<-"Average lifespan"
names(CompilationTablefRU)[6]<-"Median lifespan"
```

```
names(CompilationTablefRU)[7]<-"Maximum lifespan"

for (z in 1:length(survivaldatafgRU)){
  CompilationTablefRU[z,1]<-names(survivaldatafgRU)[z]
  CompilationTablefRU[z,2]<-"female"
  CompilationTablefRU[z,3]<-survivaldatafgRU[[z]][1,6]
  CompilationTablefRU[z,4]<-sum(survivaldatafgRU[[z]]$Dead)
  CompilationTablefRU[z,6]<-medfRU2[[z]][1,2]

  y<-names(survivaldatafgRU)[z]
  genpos2<-which(CompTablefRU==y, arr.in=TRUE)
  average1<-as.numeric(CompTablefRU[genpos2[1:dim(genpos2)[1]],6])
  max1<-as.numeric(CompTablefRU[genpos2[1:dim(genpos2)[1]],8])
  averagels<-mean(average1)
  averagemax<-mean(max1)
  print(averagels)
  print(averagemax)
  CompilationTablefRU[z,5]<-averagels
  CompilationTablefRU[z,7]<-averagemax

}

CompilationTablefnoRU<-data.frame(matrix(nrow =length(survivaldatafgnoRU),
ncol = 7))
names(CompilationTablefnoRU)[1]<-"Genotype"
names(CompilationTablefnoRU)[2]<-"Sex"
names(CompilationTablefnoRU)[3]<-"Treatment"
names(CompilationTablefnoRU)[4]<-"Sample size"
names(CompilationTablefnoRU)[5]<-"Average lifespan"
names(CompilationTablefnoRU)[6]<-"Median lifespan"
names(CompilationTablefnoRU)[7]<-"Maximum lifespan"

for (z in 1:length(survivaldatafgnoRU)){
  CompilationTablefnoRU[z,1]<-names(survivaldatafgnoRU)[z]
  CompilationTablefnoRU[z,2]<-"female"
  CompilationTablefnoRU[z,3]<-survivaldatafgnoRU[[z]][1,6]
  CompilationTablefnoRU[z,4]<-sum(survivaldatafgnoRU[[z]]$Dead)
  CompilationTablefnoRU[z,6]<-medfnoRU2[[z]][1,2]

  y<-names(survivaldatafgnoRU)[z]
  genpos2<-which(CompTablefnoRU==y, arr.in=TRUE)
  average1<-as.numeric(CompTablefnoRU[genpos2[1:dim(genpos2)[1]],6])
  max1<-as.numeric(CompTablefnoRU[genpos2[1:dim(genpos2)[1]],8])
  averagels<-mean(average1)
  averagemax<-mean(max1)
  print(averagels)
  print(averagemax)
  CompilationTablefnoRU[z,5]<-averagels
  CompilationTablefnoRU[z,7]<-averagemax

}

CompilationTablefnone<-data.frame(matrix(nrow =length(survivaldatafgnone),
ncol = 7))
```

```
names(CompilationTablefnone)[1]<-"Genotype"
names(CompilationTablefnone)[2]<-"Sex"
names(CompilationTablefnone)[3]<-"Treatment"
names(CompilationTablefnone)[4]<-"Sample size"
names(CompilationTablefnone)[5]<-"Average lifespan"
names(CompilationTablefnone)[6]<-"Median lifespan"
names(CompilationTablefnone)[7]<-"Maximum lifespan"

for (z in 1:length(survivaldatafgnone)){
  CompilationTablefnone[z,1]<-names(survivaldatafgnone)[z]
  CompilationTablefnone[z,2]<-"female"
  CompilationTablefnone[z,3]<-survivaldatafgnone[[z]][1,6]
  CompilationTablefnone[z,4]<-sum(survivaldatafgnone[[z]]$Dead)
  CompilationTablefnone[z,6]<-medfnone2[[z]][1,2]

  y<-names(survivaldatafgnone)[z]
  genpos2<-which(CompTablefnone==y, arr.in=TRUE)
  average1<-as.numeric(CompTablefnone[genpos2[1:dim(genpos2)[1]],6])
  max1<-as.numeric(CompTablefnone[genpos2[1:dim(genpos2)[1]],8])
  averagels<-mean(average1)
  averagemax<-mean(max1)
  print(averagels)
  print(averagemax)
  CompilationTablefnone[z,5]<-averagels
  CompilationTablefnone[z,7]<-averagemax

}
CompilationTableavg<-rbind(CompilationTablemRU, CompilationTablemnoRU,
CompilationTablemnone, CompilationTablefRU, CompilationTablefnoRU,
CompilationTablefnone)
write.xlsx(CompilationTable, file="CompilationTable.xlsx",asTable=TRUE)
write.xlsx(CompilationTableavg, file="CompilationTableavg.xlsx",asTable =
TRUE)

#Plotting graphs, will depend on what you intend to graph

#pairwise comparisons, all p values, will reference to manually add to graphs
in affinity designer
pairedmRU<-pairwise_survdiff(Surv(Time,Dead)~Genotype, data=survivaldatamRU,
p.adjust.method = "bonferroni", rho=0)
pairedmnoRU<-pairwise_survdiff(Surv(Time,Dead)~Genotype,
data=survivaldatamnoRU, p.adjust.method = "bonferroni", rho=0)
pairedmnone<-pairwise_survdiff(Surv(Time,Dead)~Genotype,
data=survivaldatamnone, p.adjust.method = "bonferroni", rho=0)
pairedfRU<-pairwise_survdiff(Surv(Time,Dead)~Genotype, data=survivaldatafRU,
p.adjust.method = "bonferroni", rho=0)
pairedfnoRU<-pairwise_survdiff(Surv(Time,Dead)~Genotype,
data=survivaldatafnoRU, p.adjust.method = "bonferroni", rho=0)
pairedfnone<-pairwise_survdiff(Surv(Time,Dead)~Genotype,
data=survivaldatafnone, p.adjust.method = "bonferroni", rho=0)
#write pairwise comparisons to a dataframe
pairedmRUdf<-data.frame(pairedmRU$p.value)
pairedmnoRUdf<-data.frame(pairedmnoRU$p.value)
pairedmnonedf<-data.frame(pairedmnone$p.value)
```

```
pairedfRUdf<-data.frame(pairedfRU$p.value)
pairedfnoRUdf<-data.frame(pairedfnoRU$p.value)
pairedfnonedf<-data.frame(pairedfnone$p.value)
#write table of p values to a workbook
wb<-createWorkbook()
addWorksheet(wb, "Males +RU486")
addWorksheet(wb, "Males -RU486")
addWorksheet(wb, "Males none")
addWorksheet(wb, "Females +RU486")
addWorksheet(wb, "Females -RU486")
addWorksheet(wb, "Females none")
writeData(wb,"Males +RU486",pairedmRU$p.value,colNames = TRUE,rowNames =
TRUE)
writeData(wb,"Males -RU486",pairedmnoRU$p.value,colNames = TRUE,rowNames =
TRUE)
writeData(wb, "Males none",pairedmnone$p.value,colNames = TRUE,rowNames =
TRUE)
writeData(wb,"Females +RU486",pairedfRU$p.value,colNames = TRUE,rowNames =
TRUE)
writeData(wb,"Females -RU486",pairedfnoRU$p.value,colNames = TRUE,rowNames =
TRUE)
writeData(wb, "Females none",pairedfnone$p.value,colNames = TRUE,rowNames =
TRUE)
saveWorkbook(wb, "pvalues.xlsx")

#make a colour palette to use in plots
mycols<-colors()[c(132,402,200,230,133,404,125,124)]

#subset and bind together the data you intend to plot together- will vary
#refer to genotype names in genotype.table to select which to compare
names(genotype.table)
#for testing effect of copy number MHC
copiesmRU<-rbind(subset(survivaldatamRU, Genotype==names(genotype.table)[5]),
subset(survivaldatamRU, Genotype==names(genotype.table)
[6]),subset(survivaldatamRU, Genotype==names(genotype.table)
[9]),subset(survivaldatamRU, Genotype==names(genotype.table)[10]))
copiesmnoRU<-rbind(subset(survivaldatamnoRU, Genotype==names(genotype.table)
[5]), subset(survivaldatamnoRU, Genotype==names(genotype.table)
[6]),subset(survivaldatamnoRU, Genotype==names(genotype.table)
[9]),subset(survivaldatamnoRU, Genotype==names(genotype.table)[10]))
copiesfRU<-rbind(subset(survivaldatafRU, Genotype==names(genotype.table)[5]),
subset(survivaldatafRU, Genotype==names(genotype.table)
[6]),subset(survivaldatafRU, Genotype==names(genotype.table)
[9]),subset(survivaldatafRU, Genotype==names(genotype.table)[10]))
copiesfnoRU<-rbind(subset(survivaldatafnoRU, Genotype==names(genotype.table)
[5]), subset(survivaldatafnoRU, Genotype==names(genotype.table)
[6]),subset(survivaldatafnoRU, Genotype==names(genotype.table)
[9]),subset(survivaldatafnoRU, Genotype==names(genotype.table)[10]))
#for testing effect of edtp MHC
jumpymRU<-rbind(subset(survivaldatamRU, Genotype==names(genotype.table)
[4]),subset(survivaldatamRU, Genotype==names(genotype.table)
[10]),subset(survivaldatamRU, Genotype==names(genotype.table)
[11]),subset(survivaldatamRU, Genotype==names(genotype.table)
[15]),subset(survivaldatamRU, Genotype==names(genotype.table)[16]))
```

```
jumpymnoRU<-rbind(subset(survivaldatamnoRU, Genotype==names(genotype.table)
[4]),subset(survivaldatamnoRU, Genotype==names(genotype.table)
[10]),subset(survivaldatamnoRU, Genotype==names(genotype.table)
[11]),subset(survivaldatamnoRU, Genotype==names(genotype.table)
[15]),subset(survivaldatamnoRU, Genotype==names(genotype.table)[16]))
jumpyfRU<-rbind(subset(survivaldatafRU, Genotype==names(genotype.table)
[4]),subset(survivaldatafRU, Genotype==names(genotype.table)
[10]),subset(survivaldatafRU, Genotype==names(genotype.table)
[11]),subset(survivaldatafRU, Genotype==names(genotype.table)
[15]),subset(survivaldatafRU, Genotype==names(genotype.table)[16]))
jumpyfnoRU<-rbind(subset(survivaldatafnoRU, Genotype==names(genotype.table)
[4]),subset(survivaldatafnoRU, Genotype==names(genotype.table)
[10]),subset(survivaldatafnoRU, Genotype==names(genotype.table)
[11]),subset(survivaldatafnoRU, Genotype==names(genotype.table)
[15]),subset(survivaldatafnoRU, Genotype==names(genotype.table)[16]))
#for testing effect of copy number DJ694
copiesmnone<-rbind(subset(survivaldatamnone, Genotype==names(genotype.table)
[7]),subset(survivaldatamnone, Genotype==names(genotype.table)
[8]),subset(survivaldatamnone, Genotype==names(genotype.table)
[12]),subset(survivaldatamnone, Genotype==names(genotype.table)[14]))
copiesfnone<-rbind(subset(survivaldatafnone, Genotype==names(genotype.table)
[7]),subset(survivaldatafnone, Genotype==names(genotype.table)
[8]),subset(survivaldatafnone, Genotype==names(genotype.table)
[12]),subset(survivaldatafnone, Genotype==names(genotype.table)[14]))
#for testing effect of edtp DJ694
jumpymnone<-rbind(subset(survivaldatamnone, Genotype==names(genotype.table)
[1]),subset(survivaldatamnone, Genotype==names(genotype.table)
[2]),subset(survivaldatamnone, Genotype==names(genotype.table)
[3]),subset(survivaldatamnone, Genotype==names(genotype.table)
[12]),subset(survivaldatamnone, Genotype==names(genotype.table)
[13]),subset(survivaldatamnone, Genotype==names(genotype.table)[15]))
jumpyfnone<-rbind(subset(survivaldatafnone, Genotype==names(genotype.table)
[1]),subset(survivaldatafnone, Genotype==names(genotype.table)
[2]),subset(survivaldatafnone, Genotype==names(genotype.table)
[3]),subset(survivaldatafnone, Genotype==names(genotype.table)
[12]),subset(survivaldatafnone, Genotype==names(genotype.table)
[13]),subset(survivaldatafnone, Genotype==names(genotype.table)[15]))
#fit all the subsets
copiesmRUfit<-survfit(Surv(Time,Dead)~Genotype, data=copiesmRU)
copiesmnoRUfit<-survfit(Surv(Time,Dead)~Genotype, data=copiesmnoRU)
copiesfRUfit<-survfit(Surv(Time,Dead)~Genotype, data=copiesfRU)
copiesfnoRUfit<-survfit(Surv(Time,Dead)~Genotype, data=copiesfnoRU)

jumpymRUfit<-survfit(Surv(Time,Dead)~Genotype, data=jumpymRU)
jumpymnoRUfit<-survfit(Surv(Time,Dead)~Genotype, data=jumpymnoRU)
jumpyfRUfit<-survfit(Surv(Time,Dead)~Genotype, data=jumpyfRU)
jumpyfnoRUfit<-survfit(Surv(Time,Dead)~Genotype, data=jumpyfnoRU)

copiesmnonefit<-survfit(Surv(Time,Dead)~Genotype, data=copiesmnone)
copiesfnonefit<-survfit(Surv(Time,Dead)~Genotype, data=copiesfnone)

jumpymnonefit<-survfit(Surv(Time,Dead)~Genotype, data=jumpymnone)
jumpyfnonefit<-survfit(Surv(Time,Dead)~Genotype, data=jumpyfnone)
#plot all fits
```

```
copiesmRUplot<-ggsurvplot(copiesmRUfit, data=copiesmRU, fun="pct",
conf.int=TRUE, legend.title="",palette=mycols)
copiesmnoRUplot<-ggsurvplot(copiesmnoRUfit, data=copiesmnoRU, fun="pct",
conf.int=TRUE, legend.title="",palette=mycols)
copiesfRUplot<-ggsurvplot(copiesfRUfit, data=copiesfRU, fun="pct",
conf.int=TRUE, legend.title="",palette=mycols)
copiesfnoRUplot<-ggsurvplot(copiesfnoRUfit, data=copiesfnoRU, fun="pct",
conf.int=TRUE, legend.title="",palette=mycols)

jumpymRUplot<-ggsurvplot(jumpymRUfit, data=jumpymRU, fun="pct",conf.int =
TRUE, legend.title="",palette=mycols)
jumpymnoRUplot<-ggsurvplot(jumpymnoRUfit, data=jumpymnoRU, fun="pct",conf.int
= TRUE, legend.title="",palette=mycols)
jumpyfRUplot<-ggsurvplot(jumpyfRUfit, data=jumpyfRU, fun="pct",conf.int =
TRUE, legend.title="",palette=mycols)
jumpyfnoRUplot<-ggsurvplot(jumpyfnoRUfit, data=jumpyfnoRU, fun="pct",conf.int
= TRUE, legend.title="",palette=mycols)

copiesmnoneplot<-ggsurvplot(copiesmnonefit, data=copiesmnone,
fun="pct",conf.int = TRUE,legend.title="",palette = mycols)
copiesfnoneplot<-ggsurvplot(copiesfnonefit, data=copiesfnone,
fun="pct",conf.int = TRUE,legend.title="",palette = mycols)

jumpymnoneplot<-ggsurvplot(jumpymnonefit, data=jumpymnone, fun =
"pct",conf.int = TRUE, legend.title="", palette = mycols)
jumpyfnoneplot<-ggsurvplot(jumpyfnonefit, data=jumpyfnone, fun =
"pct",conf.int = TRUE, legend.title="", palette = mycols)

copiesmRUplot
copiesmnoRUplot
copiesfRUplot
copiesfnoRUplot
jumpymRUplot
jumpymnoRUplot
jumpyfRUplot
jumpyfnoRUplot
copiesmnoneplot
copiesfnoneplot
jumpymnoneplot
jumpyfnoneplot

pairedmRU
pairedmnoRU
pairedmnone
pairedfRU
pairedfnoRU
pairedfnone
```
